# Experimental evaluation of a direct fitness effect of the *de novo* evolved mouse gene *Pldi*

**DOI:** 10.1101/2024.01.13.575362

**Authors:** Miriam Linnenbrink, Gwenna Breton, Pallavi Misra, Christine Pfeifle, Julien Y. Dutheil, Diethard Tautz

## Abstract

*De novo* evolved genes emerge from random non-coding sequences and have, therefore, no homologs from which a function could be inferred. While expression analysis and knockout experiments can provide insights into the function, they do not directly test whether the gene is beneficial for its carrier. Here, we have used a seminatural environment experiment to test the fitness of the previously identified de *novo* evolved mouse gene *Pldi*, which is thought to be involved in sperm differentiation. We used a knockout mouse strain for this gene and competed it against its parental wildtype strain for several generations of free reproduction. We found that the knockout (ko) allele frequency decreased consistently across three replicates of the experiment. Using an approximate Bayesian computation framework that simulated the data under a demographic scenario mimicking the experiment’s demography, we could estimate a fitness coefficient ranging between 0.15 to 0.67 for the wildtype allele compared to the ko allele in males. We conclude that a gene that has evolved *de novo* from a random intergenic sequence can have a measurable fitness benefit.

## Introduction

Genes can evolve *de novo* from non-coding regions of the genome, representing more or less random DNA sequences (Tautz and Domazet-Loso, 2011; Van Oss and Carvunis, 2019; Cherezov et al., 2021). Unequivocal identification of such genes is only possible among very closely related species, where the history of mutations that have led to the emergence of the gene can be traced. Such de novo evolved genes have no homolog elsewhere from which possible functional information could be inferred. While their genetic function can be studied through knockout approaches, e. g. in mice (Xie et al., 2019; Xie et al., 2020), their relative importance for the species in which they arose is much more difficult to assess. For older genes that occur in multiple lineages, one can use substitution rate comparisons between coding and non-coding positions to determine the degree of selection acting on the gene. However, this is not possible in the very young *de novo* evolved genes since they will usually not have acquired enough new substitutions to allow such ratio estimates. Furthermore, it is often unclear whether young genes already encode a protein or whether the expressed RNA carries the gene function (Ruiz-Orera et al., 2020).

*Pldi* was one of the first fully described de *novo* genes, including the identification of possibly enabling mutations and functional information from a knockout study (Heinen et al., 2009). The transcript emerged a few million years ago as a testis-specific transcript in the lineage leading to the house mouse, apparently triggered through an indel mutation in the upstream region that led to a new testis-specific expression (Heinen et al., 2009). The *Pldi* transcript includes three exons derived from previously existing cryptic splice sites and two potential ORFs but with no direct evidence for translation of these ORFs. Knockout animals that lacked the second and the third exon showed lowered testis weights and reduced numbers of rapidly moving sperm (21.6% vs. 29%). Testis transcriptome comparisons revealed several differentially expressed regulatory genes. Among the top down-regulated genes were *Hmgb2* and *Arid4b. Hmgb2* causes seminiferous tubule atrophy via aberrant expression of androgen and estrogen receptors when depleted in mouse testis (Sugita et al., 2021). *Arid4b* is a chromatin remodeling gene; mice with haploinsufficiency of this gene show, in conjunction with a *Arid4a* ko, a loss of male fertility with a spermatogenic arrest at the stages of meiotic spermatocytes and postmeiotic haploid spermatids (Wu et al., 2013) (Heinen et al., 2009). This implies that Peg13 is involved in pathways that regulate spermatogenesis and fertility.

Here, we ask whether the subtle phenotype, especially the differences in sperm mobility in the range of individual variation between wildtype animals, could have a measurable fitness effect. Therefore, we ran a seminatural environment experiment, where knockout and wildtype mice lived together and could reproduce freely for several overlapping generations. We found reproducible shifts in allele frequencies with an overall drop of the knockout allele between the start and the end of the experiment. By simulating allele evolution under the exact demography of the experiment and under the premise that only males are affected, we infer that the fitness difference of the knockout allele is in the order of 0.60 (95% posterior interval: [0.05, 1.33]). We conclude that a de *novo* evolved transcript can have a direct fitness effect on the species in which it evolved.

## Methods

### Permissions

The seminatural environment rooms were run in 2012 and 2013. Since there was no experimental interference with the animals, only a permit for keeping the mice was required. This was obtained from the local veterinary office “Veterinäramt Kreis Plön” (permit number: 1401-153). The Government of Schleswig-Holstein provided permission to sacrifice animals under permit number V312-72241.123-34.

### Mouse strains

The *Pldi* knockout construct is described in (Heinen et al., 2009). It was generated in the C57Bl6/J inbred strain, and it is fully viable as a homozygous knockout. C57Bl6/J served as the competing wildtype (WT) strain in the experiment. Hence the mice used are expected to differ only for the *Pldi* allele. However, while it is well known that even inbred strains are not isogenic due to a continuous accumulation of new mutations (Chebib et al., 2021), the design of the experiment included mixing of the founder stocks. Hence, possible background differences between them are not expected to influence the conclusion on the change of the *Pldi* allele frequency after several generations of mixing.

### Experimental setup

The experiment was run in three replicate rooms, each for approximately 12 months, during which multiple overlapping generations formed. Rooms 1 and 2 were started in parallel in mid-November, and room 3 was started three months later. The experiments were initiated with equal numbers (n=10) of *Pldi* homozygous knockout and WT mice in an equal sex ratio at a density of about one mouse/m^2^ . All founder animals, as well as subsequently sampled live animals, were individually tagged. Water and food (Altromin 1324) were supplied ad libitum. The light/dark cycle was 12/12h. The temperature was kept constant at around 20°C; humidity was between 45 and 50%. The enclosures were equipped with bedding (Rettenmaier FS14), straw, paper, and housing. Divider walls and plastic tubes provided structural variation (see suppl File 1).

### Daily check and regular monitoring

The rooms were checked every day, and dead mice (mostly newborn or very young animals) were removed. Straw and paper were supplemented if necessary. Every 4-8 weeks, complete monitoring was performed. All mice were caught and checked for individual conditions (weight, pregnancy, marks of bites). Tissue samples (ear clips) were taken, and each adult mouse was marked with a tattoo on the tail or the ear. Mice of older generations were removed at defined times (see Figure 1). Samples for genotyping also included the dead animals found during the daily monitoring, as well as embryos from dead mothers, especially at the end of the experiment.

**Figure 1:**
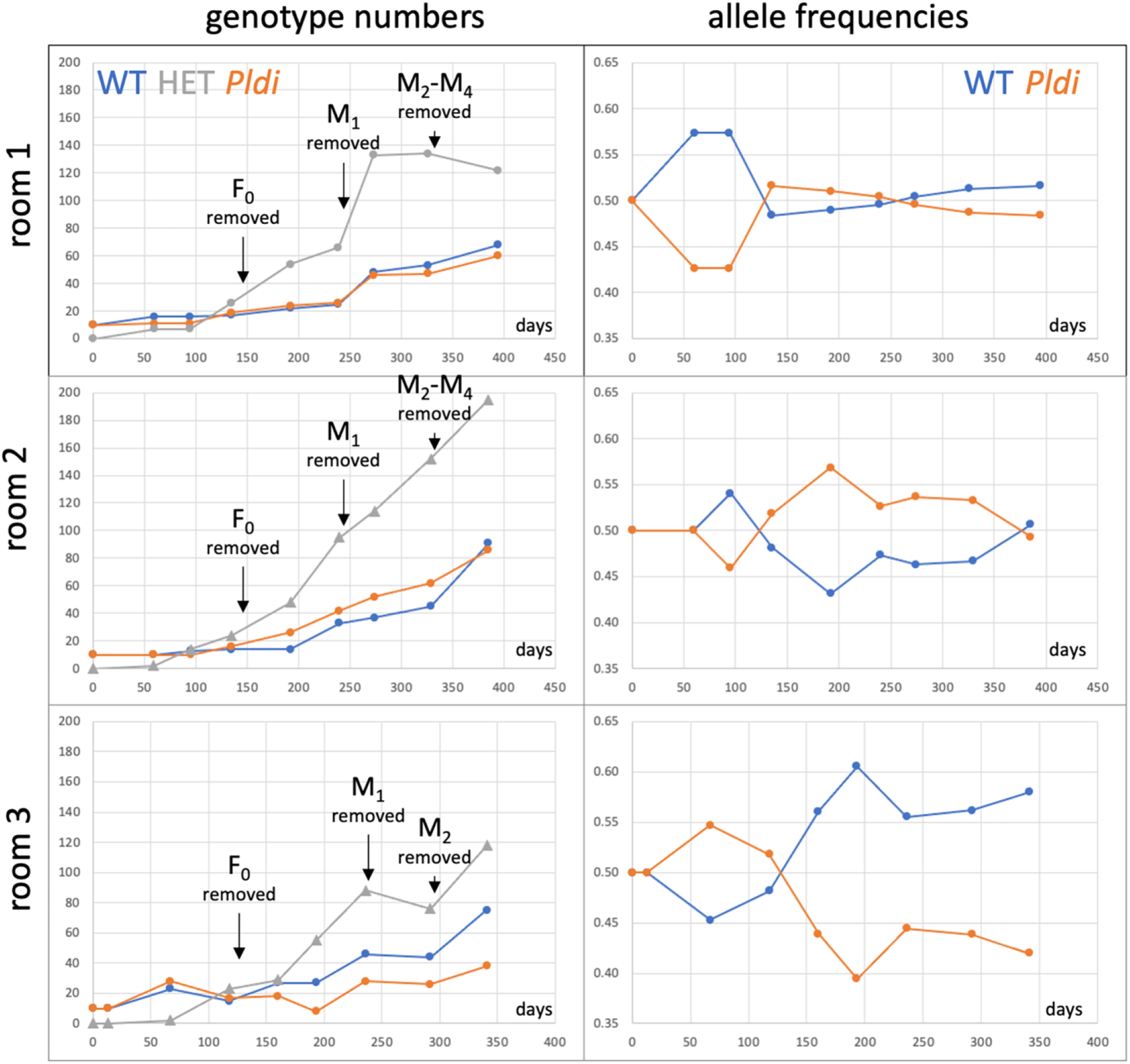
Development of animal numbers and allele frequencies during the experiment. Only the adult live animals present at the respective monitoring time point are depicted. Older animals were removed at the times indicated in the figure to mimic a more natural population growth, where older animals are frequently lost due to predation. The founder animals (F_0_) were removed at monitoring point 3 in each room. Animals first marked at later monitoring time points (M_1_-M_4_) were removed as indicated in the figure. The data for this figure are provided in suppl. Table S2.

### DNA extraction and typing

DNA from tissue samples was extracted with DNeasy 96 Blood & Tissue Kit (QIAGEN) following the “Purification of Total DNA from Animal Tissues” protocol. The amount of DNA was measured using a Nanodrop 1000 Spectrophotometer, and the samples were diluted to 5ng/μL. Genotyping was done with genotype-specific PCR primers (LOHfw: 5`-TGCCAAATCACCCTGCTTGC-3`; LOHrev: 5`-TGTGCAAGCTGTAACCATCC-3`; ROHfw: 5`-AGCCATAGCCTTGTCCAGAG-3`; ROHrev: 5`-CAGCTGCTTCTATTGGAAAGG-3`) to classify each sample into WT (LOHfw+LOHrev =351bp and ROHfw+ ROHrev=326bp), heterozygous or homozygous knockout (LOHfw+ ROHrev=420bp). The PCR was performed using the standard protocol of the QIAGEN Multiplex PCR Kit. The PCR products were loaded with the 6X DNA loading dye of Fermentas and run on a 1% agarose gel with the FastRuler DNA Ladder, Low Range, from Fermentas.

### Parameter estimation using Approximate Bayesian Computation

To assess the selective coefficient of the *Pldi* mutation, we considered a model that accounts for the exact population size and sex ratio for each monitoring point in each population. The fitness of the homozygote for the *Pldi* knockout, which mimics the ancestral genotype, was set to 1. The wildtype male homozygote containing two copies of *Pldi* was set to 1 + s, where s is the selection coefficient. Heterozygotes, which have one copy of the knockout and one copy of the wildtype, have a fitness of 1 + hs, where h is the dominance coefficient (h in [0, 1]). Individuals from a new generation were created by picking one female and one male parent, sampled with a probability proportional to their relative fitness. We first considered a model where *Pldi* is expressed in males only (that is, the fitness of females is equal to 1, independently of their genotypes).

We used two sets of observed statistics to infer values for the model parameters using approximate Bayesian computation (ABC). The models were fitted on either the set of frequencies of the two alleles or the three possible genotypes in the three populations at all relevant time points. We then simulated 100,000 replicates of the three populations, using a displaced Gamma prior for s and a uniform prior for h. More precisely, for each simulation replicate of each population, a value for s was taken randomly from a Gamma distribution with both shape and scale parameters equal to 3. The resulting value was then subtracted by one so that the distribution effectively comprises -1 and +infinity, with a mean value of 0. Similarly, the h coefficient was sampled randomly from a uniform distribution between 0 and 1. These simulations were then used to compute the posterior distributions of these parameters, using ridge regression with correction for heteroscedasticity, as implemented in the ‘abc’ package for R (Csillery et al., 2012). Posterior model probabilities were conducted using a multinomial logistic regression.

A cross-validation procedure was then performed by splitting the simulated datasets 1,000 times between validation and training sets. One simulation was randomly selected as validation data (leave-one-out procedure); the other 99,999 simulations were used as a training set to infer the corresponding values of s and h from the pseudo-data. The inferred values were then compared to the actual values used in the simulations, allowing prediction errors to be computed (Csillery et al., 2012). Three tolerance parameters, 0.01, 0.1, and 0.2, were tested, leading to highly similar prediction errors. A tolerance of 0.1 (10% of the closest points to the target values were kept) was then used for all analyses.

Model misclassification errors were calculated by simulating an extra 100,000 replicates using the posterior estimate for s (selection model Mod1), which were then compared to 100,000 replicates where s and h were set equal to 0 (neutral model Mod0). Because h could not be reliably estimated, it was also sampled from a uniform distribution in the Mod1 model. One replicate was then selected randomly, and each model’s posterior probabilities were computed using the remaining simulations (leave-one-out procedure, repeated 1,000 times). A confusion matrix was calculated by counting the proportion of cases out of 1,000 when the model with the highest posterior probability was distinct from the actual model used to generate the pseudo-data. A detailed description for estimating the selection coefficient is provided in Suppl. File 2.

Following a similar protocol, we contrasted the estimated values of s under distinct models. In the first one, the *Pldi* allele was allowed to have a fitness effect in both males and females so that the fitness of the individuals was independent of their sex (see Suppl. File 3 for details). We also tested a model where the *Pldi* allele was allowed to have a distinct fitness effect in both males and females (Suppl. File 4). We then considered a class of models where non-genetic variance was added. The fitness of an individual was randomly sampled from a normal distribution with mean 1 (homozygous male for the deletion or female), 1 + s (homozygous male for the *Pldi* allele, or 1 + h.s (heterozygous male). The standard deviation of the distribution was set to a parameter, g, which is independent of the individual’s genotype and sex. The prior of g was assigned a uniform distribution between 0 and 0.5. Because of the additional parameter, 500,000 simulations were generated and used in the ABC estimation. As g could not be reliably estimated from the data, it was integrated over its prior distribution when comparing models (Suppl. File 5). Finally, we considered a class of model with non-random mating by adding a parameter *λ* (*λ* in [0,1]) so that parents are chosen according to their fitness multiplied by (1 – *λ*) if they have identical alleles, (1 – *λ*/2) if they have only one allele in common, or 1 if both alleles are different (Suppl. Files 6-8). This model was fitted on genotype frequencies only.

The code for simulations and model fitting is available under https://gitlab.gwdg.de/molsysevol/poldi

## Results

The overall analysis across all genotyped individuals in the rooms is summarized in Table 1 (full data in suppl. Table S1). While almost double as many animals were generated in rooms 1 and 2 compared to room 3, the overall allele frequency changes are congruent. Genotypes of the offspring are compatible with Hardy-Weinberg expectations in all three rooms. The cumulative WT allele frequency (including the dead and removed animals) increased from 0.5 to 0.52-0.6 in all three rooms, implying an overall advantage of the WT over the *Pldi* ko allele.

**Table 1:** Cumulative analysis of all genotyped samples.

|  | total (N) | WT* | HET | *Pldi* ko* | Chi<sup>2</sup># | f<sub>WT</sub> | f<sub>*Pldi* ko</sub> |
| --- | --- | --- | --- | --- | --- | --- | --- |
| Room 1 | 1181 | 338 | 576 | 267 | 0.52 | 0.53 | 0.47 |
| Room 2 | 1205 | 329 | 584 | 292 | 1.07 | 0.52 | 0.48 |
| Room 3 | 578 | 216 | 264 | 98 | 1.32 | 0.6 | 0.4 |

\*founder animals not included
#deviation from Hardy-Weinberg expectations – all non-significant at $p > 0.05$ (1 d.f.)

We also created a sub-data set tracing all adults and alive animals at each monitoring time point, allowing us to visualize the population growth and shift in allele frequency (Figure 1; full data in suppl. Table S2). The first three time points represent primarily the founder animals, which were removed at monitoring time point 4. Note that the population sizes are initially very small and biased through the presence of the founder animals, leading to increased stochasticity with respect to allele frequency fluctuations. At the last sampling time, homozygous WT animals exceed the number of homozygous *Pldi* animals in all rooms.

### Estimation of fitness effect

We used an approximate Bayesian computation framework to estimate the selection coefficient of the *Pldi* allele. We simulated data under a demographic scenario mimicking the demography of the experiment (population size at each generation, sex ratio, and life expectancy), considering the actual numbers of adult animals from the three replicate populations as they are depicted in Figure 1. We conducted estimation procedures with either the frequencies of the alleles or of the three genotypes in the three populations as summary statistics and estimated two parameters, following Wright’s notations: s, the difference of fitness between the WT homozygote (fitness 1 + s) and the “ancestral” genotype (*Pldi* knockout homozygote, fitness of 1), and h, the dominance coefficient so that heterozygotes have a fitness of 1 + hs. Because the *Pldi* gene is expected to have an effect only in males, the h and s parameters were set to 0 in females, so that all genotypes have a fitness of 1 in females.

Because the founding animals were only homozygotes, they were not at Hardy-Weinberg equilibrium and could have been influenced by strain background (see Methods). Hence, we excluded the first two time points and started the modeling from the third time point, using the corresponding observed frequencies as initial conditions. The posterior distribution of h was flat, indicating that the dominance parameter could not be reliably estimated. Conversely, the posterior distribution of the selection coefficient, s, was different from the prior and notably shifted toward positive values. The posterior average was estimated to be 0.595, with a 95% posterior interval of [0.054, 1.326] using the observed allelic frequencies (Figure 2A and Table 2), and 0.343, with a 95% posterior interval of [0.015, 0.788] using the observed genotype frequencies (Table 2). We then conducted a series of posterior predictive tests, as recommended in (Csillery et al., 2012). We first conducted a cross-validation experiment to assess the accuracy of the parameter estimation. A leave-one-out procedure was used, where one simulation was selected randomly and the corresponding summary statistics were used as pseudo-observations. The other simulations were then used to estimate the underlying parameters, which were compared to their true values used in the simulation (Figure 2B). The results show that the s parameter could be estimated reliably (prediction error of 0.14 when allele frequencies are used, 0.17 for genotype frequencies), while the h parameters could not (prediction error of 0.94 and 0.95, for allele and genotype frequencies, respectively), concurring with the observed posterior distributions. The posterior model probability for the selection model was calculated to be 95% in both cases (Figure 2C and Table 2). Classification errors were assessed by a leave-one-out procedure, comparing simulations generated under a purely neutral model and a selection model where the selection coefficient in males, s, was set to the value inferred from the data, while h was sampled from a uniform distribution. One simulation was then selected randomly and used as pseudo-observed data and posterior probabilities of each model were computed using the remaining simulations. One thousand random selections were performed for each model, allowing the computation of the so-called confusion matrix, which contains the frequency of cases where the incorrect model had a higher posterior probability than the true one. This cross-validation test shows that the model with selection is distinguishable from the neutral model (Figure 2D).

**Table 2:**
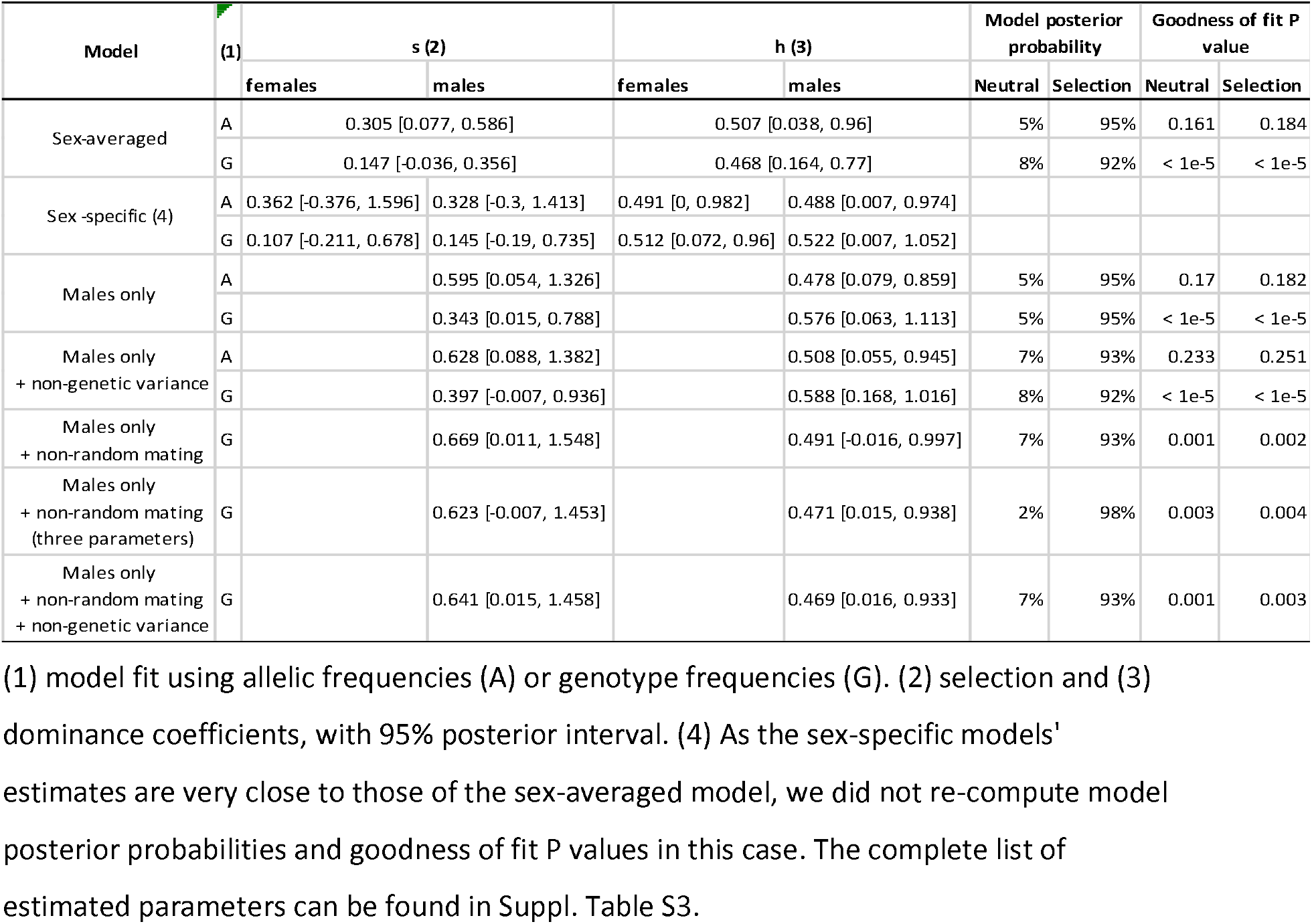
Estimation of selection coefficient under distinct models.

**Figure 2:**
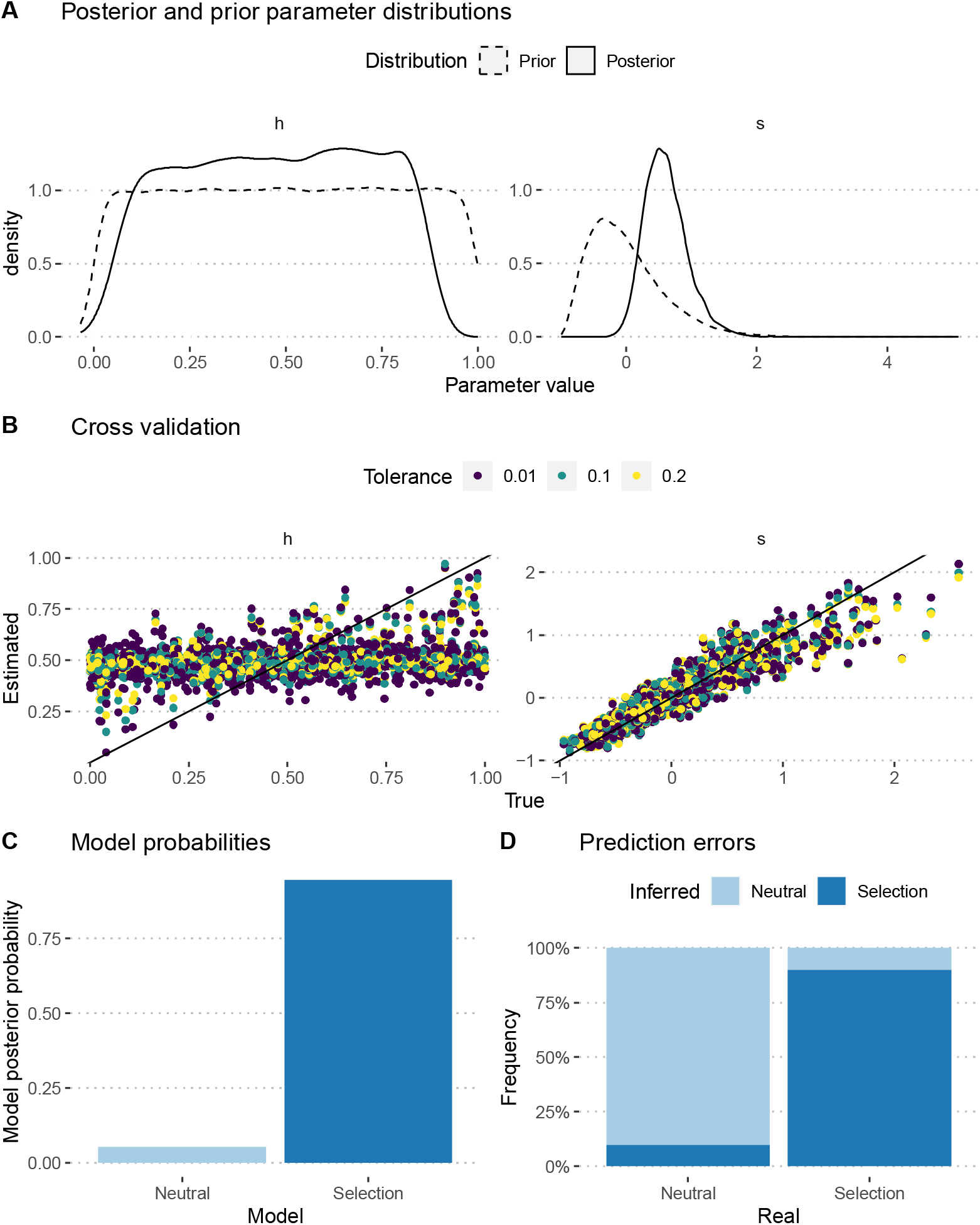
Estimation of the *Pldi* fitness effect by Approximate Bayesian Computation using observed allelic frequencies. A: posterior distributions of the h and s parameters, compared with their prior. B: cross-validation analysis showing the estimation accuracy of the two parameters. C: Posterior model probabilities of the data. D: confusion matrix of the cross-validation experiment, based on 1,000 sampled simulations. The x-axis indicates the model used for simulations. The y-axis is the proportion of cases where each model was the best fit based on their posterior model probabilities. A similar figure for the estimation using genotype frequencies is provided in Suppl. File 9.

Using a sex-specific fitness model did not allow us to estimate distinct effect effects of *Pldi* in males and females (Table 2). The estimated s values are roughly half that of the male-only model, yet with a large estimation variance, the 95% posterior interval including 0. A model where the effect of *Pldi* is independent of the sex leads to similar estimates to those of the sex-specific model. Yet, each of these models favors a model where *Pldi* has a positive effect with a probability higher than 90%, even when adding non-genetic variance (Table 2). In each of these models, however, we observe that the model fit to observed genotype frequencies was poor (goodness of fit P value < 1e-5, Table 2), which might be explained by a departure from the random mating hypothesis. We further fitted models relaxing this hypothesis, which only marginally improved the fit. The estimated selection coefficients were very similar when non-random mating was incorporated, and the posterior probability of the model with selection was systematically above 90% (Table 2).

While these analyses did not allow for the estimation of the dominance coefficient, they point to a beneficial effect of the *Pldi* WT allele. The observed increases in allele frequencies in the three populations of the experiment are better described by a model with positive selection than by a purely neutral model.

## Discussion

Our experimental setup was geared towards testing whether a knockout allele of the *de novo* evolved gene *Pldi* in house mice would show a fitness effect under seminatural conditions of reproduction over several overlapping generations. Given that *Pldi* is a *de novo* evolved gene, it is the appropriate comparison that mimics the time point at which it has evolved, i.e., a comparison between populations that have the new gene versus populations that do not have it. *Pldi* is expressed in the testis and affects sperm mobility; hence, it could directly affect the allele transmission into the next generation via a fitness effect in males. For all three replicate rooms in which the experiment was conducted, we found a reduction of the knockout allele after several generations of free mating with overlapping generations. Explicit modeling of the demographic situation in the rooms allowed us to infer a selection coefficient ranging from 0.15 to 0.67 for homozygote WT males across models, and posterior predictive analyses consistently strongly favored the model with selection in all tested models. These results imply that the knockout actually has a relatively strong negative fitness effect on its carriers. Whether one can also conclude the converse, i.e., that the fitness advantage of the newly evolved gene would have been in the same order is open since the emergence of the gene happened several million years ago in an unknown genetic background. Still, our experimental analysis implies that the fitness effects of *de novo* evolved genes can be within evolutionarily relevant dimensions. In fact, natural selection coefficients associated with strong selective sweeps are even much smaller (in the order of s=0.01) in natural mouse populations (Teschke et al., 2008).

Our modeling did not allow us to estimate the dominance coefficient and could not establish whether *Pldi* had an exclusive fitness effect in males. Relaxing the hypothesis of a male-specific effect resulted in an estimated selection coefficient half of that in models with a males-only fitness effect but did not significantly alter the support for the scenario where the *Pldi* allele is positively selected. While allelic frequency trajectories are well-predicted under our model (Suppl. File 9), this is not the case of genotype frequencies, where an excess of heterozygotes is observed in some rooms (suppl Files 2-9). This excess may stem from a deviation from the random mating assumption, an effect that we could only partially capture by adding extra parameters controlling mating choice. These results suggest that taking into account the spatial organization of the populations may be needed to improve the model further.

Tests of fitness effects in seminatural enclosure experiments were previously reported for two mouse mutant alleles for genes involved in circadian timing, namely *Per2* (Daan et al., 2011) and *tau* (= Csnk1e: casein kinase 1, epsilon) (Spoelstra et al., 2016). Both were done in outdoor enclosures under natural weather and predator conditions. Both showed major frequency shifts of the mutant alleles. *Per2*’s overall frequency dropped from 0.54 to 0.41 in the first year but increased again in the second year, in conjunction with a shift in the sex ratio (Daan et al., 2011). Hence, the allele frequency changed in response to overall conditions but did not generate a general fitness disadvantage within the two years of the experiment. In the case of the *tau* experiment, the overall allele frequency dropped consistently from 0.5 to 0.2 within 14 months, implying a negative selection effect (Spoelstra et al., 2016). This drop in allele frequency is stronger than what we found for the *Pldi* ko allele in our experiments, implying that the effect of a loss of a *de novo* evolved gene is in a lower fitness range than the loss of an ancient conserved gene, such as tau.

*Pldi* has emerged from a previously non-coding intergenic region of the mouse genome, i.e., its sequence composition is nearly random. Sequence comparisons between the outgroups that do not express the gene and the ingroup species that express the gene are shown in Suppl. File 10. It is evident that the RNA-coding region has acquired only a few new mutations compared to the outgroups. At most, seven substitutions have occurred in the case of *M. m. domesticus* GER, which represents the C57Bl6/J sequence. Five of them may have been adaptive since they are shared with at least some of the other ingroup species. However, these are less than 1% of the total sequence, which is also within the range of neutral evolution for these species (Harr et al., 2016). Hence, the *Pldi* RNA still represents primarily the essentially random intergenic sequence from which it evolved, although some optimization through adaptive mutations may have occurred.

Initially, it was thought to be very unlikely that random sequences could have a genetic function (Jacob, 1977). However, this view has changed after the discovery of *de novo* gene evolution (Tautz, 2014). Direct experiments with the expression of random sequences have also shown that a substantial fraction of them can exert phenotypic and fitness effects (Bao et al., 2017; Neme et al., 2017; Castro and Tautz, 2021; Bhave and Tautz, 2021; Shuipys et al., 2019). Our results add to this insight by showing that a naturally evolved *de novo* gene has a direct fitness effect on its carriers.

## Supporting information

combined suppl files

## Acknowledgments

We thank Heike Harre for her invaluable help with conducting the experiment and Prof. Mathieu Emily for his advice on the ABC analysis. Many thanks also to Maren Volquardsen, who took care of the daily control of the mice and Cornelia Burghardt and Heinke Buhtz for their support in the lab. The work was financed through institutional funds of the MPG to DT.

