## Supplementary material for "Experimental evaluation of a direct fitness effect of the *de novo* evolved mouse gene *Pldi*": combined suppl files: SupplementaryFile1.pdf

**Supplementary Figure 1**

Setup of the seminatural environment enclosures

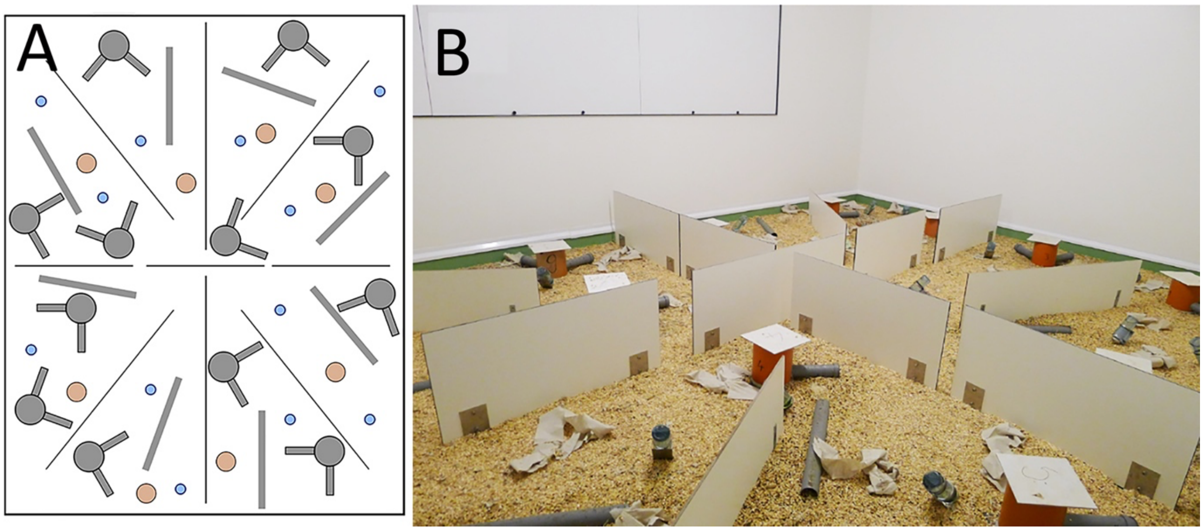


**A** Schematic representation of an enclosure. 4 quadrants were delimited with dividers. Each quadrant contained three houses with two tubes, three water bottles (blue circles), two feeding stations, and two additional tubes where mice could hide. **B** View into an enclosure room.
