## Supplementary material for "Experimental evaluation of a direct fitness effect of the *de novo* evolved mouse gene *Pldi*": combined suppl files: SupplementaryFile3.pdf

### Estimating poldi's selection coefficient

Julien Y. Dutheil

27/01/2023

### 1 Model parameterization

The model includes two parameters: $s$ the selection coefficient of the *poldi* allele compared to the knockout strain, which mimics the ancestal state, and $h$ the heterozygosity. Fitness values of each genotype are parameterized as follow:

$$\begin{aligned} poldi/poldi &: 1 \\ BL6/poldi &: 1 + h \cdot s \\ BL6/BL6 &: 1 + s \end{aligned}$$

where BL6 denotes the wild strain, and poldi the strain where the *poldi* gene is knocked out. Here, we assume that the fitness effect of an allele is independent of the sex of the individual.

We performed 100,000 simulations, using a gamma prior for $s$ and a uniform prior between 0 and 1 for $h$.

### 2 Preamble

We load the data (observed and simulated):

```
sims <- read.csv("Simulations/SimulationsFromGen3.csv.gz")
param.sim <- subset(sims, select = c("h", "s"))
```

```
stat.sim <- subset(sims, select = c(2:46))
# Only consider the WT homozygotes and the heterozygotes,
# as the sum of the three genotypes is 1
stat.sim <- subset(stat.sim, select = grep(".22.", names(stat.sim), invert = TRUE))

library(plyr, quietly = TRUE)
geno <- read.csv("GenotypeFrequencies.csv")
# We select the WT homozygote and heterozygote frequencies.
# Pldi homozygotes are 1 - sum(other two).
stat.obs <- subset(geno, TimePoint %in% 4:8, c("WT", "HET", "Replicate"))
stat.obs <- ddply(stat.obs, "Replicate", function(d) unlist(d[,1:2]))
stat.obs$Replicate <- NULL
stat.obs <- as.data.frame(t(stat.obs))
colnames(stat.obs) <- c("F1", "F2", "F3")
x <- rownames(stat.obs)
x <- gsub(x, pattern = "WT", replacement = ".11.")
x <- gsub(x, pattern = "HET", replacement = ".12.")
#x <- gsub(x, pattern = "Pldi", replacement = ".22.")
rownames(stat.obs) <- x
stat.obs1 <- subset(stat.obs, select = F1)
rownames(stat.obs1) <- paste0("F1", rownames(stat.obs1))
stat.obs2 <- subset(stat.obs, select = F2)
rownames(stat.obs2) <- paste0("F2", rownames(stat.obs2))
stat.obs3 <- subset(stat.obs, select = F3)
rownames(stat.obs3) <- paste0("F3", rownames(stat.obs3))
stat.obs <- cbind(t(stat.obs1), t(stat.obs2), t(stat.obs3))[1,]
```

### 3 Using allele frequencies

First, we need to compute the allelic frequencies from the genotype frequencies:

```
compute.allelic.frequencies <- function(stat.sim) {
  stat.sim.a <- data.frame(row.names = row.names(stat.sim))
  #Pop1
  stat.sim.a$F1.1 <- stat.sim$F1.11.1 + stat.sim$F1.12.1 / 2
  stat.sim.a$F1.2 <- stat.sim$F1.11.2 + stat.sim$F1.12.2 / 2
  stat.sim.a$F1.3 <- stat.sim$F1.11.3 + stat.sim$F1.12.3 / 2
  stat.sim.a$F1.4 <- stat.sim$F1.11.4 + stat.sim$F1.12.4 / 2
  stat.sim.a$F1.5 <- stat.sim$F1.11.5 + stat.sim$F1.12.5 / 2
  #Pop2
  stat.sim.a$F2.1 <- stat.sim$F2.11.1 + stat.sim$F2.12.1 / 2
  stat.sim.a$F2.2 <- stat.sim$F2.11.2 + stat.sim$F2.12.2 / 2
  stat.sim.a$F2.3 <- stat.sim$F2.11.3 + stat.sim$F2.12.3 / 2
  stat.sim.a$F2.4 <- stat.sim$F2.11.4 + stat.sim$F2.12.4 / 2
  stat.sim.a$F2.5 <- stat.sim$F2.11.5 + stat.sim$F2.12.5 / 2
  #Pop3
  stat.sim.a$F3.1 <- stat.sim$F3.11.1 + stat.sim$F3.12.1 / 2
  stat.sim.a$F3.2 <- stat.sim$F3.11.2 + stat.sim$F3.12.2 / 2
  stat.sim.a$F3.3 <- stat.sim$F3.11.3 + stat.sim$F3.12.3 / 2
  stat.sim.a$F3.4 <- stat.sim$F3.11.4 + stat.sim$F3.12.4 / 2
  stat.sim.a$F3.5 <- stat.sim$F3.11.5 + stat.sim$F3.12.5 / 2

  return(stat.sim.a)
```

```r
}
stat.sim.a <- compute.allelic.frequencies(stat.sim)
stat.obs.a <- numeric(15)
names(stat.obs.a) <- c("F1.1", "F1.2", "F1.3", "F1.4", "F1.5",
                       "F2.1", "F2.2", "F2.3", "F2.4", "F2.5",
                       "F3.1", "F3.2", "F3.3", "F3.4", "F3.5")
#Pop1
stat.obs.a[1]  <- stat.obs["F1.11.1"] + stat.obs["F1.12.1"] / 2
stat.obs.a[2]  <- stat.obs["F1.11.2"] + stat.obs["F1.12.2"] / 2
stat.obs.a[3]  <- stat.obs["F1.11.3"] + stat.obs["F1.12.3"] / 2
stat.obs.a[4]  <- stat.obs["F1.11.4"] + stat.obs["F1.12.4"] / 2
stat.obs.a[5]  <- stat.obs["F1.11.5"] + stat.obs["F1.12.5"] / 2
#Pop2
stat.obs.a[6]  <- stat.obs["F2.11.1"] + stat.obs["F2.12.1"] / 2
stat.obs.a[7]  <- stat.obs["F2.11.2"] + stat.obs["F2.12.2"] / 2
stat.obs.a[8]  <- stat.obs["F2.11.3"] + stat.obs["F2.12.3"] / 2
stat.obs.a[9]  <- stat.obs["F2.11.4"] + stat.obs["F2.12.4"] / 2
stat.obs.a[10] <- stat.obs["F2.11.5"] + stat.obs["F2.12.5"] / 2
#Pop3
stat.obs.a[11] <- stat.obs["F3.11.1"] + stat.obs["F3.12.1"] / 2
stat.obs.a[12] <- stat.obs["F3.11.2"] + stat.obs["F3.12.2"] / 2
stat.obs.a[13] <- stat.obs["F3.11.3"] + stat.obs["F3.12.3"] / 2
stat.obs.a[14] <- stat.obs["F3.11.4"] + stat.obs["F3.12.4"] / 2
stat.obs.a[15] <- stat.obs["F3.11.5"] + stat.obs["F3.12.5"] / 2
```

We estimate parameters using the ridge regression method:

```r
library(abc, quietly = TRUE)
```

```
##
## Attaching package: 'SparseM'

## The following object is masked from 'package:base':
##
##     backsolve

## locfit 1.5-9.8    2023-06-11
```

```r
if (redo) {
  m.r.a <- abc(target = stat.obs.a, param = param.sim, sumstat = stat.sim.a, tol=.1,
               method = "ridge", transf = c("none", "none"))
  save(m.r.a, file = "Rdata/SexAveraged/backup_abc_allelic.Rdata")
} else {
  load("Rdata/SexAveraged/backup_abc_allelic.Rdata")
}
```

Now we display the results:

```r
summary(m.r.a, intvl = .95)
```

```
## Call:
## abc(target = stat.obs.a, param = param.sim, sumstat = stat.sim.a,
##     tol = 0.1, method = "ridge", transf = c("none", "none"))
## Data:
##  abc.out$adj.values (10000 posterior samples)
## Weights:
##  abc.out$weights
```

```
##
##                                h       s
## Min.:                    -0.0904 -0.1345
## Weighted 2.5 % Perc.:     0.0376  0.0768
## Weighted Median:          0.5127  0.2968
## Weighted Mean:            0.5074  0.3051
## Weighted Mode:            0.6122  0.3072
## Weighted 97.5 % Perc.:    0.9604  0.5857
## Max.:                     1.0666  1.0572
```

Distribution of parameter estimates:

```
hist(m.r.a, breaks = 30, caption = c(expression(h), expression(s)))
```

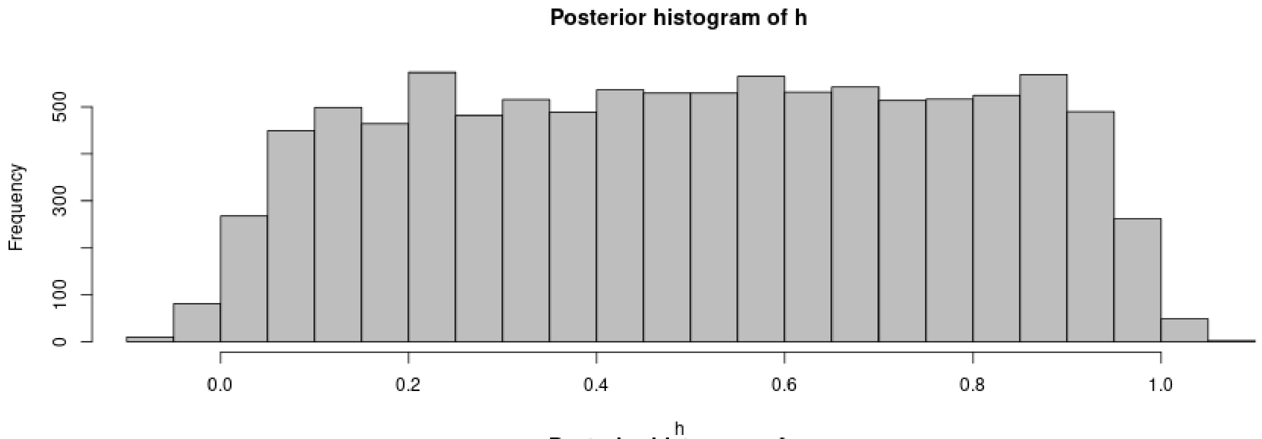

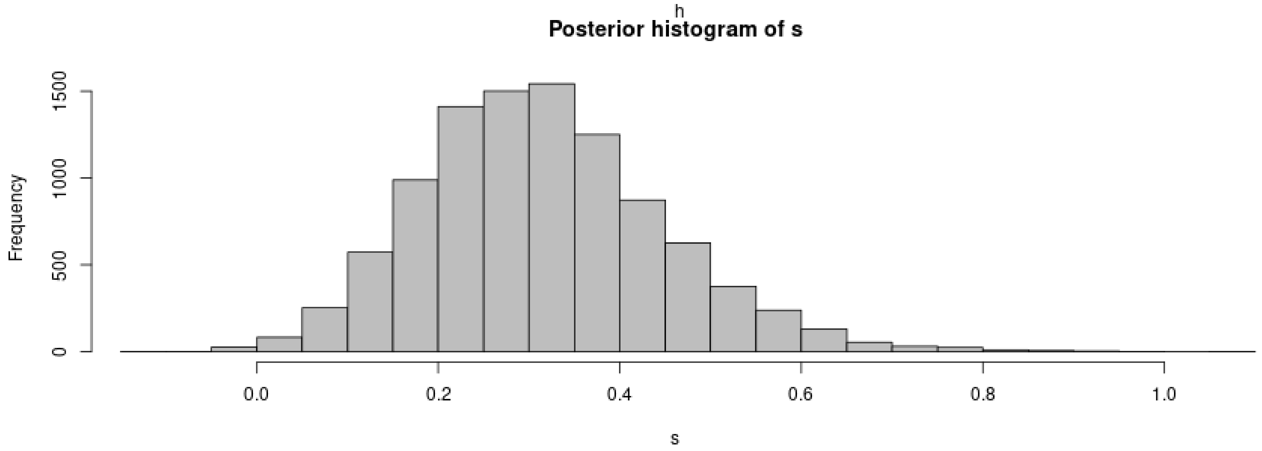

```
plot(m.r.a, param = param.sim)
```

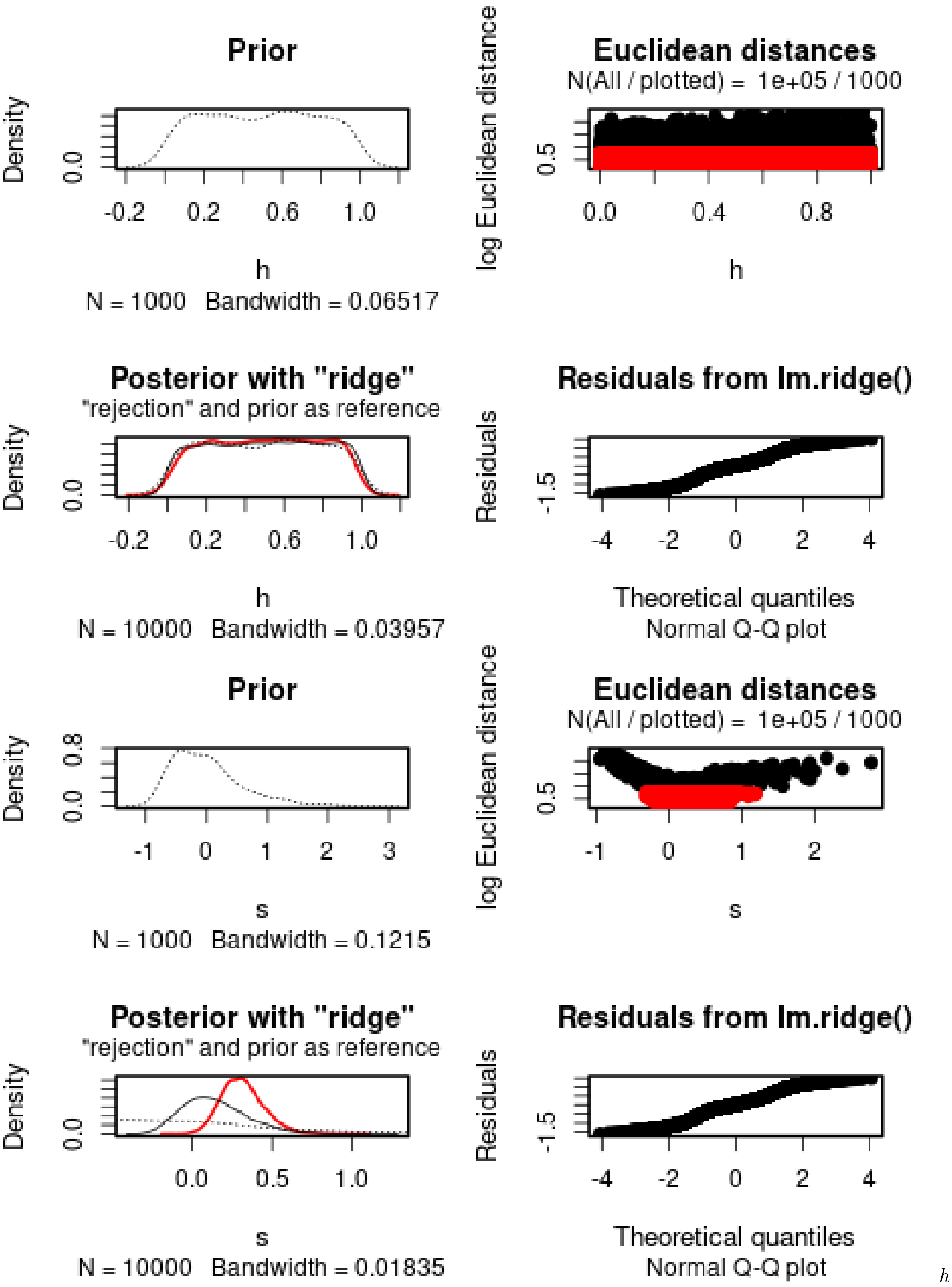

$h$ cannot properly be estimated. $s$ is positive, but has a large variance. The 95% posterior interval does not include 0.

We then check the distribution of summary statistics in the simulations:

```
library(reshape2)
library(ggplot2)
library(ggpubr)
```

```
##
## Attaching package: 'ggpubr'

## The following object is masked from 'package:plyr':
##
##     mutate
```

```
df.stats.sim <- melt(stat.sim.a, variable.name = "Statistic", value.name = "Frequency")
```

```
## No id variables; using all as measure variables
```

```
l <- strsplit(as.character(df.stats.sim$Statistic), split = ".", fixed = TRUE)
df.stats.sim$Room <- sapply(l, function(x) x[1])
df.stats.sim$Generation <- sapply(
  strsplit(as.character(df.stats.sim$Statistic), split = ".", fixed = TRUE),
  function(x) x[2])

df.stats.obs <- as.data.frame(stat.obs.a)
names(df.stats.obs) <- "Frequency"
df.stats.obs$Statistic <- row.names(df.stats.obs)
l <- strsplit(as.character(df.stats.obs$Statistic), split = ".", fixed = TRUE)
df.stats.obs$Room <- sapply(l, function(x) x[1])
df.stats.obs$Generation <- sapply(
  strsplit(as.character(df.stats.obs$Statistic), split = ".", fixed = TRUE),
  function(x) x[2])

p <- ggplot(data = df.stats.sim, aes(x = Frequency, y = after_stat(density))) +
      geom_histogram(bins = 50) +
      geom_vline(data = df.stats.obs, aes(xintercept = Frequency), color = "orange") +
      facet_grid(Room~Generation) +
      theme_pubclean()
p
```

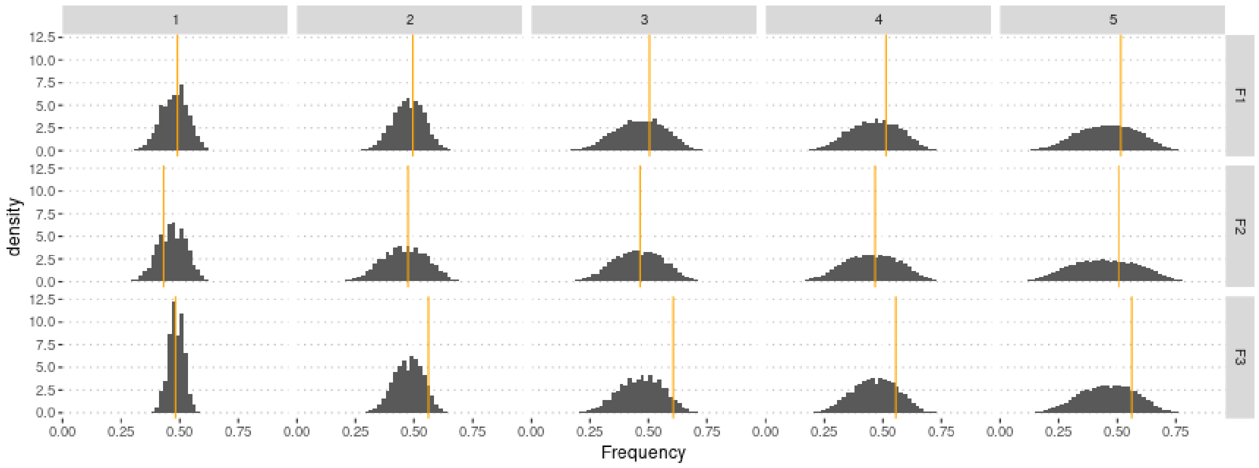

#### 3.1 Cross-Validation analysis

Compute predictions errors:

```
if (redo) {
  cv.ridge.a <- cv4abc(param = param.sim,
                       sumstat = stat.sim.a,
                       abc.out = m.r.a,
                       nval = 1000,
                       tols = c(.01,.1,.2))
  save(cv.ridge.a, file = "Rdata/SexAveraged/backup_cv_allelic.Rdata")
} else {
  load("Rdata/SexAveraged/backup_cv_allelic.Rdata")
}
```

```
summary(cv.ridge.a)
```

```
## Prediction error based on a cross-validation sample of 1000

##               h          s
## 0.01 0.90259525 0.07296150
## 0.1  0.89131439 0.07815060
## 0.2  0.89804001 0.08237657
```

```
plot(cv.ridge.a)
```

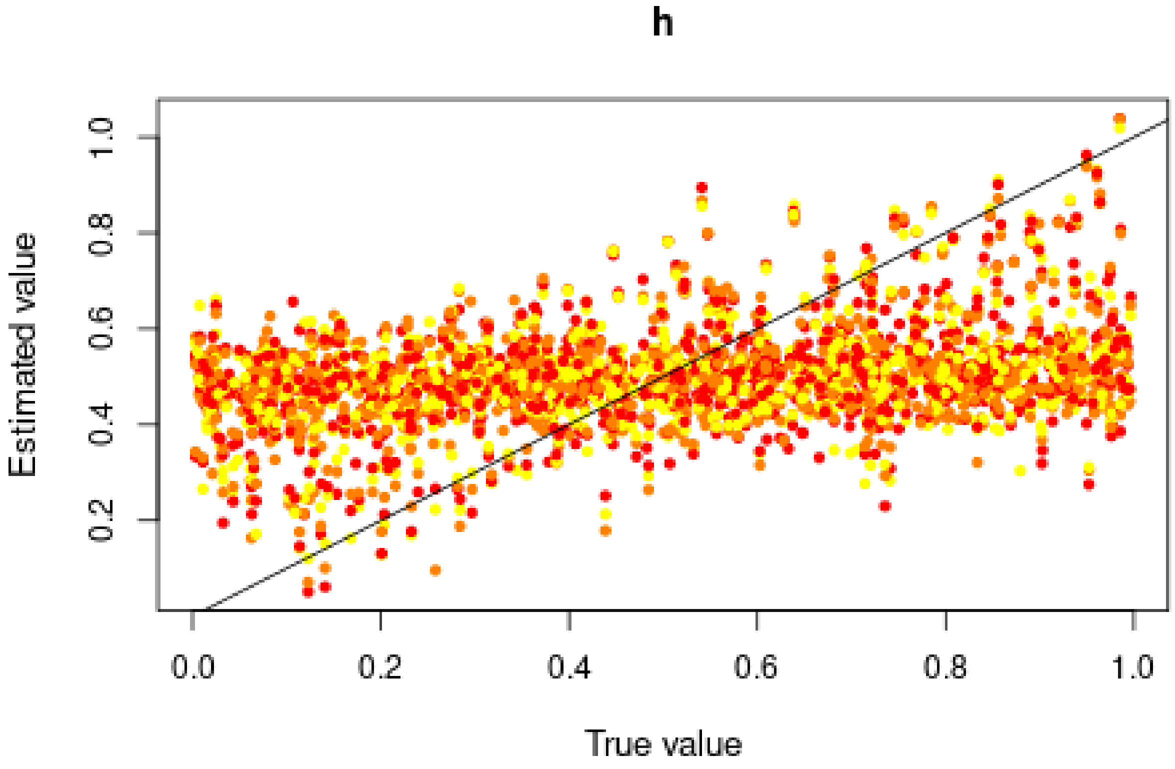

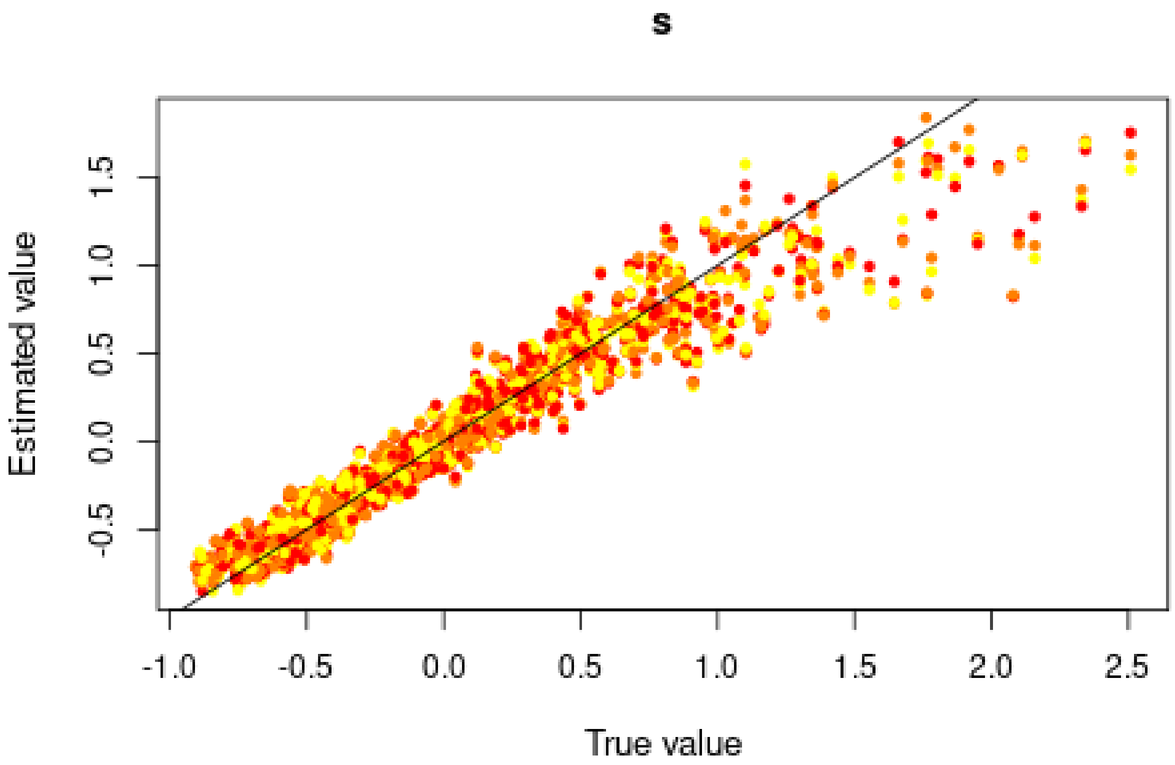

Good for $s$, but very bad for $h$, in agreement with the posterior distributions.

#### 3.2 Misclassification errors

To test the power of the approach to distinguish between models, we also conduct a cross-validation experiment. We compare model M0 ($s = 0$) and M1 ($s = 0.3051 > 0$). $h$ is sampled over its prior distribution. 100,000 simulations were conducted under each model.

```
sims0 <- read.csv("Simulations/Simulations0FromGen3.csv.gz")
param.sim0 <- subset(sims0, select = c("h", "s"))
stat.sim0 <- subset(sims0, select = 2:46)
stat.sim0.a <- compute.allelic.frequencies(stat.sim0)

sims1 <- read.csv("Simulations/Simulations1aFromGen3.csv.gz")
param.sim1 <- subset(sims1, select = c("h", "s"))
stat.sim1 <- subset(sims1, select = 2:46)
stat.sim1.a <- compute.allelic.frequencies(stat.sim1)
```

We conduct a CV analysis (this takes some time... set nval = 100 for faster results):

```
models <- rep(c("Neutral", "Selection"), each = 100000)
if (redo) {
  cv.modsel.a <- cv4postpr(models, rbind(stat.sim0.a, stat.sim1.a),
                           nval = 1000, tols = .1, method = "mnlogistic")
  save(cv.modsel.a, file = "Rdata/SexAveraged/backup_cv4postpr_allelic.Rdata")
} else {
  load("Rdata/SexAveraged/backup_cv4postpr_allelic.Rdata")
}
```

We display the results:

```
summary(cv.modsel.a)
```

```
## Confusion matrix based on 1000 samples for each model.
##
## $tol0.1
##           Neutral Selection
## Neutral       939        61
## Selection      65       935
##
##
## Mean model posterior probabilities (mnlogistic)
##
## $tol0.1
##           Neutral Selection
## Neutral    0.9112    0.0888
## Selection  0.0983    0.9017
```

```
plot(cv.modsel.a, names.arg = c("Neutral", "Selection"))
```

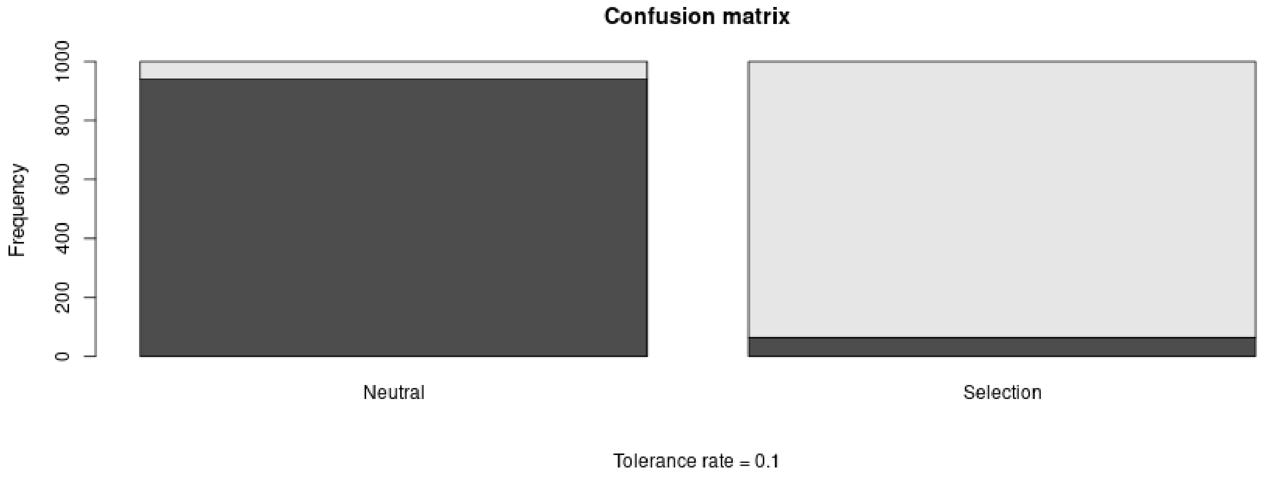

#### 3.3 Posterior prediction

```
if (redo) {
  modsel.a <- postpr(stat.obs.a, models, rbind(stat.sim0.a, stat.sim1.a),
                     tol = .1, method = "mnlogistic")
  save(modsel.a, file = "Rdata/SexAveraged/backup_postpr_allelic.Rdata")
} else {
  load("Rdata/SexAveraged/backup_postpr_allelic.Rdata")
}
summary(modsel.a)
```

```
## Call:
## postpr(target = stat.obs.a, index = models, sumstat = rbind(stat.sim0.a,
##     stat.sim1.a), tol = 0.1, method = "mnlogistic")
## Data:
##  postpr.out$values (20000 posterior samples)
## Models a priori:
##  Neutral, Selection
## Models a posteriori:
##  Neutral, Selection
##
## Proportion of accepted simulations (rejection):
```

```
##   Neutral Selection
##    0.4488    0.5512
##
## Bayes factors:
##           Neutral Selection
## Neutral    1.0000    0.8141
## Selection  1.2284    1.0000
##
##
## Posterior model probabilities (mnlogistic):
##   Neutral Selection
##    0.0533    0.9467
##
## Bayes factors:
##           Neutral Selection
## Neutral    1.0000    0.0563
## Selection 17.7476    1.0000
```

The model with selection is preferred.

##### 3.3.1 Goodness of fit

Under the neutral model:

```
if (redo) {
  res.gfit0.a <- gfit(target = stat.obs.a, sumstat = stat.sim0.a,
                      statistic = median, nb.replicate = 1000)
  save(res.gfit0.a, file = "Rdata/SexAveraged/backup_gfit0_allelic.Rdata")
} else {
  load("Rdata/SexAveraged/backup_gfit0_allelic.Rdata")
}
plot(res.gfit0.a, main = "Histogram under M0")
```

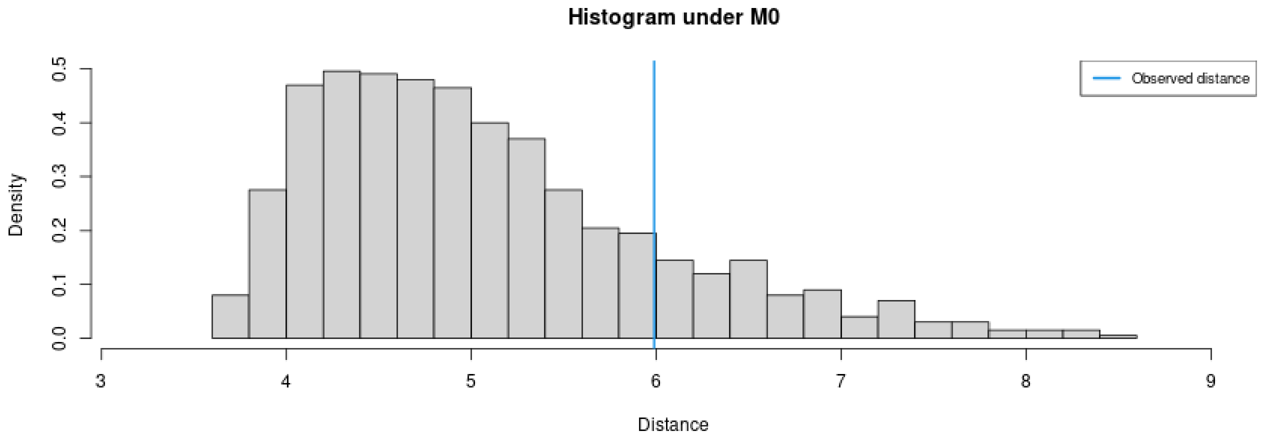

```
summary(res.gfit0.a)
```

```
## $pvalue
## [1] 0.161
##
## $s.dist.sim
##    Min. 1st Qu.  Median    Mean 3rd Qu.    Max.
##   3.665   4.375   4.886   5.079   5.568   8.527
##
```

```
## $dist.obs
## [1] 5.988453
```

Under the selection model:

```
if (redo) {
  res.gfit1.a <- gfit(target = stat.obs.a, sumstat = stat.sim1.a,
                      statistic = median, nb.replicate = 1000)
  save(res.gfit1.a, file = "Rdata/SexAveraged/backup_gfit1_allelic.Rdata")
} else {
  load("Rdata/SexAveraged/backup_gfit1_allelic.Rdata")
}
plot(res.gfit1.a, main = "Histogram under M1")
```

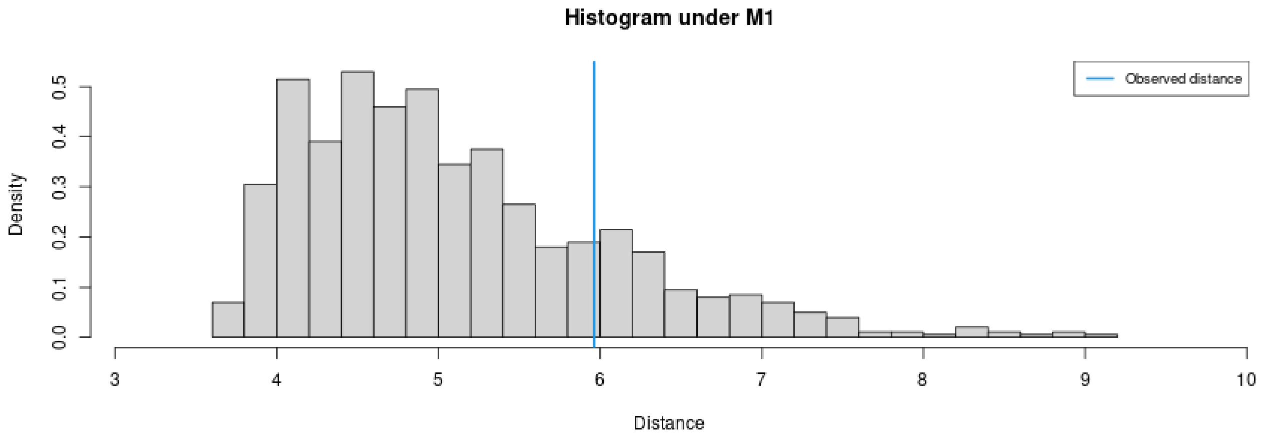

```
summary(res.gfit1.a)
```

```
## $pvalue
## [1] 0.184
##
## $s.dist.sim
##    Min. 1st Qu.  Median    Mean 3rd Qu.    Max.
##   3.668   4.376   4.891   5.095   5.600   9.196
##
## $dist.obs
## [1] 5.962703
```

Both models provide a reasonably good fit.

#### 3.4 Summary figure

Posterior distributions, with prior for comparison:

```
library(ggplot2)
library(ggpubr)
dat.prior.h <- data.frame(h = param.sim[,"h"], Distribution = "Prior")
dat.post.h <- data.frame(h = m.r.a$adj.values[,"h"], Distribution = "Posterior")
dat.prior.s <- data.frame(s = param.sim[,"s"], Distribution = "Prior")
dat.post.s <- data.frame(s = m.r.a$adj.values[,"s"], Distribution = "Posterior")
dat.h <- rbind(dat.prior.h, dat.post.h)
dat.s <- rbind(dat.prior.s, dat.post.s)
dat.h$Distribution <- factor(dat.h$Distribution, levels = c("Prior", "Posterior"))
dat.s$Distribution <- factor(dat.s$Distribution, levels = c("Prior", "Posterior"))
names(dat.h)[1] <- "value"
```

```r
dat.h$variable <- "h"
names(dat.s)[1] <- "value"
dat.s$variable <- "s"
dat.dist <- rbind(dat.h, dat.s)

p.dist <- ggplot(data = dat.dist,
                aes(x = value, linetype = Distribution, fill = Distribution)) +
          geom_density(alpha = 0.5) +
          scale_fill_brewer(type = "qual", palette = 3) +
          scale_linetype_manual(values = c(Prior = "dashed", Posterior = "solid")) +
          xlab("Parameter value") +
          facet_wrap(~variable, scales = "free") +
          ggtitle("Posterior and prior parameter distributions") +
          theme_pubclean() + theme(strip.background = element_blank())
```

Cross-validation:

```r
d1<-rbind(as.data.frame(cv.ridge.a$estim$tol0.01),
          as.data.frame(cv.ridge.a$estim$tol0.1),
          as.data.frame(cv.ridge.a$estim$tol0.2))
d2<-rbind(cv.ridge.a$true, cv.ridge.a$true, cv.ridge.a$true)
d1<-melt(d1, value.name = "Estimated")
```

```
## No id variables; using all as measure variables
```

```r
d1$Tolerance <- rep(c(0.01, 0.1, 0.2), each = 1000)
d1$Replicate <- rep(1:1000, 3)
d2<-melt(d2, value.name = "True")
```

```
## No id variables; using all as measure variables
```

```r
d2$Replicate <- rep(1:1000, 3)
dat.cv <- merge(d1, d2, by = c("variable", "Replicate"))
dat.cv$variable <- substr(dat.cv$variable, 1, 1)
```

```r
p.cv <- ggplot(dat.cv, aes(x = True, y = Estimated)) +
        geom_point(aes(col = as.ordered(Tolerance))) +
        geom_abline(slope = 1) +
        facet_wrap(~variable, scales = "free") +
        theme_pubclean() + labs(color = "Tolerance") +
        ggtitle("Cross validation") +
        theme(strip.background = element_blank())
```

Confusion matrix:

```r
library(scales)
cv.sum <- summary(cv.modsel.a)
```

```
## Confusion matrix based on 1000 samples for each model.
##
## $tol0.1
##           Neutral Selection
## Neutral       939        61
## Selection      65       935
##
##
## Mean model posterior probabilities (mnlogistic)
```

```
##
## $tol0.1
##           Neutral Selection
## Neutral    0.9112    0.0888
## Selection  0.0983    0.9017
```

```
dat.cv <- as.data.frame(cv.sum$conf.matrix$tol0.1/1000)
names(dat.cv) <- c("Real", "Inferred", "Frequency")
p.confmat <- ggplot(dat.cv, aes(x = Real, y = Frequency, fill = Inferred)) +
             geom_col() +
             scale_y_continuous(labels = scales::percent) +
             scale_fill_brewer(type = "qual", palette = 3) +
             ggtitle("Prediction errors") +
             theme_pubclean()
```

Model probabilities:

```
p.mprob <- ggplot(as.data.frame(modsel.a$pred), aes(x = Var1, y = Freq, fill = Var1)) +
           geom_col() + ylab("Model posterior probability") + xlab("Model") +
           scale_fill_brewer(type = "qual", palette = 3) +
           theme_pubclean() +
           ggtitle("Model probabilities") +
           theme(legend.position = "none")
```

```
library(cowplot)
```

```
##
## Attaching package: 'cowplot'

## The following object is masked from 'package:ggpubr':
##
##     get_legend
```

```
p1 <- plot_grid(p.mprob, p.confmat, labels = c("C", "D"), nrow = 1)
p <- plot_grid(p.dist, p.cv, p1, labels = c("A", "B", ""), nrow = 3)
p
```

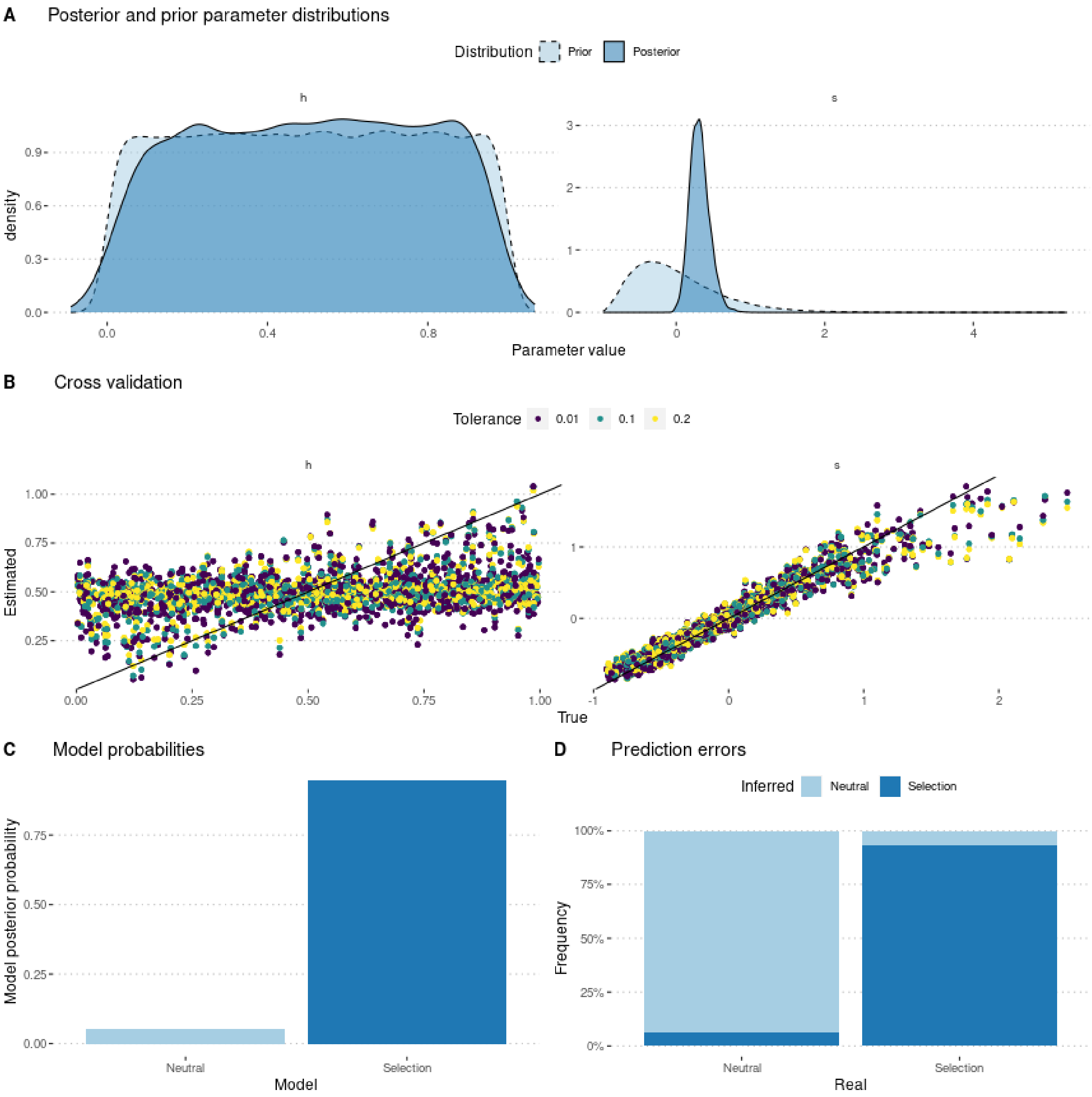

```r
ggsave(p, filename = "FigureABC-SexAveraged-Allelic.pdf", width = 8, height = 10)
```

### 4 Using genotype frequencies

We estimate parameters using the ridge regression method:

```r
if (redo) {
  m.r.g <- abc(target = stat.obs, param = param.sim, sumstat = stat.sim, tol = .1,
               method = "ridge", transf = c("none", "none"))
  save(m.r.g, file = "Rdata/SexAveraged/backup_abc_genotype.Rdata")
} else {
  load("Rdata/SexAveraged/backup_abc_genotype.Rdata")
}
```

Now we display the results:

```r
summary(m.r.g, intvl = .95)
```

```
## Call:
## abc(target = stat.obs, param = param.sim, sumstat = stat.sim,
##     tol = 0.1, method = "ridge", transf = c("none", "none"))
## Data:
##  abc.out$adj.values (10000 posterior samples)
## Weights:
##  abc.out$weights
##
##                               h       s
## Min.:                    0.0683 -0.1786
## Weighted 2.5 % Perc.:    0.1637 -0.0357
## Weighted Median:         0.4713  0.1411
## Weighted Mean:           0.4680  0.1474
## Weighted Mode:           0.5378  0.1363
## Weighted 97.5 % Perc.:   0.7699  0.3555
## Max.:                    0.8612  0.8899
```

Distribution of parameter estimates:

```r
hist(m.r.g, breaks = 30, caption = c(expression(h), expression(s)))
```

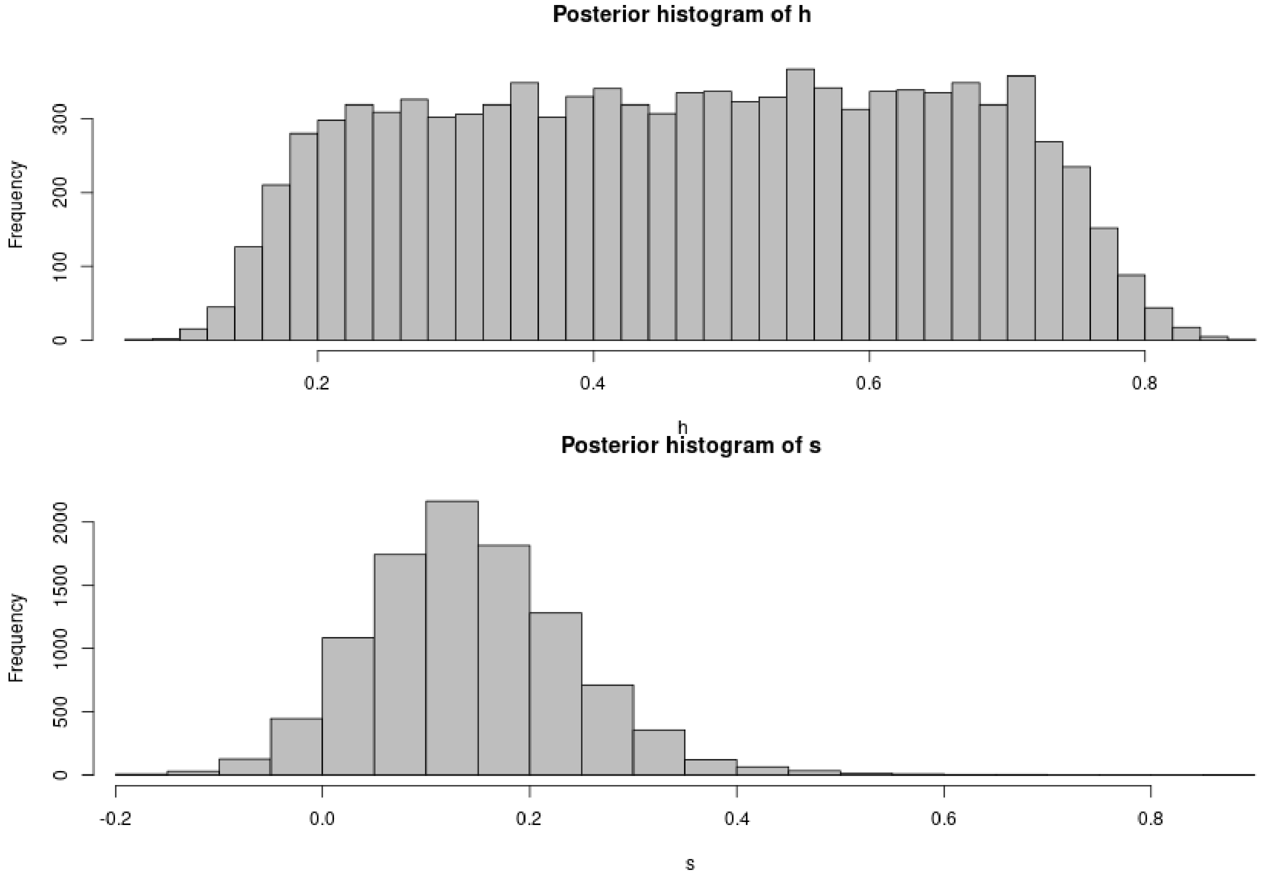

```
plot(m.r.g, param = param.sim)
```

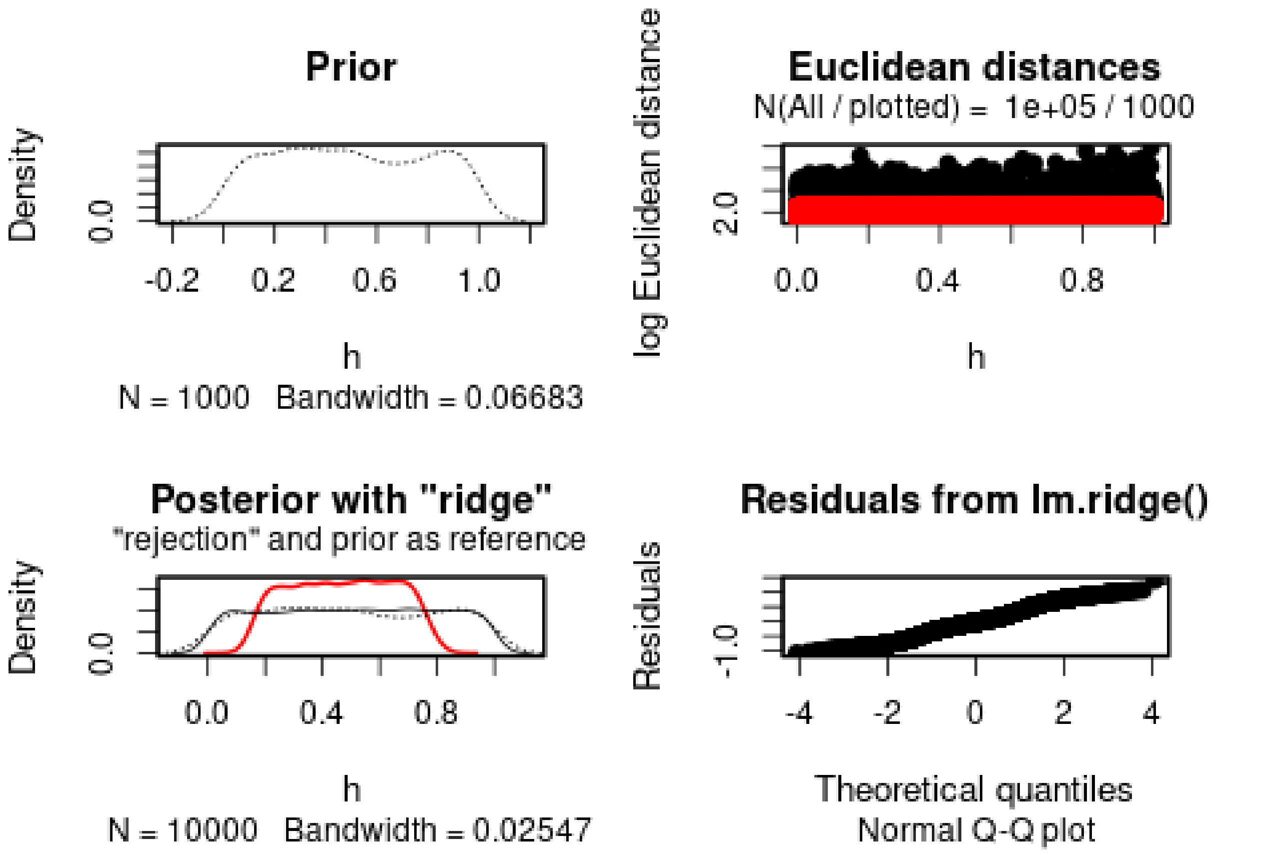

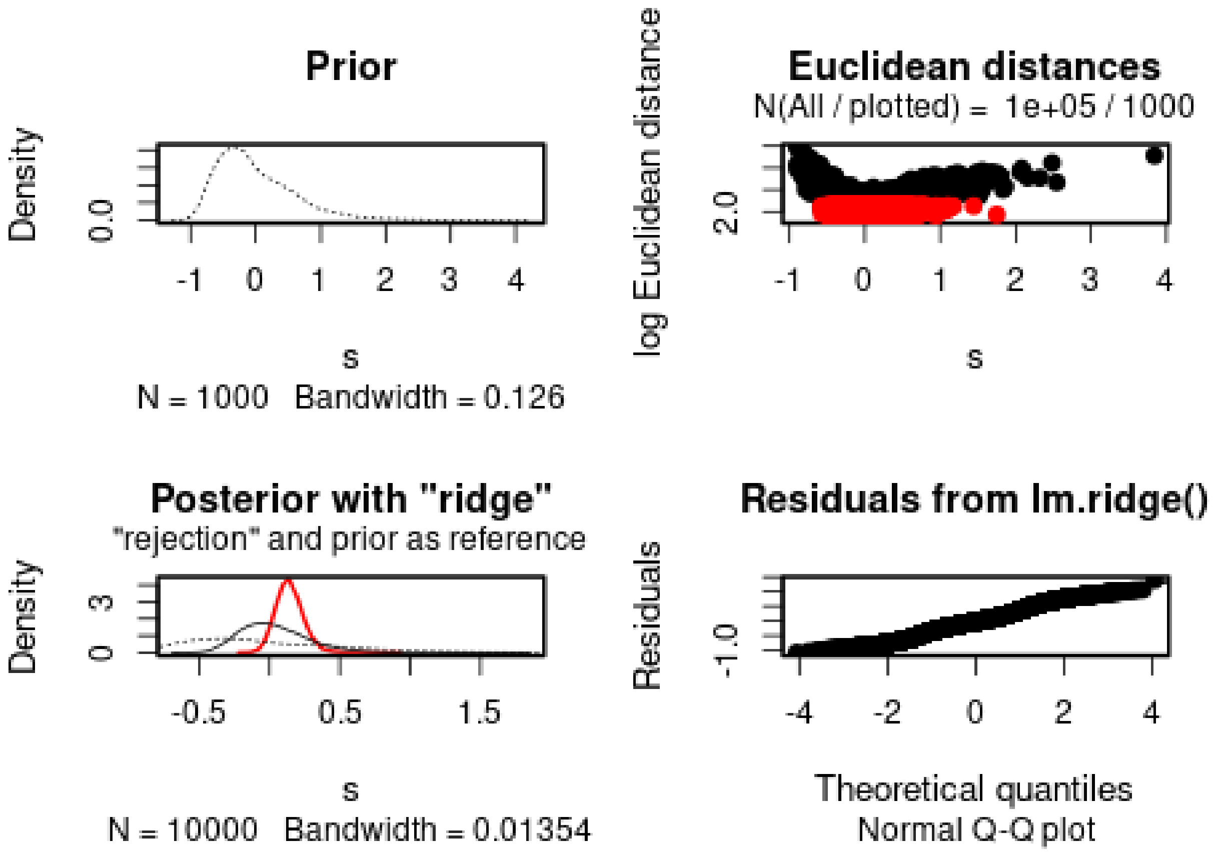

*h* cannot properly be estimated. *s* is positive (s = 0.1474), but has a large variance. The 95% posterior interval includes 0.

We then check the distribution of summary statistics in the simulations:

```
library(reshape2)
library(ggplot2)
library(ggpubr)

df.stats.sim <- melt(stat.sim, variable.name = "Statistic", value.name = "Frequency")
```

```
## No id variables; using all as measure variables
```

```
l <- strsplit(as.character(df.stats.sim$Statistic), split = ".", fixed = TRUE)
df.stats.sim$Room <- sapply(l, function(x) x[1])
df.stats.sim$Genotype <- sapply(l, function(x) x[2])
df.stats.sim$Generation <- sapply(
  strsplit(as.character(df.stats.sim$Statistic), split = ".", fixed = TRUE),
  function(x) x[3])

df.stats.obs <- as.data.frame(stat.obs)
names(df.stats.obs) <- "Frequency"
df.stats.obs$Statistic <- row.names(df.stats.obs)
l <- strsplit(as.character(df.stats.obs$Statistic), split = ".", fixed = TRUE)
df.stats.obs$Room <- sapply(l, function(x) x[1])
df.stats.obs$Genotype <- sapply(l, function(x) x[2])
df.stats.obs$Generation <- sapply(
  strsplit(as.character(df.stats.obs$Statistic), split = ".", fixed = TRUE),
  function(x) x[3])
```

```r
plot.obs <- function(gen) {
  p <- ggplot(data = subset(df.stats.sim, Generation == gen),
              aes(x = Frequency, y = ..density..)) +
    geom_histogram(bins = 50) +
    geom_vline(data = subset(df.stats.obs, Generation == gen),
               aes(xintercept = Frequency), color = "orange") +
    facet_grid(Genotype~Room) +
    theme_pubclean()
  return(p)
}
```

```r
plot.obs(1) + ggtitle("Timepoint 4")
```

```
## Warning: The dot-dot notation (`..density..`) was deprecated in ggplot2 3.4.0.
## i Please use `after_stat(density)` instead.
## This warning is displayed once every 8 hours.
## Call `lifecycle::last_lifecycle_warnings()` to see where this warning was
## generated.
```

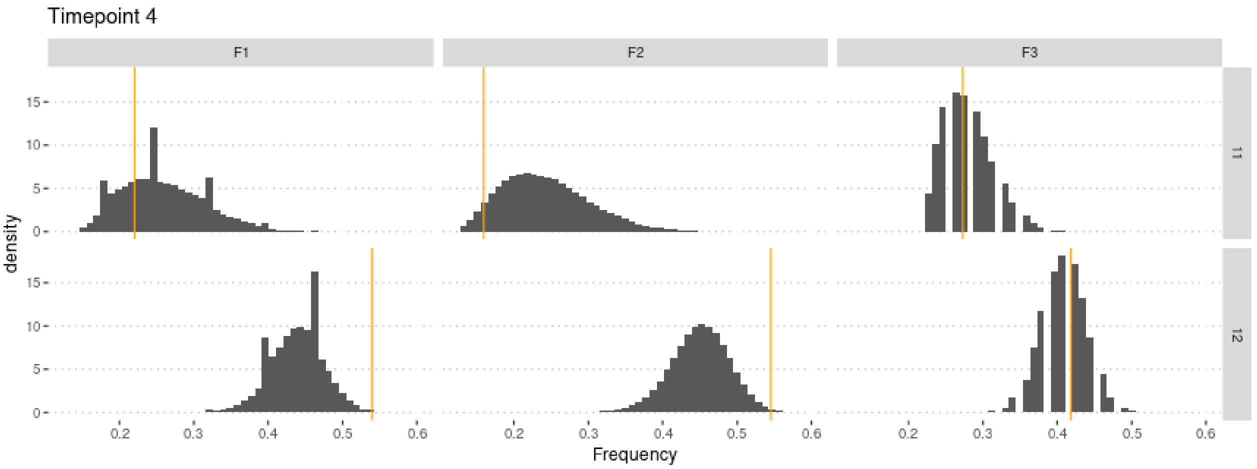

```r
plot.obs(2) + ggtitle("Timepoint 5")
```

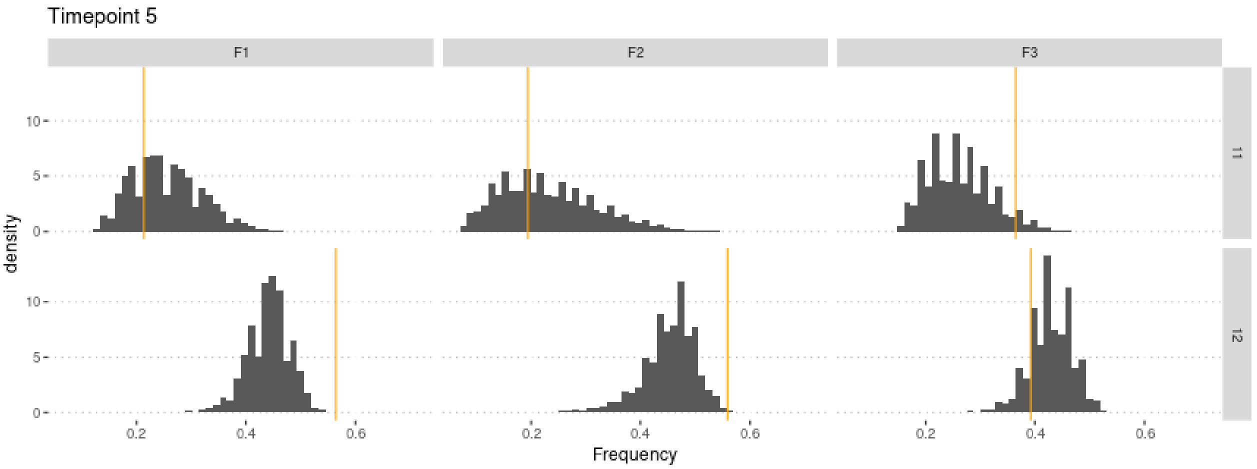

```
plot.obs(3) + ggtitle("Timepoint 6")
```

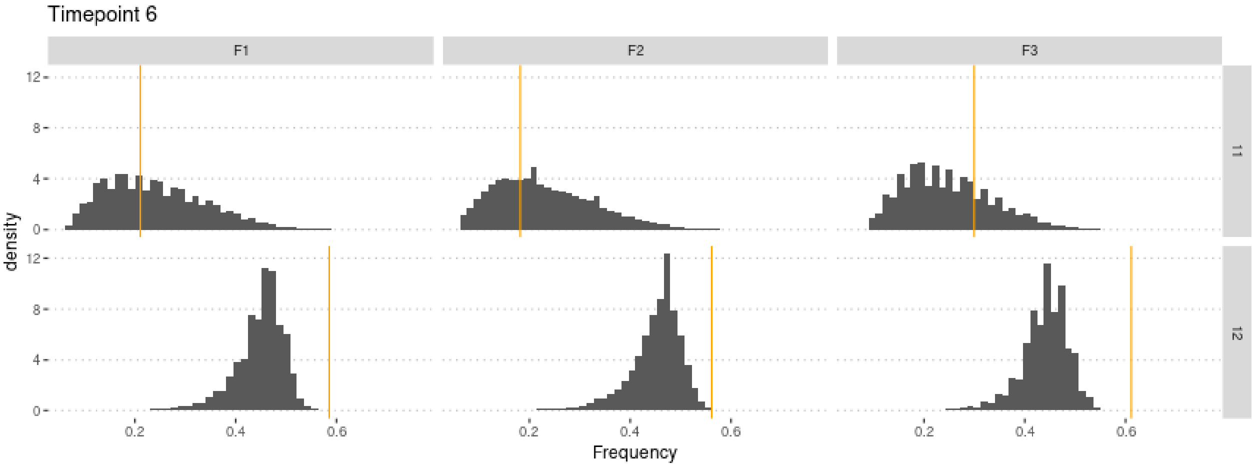

```
plot.obs(4) + ggtitle("Timepoint 7")
```

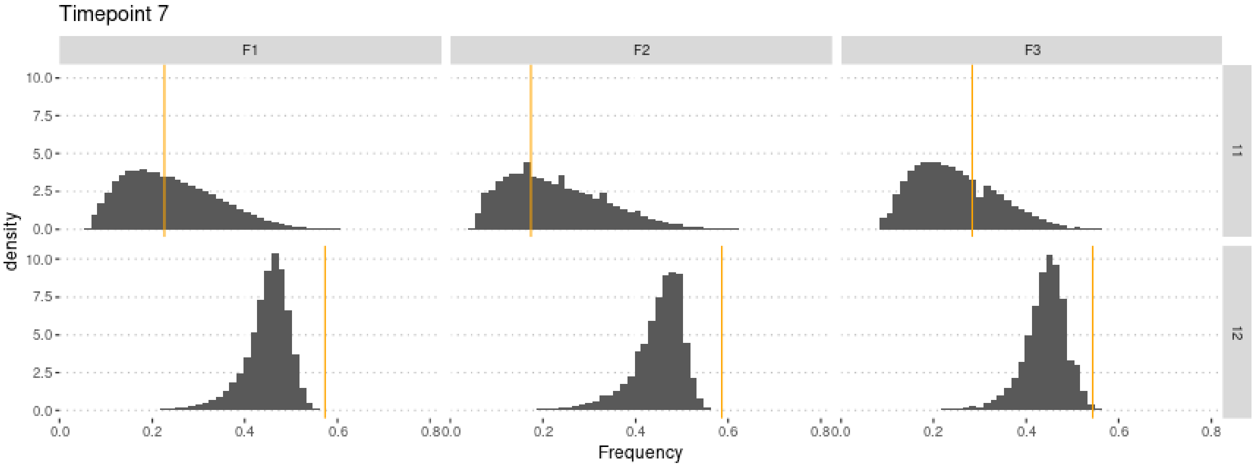

```
plot.obs(5) + ggtitle("Timepoint 8")
```

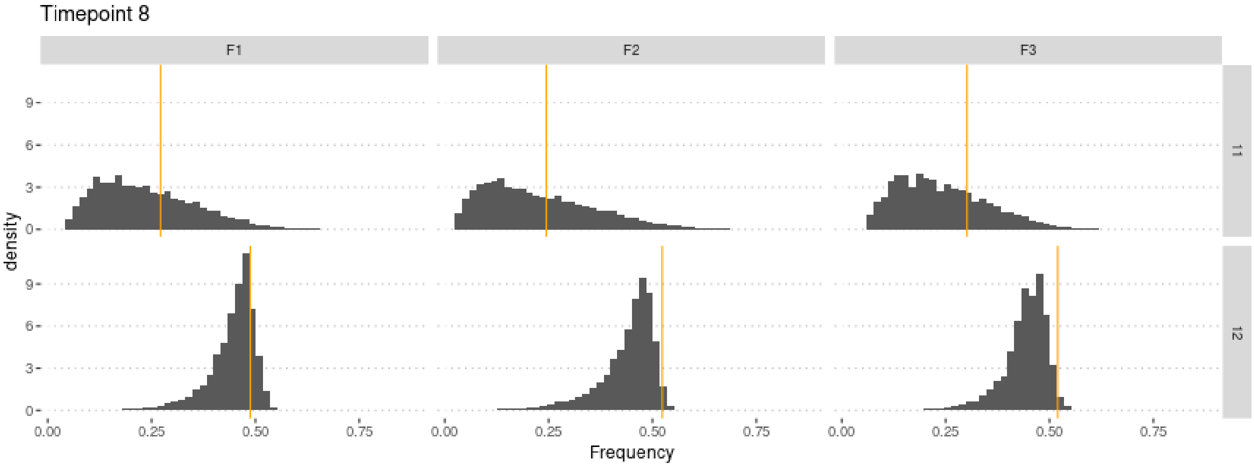

#### 4.1 Cross-Validation analysis

Compute predictions errors:

```r
if (redo) {
  cv.ridge.g <- cv4abc(
      param = param.sim,
      sumstat = stat.sim,
      abc.out = m.r.g,
      nval = 1000,
      tols = c(.01, .1, .2))
  save(cv.ridge.g, file = "Rdata/SexAveraged/backup_cv_genotype.Rdata")
} else {
  load("Rdata/SexAveraged/backup_cv_genotype.Rdata")
}
```

```r
summary(cv.ridge.g)
```

```
## Prediction error based on a cross-validation sample of 1000
```

```
##               h          s
## 0.01 0.91459359 0.07568590
## 0.1  0.86774971 0.07682655
## 0.2  0.86890486 0.07743617
```

```r
plot(cv.ridge.g)
```

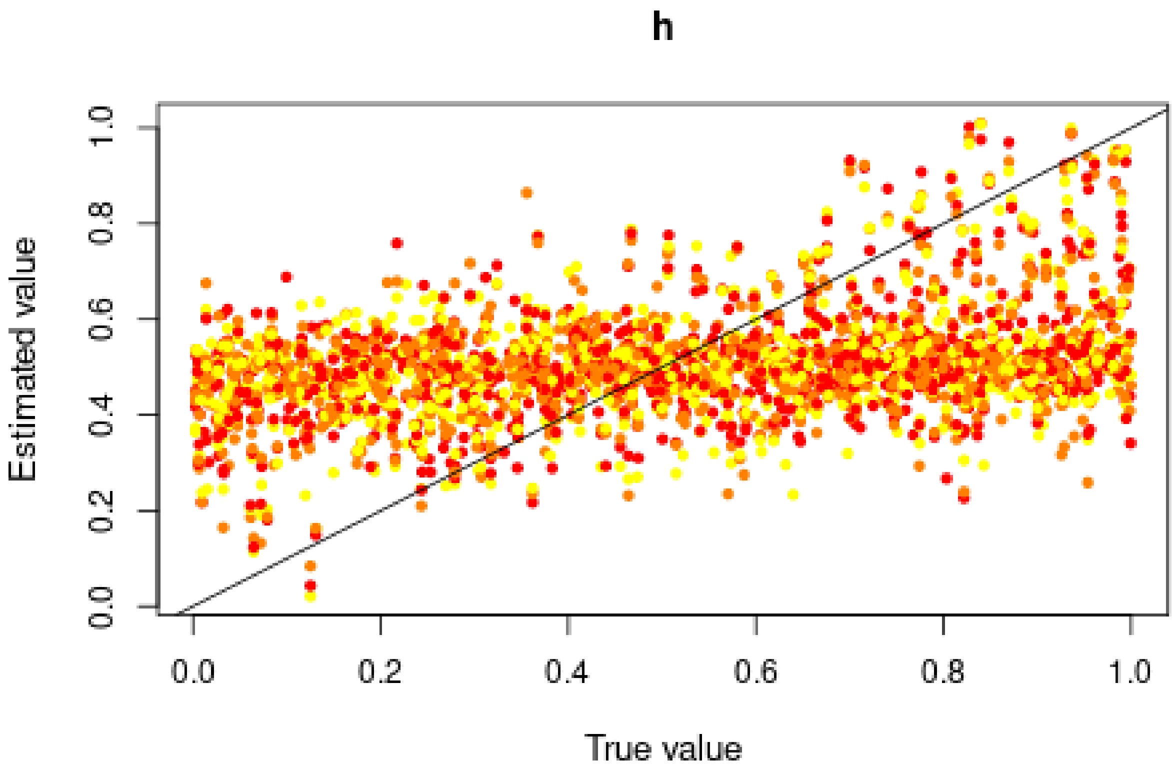

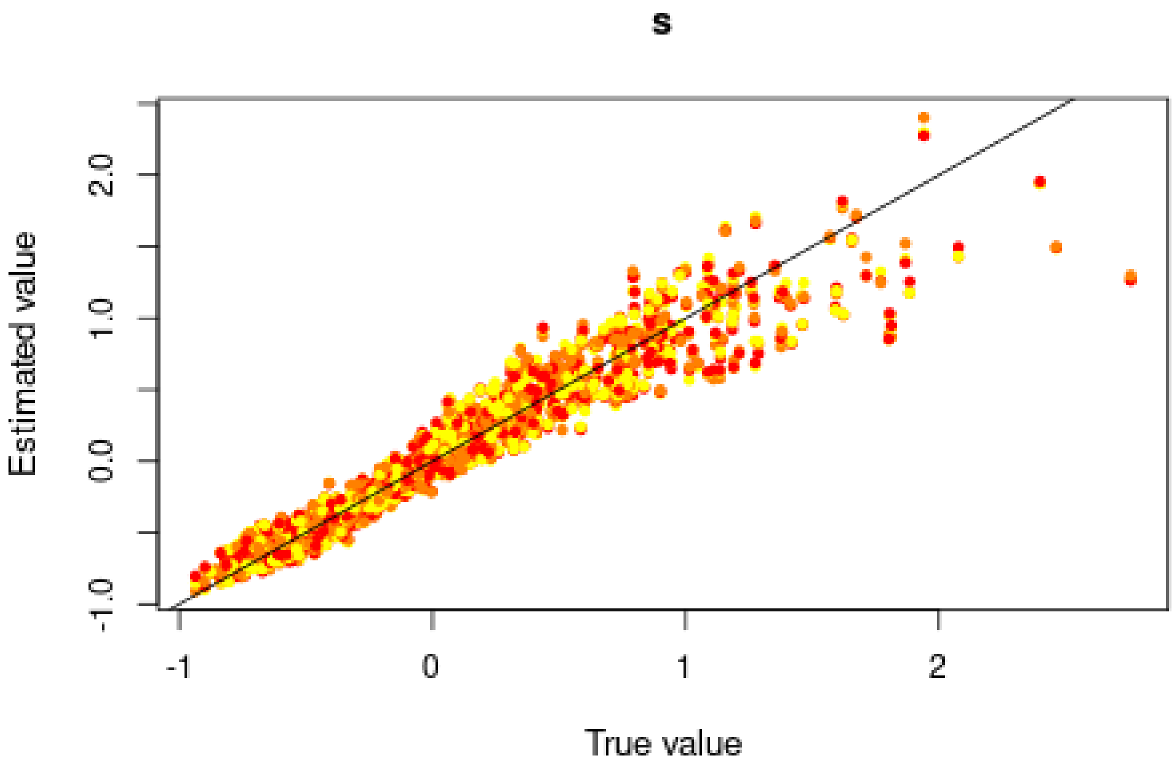

Good for $s$, but very bad for $h$, in agreement with the posterior distributions.

#### 4.2 Misclassification errors

To test the power of the approach to distinguish between models, we also conduct a cross-validation experiment. We compare model M0 ($s = 0$) and M1 ($s = 0.1474 > 0$). $h$ is sampled over its prior distribution. 100,000 simulations were conducted under each model.

```
sims0 <- read.csv("Simulations/Simulations0FromGen3.csv.gz")
param.sim0 <- subset(sims0, select = c("h", "s"))
stat.sim0 <- subset(sims0, select = 2:46)
# Only consider the WT homozygotes and the heterozygotes,
# as the sum of the three genotypes is 1
stat.sim0 <- subset(stat.sim0, select = grep(".22.", names(stat.sim0), invert = TRUE))

sims1 <- read.csv("Simulations/Simulations1bFromGen3.csv.gz")
param.sim1 <- subset(sims1, select = c("h", "s"))
stat.sim1 <- subset(sims1, select = 2:46)
# Only consider the WT homozygotes and the heterozygotes,
# as the sum of the three genotypes is 1
stat.sim1 <- subset(stat.sim1, select = grep(".22.", names(stat.sim1), invert = TRUE))
```

We conduct a CV analysis (this takes some time... set nval = 100 for faster results):

```
models <- rep(c("Neutral", "Selection"), each = 100000)
if (redo) {
  cv.modsel.g <- cv4postpr(
    models, rbind(stat.sim0, stat.sim1),
    nval = 1000, tols=.1, method = "mnlogistic")
  save(cv.modsel.g, file = "Rdata/SexAveraged/backup_cv4postpr_genotype.Rdata")
} else {
```

```r
  load("Rdata/SexAveraged/backup_cv4postpr_genotype.Rdata")
}
```

We display the results:

```r
summary(cv.modsel.g)
```

```
## Confusion matrix based on 1000 samples for each model.
##
## $tol0.1
##           Neutral Selection
## Neutral       779       221
## Selection     208       792
##
##
## Mean model posterior probabilities (mnlogistic)
##
## $tol0.1
##           Neutral Selection
## Neutral    0.7000    0.3000
## Selection  0.2892    0.7108
```

```r
plot(cv.modsel.g, names.arg=c("Neutral", "Selection"))
```

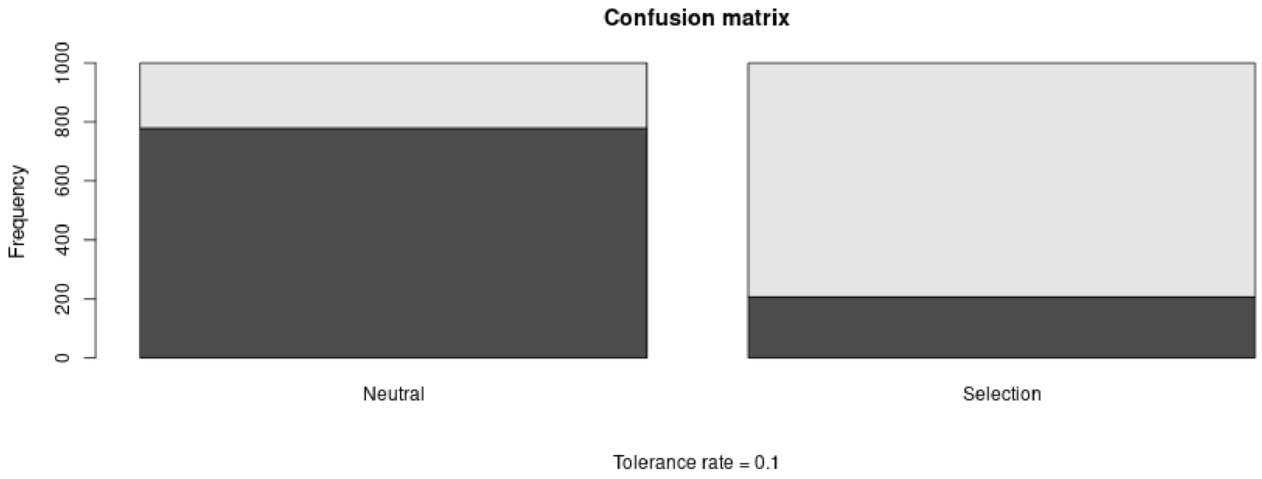

#### 4.3 Posterior prediction

```r
if (redo) {
  modsel.g <- postpr(
    stat.obs, models,
    rbind(stat.sim0, stat.sim1),
    tol = .1, method = "mnlogistic")
  save(modsel.g, file = "Rdata/SexAveraged/backup_postpr_genotype.Rdata")
} else {
  load("Rdata/SexAveraged/backup_postpr_genotype.Rdata")
}
summary(modsel.g)
```

```
## Call:
## postpr(target = stat.obs, index = models, sumstat = rbind(stat.sim0,
##     stat.sim1), tol = 0.1, method = "mnlogistic")
```

```
## Data:
##  postpr.out$values (20000 posterior samples)
## Models a priori:
##  Neutral, Selection
## Models a posteriori:
##  Neutral, Selection
##
## Proportion of accepted simulations (rejection):
##   Neutral Selection
##    0.5471    0.4528
##
## Bayes factors:
##           Neutral Selection
## Neutral    1.0000    1.2082
## Selection  0.8277    1.0000
##
##
## Posterior model probabilities (mnlogistic):
##   Neutral Selection
##    0.0766    0.9234
##
## Bayes factors:
##           Neutral Selection
## Neutral    1.0000    0.0829
## Selection 12.0558    1.0000
```

The model with selection is preferred.

#### 4.4 Goodness of fit

Under the neutral model:

```r
if (redo) {
  res.gfit0.g <- gfit(
    target = stat.obs, sumstat = stat.sim0,
    statistic = median, nb.replicate = 1000)
  save(res.gfit0.g, file = "Rdata/SexAveraged/backup_gfit0_genotype.Rdata")
} else {
  load("Rdata/SexAveraged/backup_gfit0_genotype.Rdata")
}
plot(res.gfit0.g, main = "Histogram under M0")
```

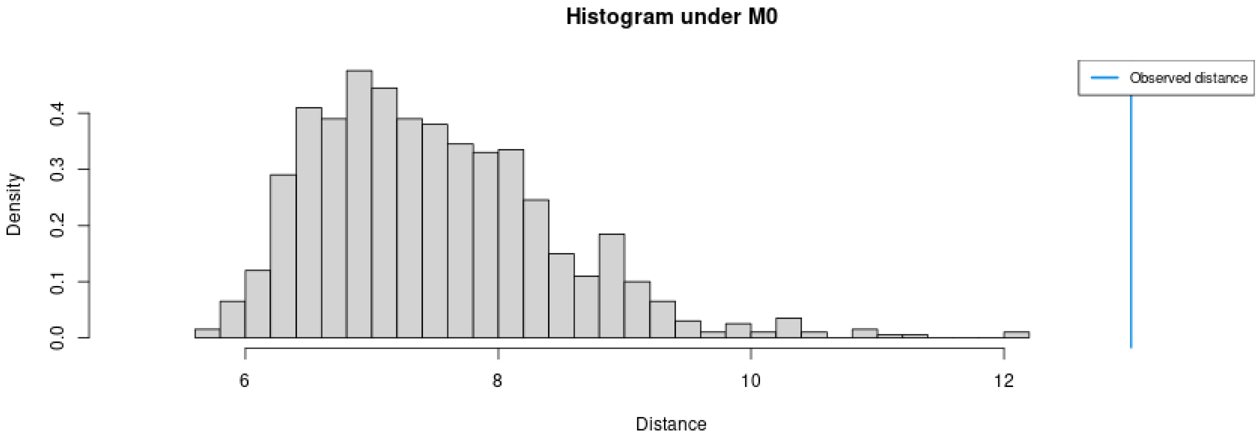

```r
summary(res.gfit0.g)
```

```
## $pvalue
## [1] 0
##
## $s.dist.sim
##    Min. 1st Qu.  Median    Mean 3rd Qu.    Max.
##   5.740   6.783   7.317   7.489   8.049  12.086
##
## $dist.obs
## [1] 13.00821
```

Under the selection model:

```r
if (redo) {
  res.gfit1.g <- gfit(
    target = stat.obs, sumstat = stat.sim1,
    statistic = median, nb.replicate = 1000)
  save(res.gfit1.g, file = "Rdata/SexAveraged/backup_gfit1_genotype.Rdata")
} else {
  load("Rdata/SexAveraged/backup_gfit1_genotype.Rdata")
}
plot(res.gfit1.g, main = "Histogram under M1")
```

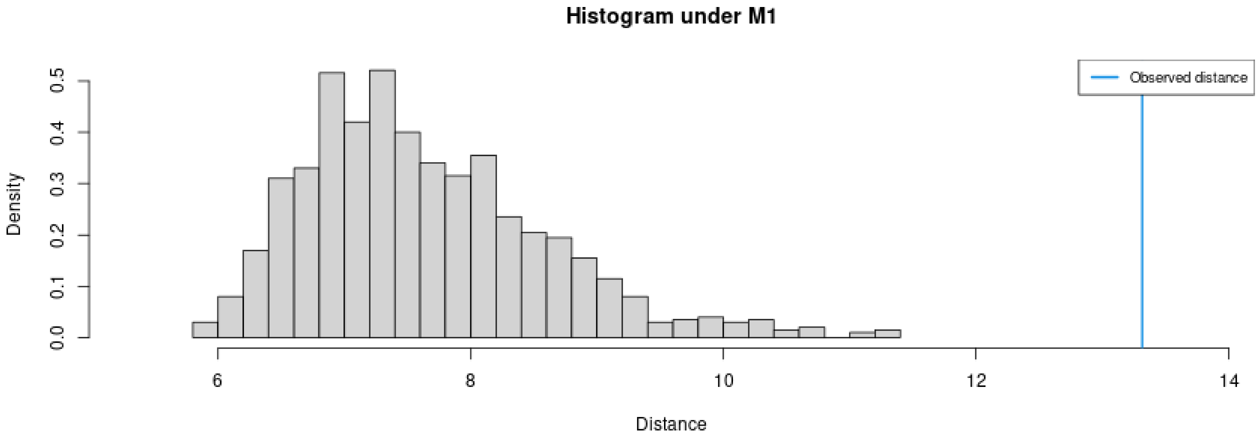

```r
summary(res.gfit1.g)
```

```
## $pvalue
## [1] 0
##
## $s.dist.sim
##    Min. 1st Qu.  Median    Mean 3rd Qu.    Max.
##   5.868   6.941   7.452   7.635   8.188  11.333
##
## $dist.obs
## [1] 13.31615
```

Both models provide a bad fit.

#### 4.5 Summary figure

Posterior distributions, with prior for comparison:

Confusion matrix:

```
library(scales)
cv.sum <- summary(cv.modsel.g)
```

```
## Confusion matrix based on 1000 samples for each model.
##
## $tol0.1
##           Neutral Selection
## Neutral       779       221
## Selection     208       792
##
##
## Mean model posterior probabilities (mnlogistic)
##
## $tol0.1
##           Neutral Selection
## Neutral    0.7000    0.3000
## Selection  0.2892    0.7108
```

```
dat.cv <- as.data.frame(cv.sum$conf.matrix$tol0.1/1000)
names(dat.cv) <- c("Real", "Inferred", "Frequency")
p.confmat <- ggplot(dat.cv, aes(x = Real, y = Frequency, fill = Inferred)) +
             geom_col() +
             scale_y_continuous(labels = scales::percent) +
             scale_fill_brewer(type = "qual", palette = 3) +
             ggtitle("Prediction errors") +
             theme_pubclean()
```

Model probabilities:

```
p.mprob <- ggplot(as.data.frame(modsel.g$pred), aes(x = Var1, y = Freq, fill = Var1)) +
           geom_col() + ylab("Model posterior probability") + xlab("Model") +
           scale_fill_brewer(type = "qual", palette = 3) +
           theme_pubclean() +
           ggtitle("Model probabilities") +
           theme(legend.position = "none")
```

```
library(cowplot)
p1 <- plot_grid(p.mprob, p.confmat, labels = c("C", "D"), nrow = 1)
p <- plot_grid(p.dist, p.cv, p1, labels = c("A", "B", ""), nrow = 3)
p
```

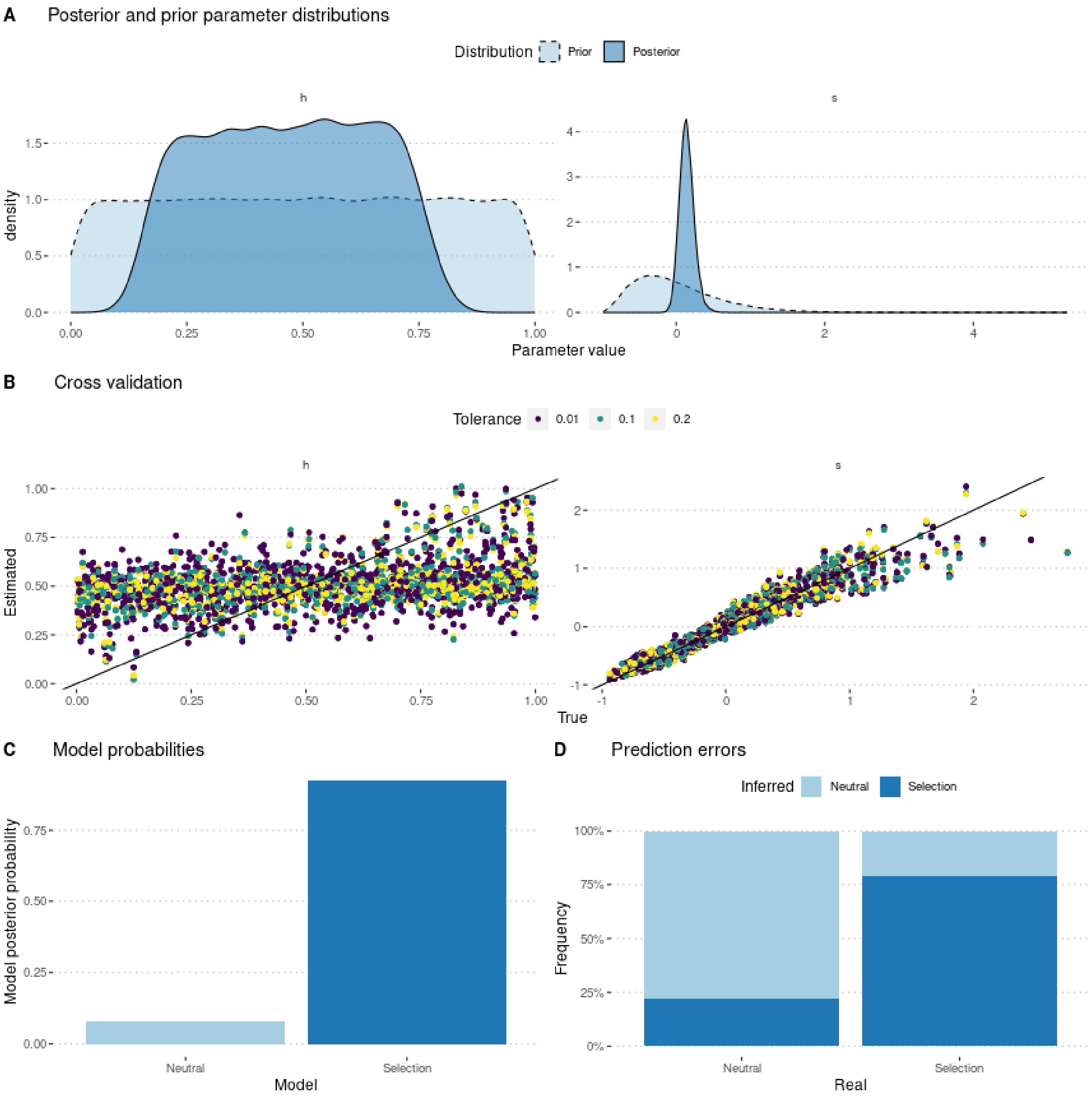

```r
ggsave(p, filename = "FigureABC-SexAveraged-Genotype.pdf", width = 8, height = 10)
```
