## Supplementary material for "Experimental evaluation of a direct fitness effect of the *de novo* evolved mouse gene *Pldi*": combined suppl files: SupplementaryFile4.pdf

### Estimating poldi's selection coefficient

Julien Y. Dutheil

01/02/2023

#### Contents


1 **Model parameterization** 1

2 **Preamble** 1

3 **Using allele frequencies** 2

4 **Using genotype frequencies** 8


### 1 Model parameterization

The model includes two parameters: $s$ the selection coefficient of the *poldi* allele compared to the knockout strain, which mimics the ancestral state, and $h$ the heterozygosity. Fitness values of each genotype are parameterized as follow:

$$\begin{aligned} poldi/poldi &: 1 \\ BL6/poldi &: 1 + h \cdot s \\ BL6/BL6 &: 1 + s \end{aligned}$$

where BL6 denotes the wild strain, and poldi the strain where the *poldi* gene is knocked out. Furthermore, we consider that the allele has a distinct effect in males and in females, so that we have four parameters *hf*, *sf*, *hm*, and *sm*.

We performed 400,000 simulations, using a gamma prior for *sf* and *sm* and a uniform prior between 0 and 1 for *hf* and *hm*. Male and female coefficients are distinct and drawn independently.

### 2 Preamble

We load the data (observed and simulated):

```
sims <- read.csv("Simulations/SimulationsSexSpecificFromGen3.csv.gz")
param.sim <- subset(sims, select = c("hf", "sf", "hm", "sm"))
stat.sim <- subset(sims, select = c(2:46))
# Only consider the WT homozygotes and the heterozygotes,
# as the sum of the three genotypes is 1
stat.sim <- subset(stat.sim, select = grep(".22.", names(stat.sim), invert = TRUE))

library(plyr, quietly = TRUE)
geno <- read.csv("GenotypeFrequencies.csv")
```

```r
stat.obs.a[1]  <- stat.obs["F1.11.1"] + stat.obs["F1.12.1"] / 2
stat.obs.a[2]  <- stat.obs["F1.11.2"] + stat.obs["F1.12.2"] / 2
stat.obs.a[3]  <- stat.obs["F1.11.3"] + stat.obs["F1.12.3"] / 2
stat.obs.a[4]  <- stat.obs["F1.11.4"] + stat.obs["F1.12.4"] / 2
stat.obs.a[5]  <- stat.obs["F1.11.5"] + stat.obs["F1.12.5"] / 2
#Pop2
stat.obs.a[6]  <- stat.obs["F2.11.1"] + stat.obs["F2.12.1"] / 2
stat.obs.a[7]  <- stat.obs["F2.11.2"] + stat.obs["F2.12.2"] / 2
stat.obs.a[8]  <- stat.obs["F2.11.3"] + stat.obs["F2.12.3"] / 2
stat.obs.a[9]  <- stat.obs["F2.11.4"] + stat.obs["F2.12.4"] / 2
stat.obs.a[10] <- stat.obs["F2.11.5"] + stat.obs["F2.12.5"] / 2
#Pop3
stat.obs.a[11] <- stat.obs["F3.11.1"] + stat.obs["F3.12.1"] / 2
stat.obs.a[12] <- stat.obs["F3.11.2"] + stat.obs["F3.12.2"] / 2
stat.obs.a[13] <- stat.obs["F3.11.3"] + stat.obs["F3.12.3"] / 2
stat.obs.a[14] <- stat.obs["F3.11.4"] + stat.obs["F3.12.4"] / 2
stat.obs.a[15] <- stat.obs["F3.11.5"] + stat.obs["F3.12.5"] / 2
```

We estimate parameters using the ridge regression method:

```r
library(abc, quietly = TRUE)
```

```
##
## Attaching package: 'SparseM'

## The following object is masked from 'package:base':
##
##     backsolve

## locfit 1.5-9.8    2023-06-11
```

```r
if (redo) {
  m.r.a <- abc(target = stat.obs.a, param = param.sim, sumstat = stat.sim.a, tol=.1,
             method = "ridge", transf = c("none", "none", "none", "none"))
  save(m.r.a, file = "Rdata/SexSpecific/backup_abc_allelic.Rdata")
} else {
  load("Rdata/SexSpecific/backup_abc_allelic.Rdata")
}
```

Now we display the results:

```r
summary(m.r.a, intvl = .95)
```

```
## Call:
## abc(target = stat.obs.a, param = param.sim, sumstat = stat.sim.a,
##     tol = 0.1, method = "ridge", transf = c("none", "none", "none",
##         "none"))
## Data:
##  abc.out$adj.values (40000 posterior samples)
## Weights:
##  abc.out$weights
##
##                              hf      sf      hm      sm
## Min.:                   -0.0816 -0.8177 -0.0693 -0.7348
## Weighted 2.5 % Perc.:   -0.0001 -0.3762  0.0068 -0.2997
## Weighted Median:         0.4910  0.2705  0.4875  0.2490
## Weighted Mean:           0.4906  0.3616  0.4879  0.3284
```

```
## Weighted Mode:           0.2635  0.1082  0.0936  0.0879
## Weighted 97.5 % Perc.:   0.9821  1.5958  0.9742  1.4126
## Max.:                    1.0613  4.3444  1.0212  3.4356
```

Distribution of parameter estimates:

```r
hist(m.r.a, breaks = 30, caption = c(expression(h), expression(s)))
```

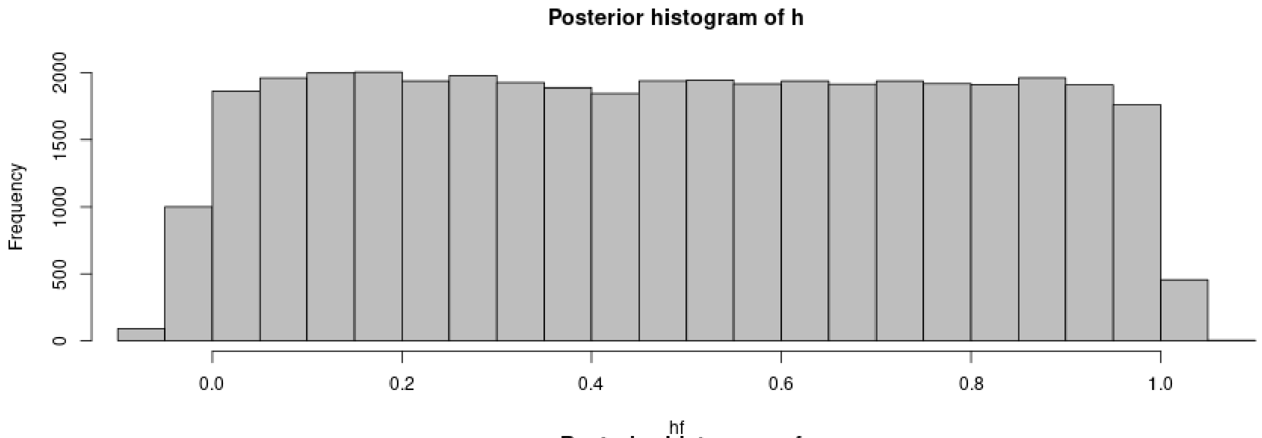


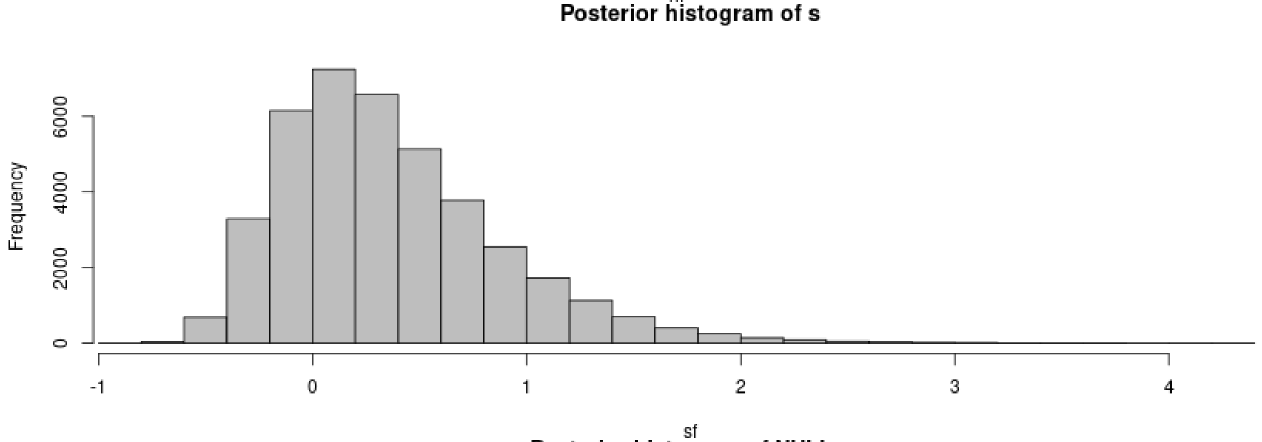


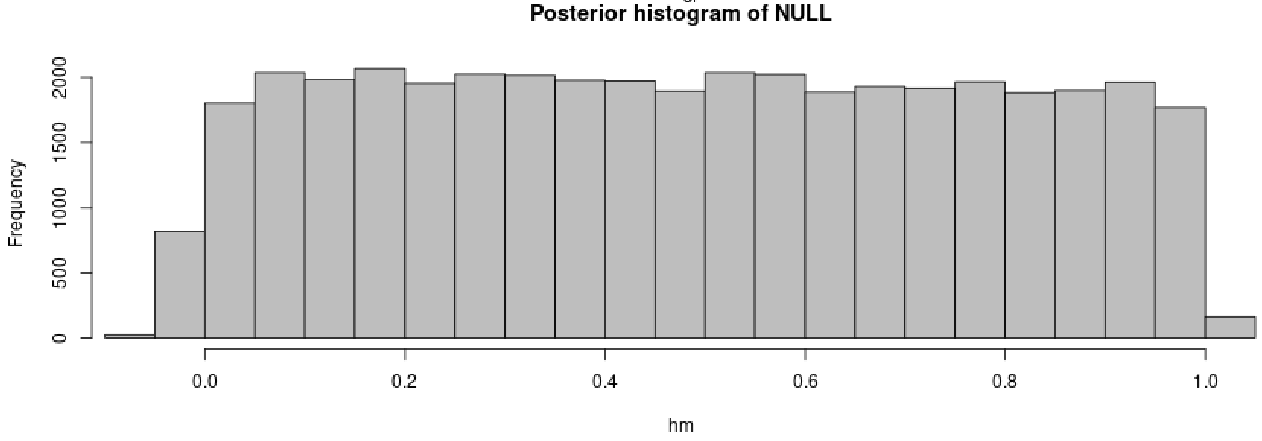

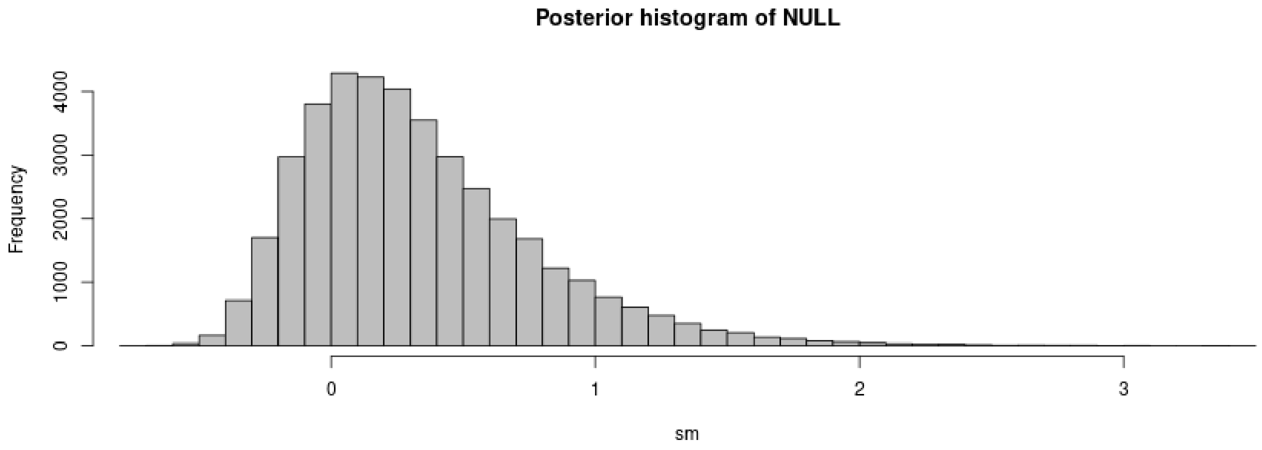


```r
plot(m.r.a, param = param.sim)
```

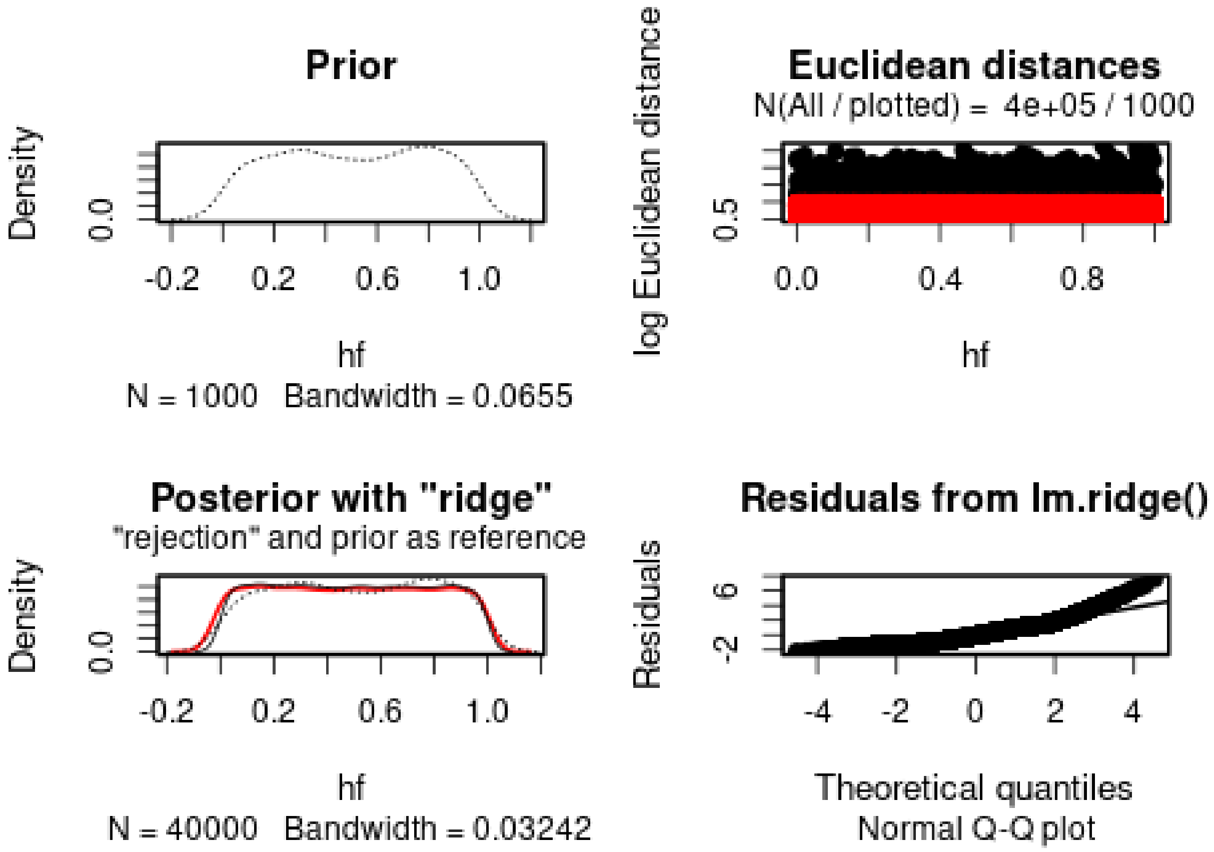

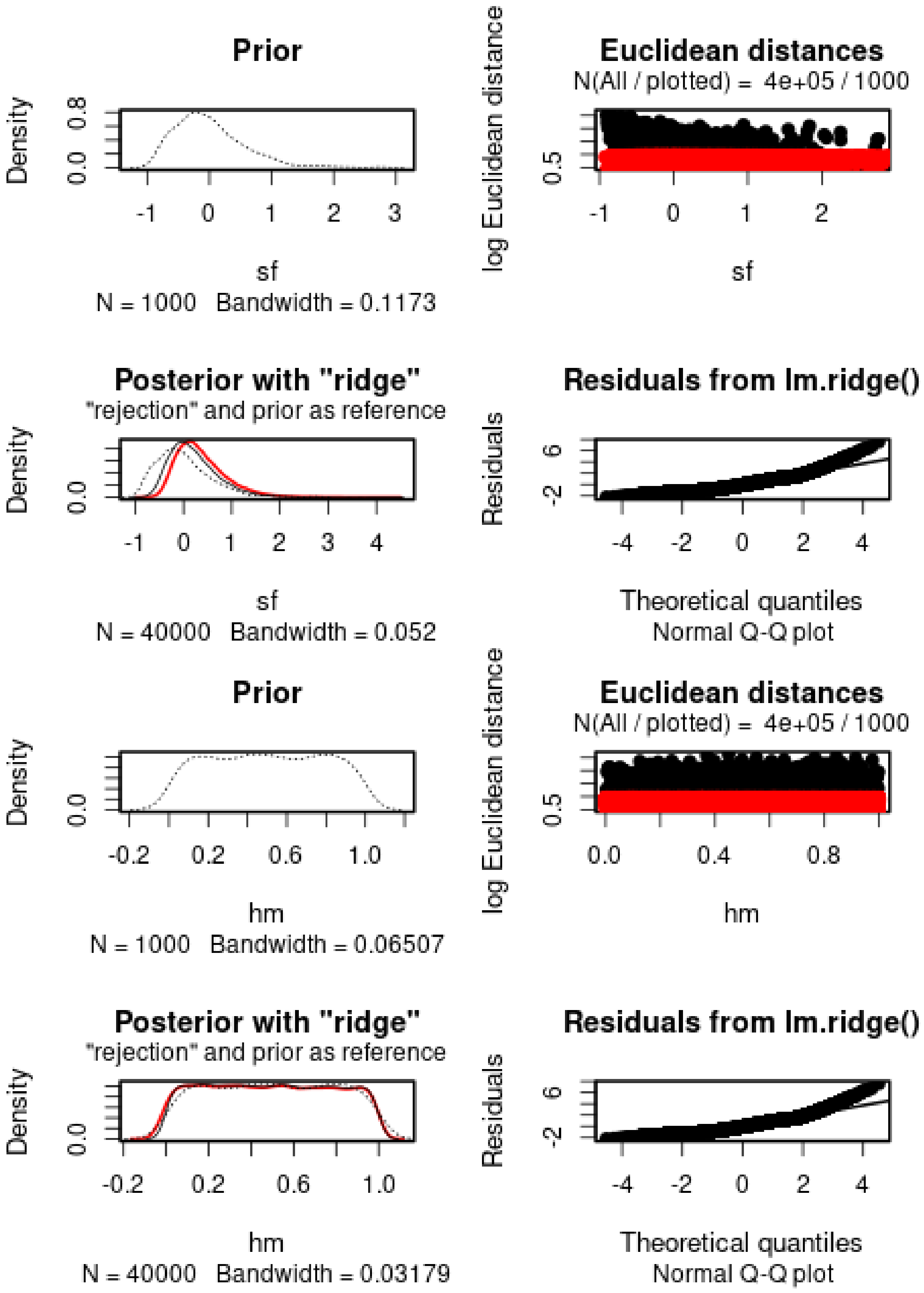
Prior
Density
0.0
0.8
-1
0
1
2
3
sf
N = 1000   Bandwidth = 0.1173
Euclidean distances
N(All / plotted) = 4e+05 / 1000
log Euclidean distance
0.5
-1
0
1
2
sf
Posterior with "ridge"
"rejection" and prior as reference
Density
0.0
-1
0
1
2
3
4
sf
N = 40000   Bandwidth = 0.052
Residuals from lm.ridge()
Residuals
-2
6
-4
-2
0
2
4
Theoretical quantiles
Normal Q-Q plot
Prior
Density
0.0
-0.2
0.2
0.6
1.0
hm
N = 1000   Bandwidth = 0.06507
Euclidean distances
N(All / plotted) = 4e+05 / 1000
log Euclidean distance
0.5
0.0
0.4
0.8
hm
Posterior with "ridge"
"rejection" and prior as reference
Density
0.0
-0.2
0.2
0.6
1.0
hm
N = 40000   Bandwidth = 0.03179
Residuals from lm.ridge()
Residuals
-2
6
-4
-2
0
2
4
Theoretical quantiles
Normal Q-Q plot

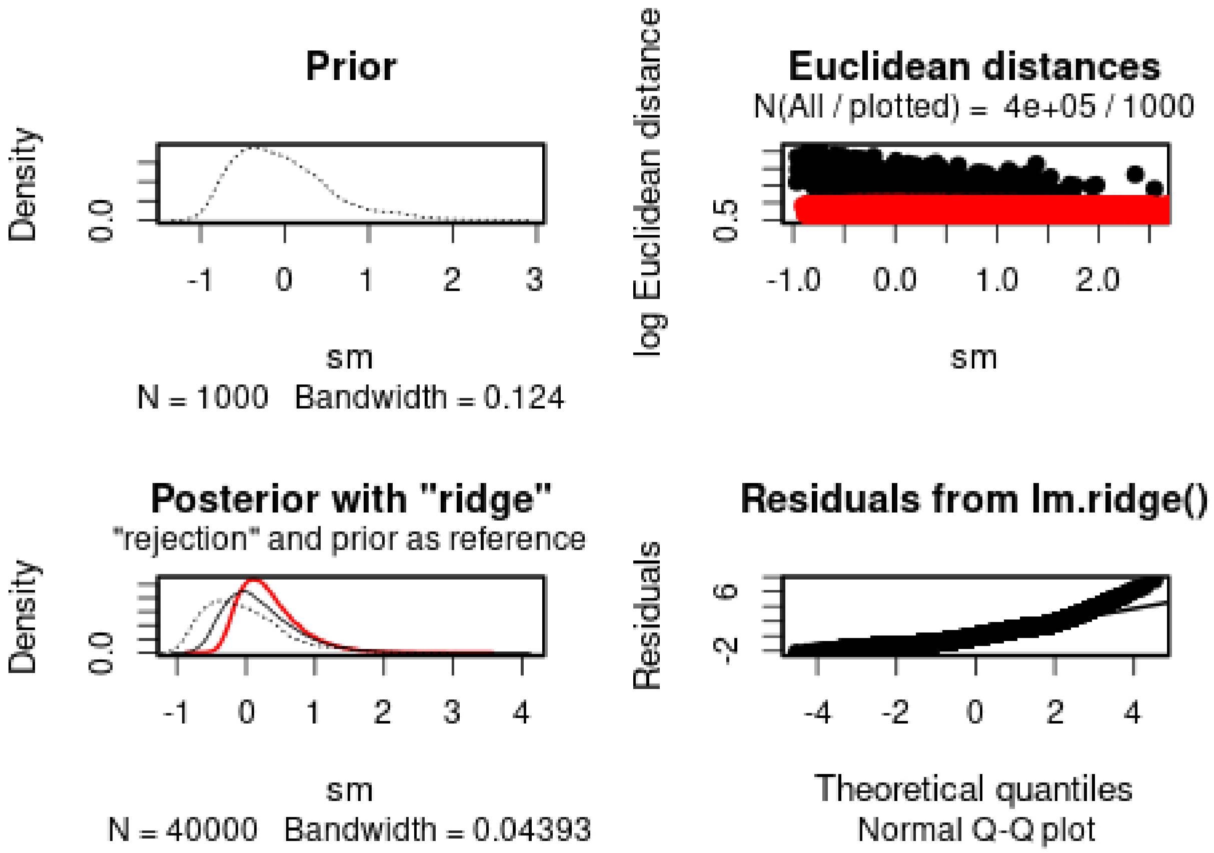


The *h*s cannot properly be estimated. The *s*s are positive, but have a large variance. The 95% posterior intervals include 0. Similar values are inferred for males and females.

We then check the distribution of summary statistics in the simulations:

```r
library(reshape2)
library(ggplot2)
library(ggpubr)
```

```
##
## Attaching package: 'ggpubr'

## The following object is masked from 'package:plyr':
##
##     mutate
```

```r
df.stats.sim <- melt(stat.sim.a, variable.name = "Statistic", value.name = "Frequency")
```

```
## No id variables; using all as measure variables
```

```r
l <- strsplit(as.character(df.stats.sim$Statistic), split = ".", fixed = TRUE)
df.stats.sim$Room <- sapply(l, function(x) x[1])
df.stats.sim$Generation <- sapply(
  strsplit(as.character(df.stats.sim$Statistic), split = ".", fixed = TRUE),
  function(x) x[2])

df.stats.obs <- as.data.frame(stat.obs.a)
names(df.stats.obs) <- "Frequency"
df.stats.obs$Statistic <- row.names(df.stats.obs)
l <- strsplit(as.character(df.stats.obs$Statistic), split = ".", fixed = TRUE)
```

```
df.stats.obs$Room <- sapply(l, function(x) x[1])
df.stats.obs$Generation <- sapply(
  strsplit(as.character(df.stats.obs$Statistic), split = ".", fixed = TRUE),
  function(x) x[2])

p <- ggplot(data = df.stats.sim, aes(x = Frequency, y = after_stat(density))) +
      geom_histogram(bins = 50) +
      geom_vline(data = df.stats.obs, aes(xintercept = Frequency), color = "orange") +
      facet_grid(Room~Generation) +
      theme_pubclean()
p
```

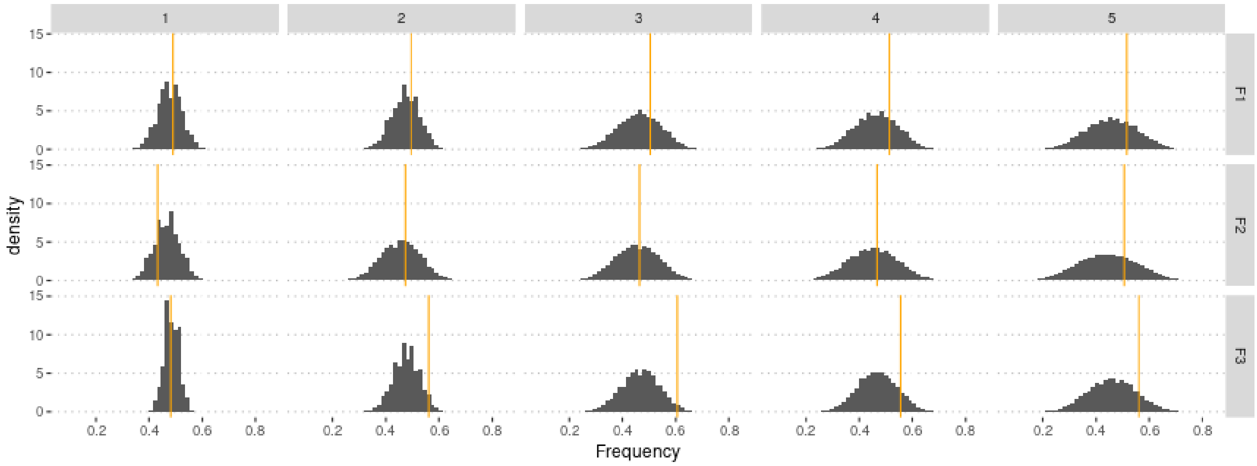


Similar to the sex-averaged model.

### 4 Using genotype frequencies

We estimate parameters using the ridge regression method:

```
if (redo) {
  m.r.g <- abc(target = stat.obs, param = param.sim, sumstat = stat.sim, tol = .1,
             method = "ridge", transf = c("none", "none", "none", "none"))
  save(m.r.g, file = "Rdata/SexSpecific/backup_abc_genotype.Rdata")
} else {
  load("Rdata/SexSpecific/backup_abc_genotype.Rdata")
}
```

Now we display the results:

```
summary(m.r.g, intvl = .95)
```

```
## Call:
## abc(target = stat.obs, param = param.sim, sumstat = stat.sim,
##     tol = 0.1, method = "ridge", transf = c("none", "none", "none",
##         "none"))
## Data:
##  abc.out$adj.values (40000 posterior samples)
## Weights:
##  abc.out$weights
##
##                              hf       sf       hm       sm
```

```
## Min.:                      -0.0146 -0.5083 -0.1095 -0.5301
## Weighted 2.5 % Perc.:       0.0722 -0.2114  0.0070 -0.1903
## Weighted Median:            0.5088  0.0503  0.5205  0.0864
## Weighted Mean:              0.5124  0.1069  0.5221  0.1452
## Weighted Mode:              0.3508 -0.0872  0.3641 -0.0573
## Weighted 97.5 % Perc.:      0.9596  0.6785  1.0520  0.7346
## Max.:                       1.0411  1.8201  1.1571  2.4002
```

Distribution of parameter estimates:

```
hist(m.r.g, breaks = 30, caption = c(expression(h), expression(s)))
```

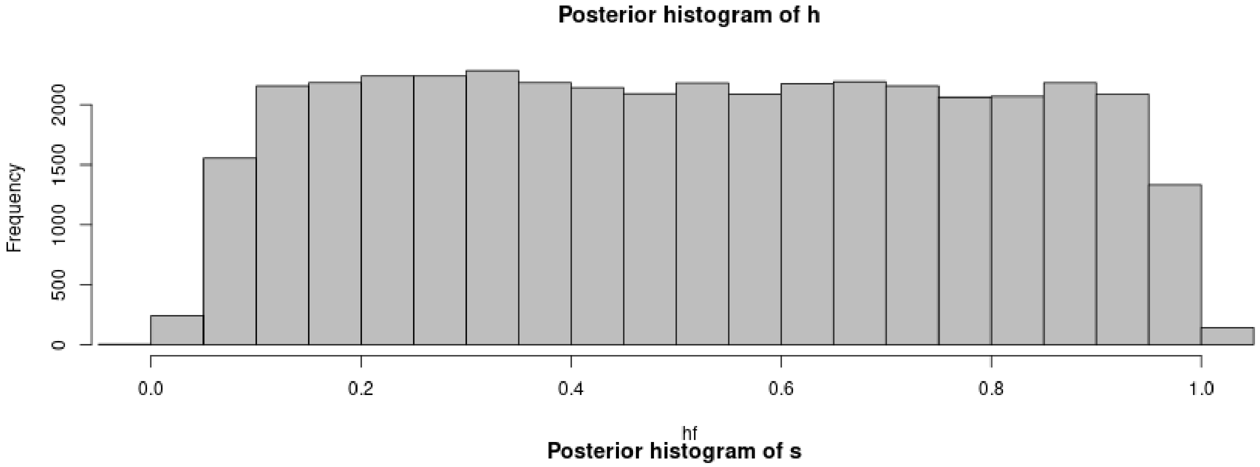


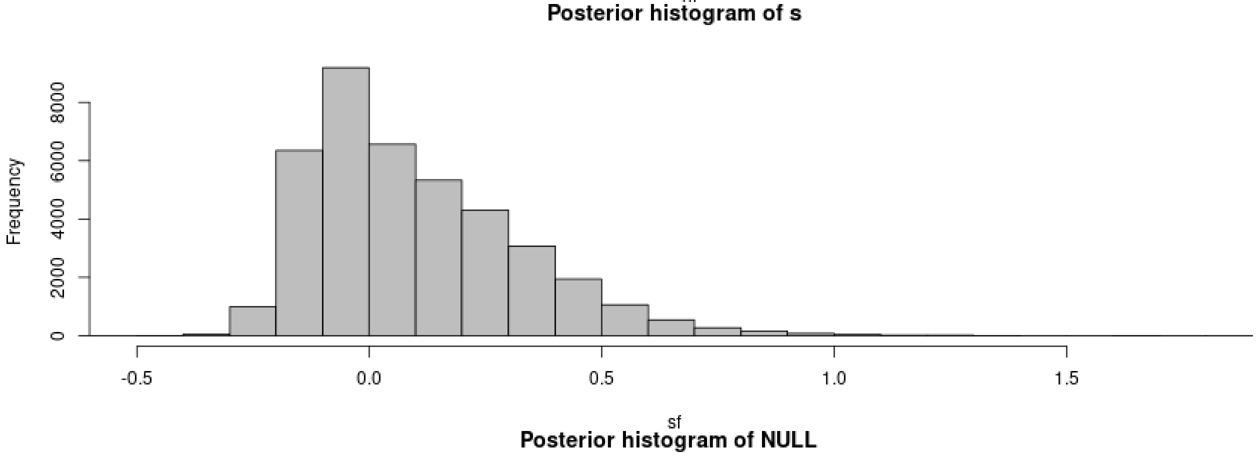


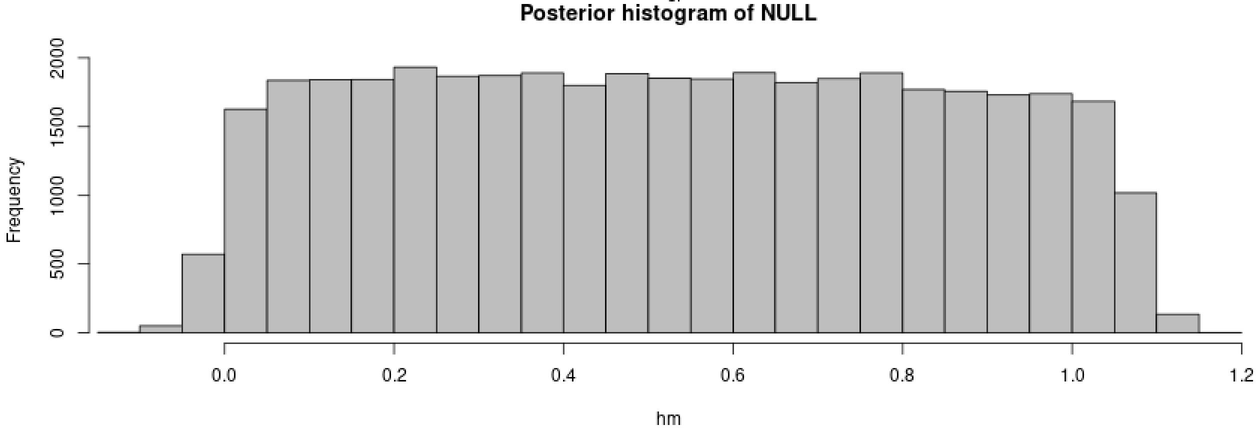

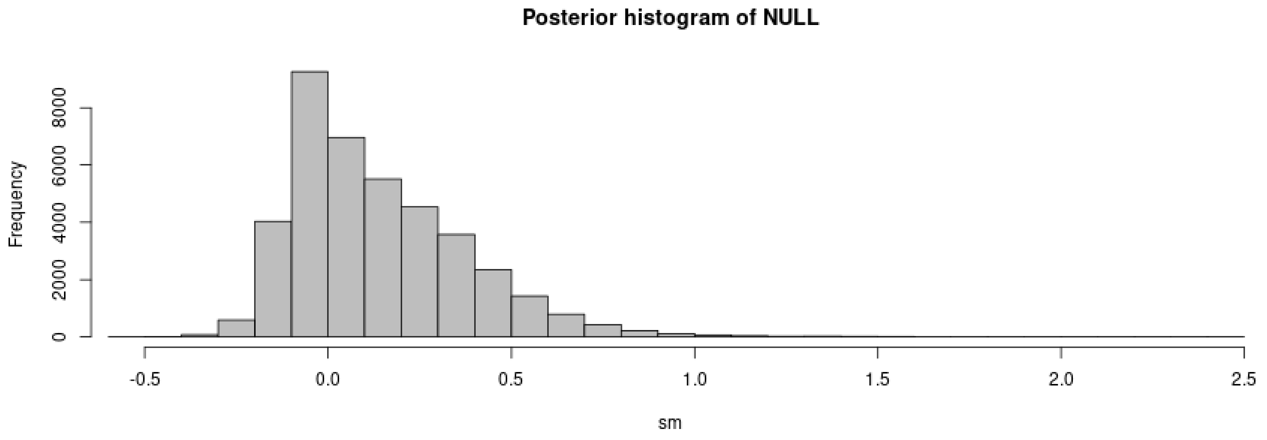


```
plot(m.r.g, param = param.sim)
```

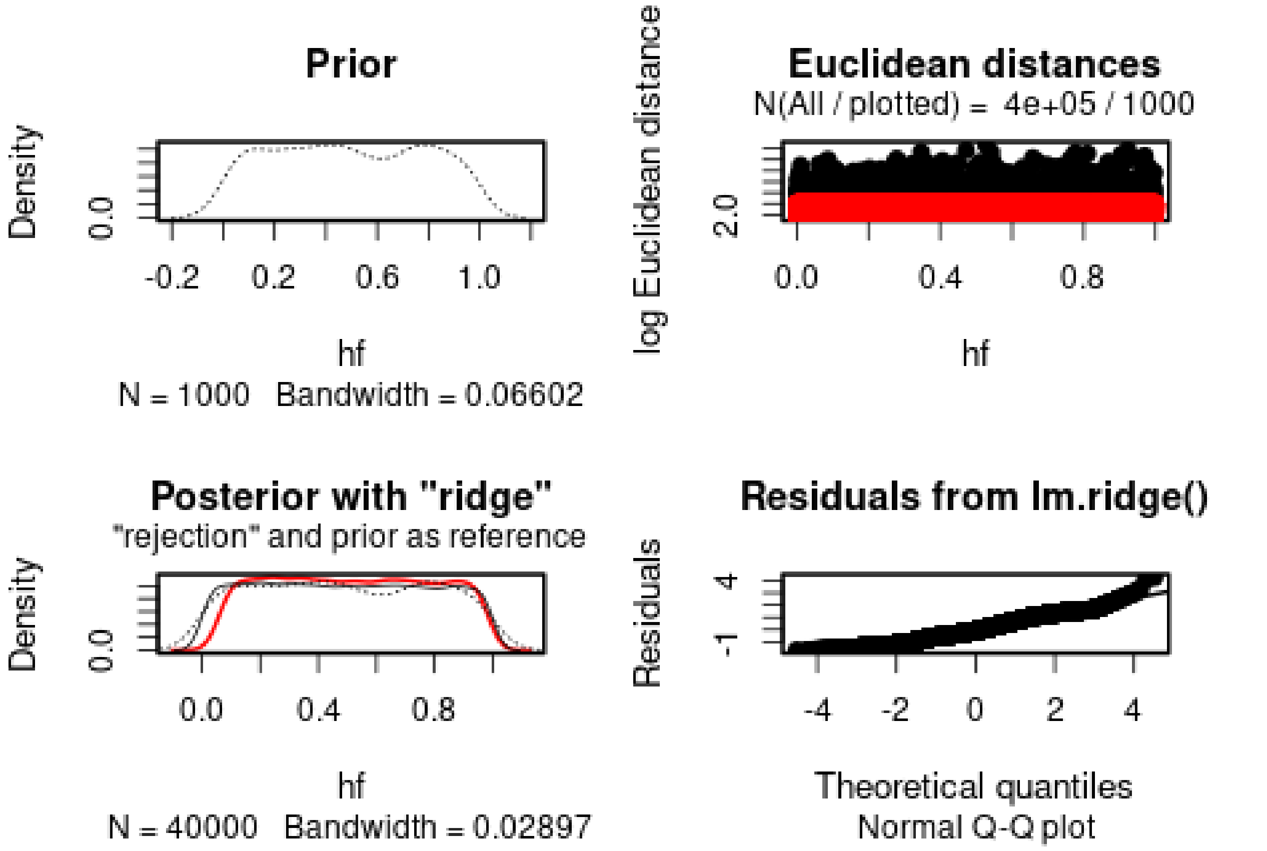

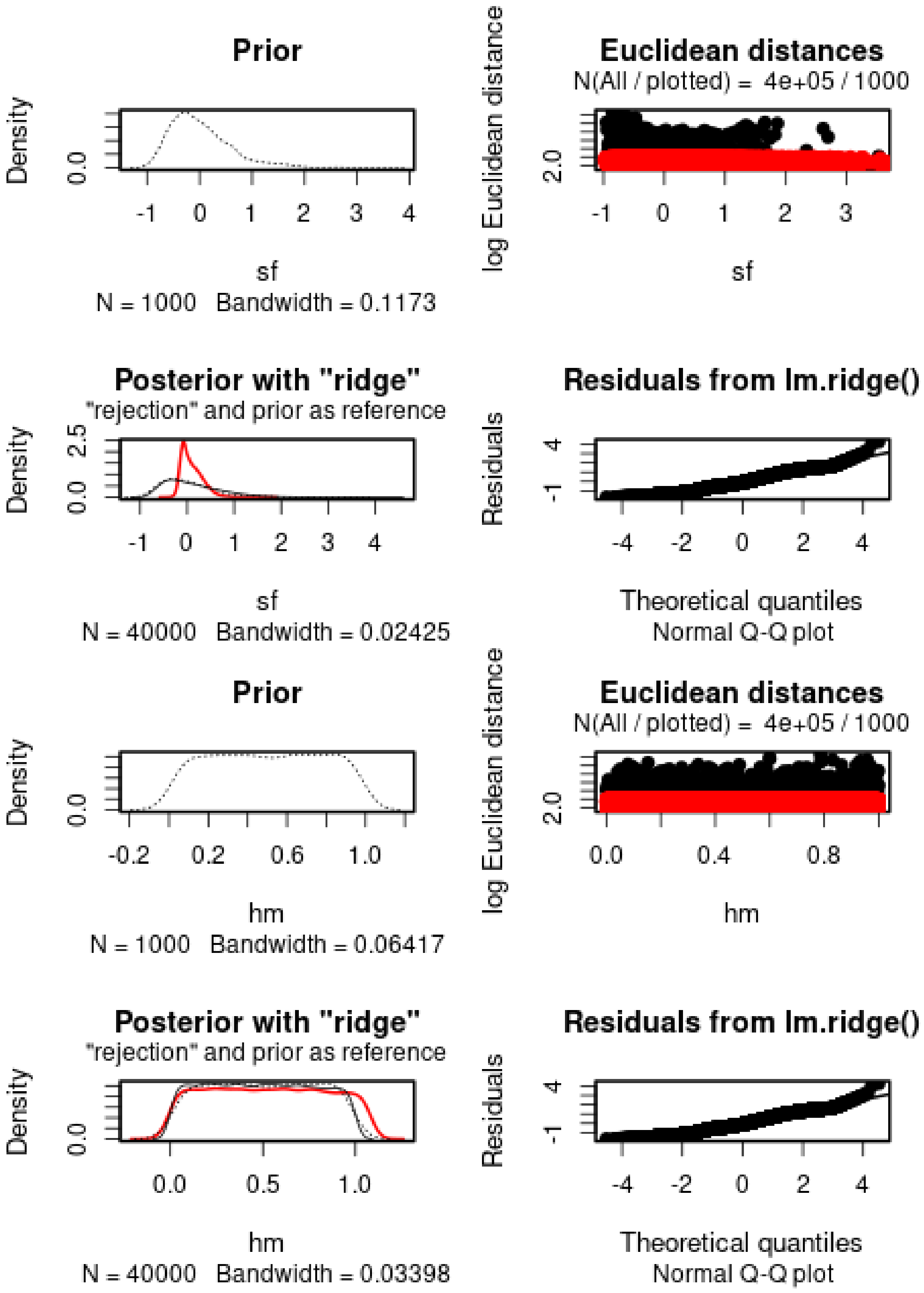
Prior
Density
0.0
-1 0 1 2 3 4
sf
N = 1000 Bandwidth = 0.1173
Euclidean distances
N(All / plotted) = 4e+05 / 1000
log Euclidean distance
2.0
-1 0 1 2 3
sf
Posterior with "ridge"
"rejection" and prior as reference
Density
0.0 2.5
-1 0 1 2 3 4
sf
N = 40000 Bandwidth = 0.02425
Residuals from lm.ridge()
Residuals
-1 4
-4 -2 0 2 4
Theoretical quantiles
Normal Q-Q plot
Prior
Density
0.0
-0.2 0.2 0.6 1.0
hm
N = 1000 Bandwidth = 0.06417
Euclidean distances
N(All / plotted) = 4e+05 / 1000
log Euclidean distance
2.0
0.0 0.4 0.8
hm
Posterior with "ridge"
"rejection" and prior as reference
Density
0.0
0.0 0.5 1.0
hm
N = 40000 Bandwidth = 0.03398
Residuals from lm.ridge()
Residuals
-1 4
-4 -2 0 2 4
Theoretical quantiles
Normal Q-Q plot

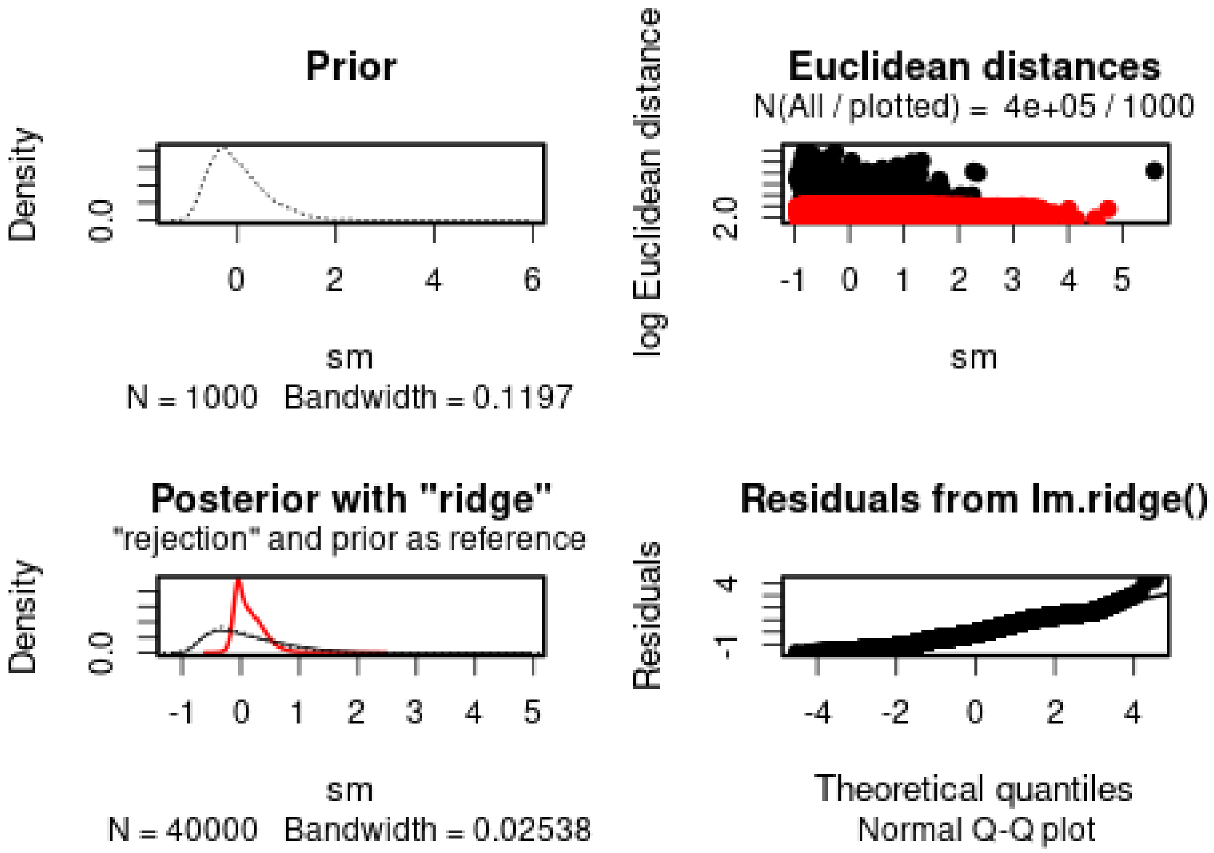


Similar to the model based on allelic frequencies, two *s* coefficients very similar and with large variance.

We then check the distribution of summary statistics in the simulations:

```r
library(reshape2)
library(ggplot2)
library(ggpubr)

df.stats.sim <- melt(stat.sim, variable.name = "Statistic", value.name = "Frequency")
```

```
## No id variables; using all as measure variables
```

```r
l <- strsplit(as.character(df.stats.sim$Statistic), split = ".", fixed = TRUE)
df.stats.sim$Room <- sapply(l, function(x) x[1])
df.stats.sim$Genotype <- sapply(l, function(x) x[2])
df.stats.sim$Generation <- sapply(
  strsplit(as.character(df.stats.sim$Statistic), split = ".", fixed = TRUE),
  function(x) x[3])

df.stats.obs <- as.data.frame(stat.obs)
names(df.stats.obs) <- "Frequency"
df.stats.obs$Statistic <- row.names(df.stats.obs)
l <- strsplit(as.character(df.stats.obs$Statistic), split = ".", fixed = TRUE)
df.stats.obs$Room <- sapply(l, function(x) x[1])
df.stats.obs$Genotype <- sapply(l, function(x) x[2])
df.stats.obs$Generation <- sapply(
  strsplit(as.character(df.stats.obs$Statistic), split = ".", fixed = TRUE),
  function(x) x[3])
```

```r
plot.obs <- function(gen) {
  p <- ggplot(data = subset(df.stats.sim, Generation == gen),
              aes(x = Frequency, y = ..density..)) +
    geom_histogram(bins = 50) +
    geom_vline(data = subset(df.stats.obs, Generation == gen),
               aes(xintercept = Frequency), color = "orange") +
    facet_grid(Genotype~Room) +
    theme_pubclean()
  return(p)
}
```

```r
plot.obs(1) + ggtitle("Timepoint 4")
```

```
## Warning: The dot-dot notation (`..density..`) was deprecated in ggplot2 3.4.0.
## i Please use `after_stat(density)` instead.
## This warning is displayed once every 8 hours.
## Call `lifecycle::last_lifecycle_warnings()` to see where this warning was
## generated.
```

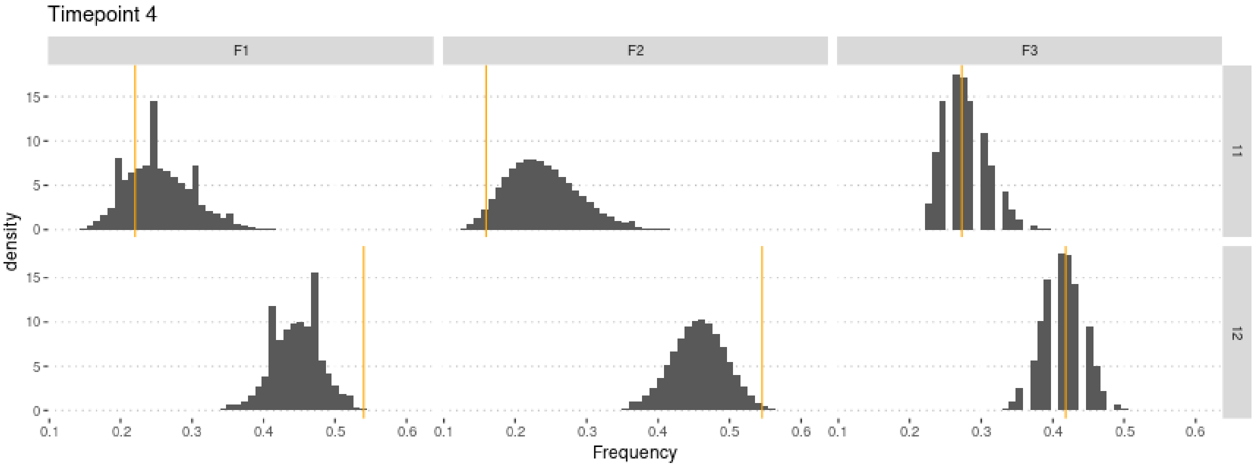


```r
plot.obs(2) + ggtitle("Timepoint 5")
```

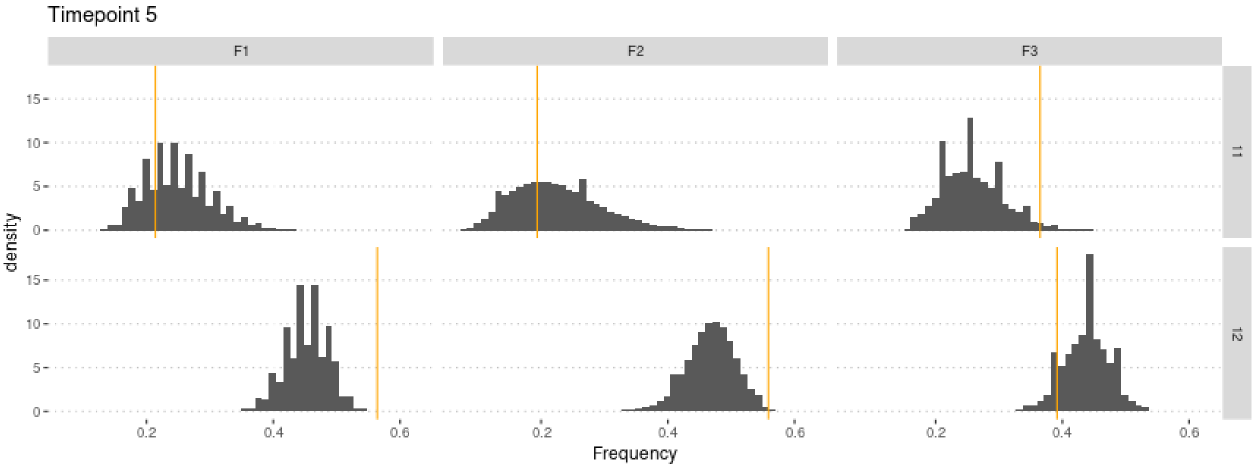

```r
plot.obs(3) + ggtitle("Timepoint 6")
```

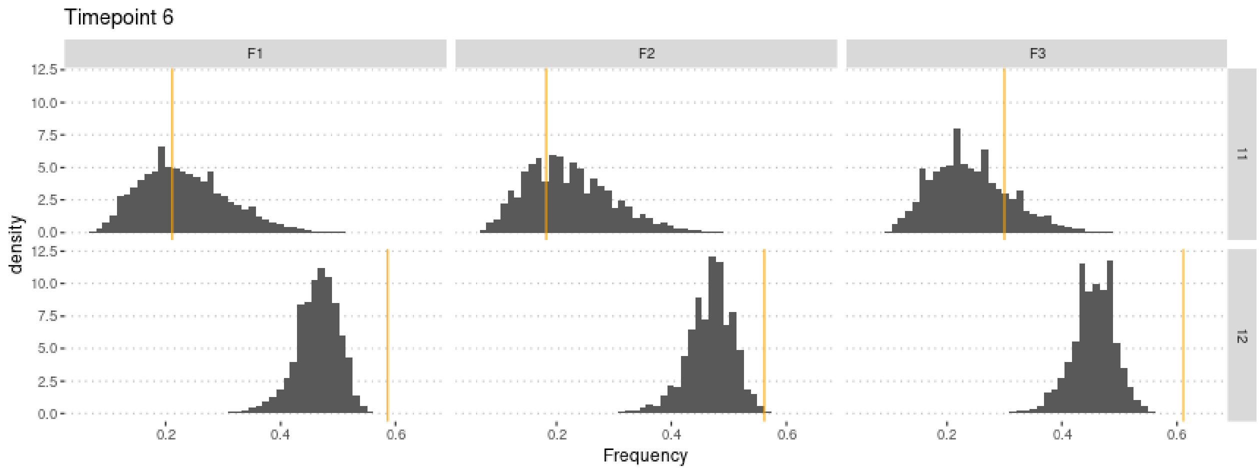


```r
plot.obs(4) + ggtitle("Timepoint 7")
```

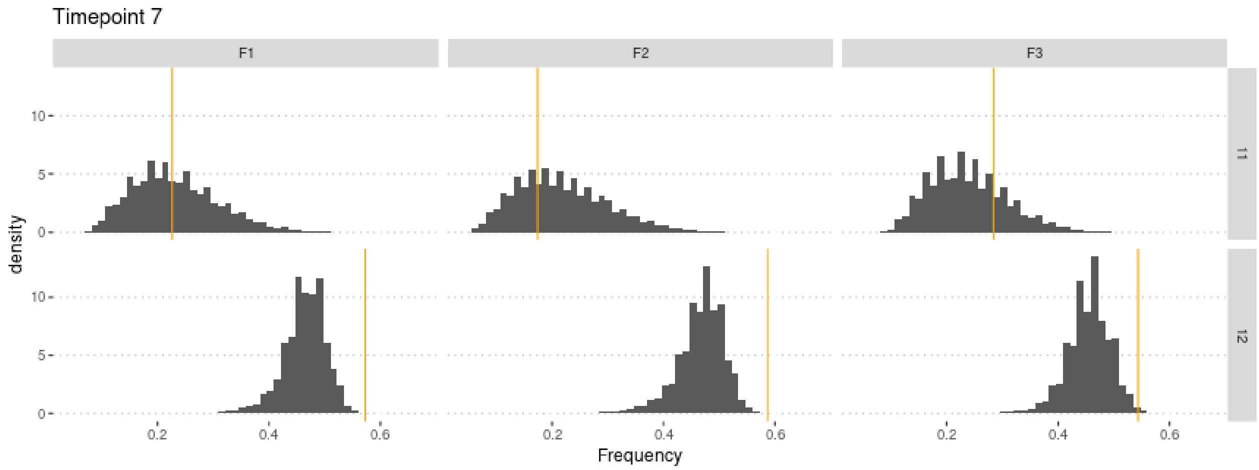


```r
plot.obs(5) + ggtitle("Timepoint 8")
```

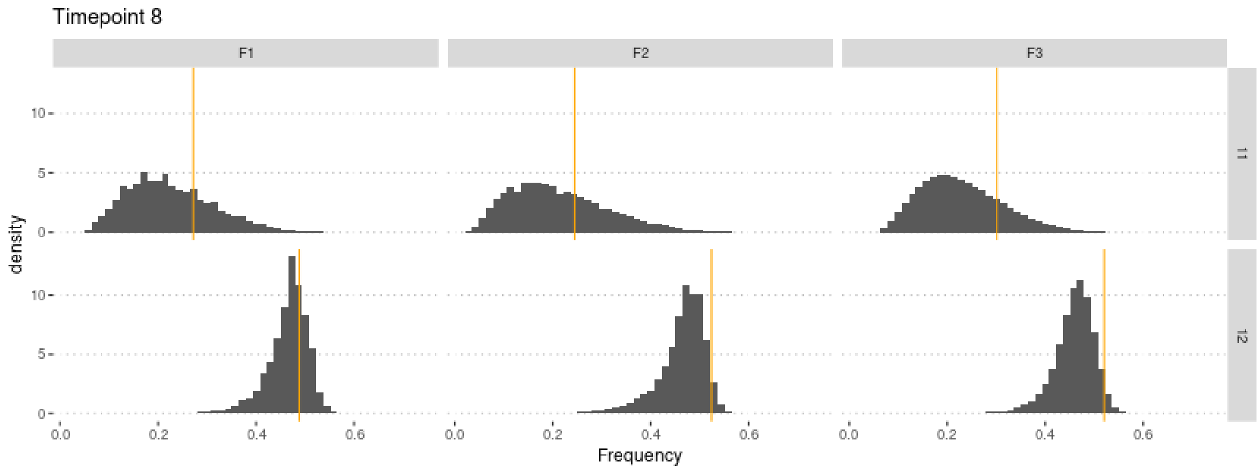
