## Supplementary material for "Experimental evaluation of a direct fitness effect of the *de novo* evolved mouse gene *Pldi*": combined suppl files: SupplementaryFile5.pdf

### Estimating poldi's selection coefficient

Julien Y. Dutheil

06/02/2023

### 1 Model parameterization

The model includes two parameters: $s$ the selection coefficient of the *poldi* allele compared to the knockout strain, which mimics the ancestral state, and $h$ the heterozygosity. Fitness values of each genotype are parameterized as follow:

$$\begin{aligned} poldi/poldi &: 1 \\ BL6/poldi &: 1 + h \cdot s \\ BL6/BL6 &: 1 + s \end{aligned}$$

where BL6 denotes the wild strain, and poldi the strain where the *poldi* gene is knocked out. Furthermore, we consider that the allele only has an effect in males, so that $h$ and $s$ are 0 in females (the two alleles have the same fitness). In this stochastic version, we account for non-genetic variance by modeling the fitness of an individual as a normal distribution with standard deviation $g$, and mean 1, $1 + h.s$, or $1 + s$, depending on its genotype.

We performed 500,000 simulations, using a gamma prior for $s$, a uniform prior between 0 and 1 for $h$, and a uniform prior between 0 and 0.5 for $g$.

#### 2 Preamble

We load the data (observed and simulated):

```
sims <- read.csv("Simulations/SimulationsWithNonGeneticVarianceMalesOnlyFromGen3.csv.gz")
param.sim <- subset(sims, select = c("hm", "sm", "g"))
stat.sim <- subset(sims, select = c(2:46))
# Only consider the WT homozygotes and the heterozygotes,
# as the sum of the three genotypes is 1
stat.sim <- subset(stat.sim, select = grep(".22.", names(stat.sim), invert = TRUE))

library(plyr, quietly = TRUE)
geno <- read.csv("GenotypeFrequencies.csv")
# We select the WT homozygote and heterozygote frequencies.
# Pldi homozygotes are 1 - sum(other two).
stat.obs <- subset(geno, TimePoint %in% 4:8, c("WT", "HET", "Replicate"))
stat.obs <- ddply(stat.obs, "Replicate", function(d) unlist(d[,1:2]))
stat.obs$Replicate <- NULL
stat.obs <- as.data.frame(t(stat.obs))
colnames(stat.obs) <- c("F1", "F2", "F3")
x <- rownames(stat.obs)
x <- gsub(x, pattern = "WT", replacement = ".11.")
x <- gsub(x, pattern = "HET", replacement = ".12.")
#x <- gsub(x, pattern = "Pldi", replacement = ".22.")
rownames(stat.obs) <- x
stat.obs1 <- subset(stat.obs, select = F1)
rownames(stat.obs1) <- paste0("F1", rownames(stat.obs1))
stat.obs2 <- subset(stat.obs, select = F2)
rownames(stat.obs2) <- paste0("F2", rownames(stat.obs2))
stat.obs3 <- subset(stat.obs, select = F3)
rownames(stat.obs3) <- paste0("F3", rownames(stat.obs3))
stat.obs <- cbind(t(stat.obs1), t(stat.obs2), t(stat.obs3))[1,]
```

We estimate parameters using the ridge regression method:

```
library(abc, quietly = TRUE)
```

```
##
## Attaching package: 'SparseM'

## The following object is masked from 'package:base':
##
##     backsolve

## locfit 1.5-9.8    2023-06-11
```

```
if (redo) {
  m.r.a <- abc(target = stat.obs.a, param = param.sim, sumstat = stat.sim.a, tol=.1,
            method = "ridge", transf = c("none", "none", "none"))
  save(m.r.a, file = "Rdata/MalesOnlyNonGeneticVariance/backup_abc_allelic.Rdata")
} else {
  load("Rdata/MalesOnlyNonGeneticVariance/backup_abc_allelic.Rdata")
}
```

Now we display the results:

```
summary(m.r.a, intvl = .95)
```

```
## Call:
## abc(target = stat.obs.a, param = param.sim, sumstat = stat.sim.a,
```

```
##     tol = 0.1, method = "ridge", transf = c("none", "none", "none"))
## Data:
##  abc.out$adj.values (50000 posterior samples)
## Weights:
##  abc.out$weights
##
##                               hm      sm       g
## Min.:                    -0.0261 -0.4756 -0.0558
## Weighted 2.5 % Perc.:     0.0553  0.0883  0.0054
## Weighted Median:          0.5123  0.5919  0.2484
## Weighted Mean:            0.5080  0.6285  0.2489
## Weighted Mode:            0.7039  0.5344  0.0554
## Weighted 97.5 % Perc.:    0.9453  1.3825  0.4928
## Max.:                     0.9977  3.6490  0.5327
```

Distribution of parameter estimates:

```
hist(m.r.a, breaks = 30, caption = c(expression(h), expression(s), expression(g)))
```

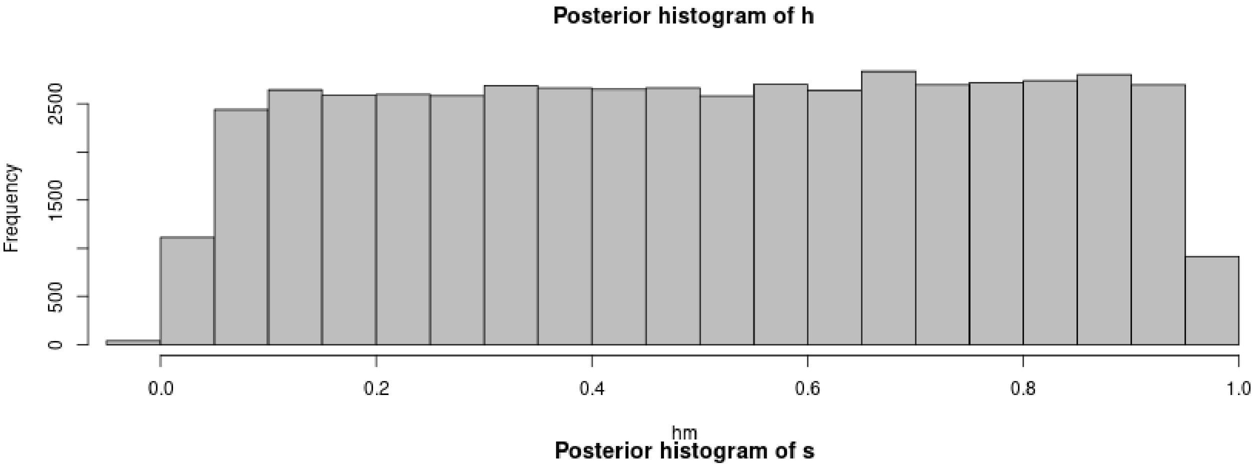

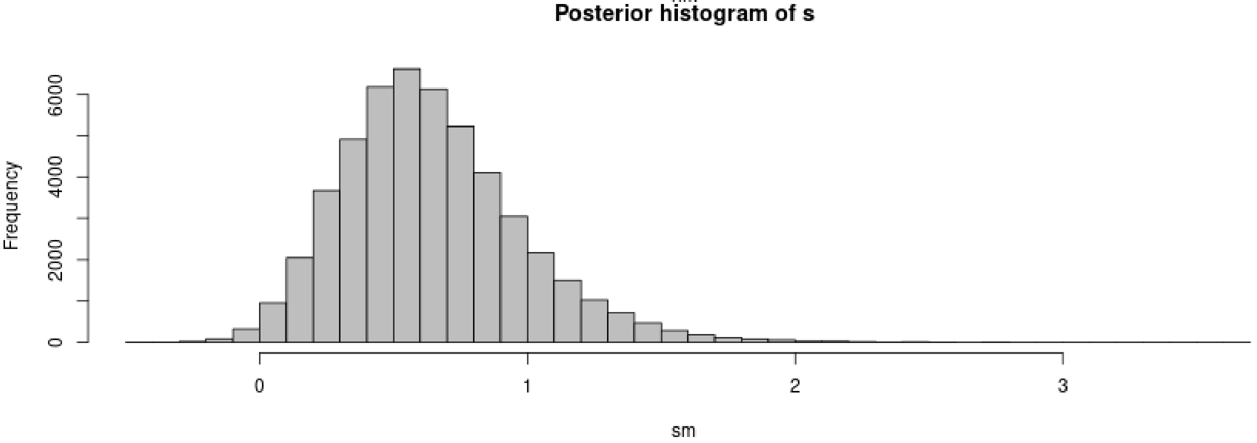

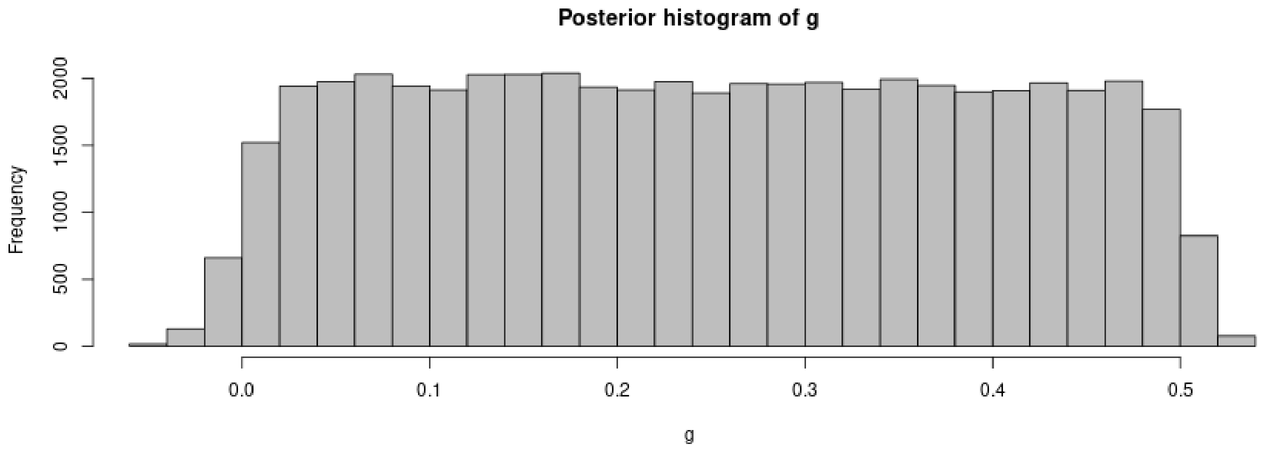

```r
plot(m.r.a, param = param.sim)
```

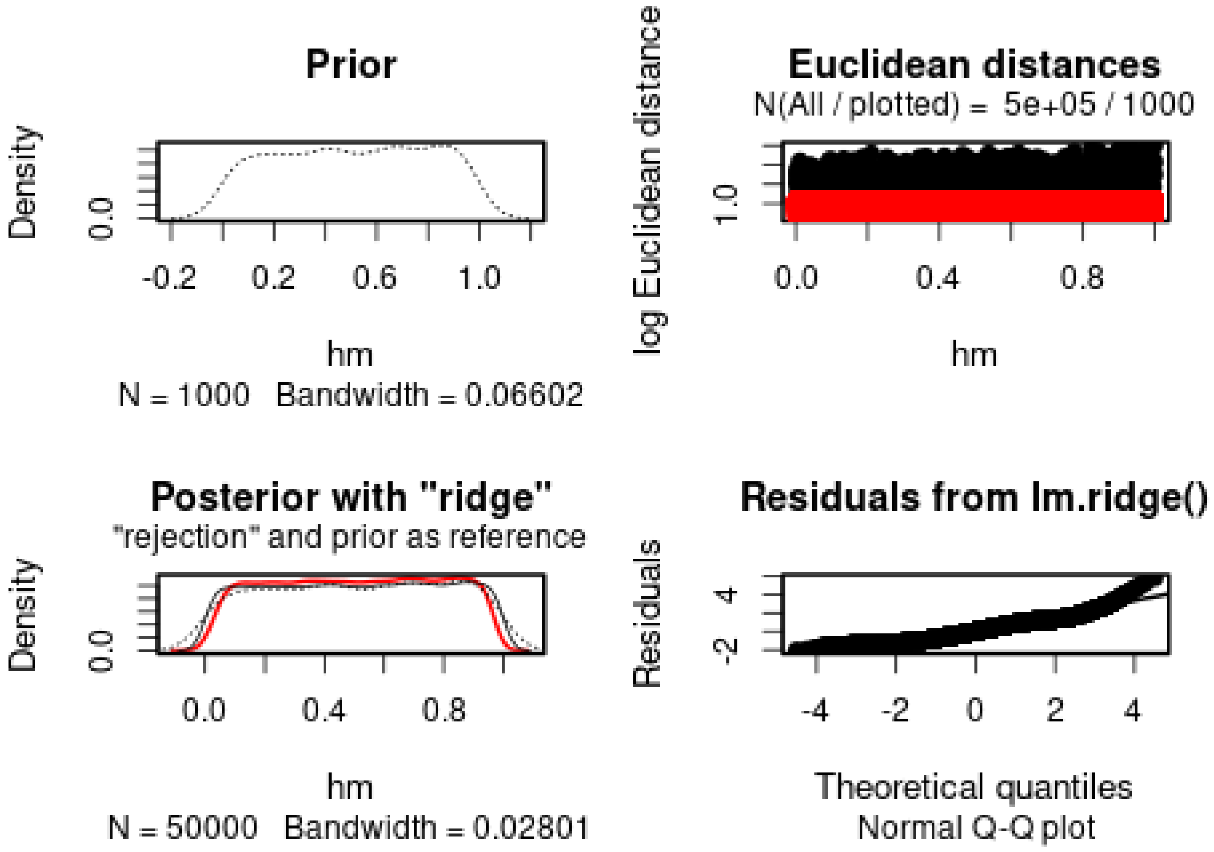

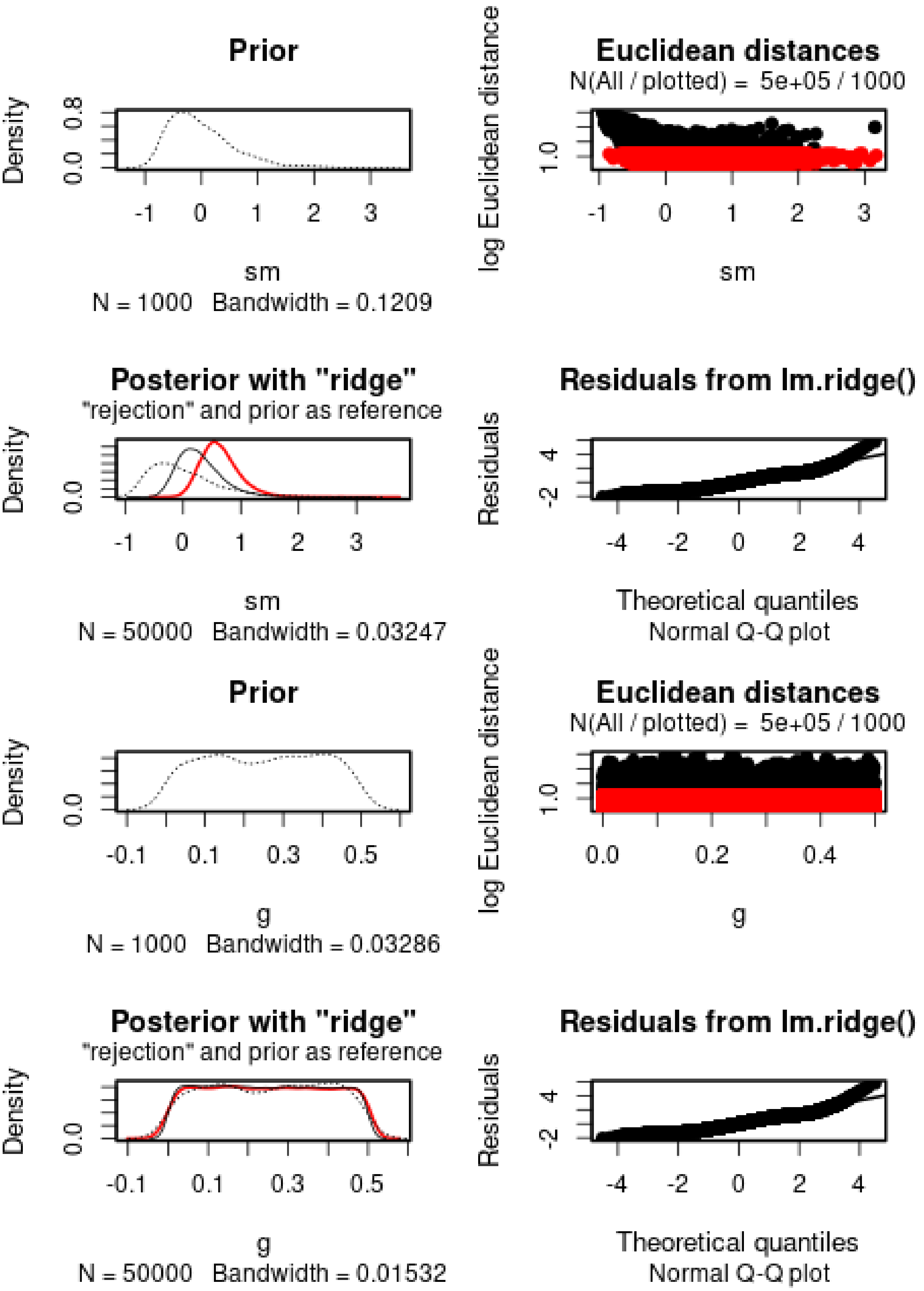

$h$ cannot properly be estimated. $s$ is positive, but has a large variance. The 95% posterior interval does not include 0.

We then check the distribution of summary statistics in the simulations:

```
library(reshape2)
library(ggplot2)
library(ggpubr)
```

```
##
## Attaching package: 'ggpubr'

## The following object is masked from 'package:plyr':
##
##     mutate
```

```
df.stats.sim <- melt(stat.sim.a, variable.name = "Statistic", value.name = "Frequency")
```

```
## No id variables; using all as measure variables
```

```
l <- strsplit(as.character(df.stats.sim$Statistic), split = ".", fixed = TRUE)
df.stats.sim$Room <- sapply(l, function(x) x[1])
df.stats.sim$Generation <- sapply(
  strsplit(as.character(df.stats.sim$Statistic), split = ".", fixed = TRUE),
  function(x) x[2])

df.stats.obs <- as.data.frame(stat.obs.a)
names(df.stats.obs) <- "Frequency"
df.stats.obs$Statistic <- row.names(df.stats.obs)
l <- strsplit(as.character(df.stats.obs$Statistic), split = ".", fixed = TRUE)
df.stats.obs$Room <- sapply(l, function(x) x[1])
df.stats.obs$Generation <- sapply(
  strsplit(as.character(df.stats.obs$Statistic), split = ".", fixed = TRUE),
  function(x) x[2])

p <- ggplot(data = df.stats.sim, aes(x = Frequency, y = after_stat(density))) +
      geom_histogram(bins = 50) +
      geom_vline(data = df.stats.obs, aes(xintercept = Frequency), color = "orange") +
      facet_grid(Room~Generation) +
      theme_pubclean()
p
```

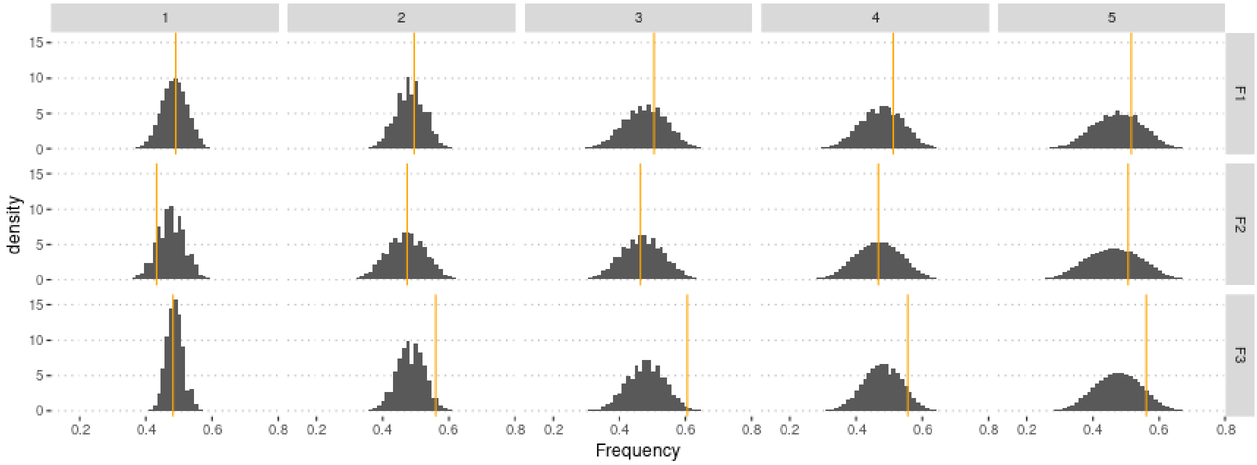

#### 3.1 Cross-Validation analysis

Compute predictions errors. For memory-saving reasons, we only use 100,000 simulations in the following.

```
sims2 <- sims[1:1e5,] #Not enough memory to use all simulations
param.sim2 <- param.sim[1:1e5,]
stat.sim.a2 <- stat.sim.a[1:1e5,]
m.r.a2 <- abc(target = stat.obs.a, param = param.sim2, sumstat = stat.sim.a2, tol = .1,
              method = "ridge", transf = c("none", "none", "none"))
```

Check that this does not affect the estimates too much:

```
summary(m.r.a2, intvl = .95)
```

```
## Call:
## abc(target = stat.obs.a, param = param.sim2, sumstat = stat.sim.a2,
##     tol = 0.1, method = "ridge", transf = c("none", "none", "none"))
## Data:
##  abc.out$adj.values (10000 posterior samples)
## Weights:
##  abc.out$weights
##
##                              hm      sm       g
## Min.:                    0.0135 -0.4866 -0.0472
## Weighted 2.5 % Perc.:    0.1148  0.0350  0.0082
## Weighted Median:         0.5024  0.5565  0.2396
## Weighted Mean:           0.5038  0.5973  0.2412
## Weighted Mode:           0.3494  0.4821  0.0602
## Weighted 97.5 % Perc.:   0.8866  1.3872  0.4808
## Max.:                    0.9499  3.5982  0.5202
```

Similar estimate, but slightly lower variance/

```
if (redo) {
  cv.ridge.a <- cv4abc(param = param.sim2,
                       sumstat = stat.sim.a2,
                       abc.out = m.r.a2,
                       nval = 1000,
                       tols = c(.01, .1, .2))
  save(cv.ridge.a, file = "Rdata/MalesOnlyNonGeneticVariance/backup_cv_allelic.Rdata")
} else {
  load("Rdata/MalesOnlyNonGeneticVariance/backup_cv_allelic.Rdata")
}
summary(cv.ridge.a)
```

```
## Prediction error based on a cross-validation sample of 1000

##             hm        sm         g
## 0.01 0.9436670 0.1774027 1.0236665
## 0.1  0.9273673 0.1763113 0.9887152
## 0.2  0.9281779 0.1789099 0.9874353
```

```
plot(cv.ridge.a)
```

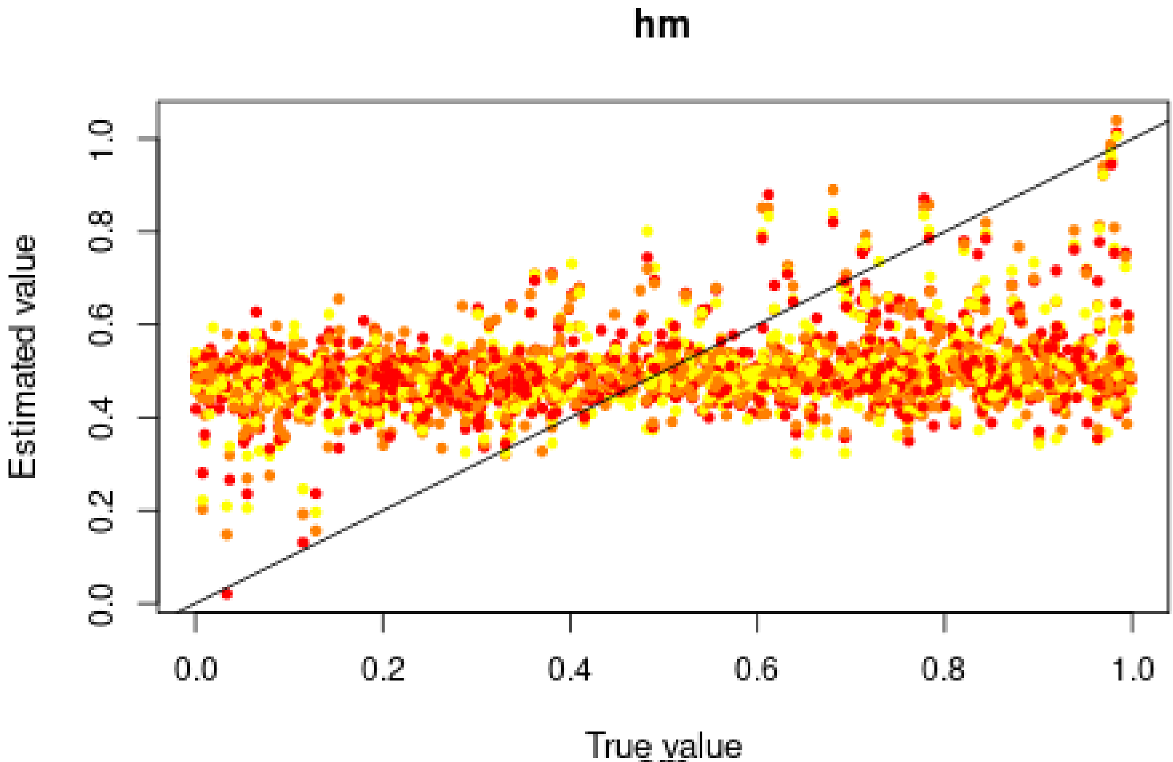
hm
Estimated value
0.0
0.2
0.4
0.6
0.8
1.0
True value

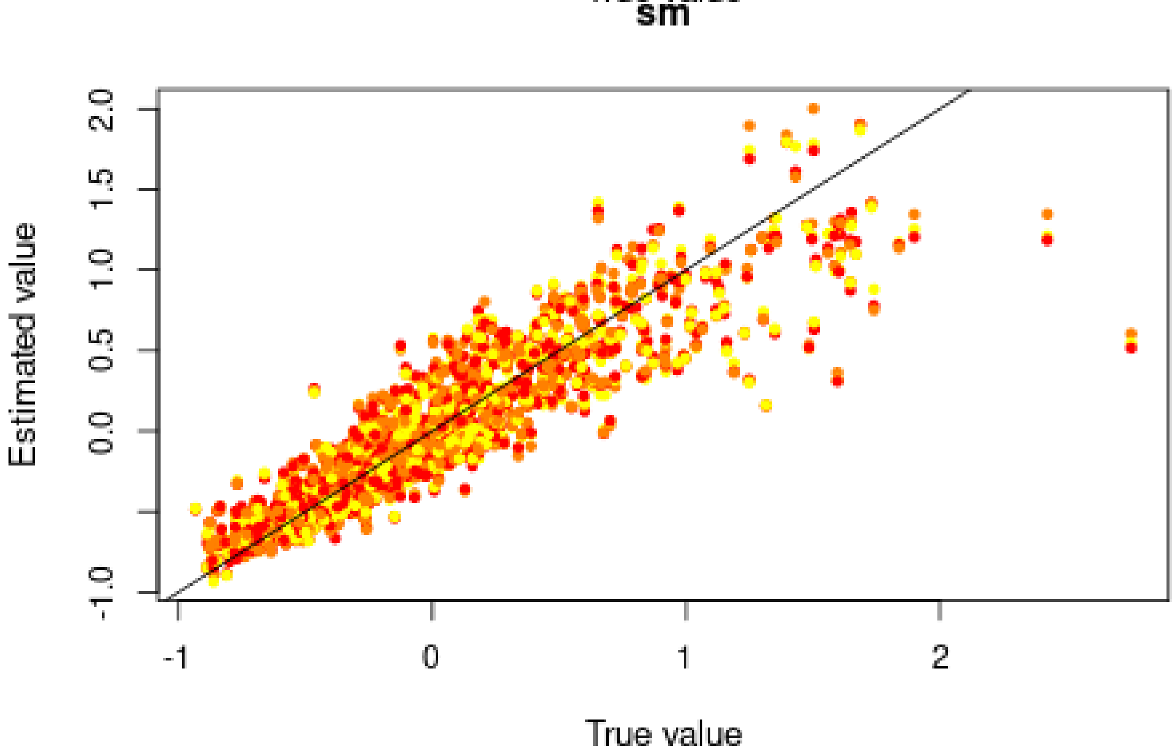
sm
Estimated value
-1.0
0.0
0.5
1.0
1.5
2.0
-1
0
1
2
True value

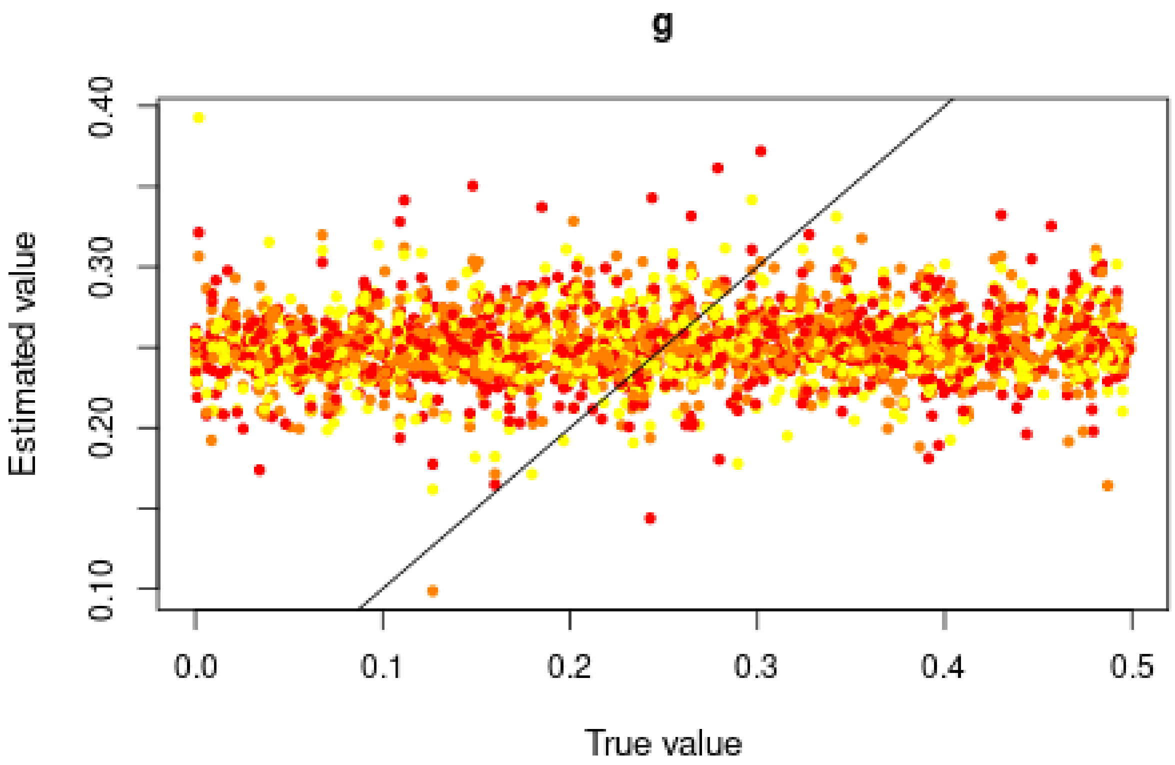

Good for $s$, but very bad for $h$, in agreement with the posterior distributions.

##### 3.2 Misclassification errors

To test the power of the approach to distinguish between models, we also conduct a cross-validation experiment. We compare model M0 ($s = 0$) and M1 ($s = 0.6151 > 0$). $h$ is sampled over its prior distribution. 100,000 simulations were conducted under each model.

```
sims0 <- read.csv("Simulations/Simulations0WithNonGeneticVarianceFromGen3.csv.gz")
param.sim0 <- subset(sims0, select = c("hm", "sm", "g"))
stat.sim0 <- subset(sims0, select = 2:46)
stat.sim0.a <- compute.allelic.frequencies(stat.sim0)

sims1 <- read.csv("Simulations/Simulations1aWithNonGeneticVarianceMalesOnlyFromGen3.csv.gz")
param.sim1 <- subset(sims1, select = c("hm", "sm", "g"))
stat.sim1 <- subset(sims1, select = 2:46)
stat.sim1.a <- compute.allelic.frequencies(stat.sim1)
```

We conduct a CV analysis (this takes some time... set nval = 100 for faster results):

```
models <- rep(c("Neutral", "Selection"), each = 100000)
if (redo) {
  cv.modsel.a <- cv4postpr(models, rbind(stat.sim0.a, stat.sim1.a),
                           nval = 1000, tols = .1, method = "mnlogistic")
  save(cv.modsel.a, file = "Rdata/MalesOnlyNonGeneticVariance/backup_cv4postpr_allelic.Rdata")
} else {
  load("Rdata/MalesOnlyNonGeneticVariance/backup_cv4postpr_allelic.Rdata")
}
```

We display the results:

```
summary(cv.modsel.a)
```

```
## Confusion matrix based on 1000 samples for each model.
##
## $tol0.1
##           Neutral Selection
## Neutral       897       103
## Selection     108       892
##
##
## Mean model posterior probabilities (mnlogistic)
##
## $tol0.1
##           Neutral Selection
## Neutral    0.8476    0.1524
## Selection  0.1567    0.8433
```

```
plot(cv.modsel.a, names.arg = c("Neutral", "Selection"))
```

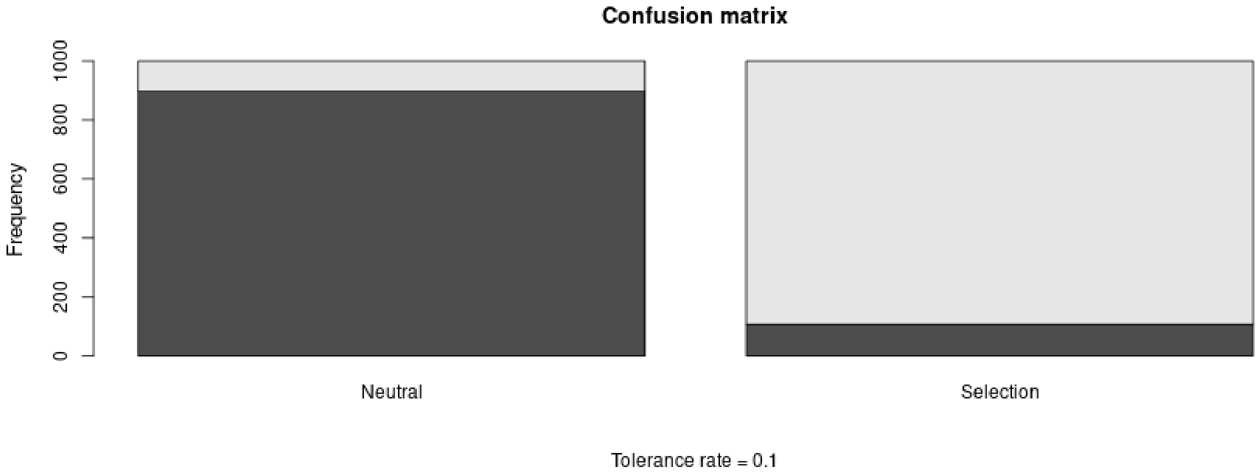

##### 3.3 Posterior prediction

```
if (redo) {
  modsel.a <- postpr(stat.obs.a, models, rbind(stat.sim0.a, stat.sim1.a),
                     tol = .1, method = "mnlogistic")
  save(modsel.a, file = "Rdata/MalesOnlyNonGeneticVariance/backup_postpr_allelic.Rdata")
} else {
  load("Rdata/MalesOnlyNonGeneticVariance/backup_postpr_allelic.Rdata")
}
summary(modsel.a)
```

```
## Call:
## postpr(target = stat.obs.a, index = models, sumstat = rbind(stat.sim0.a,
##     stat.sim1.a), tol = 0.1, method = "mnlogistic")
## Data:
##  postpr.out$values (20000 posterior samples)
## Models a priori:
##  Neutral, Selection
## Models a posteriori:
##  Neutral, Selection
```

```
##
## Proportion of accepted simulations (rejection):
##   Neutral Selection
##    0.4186    0.5813
##
## Bayes factors:
##           Neutral Selection
## Neutral    1.0000    0.7201
## Selection  1.3886    1.0000
##
##
## Posterior model probabilities (mnlogistic):
##   Neutral Selection
##    0.0691    0.9309
##
## Bayes factors:
##           Neutral Selection
## Neutral    1.0000    0.0742
## Selection 13.4811    1.0000
```

The model with selection is preferred.

##### 3.4 Goodness of fit

Under the neutral model:

```
if (redo) {
  res.gfit0.a <- gfit(target = stat.obs.a, sumstat = stat.sim0.a,
                      statistic = median, nb.replicate = 1000)
  save(res.gfit0.a, file = "Rdata/MalesOnlyNonGeneticVariance/backup_gfit0_allelic.Rdata")
} else {
  load("Rdata/MalesOnlyNonGeneticVariance/backup_gfit0_allelic.Rdata")
}
plot(res.gfit0.a, main = "Histogram under M0")
```

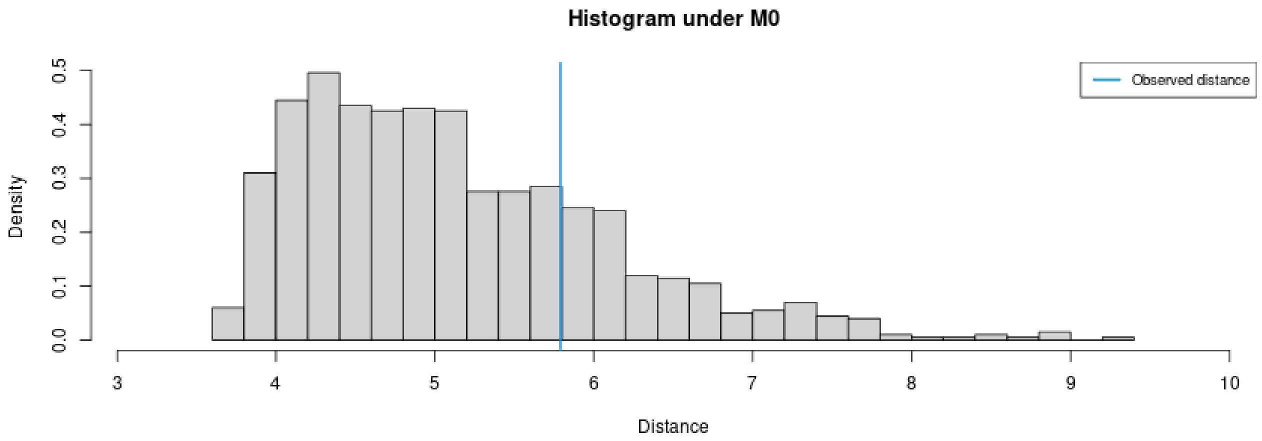

```
summary(res.gfit0.a)
```

```
## $pvalue
## [1] 0.233
##
## $s.dist.sim
##    Min. 1st Qu.  Median    Mean 3rd Qu.    Max.
```

```
##    3.675    4.363    4.930    5.134    5.729    9.333
##
## $dist.obs
## [1] 5.789998
```

Under the selection model:

```
if (redo) {
  res.gfit1.a <- gfit(target = stat.obs.a, sumstat = stat.sim1.a,
                      statistic = median, nb.replicate = 1000)
  save(res.gfit1.a, file = "Rdata/MalesOnlyNonGeneticVariance/backup_gfit1_allelic.Rdata")
} else {
  load("Rdata/MalesOnlyNonGeneticVariance/backup_gfit1_allelic.Rdata")
}
plot(res.gfit1.a, main = "Histogram under M1")
```

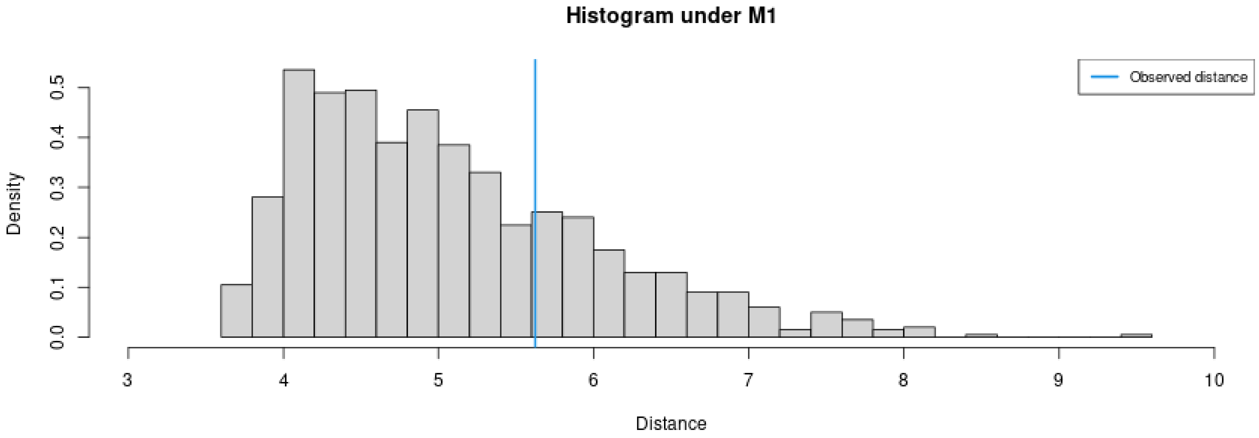

```
summary(res.gfit1.a)
```

```
## $pvalue
## [1] 0.251
##
## $s.dist.sim
##     Min. 1st Qu.  Median    Mean 3rd Qu.    Max.
##    3.607    4.341    4.882    5.073    5.624    9.402
##
## $dist.obs
## [1] 5.623276
```

Both models provide a reasonably good fit.

#### 3.5 Summary figure

Posterior distributions, with prior for comparison:

```
library(ggplot2)
library(ggpubr)
dat.prior.h <- data.frame(h = param.sim[,"hm"], Distribution = "Prior")
dat.post.h <- data.frame(h = m.r.a$adj.values[,"hm"], Distribution = "Posterior")
dat.prior.s <- data.frame(s = param.sim[,"sm"], Distribution = "Prior")
dat.post.s <- data.frame(s = m.r.a$adj.values[,"sm"], Distribution = "Posterior")
dat.prior.g <- data.frame(s = param.sim[,"g"], Distribution = "Prior")
dat.post.g <- data.frame(s = m.r.a$adj.values[,"g"], Distribution = "Posterior")
dat.h <- rbind(dat.prior.h, dat.post.h)
```

```
dat.s <- rbind(dat.prior.s, dat.post.s)
dat.g <- rbind(dat.prior.g, dat.post.g)
dat.h$Distribution <- factor(dat.h$Distribution, levels = c("Prior", "Posterior"))
dat.s$Distribution <- factor(dat.s$Distribution, levels = c("Prior", "Posterior"))
dat.g$Distribution <- factor(dat.g$Distribution, levels = c("Prior", "Posterior"))
names(dat.h)[1] <- "value"
dat.h$variable <- "h"
names(dat.s)[1] <- "value"
dat.s$variable <- "s"
names(dat.g)[1] <- "value"
dat.g$variable <- "g"
dat.dist <- rbind(dat.h, dat.s, dat.g)
dat.dist$variable <- factor(dat.dist$variable, levels = c("h", "s", "g"))

p.dist <- ggplot(data = dat.dist,
                 aes(x = value, linetype = Distribution, fill = Distribution)) +
          geom_density(alpha = 0.5) +
          scale_fill_brewer(type = "qual", palette = 3) +
          scale_linetype_manual(values = c(Prior = "dashed", Posterior = "solid")) +
          xlab("Parameter value") +
          facet_wrap(~variable, scales = "free") +
          ggtitle("Posterior and prior parameter distributions") +
          theme_pubclean() + theme(strip.background = element_blank())
```

Cross-validation:

```
d1<-rbind(as.data.frame(cv.ridge.a$estim$tol0.01),
          as.data.frame(cv.ridge.a$estim$tol0.1),
          as.data.frame(cv.ridge.a$estim$tol0.2))
d2<-rbind(cv.ridge.a$true, cv.ridge.a$true, cv.ridge.a$true)
d1<-melt(d1, value.name = "Estimated")
```

```
## No id variables; using all as measure variables
```

```
d1$Tolerance <- rep(c(0.01, 0.1, 0.2), each = 1000)
d1$Replicate <- rep(1:1000, 3)
d2<-melt(d2, value.name = "True")
```

```
## No id variables; using all as measure variables
```

```
d2$Replicate <- rep(1:1000, 3)
dat.cv <- merge(d1, d2, by = c("variable", "Replicate"))
dat.cv$variable <- substr(dat.cv$variable, 1, 1)
dat.cv$variable <- factor(dat.cv$variable, levels = c("h", "s", "g"))
```

```
p.cv <- ggplot(dat.cv, aes(x = True, y = Estimated)) +
        geom_point(aes(col = as.ordered(Tolerance))) +
        geom_abline(slope = 1) +
        facet_wrap(~variable, scales = "free") +
        theme_pubclean() + labs(color = "Tolerance") +
        ggtitle("Cross validation") +
        theme(strip.background = element_blank())
```

Confusion matrix:

```
library(scales)
cv.sum <- summary(cv.modsel.a)
```

```
## Confusion matrix based on 1000 samples for each model.
##
## $tol0.1
##           Neutral Selection
## Neutral       897       103
## Selection     108       892
##
##
## Mean model posterior probabilities (mnlogistic)
##
## $tol0.1
##           Neutral Selection
## Neutral    0.8476    0.1524
## Selection  0.1567    0.8433
```

```
dat.cv <- as.data.frame(cv.sum$conf.matrix$tol0.1/1000)
names(dat.cv) <- c("Real", "Inferred", "Frequency")
p.confmat <- ggplot(dat.cv, aes(x = Real, y = Frequency, fill = Inferred)) +
             geom_col() +
             scale_y_continuous(labels = scales::percent) +
             scale_fill_brewer(type = "qual", palette = 3) +
             ggtitle("Prediction errors") +
             theme_pubclean()
```

Model probabilities:

```
p.mprob <- ggplot(as.data.frame(modsel.a$pred), aes(x = Var1, y = Freq, fill = Var1)) +
           geom_col() + ylab("Model posterior probability") + xlab("Model") +
           scale_fill_brewer(type = "qual", palette = 3) +
           theme_pubclean() +
           ggtitle("Model probabilities") +
           theme(legend.position = "none")
```

```
library(cowplot)
```

```
##
## Attaching package: 'cowplot'

## The following object is masked from 'package:ggpubr':
##
##     get_legend
```

```
p1 <- plot_grid(p.mprob, p.confmat, labels = c("C", "D"), nrow = 1)
p <- plot_grid(p.dist, p.cv, p1, labels = c("A", "B", ""), nrow = 3)
p
```

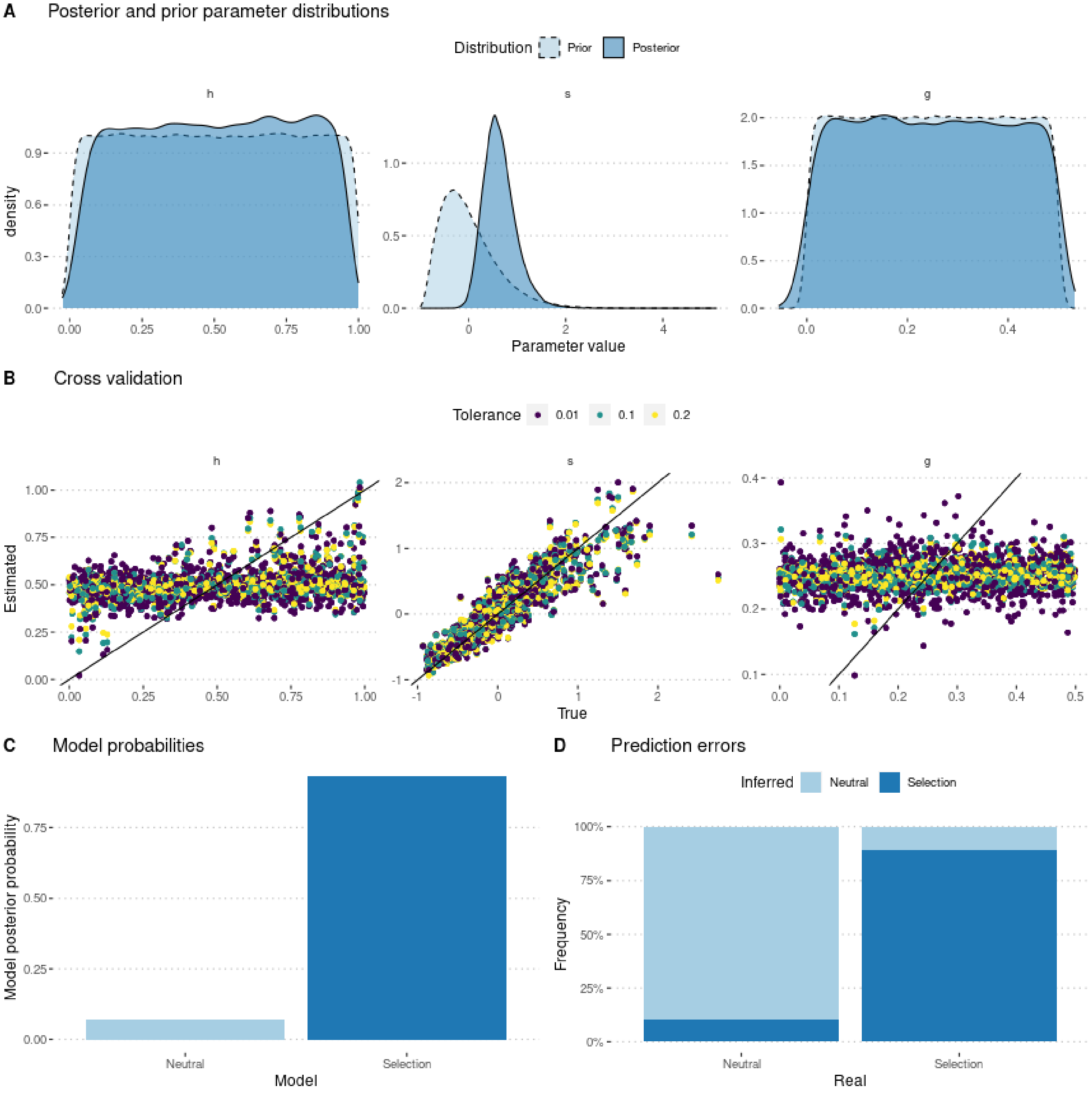

```
ggsave(p, filename = "FigureABC-MalesOnlyNonGeneticVariance-Allelic.pdf", width = 8, height = 10)
```

### 4 Using genotype frequencies

We estimate parameters using the ridge regression method:

```
if (redo) {
  m.r.g <- abc(target = stat.obs, param = param.sim, sumstat = stat.sim, tol = .1,
               method = "ridge", transf = c("none", "none", "none"))
  save(m.r.g, file = "Rdata/MalesOnlyNonGeneticVariance/backup_abc_genotype.Rdata")
} else {
  load("Rdata/MalesOnlyNonGeneticVariance/backup_abc_genotype.Rdata")
}
```

Now we display the results:

```r
summary(m.r.g, intvl = .95)
```

```
## Call:
## abc(target = stat.obs, param = param.sim, sumstat = stat.sim,
##     tol = 0.1, method = "ridge", transf = c("none", "none", "none"))
## Data:
##  abc.out$adj.values (50000 posterior samples)
## Weights:
##  abc.out$weights
##
##                                  hm      sm       g
## Min.:                        0.0563 -0.4098 -0.0077
## Weighted 2.5 % Perc.:        0.1683 -0.0069  0.0456
## Weighted Median:             0.5851  0.3725  0.2962
## Weighted Mean:               0.5883  0.3973  0.2973
## Weighted Mode:               0.4652  0.3230  0.2902
## Weighted 97.5 % Perc.:       1.0160  0.9360  0.5526
## Max.:                        1.1029  2.2988  0.6061
```

Distribution of parameter estimates:

```r
hist(m.r.g, breaks = 30, caption = c(expression(h), expression(s), expression(g)))
```

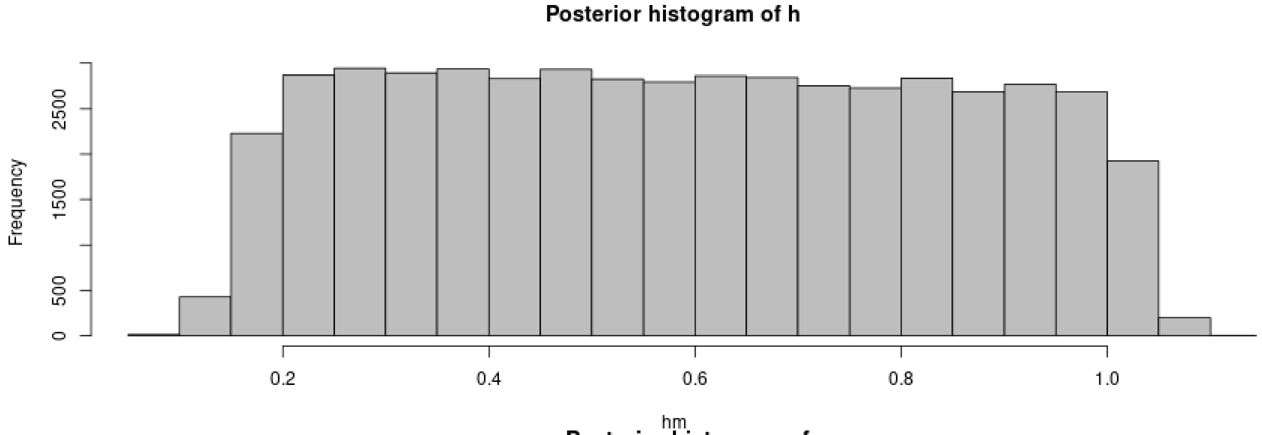

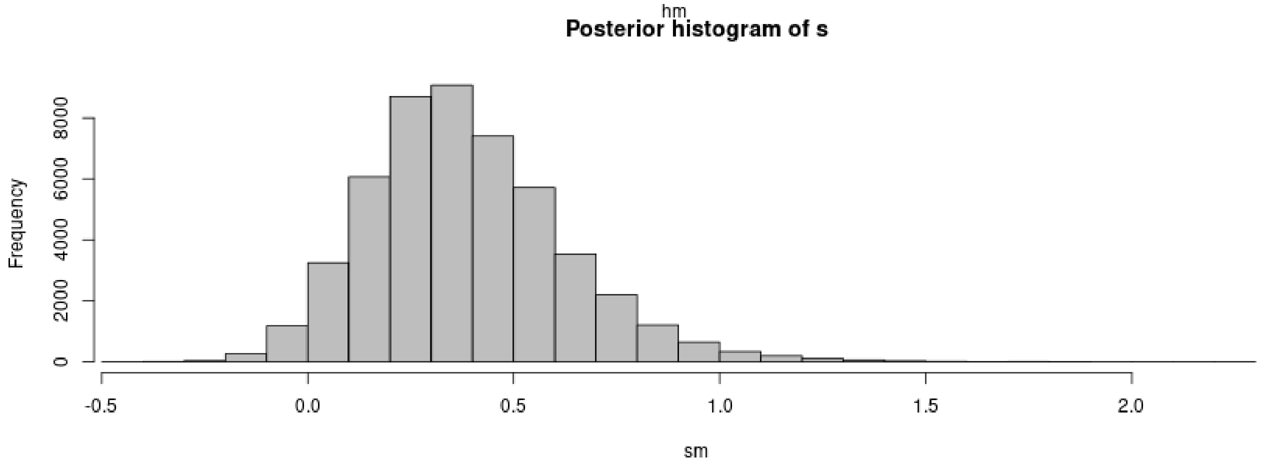

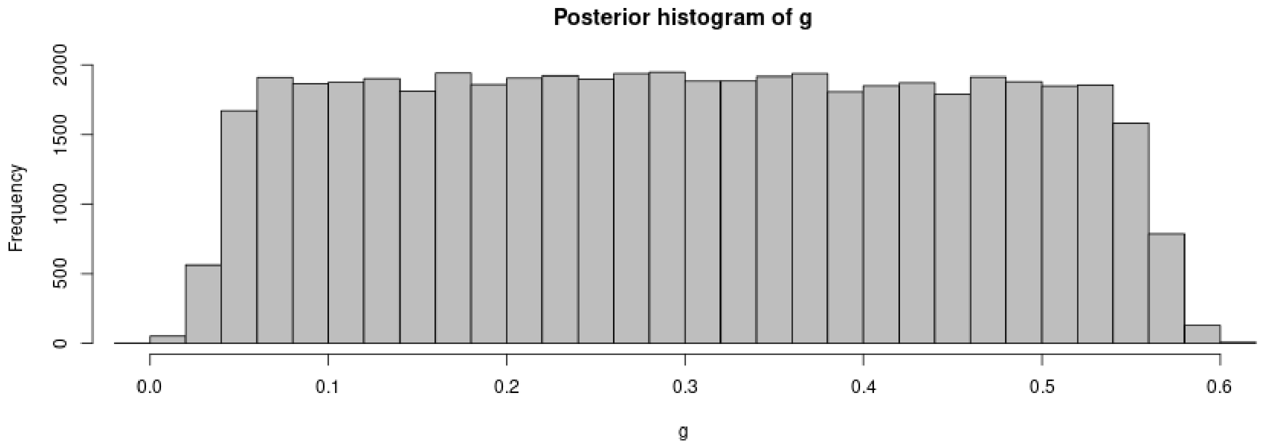

```
plot(m.r.g, param = param.sim)
```

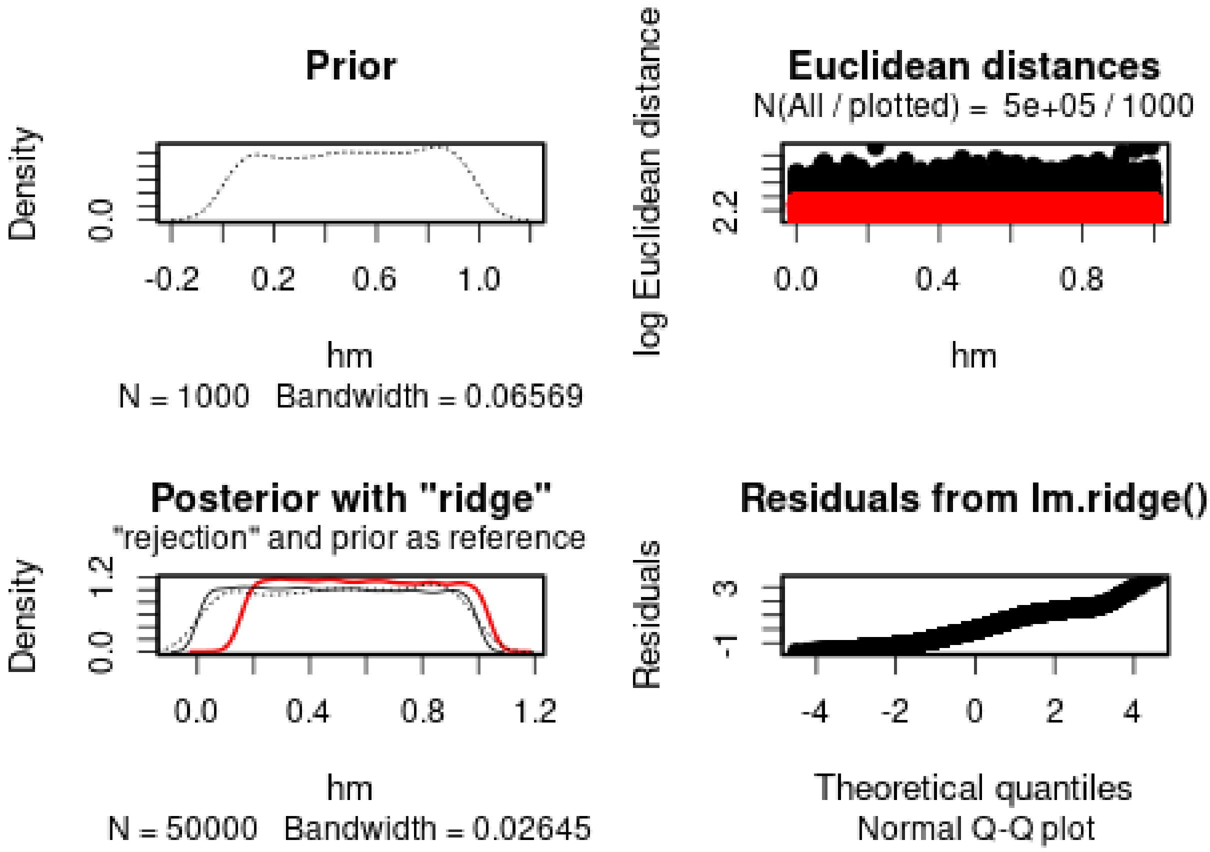

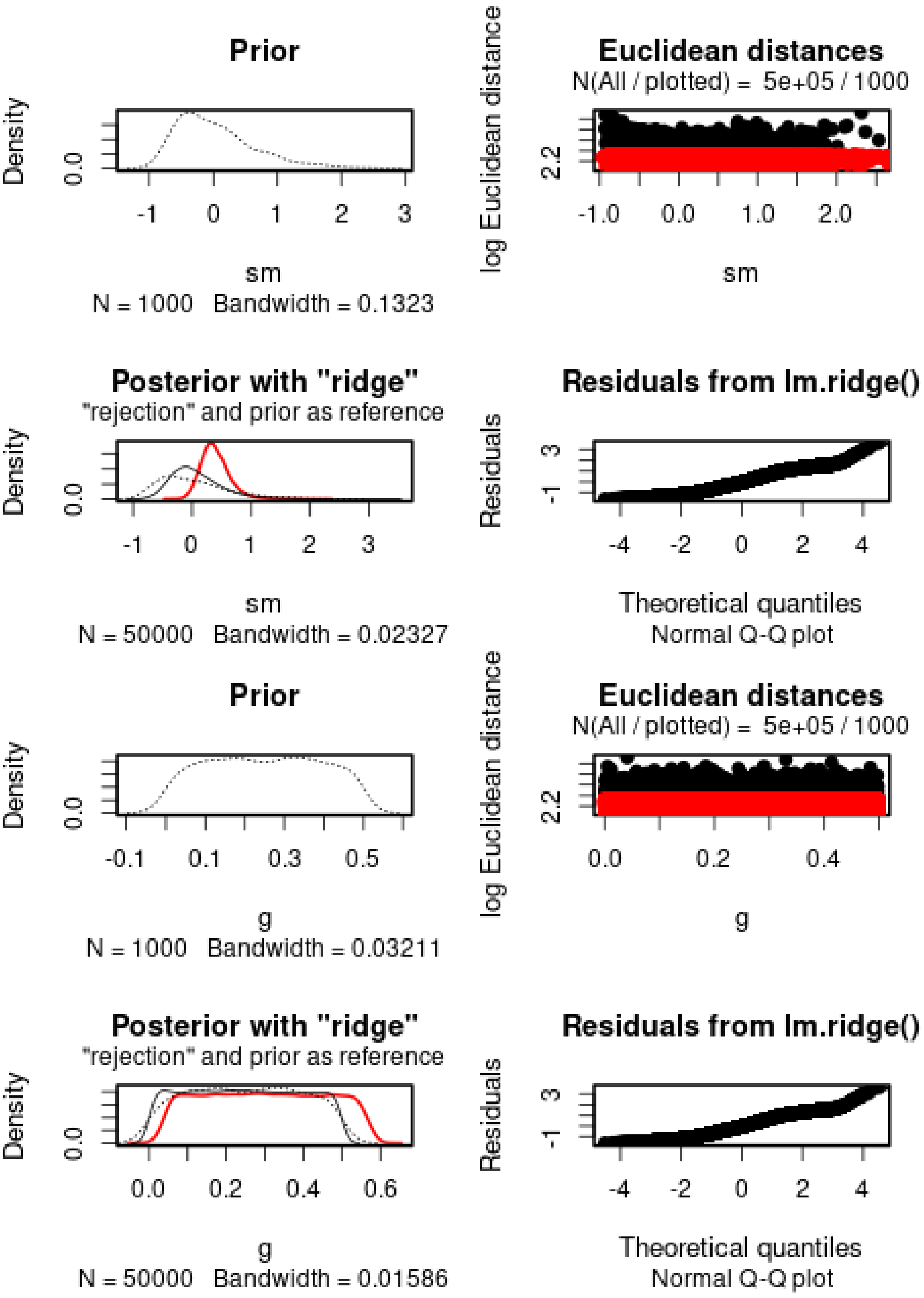
Prior
Density
0.0
-1
0
1
2
3
sm
N = 1000 Bandwidth = 0.1323
Euclidean distances
N(All / plotted) = 5e+05 / 1000
log Euclidean distance
2.2
-1.0
0.0
1.0
2.0
sm
Posterior with "ridge"
"rejection" and prior as reference
Density
0.0
-1
0
1
2
3
sm
N = 50000 Bandwidth = 0.02327
Residuals from lm.ridge()
Residuals
-1
3
-4
-2
0
2
4
Theoretical quantiles
Normal Q-Q plot
Prior
Density
0.0
-0.1
0.1
0.3
0.5
g
N = 1000 Bandwidth = 0.03211
Euclidean distances
N(All / plotted) = 5e+05 / 1000
log Euclidean distance
2.2
0.0
0.2
0.4
g
Posterior with "ridge"
"rejection" and prior as reference
Density
0.0
0.0
0.2
0.4
0.6
g
N = 50000 Bandwidth = 0.01586
Residuals from lm.ridge()
Residuals
-1
3
-4
-2
0
2
4
Theoretical quantiles
Normal Q-Q plot

$h$ cannot properly be estimated. $s$ is positive (s = 0.3973), but has a large variance. The 95% posterior interval includes 0. $g$ cannot be estimated.

We then check the distribution of summary statistics in the simulations:

```
library(reshape2)
library(ggplot2)
library(ggpubr)

df.stats.sim <- melt(stat.sim, variable.name = "Statistic", value.name = "Frequency")
```

```
## No id variables; using all as measure variables
```

```
l <- strsplit(as.character(df.stats.sim$Statistic), split = ".", fixed = TRUE)
df.stats.sim$Room <- sapply(l, function(x) x[1])
df.stats.sim$Genotype <- sapply(l, function(x) x[2])
df.stats.sim$Generation <- sapply(
  strsplit(as.character(df.stats.sim$Statistic), split = ".", fixed = TRUE),
  function(x) x[3])

df.stats.obs <- as.data.frame(stat.obs)
names(df.stats.obs) <- "Frequency"
df.stats.obs$Statistic <- row.names(df.stats.obs)
l <- strsplit(as.character(df.stats.obs$Statistic), split = ".", fixed = TRUE)
df.stats.obs$Room <- sapply(l, function(x) x[1])
df.stats.obs$Genotype <- sapply(l, function(x) x[2])
df.stats.obs$Generation <- sapply(
  strsplit(as.character(df.stats.obs$Statistic), split = ".", fixed = TRUE),
  function(x) x[3])

plot.obs <- function(gen) {
  p <- ggplot(data = subset(df.stats.sim, Generation == gen),
            aes(x = Frequency, y = ..density..)) +
    geom_histogram(bins = 50) +
    geom_vline(data = subset(df.stats.obs, Generation == gen),
             aes(xintercept = Frequency), color = "orange") +
    facet_grid(Genotype~Room) +
    theme_pubclean()
  return(p)
}
```

```
plot.obs(1) + ggtitle("Timepoint 4")
```

```
## Warning: The dot-dot notation (`..density..`) was deprecated in ggplot2 3.4.0.
## i Please use `after_stat(density)` instead.
## This warning is displayed once every 8 hours.
## Call `lifecycle::last_lifecycle_warnings()` to see where this warning was
## generated.
```

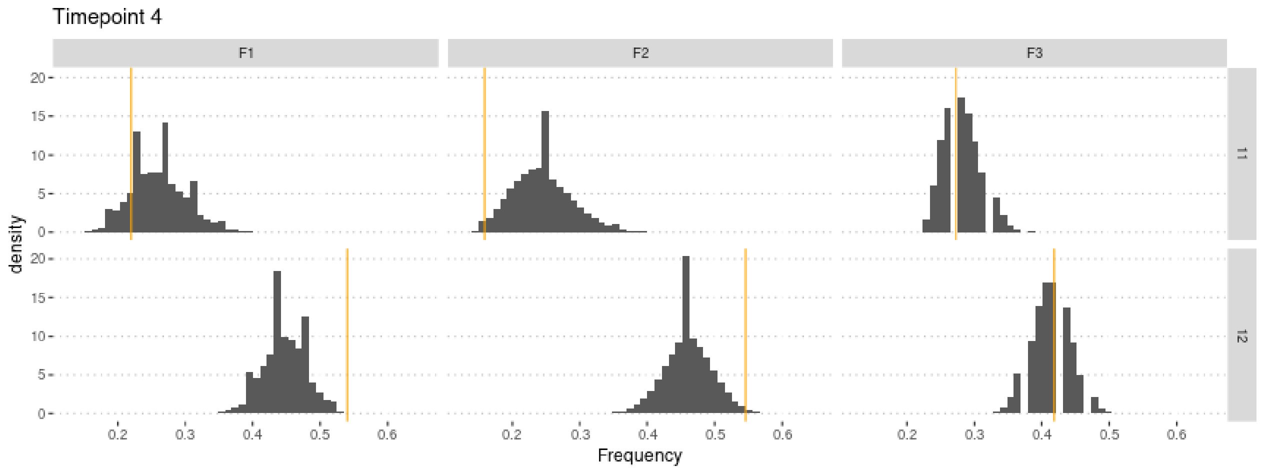

```
plot.obs(2) + ggtitle("Timepoint 5")
```

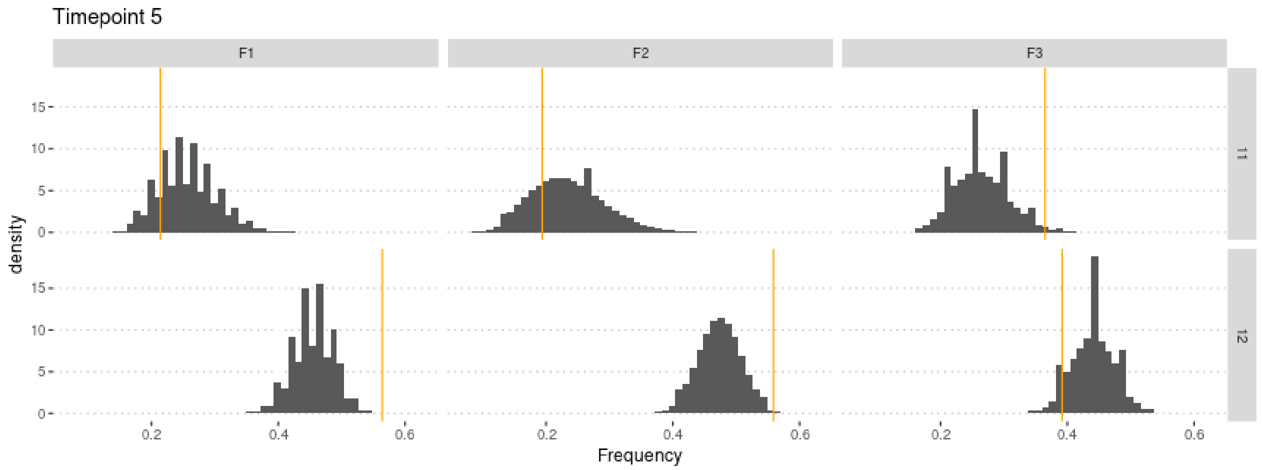

```
plot.obs(3) + ggtitle("Timepoint 6")
```

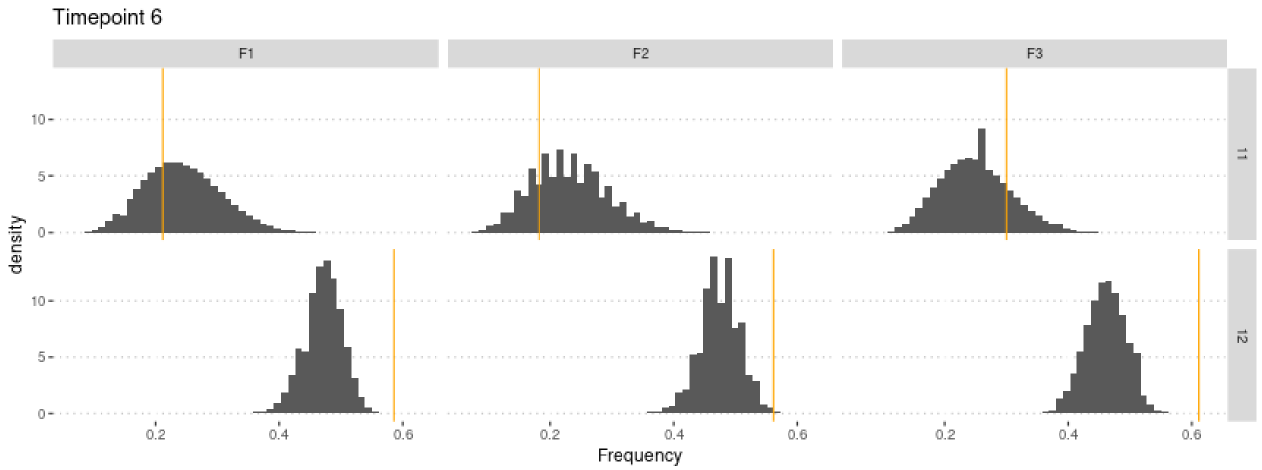

```
plot.obs(4) + ggtitle("Timepoint 7")
```

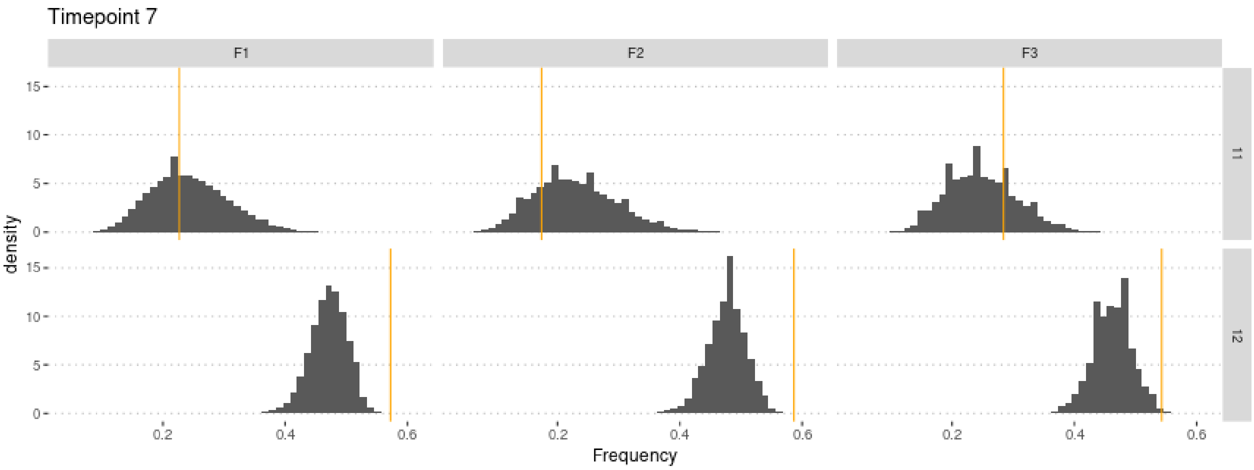

```
plot.obs(5) + ggtitle("Timepoint 8")
```

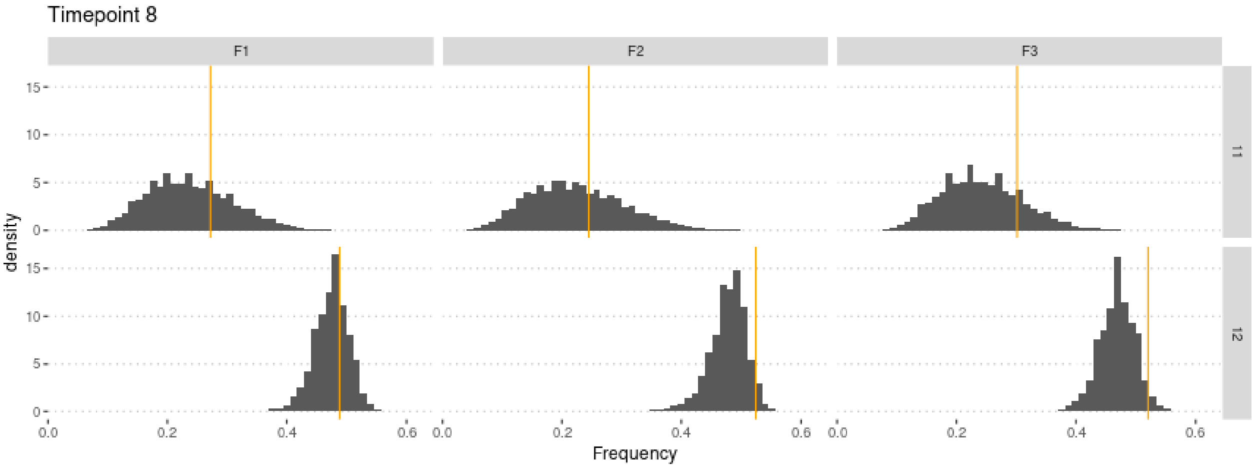

#### 4.1 Cross-Validation analysis

For memory-saving reasons, we only use 100,000 simulations in the following.

```
stat.sim2 <- stat.sim[1:1e5,]
m.r.g2 <- abc(target = stat.obs, param = param.sim2, sumstat = stat.sim2, tol = .1,
              method = "ridge", transf = c("none", "none", "none"))
```

Check that this does not affect the estimates too much:

```
summary(m.r.g2, intvl = .95)
```

```
## Call:
## abc(target = stat.obs, param = param.sim2, sumstat = stat.sim2,
##     tol = 0.1, method = "ridge", transf = c("none", "none", "none"))
## Data:
##  abc.out$adj.values (10000 posterior samples)
## Weights:
##  abc.out$weights
##
##                               hm      sm       g
## Min.:                    -0.0740 -1.0214 -0.0238
## Weighted 2.5 % Perc.:     0.0417 -0.2824  0.0205
```

```
## Weighted Median:         0.5135  0.3607  0.2300
## Weighted Mean:           0.5204  0.4065  0.2314
## Weighted Mode:           0.3382  0.2543  0.1308
## Weighted 97.5 % Perc.:   1.0072  1.3292  0.4437
## Max.:                    1.1126  3.1934  0.5228
```

Similar estimate, but slightly lower variance!

Compute predictions errors:

```
if (redo) {
  cv.ridge.g <- cv4abc(
      param = param.sim2,
      sumstat = stat.sim2,
      abc.out = m.r.g2,
      nval = 1000,
      tols = c(.01, .1, .2))
  save(cv.ridge.g, file = "Rdata/MalesOnlyNonGeneticVariance/backup_cv_genotype.Rdata")
} else {
  load("Rdata/MalesOnlyNonGeneticVariance/backup_cv_genotype.Rdata")
}
summary(cv.ridge.g)
```

```
## Prediction error based on a cross-validation sample of 1000
```

```
##             hm        sm         g
## 0.01 1.0068074 0.1562883 1.0436777
## 0.1  0.9647427 0.1578908 0.9895736
## 0.2  0.9689246 0.1607804 0.9885417
```

```
plot(cv.ridge.g)
```

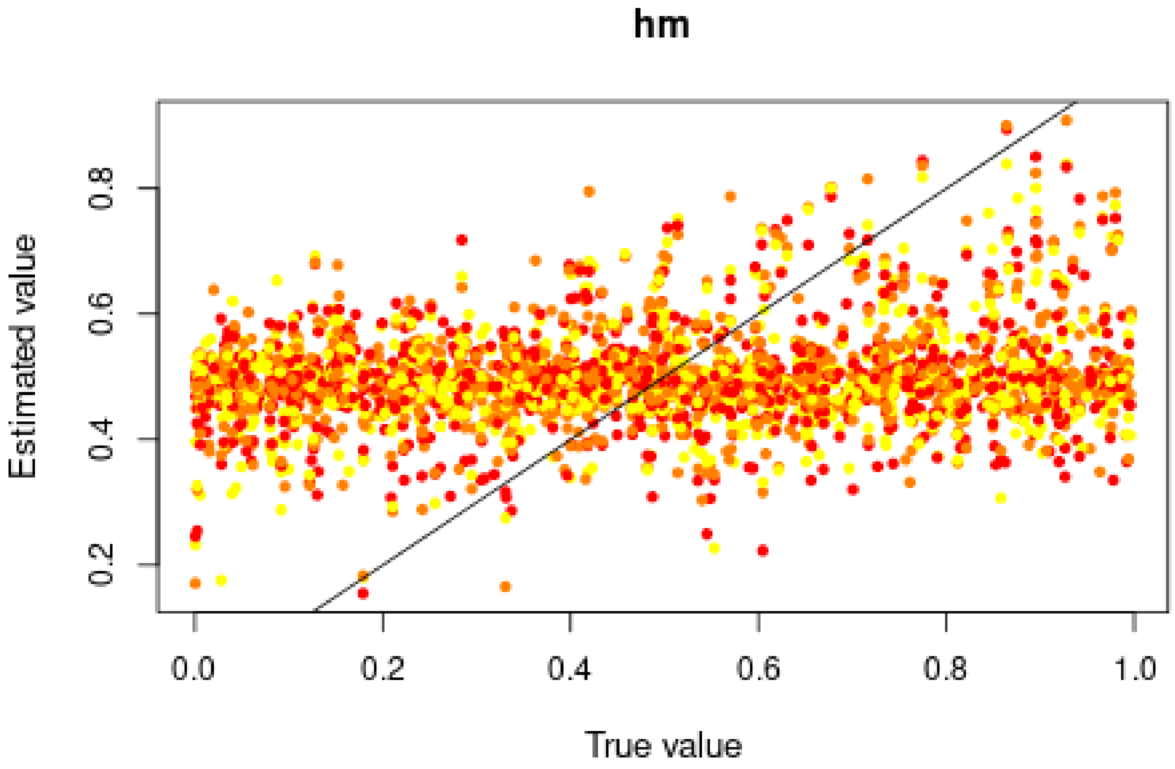

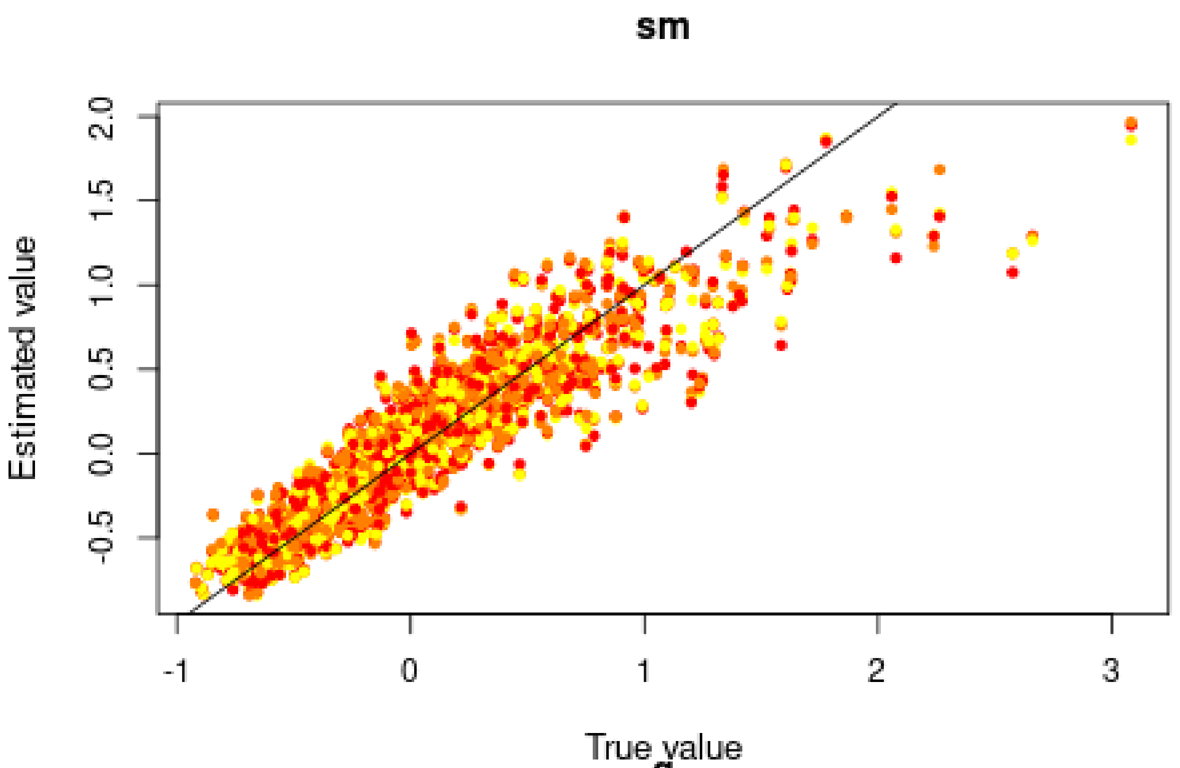

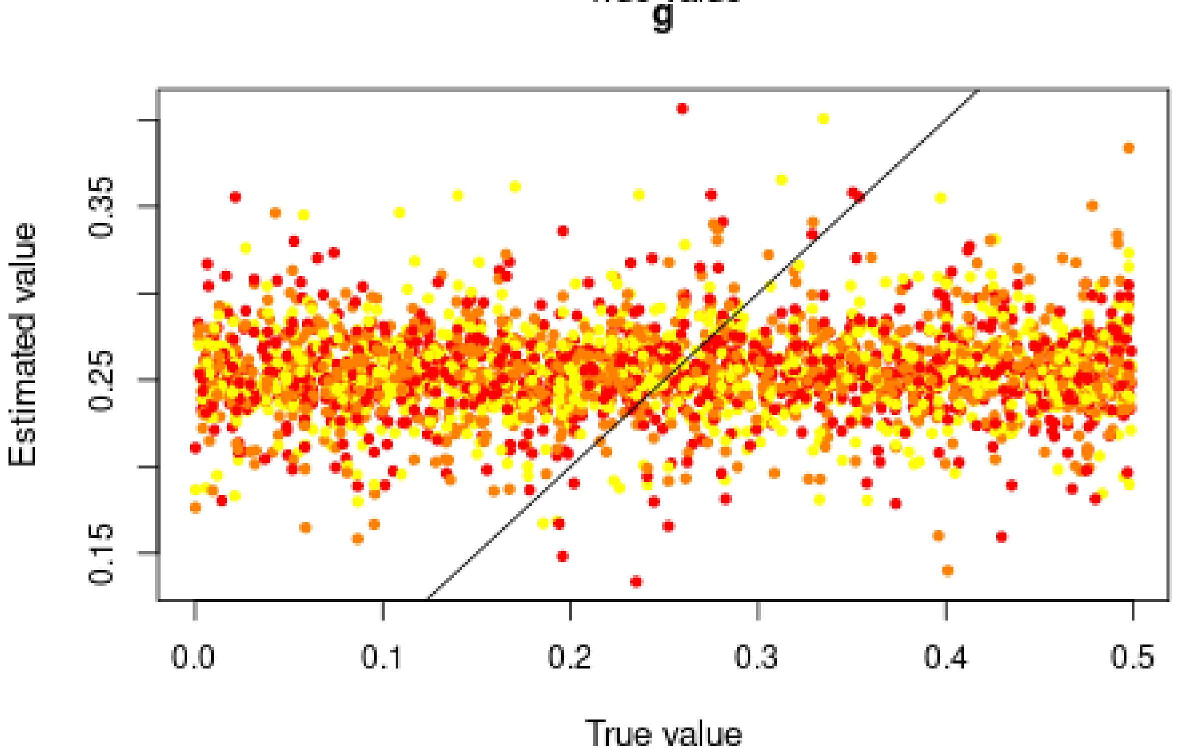

Good for $s$, but very bad for $h$ and $g$, in agreement with the posterior distributions.

#### 4.2 Misclassification errors

To test the power of the approach to distinguish between models, we also conduct a cross-validation experiment. We compare model M0 ($s = 0$) and M1 ($s = 0.3973 > 0$). $h$ and $g$ are sampled over their prior distributions. 100,000 simulations were conducted under each model.

```r
sims0 <- read.csv("Simulations/Simulations0WithNonGeneticVarianceFromGen3.csv.gz")
param.sim0 <- subset(sims0, select = c("hm", "sm", "g"))
stat.sim0 <- subset(sims0, select = 2:46)
# Only consider the WT homozygotes and the heterozygotes,
# as the sum of the three genotypes is 1
stat.sim0 <- subset(stat.sim0, select = grep(".22.", names(stat.sim0), invert = TRUE))

sims1 <- read.csv("Simulations/Simulations1bWithNonGeneticVarianceMalesOnlyFromGen3.csv.gz")
param.sim1 <- subset(sims1, select = c("hm", "sm", "g"))
stat.sim1 <- subset(sims1, select = 2:46)
# Only consider the WT homozygotes and the heterozygotes,
# as the sum of the three genotypes is 1
stat.sim1 <- subset(stat.sim1, select = grep(".22.", names(stat.sim1), invert = TRUE))
```

We conduct a CV analysis (this takes some time… set nval = 100 for faster results):

```r
models <- rep(c("Neutral", "Selection"), each = 100000)
if (redo) {
  cv.modsel.g <- cv4postpr(
    models, rbind(stat.sim0, stat.sim1),
    nval = 1000, tols=.1, method = "mnlogistic")
  save(cv.modsel.g, file = "Rdata/MalesOnlyNonGeneticVariance/backup_cv4postpr_genotype.Rdata")
} else {
  load("Rdata/MalesOnlyNonGeneticVariance/backup_cv4postpr_genotype.Rdata")
}
```

We display the results:

```r
summary(cv.modsel.g)
```

```
## Confusion matrix based on 1000 samples for each model.
##
## $tol0.1
##           Neutral Selection
## Neutral       811       189
## Selection     203       797
##
##
## Mean model posterior probabilities (mnlogistic)
##
## $tol0.1
##           Neutral Selection
## Neutral    0.7387    0.2613
## Selection  0.2688    0.7312
```

```r
plot(cv.modsel.g, names.arg = c("Neutral", "Selection"))
```

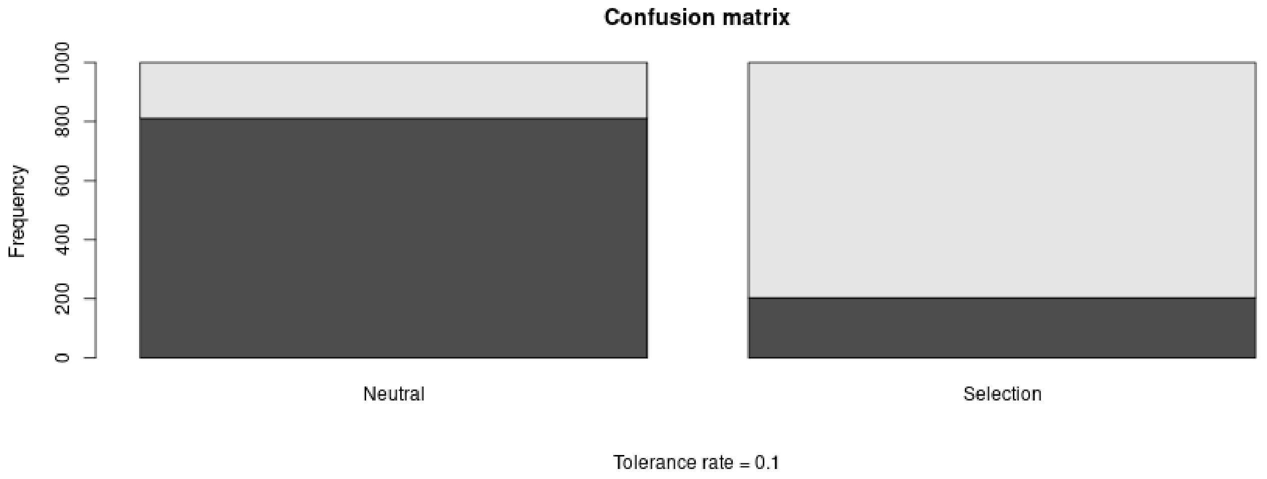

#### 4.3 Posterior prediction

```
if (redo) {
  modsel.g <- postpr(
    stat.obs, models,
    rbind(stat.sim0, stat.sim1),
    tol = .1, method = "mnlogistic")
  save(modsel.g, file = "Rdata/MalesOnlyNonGeneticVariance/backup_postpr_genotype.Rdata")
} else {
  load("Rdata/MalesOnlyNonGeneticVariance/backup_postpr_genotype.Rdata")
}
summary(modsel.g)
```

```
## Call:
## postpr(target = stat.obs, index = models, sumstat = rbind(stat.sim0,
##     stat.sim1), tol = 0.1, method = "mnlogistic")
## Data:
##  postpr.out$values (20000 posterior samples)
## Models a priori:
##  Neutral, Selection
## Models a posteriori:
##  Neutral, Selection
##
## Proportion of accepted simulations (rejection):
##   Neutral Selection
##    0.5352    0.4648
##
## Bayes factors:
##           Neutral Selection
## Neutral    1.0000    1.1517
## Selection  0.8683    1.0000
##
##
## Posterior model probabilities (mnlogistic):
##   Neutral Selection
##     0.081     0.919
##
## Bayes factors:
```

```
##           Neutral Selection
## Neutral    1.0000    0.0882
## Selection 11.3407    1.0000
```

The model with selection is preferred.

#### 4.4 Goodness of fit

Under the neutral model:

```
if (redo) {
  res.gfit0.g <- gfit(
    target = stat.obs, sumstat = stat.sim0,
    statistic = median, nb.replicate = 1000)
  save(res.gfit0.g, file = "Rdata/MalesOnlyNonGeneticVariance/backup_gfit0_genotype.Rdata")
} else {
  load("Rdata/MalesOnlyNonGeneticVariance/backup_gfit0_genotype.Rdata")
}
plot(res.gfit0.g, main = "Histogram under M0")
```

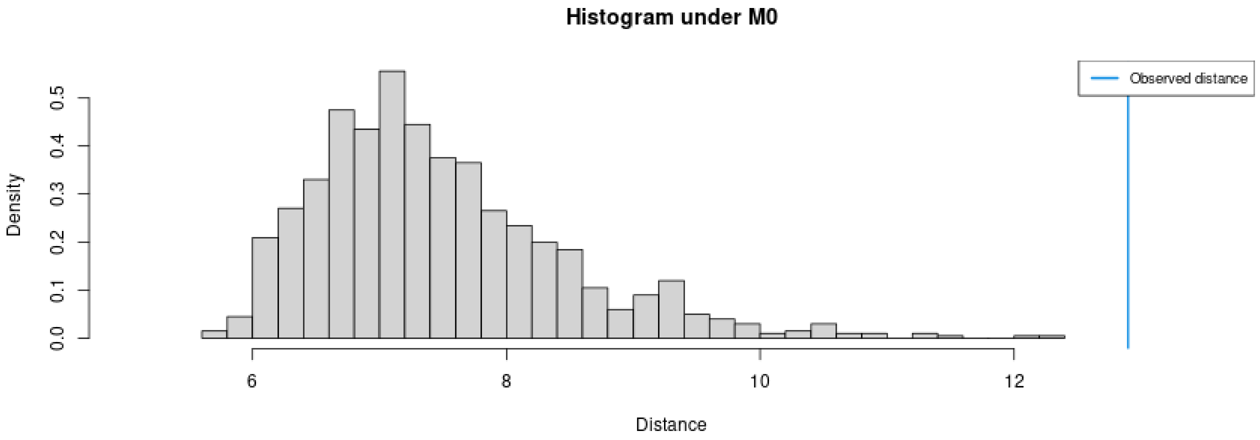

```
summary(res.gfit0.g)
```

```
## $pvalue
## [1] 0
##
## $s.dist.sim
##    Min. 1st Qu.  Median    Mean 3rd Qu.    Max.
##   5.708   6.764   7.287   7.466   7.966  12.271
##
## $dist.obs
## [1] 12.9008
```

Under the selection model:

```
if (redo) {
  res.gfit1.g <- gfit(
    target = stat.obs, sumstat = stat.sim1,
    statistic = median, nb.replicate = 1000)
  save(res.gfit1.g, file = "Rdata/MalesOnlyNonGeneticVariance/backup_gfit1_genotype.Rdata")
} else {
  load("Rdata/MalesOnlyNonGeneticVariance/backup_gfit1_genotype.Rdata")
}
plot(res.gfit1.g, main = "Histogram under M1")
```

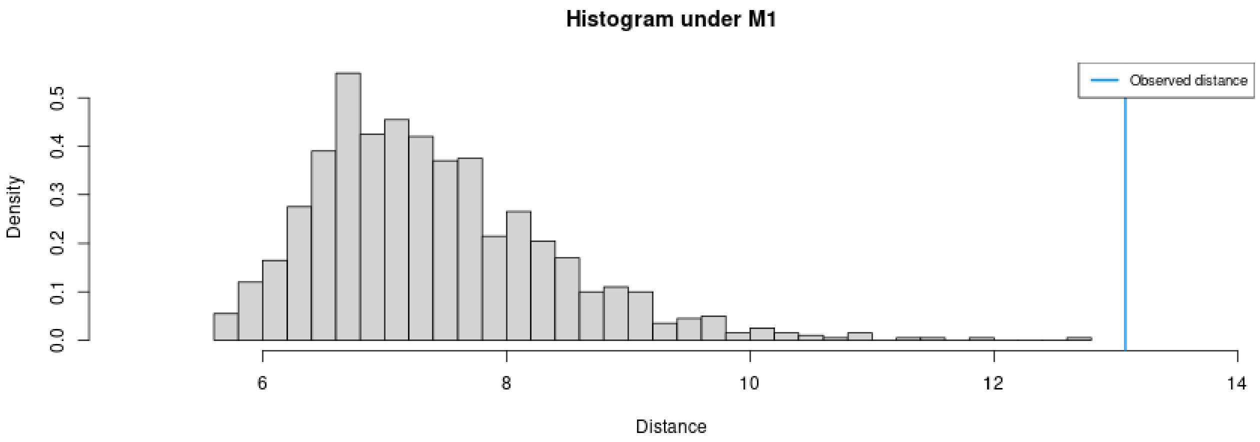

```
summary(res.gfit1.g)
```

```
## $pvalue
## [1] 0
##
## $s.dist.sim
##    Min. 1st Qu.  Median    Mean 3rd Qu.    Max.
##   5.636   6.685   7.238   7.395   7.927  12.691
##
## $dist.obs
## [1] 13.08
```

Both models provide a bad fit.

#### 4.5 Summary figure

Posterior distributions, with prior for comparison:

```
library(ggplot2)
library(ggpubr)
dat.prior.h <- data.frame(h = param.sim[,"hm"], Distribution = "Prior")
dat.post.h <- data.frame(h = m.r.g$adj.values[,"hm"], Distribution = "Posterior")
dat.prior.s <- data.frame(s = param.sim[,"sm"], Distribution = "Prior")
dat.post.s <- data.frame(s = m.r.g$adj.values[,"sm"], Distribution = "Posterior")
dat.prior.g <- data.frame(s = param.sim[,"g"], Distribution = "Prior")
dat.post.g <- data.frame(s = m.r.g$adj.values[,"g"], Distribution = "Posterior")
dat.h <- rbind(dat.prior.h, dat.post.h)
dat.s <- rbind(dat.prior.s, dat.post.s)
dat.g <- rbind(dat.prior.g, dat.post.g)
dat.h$Distribution <- factor(dat.h$Distribution, levels = c("Prior", "Posterior"))
dat.s$Distribution <- factor(dat.s$Distribution, levels = c("Prior", "Posterior"))
dat.g$Distribution <- factor(dat.g$Distribution, levels = c("Prior", "Posterior"))
names(dat.h)[1] <- "value"
dat.h$variable <- "h"
names(dat.s)[1] <- "value"
dat.s$variable <- "s"
names(dat.g)[1] <- "value"
dat.g$variable <- "g"
dat.dist <- rbind(dat.h, dat.s, dat.g)
dat.dist$variable <- factor(dat.dist$variable, levels = c("h", "s", "g"))
```

```r
p.dist <- ggplot(data = dat.dist,
              aes(x = value, linetype = Distribution, fill = Distribution)) +
          geom_density(alpha = 0.5) +
          scale_fill_brewer(type = "qual", palette = 3) +
          scale_linetype_manual(values = c(Prior = "dashed", Posterior = "solid")) +
          xlab("Parameter value") +
          facet_wrap(~variable, scales = "free") +
          ggtitle("Posterior and prior parameter distributions") +
          theme_pubclean() + theme(strip.background = element_blank())
```

Cross-validation:

```r
d1<-rbind(as.data.frame(cv.ridge.g$estim$tol0.01),
          as.data.frame(cv.ridge.g$estim$tol0.1),
          as.data.frame(cv.ridge.g$estim$tol0.2))
d2<-rbind(cv.ridge.g$true, cv.ridge.g$true, cv.ridge.g$true)
d1<-melt(d1, value.name = "Estimated")
```

```
## No id variables; using all as measure variables
```

```r
d1$Tolerance <- rep(c(0.01, 0.1, 0.2), each = 1000)
d1$Replicate <- rep(1:1000, 3)
d2<-melt(d2, value.name = "True")
```

```
## No id variables; using all as measure variables
```

```r
d2$Replicate <- rep(1:1000, 3)
dat.cv <- merge(d1, d2, by = c("variable", "Replicate"))
dat.cv$variable <- substr(dat.cv$variable, 1, 1)
dat.cv$variable <- factor(dat.cv$variable, levels = c("h", "s", "g"))
```

```r
p.cv <- ggplot(dat.cv, aes(x = True, y = Estimated)) +
        geom_point(aes(col = as.ordered(Tolerance))) +
        geom_abline(slope = 1) +
        facet_wrap(~variable, scales = "free") +
        theme_pubclean() + labs(color = "Tolerance") +
        ggtitle("Cross validation") +
        theme(strip.background = element_blank())
```

Confusion matrix:

```r
library(scales)
cv.sum <- summary(cv.modsel.g)
```

```
## Confusion matrix based on 1000 samples for each model.
##
## $tol0.1
##           Neutral Selection
## Neutral       811       189
## Selection     203       797
##
##
## Mean model posterior probabilities (mnlogistic)
##
## $tol0.1
##           Neutral Selection
```

```
## Neutral    0.7387    0.2613
## Selection  0.2688    0.7312
```

```
dat.cv <- as.data.frame(cv.sum$conf.matrix$tol0.1/1000)
names(dat.cv) <- c("Real", "Inferred", "Frequency")
p.confmat <- ggplot(dat.cv, aes(x = Real, y = Frequency, fill = Inferred)) +
             geom_col() +
             scale_y_continuous(labels = scales::percent) +
             scale_fill_brewer(type = "qual", palette = 3) +
             ggtitle("Prediction errors") +
             theme_pubclean()
```

Model probabilities:

```
p.mprob <- ggplot(as.data.frame(modsel.g$pred), aes(x = Var1, y = Freq, fill = Var1)) +
           geom_col() + ylab("Model posterior probability") + xlab("Model") +
           scale_fill_brewer(type = "qual", palette = 3) +
           theme_pubclean() +
           ggtitle("Model probabilities") +
           theme(legend.position = "none")
```

```
library(cowplot)
p1 <- plot_grid(p.mprob, p.confmat, labels = c("C", "D"), nrow = 1)
p <- plot_grid(p.dist, p.cv, p1, labels = c("A", "B", ""), nrow = 3)
p
```

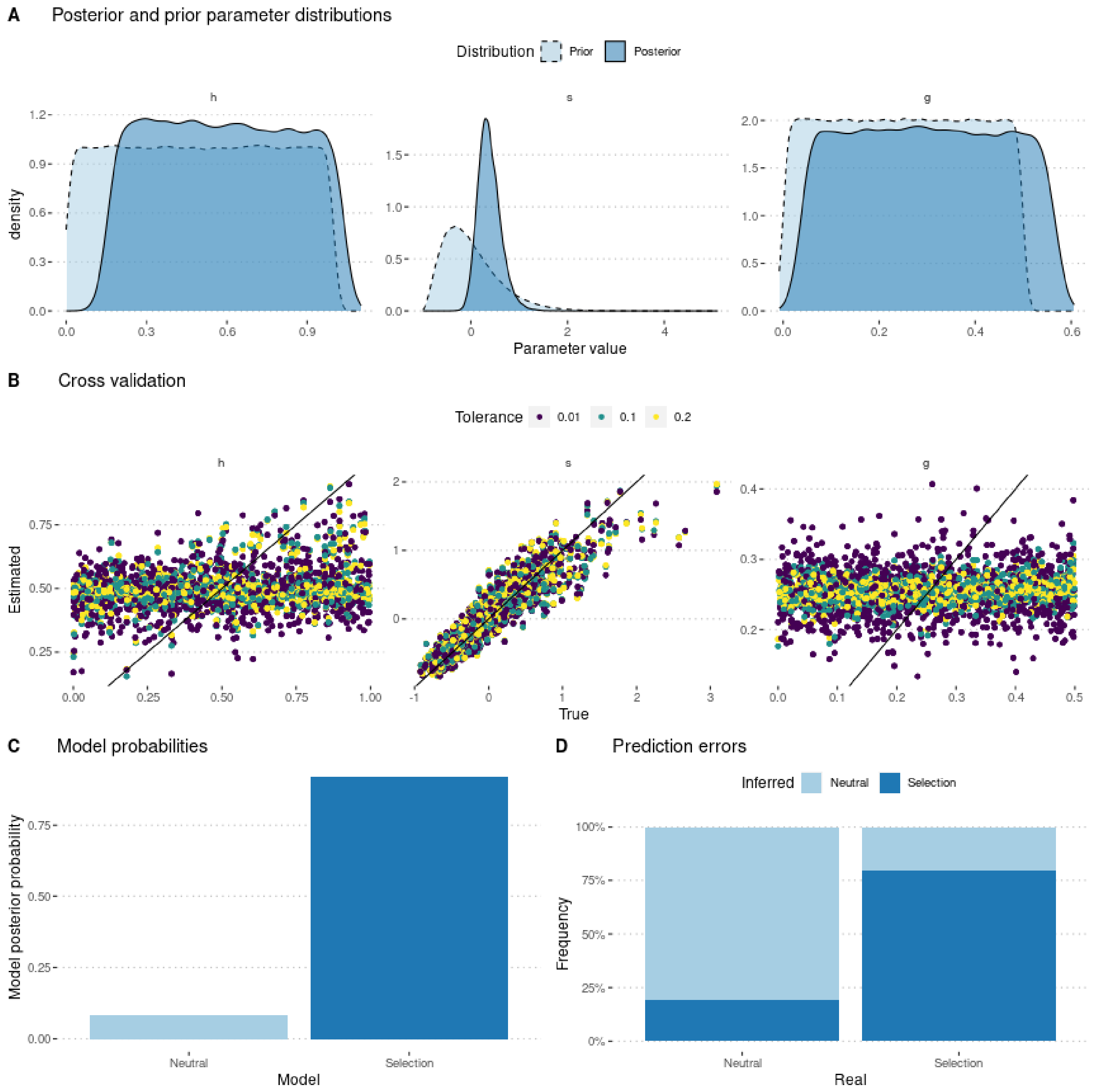

```
ggsave(p, filename = "FigureABC-MalesOnlyNonGeneticVariance-Genotype.pdf", width = 8, height = 10)
```
