## Supplementary material for "Experimental evaluation of a direct fitness effect of the *de novo* evolved mouse gene *Pldi*": combined suppl files: SupplementaryFile6.pdf

### Estimating poldi's selection coefficient

Julien Y. Dutheil

17/02/2023

#### Contents

**1 Model parameterization** **1**

**2 Preamble** **1**

**3 Using allele frequencies** **2**

**4 Using genotype frequencies** **8**

### 1 Model parameterization

The model includes two parameters: $s$ the selection coefficient of the *poldi* allele compared to the knockout strain, which mimics the ancestral state, and $h$ the heterozygosity. Fitness values of each genotype are parameterized as follow:

$$\begin{aligned} poldi/poldi &: 1 \\ BL6/poldi &: 1 + h \cdot s \\ BL6/BL6 &: 1 + s \end{aligned}$$

where BL6 denotes the wild strain, and poldi the strain where the *poldi* gene is knocked out. Furthermore, we consider that the allele only has an effect in males, so that $h$ and $s$ are 0 in females (the two alleles have the same fitness). In this version, we model non random mating by adding a parameter $\lambda$, so that parents are chosen according to their fitness multiplied by $(1 - \lambda)$ if they have the same two alleles, $(1 - \lambda/2)$ if they have only one allele in common, or 1 if both alleles are different.

We performed 200,000 simulations, using a gamma prior for $s$, a uniform prior between 0 and 1 for $h$, and a uniform prior between 0 and 1 for *lambda*.

### 2 Preamble

We load the data (observed and simulated):

```
sims <- read.csv("Simulations/SimulationsNonRandomMatingMalesOnlyFromGen3.csv.gz")
param.sim <- subset(sims, select = c("hm", "sm", "lambda"))
```

We estimate parameters using the ridge regression method:

```
library(abc, quietly = TRUE)
```

```
##
## Attaching package: 'SparseM'

## The following object is masked from 'package:base':
##
##     backsolve

## locfit 1.5-9.8    2023-06-11
```

```
if (redo) {
  m.r.a <- abc(target = stat.obs.a, param = param.sim, sumstat = stat.sim.a, tol=.1,
              method = "ridge", transf = c("none", "none", "none"))
  save(m.r.a, file = "Rdata/MalesOnlyNonRandomMating/backup_abc_allelic.Rdata")
} else {
  load("Rdata/MalesOnlyNonRandomMating/backup_abc_allelic.Rdata")
}
```

Now we display the results:

```
summary(m.r.a, intvl = .95)
```

```
## Call:
## abc(target = stat.obs.a, param = param.sim, sumstat = stat.sim.a,
##     tol = 0.1, method = "ridge", transf = c("none", "none", "none"))
## Data:
##  abc.out$adj.values (20000 posterior samples)
## Weights:
##  abc.out$weights
```

```
##
##                                 hm      sm  lambda
## Min.:                      -0.2199 -0.3293 -0.1367
## Weighted 2.5 % Perc.:      -0.0966  0.0417 -0.0315
## Weighted Median:            0.5046  0.5797  0.4277
## Weighted Mean:              0.5010  0.6155  0.4269
## Weighted Mode:              0.8682  0.5202  0.5560
## Weighted 97.5 % Perc.:      1.0855  1.3879  0.8726
## Max.:                       1.1956  3.3688  0.9652
```

Distribution of parameter estimates:

```r
hist(m.r.a, breaks = 30, caption = c(expression(h), expression(s), expression(lambda)))
```

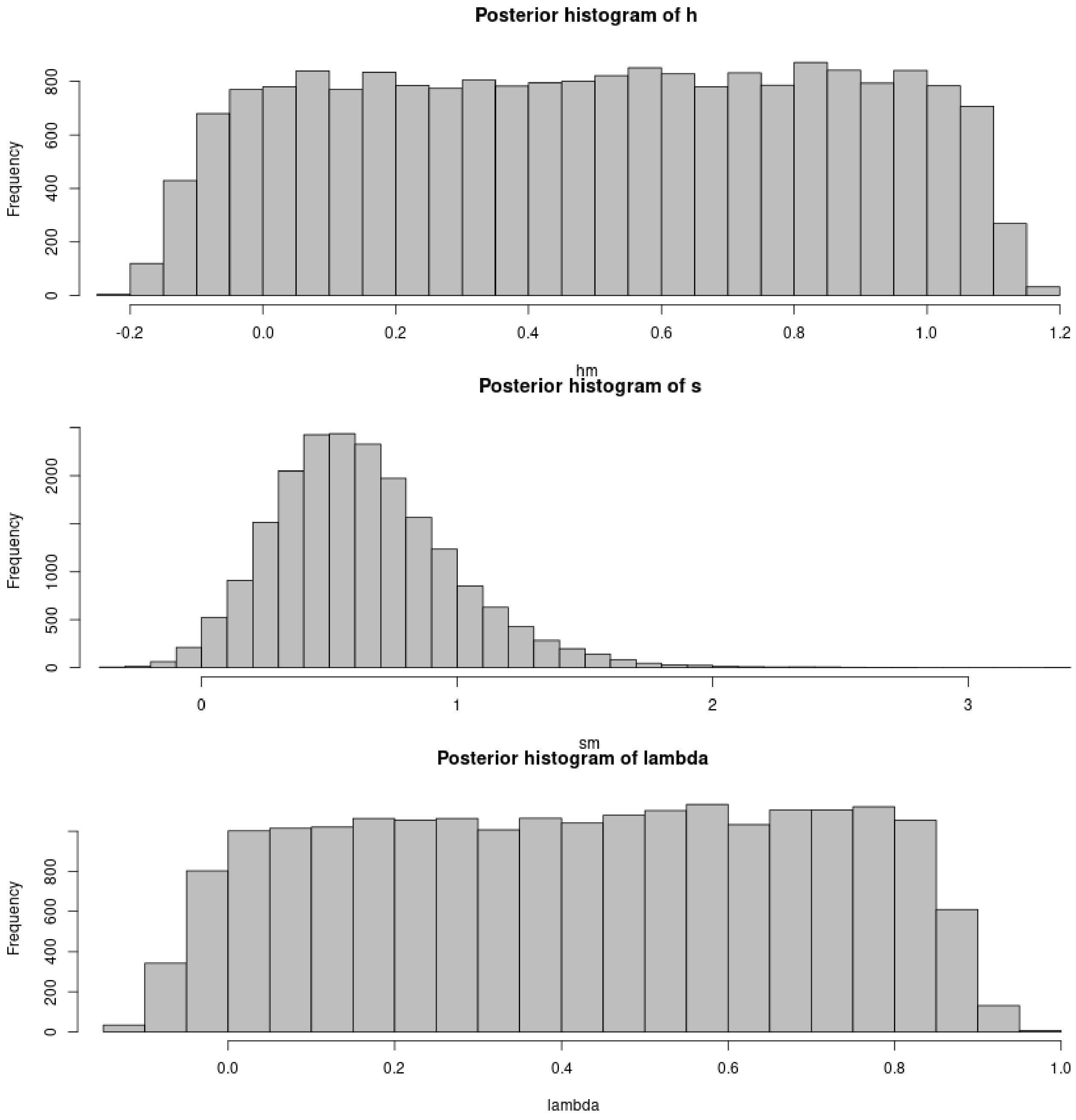

```
plot(m.r.a, param = param.sim)
```

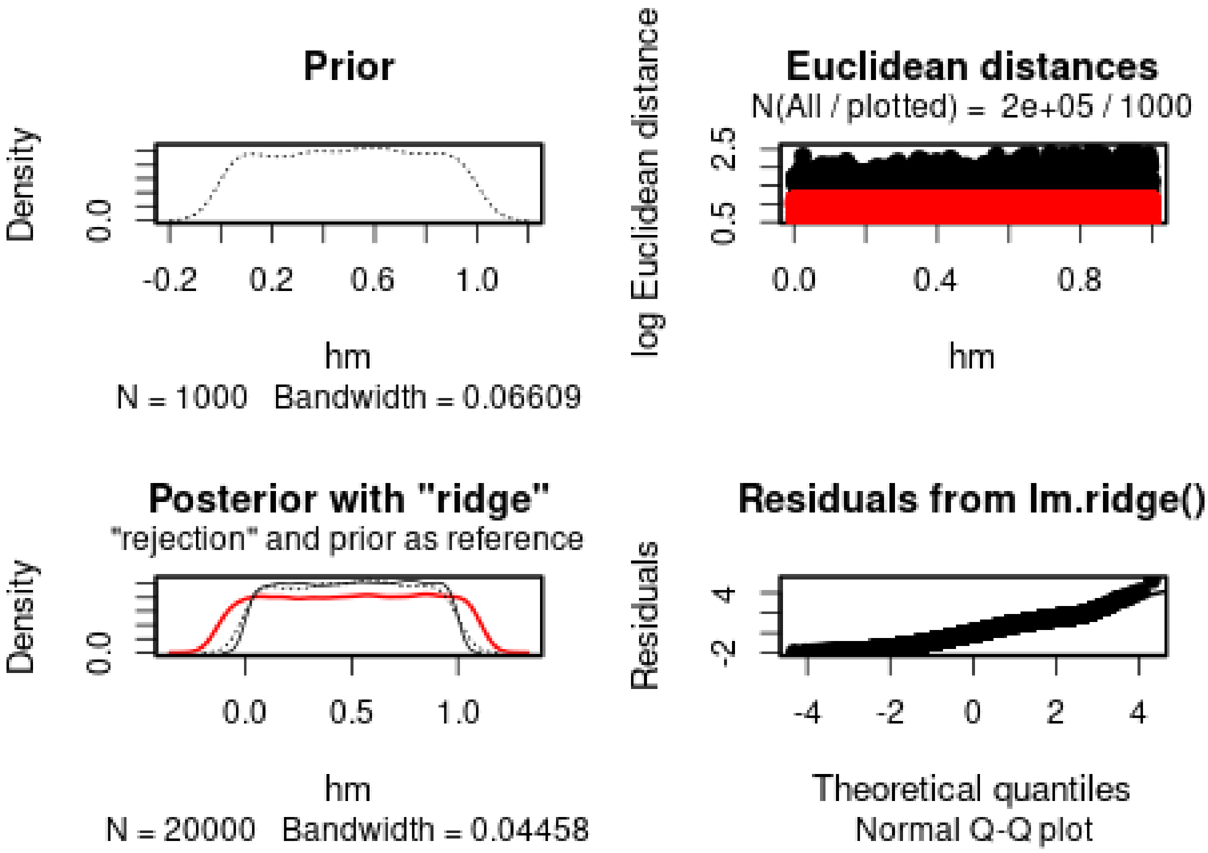

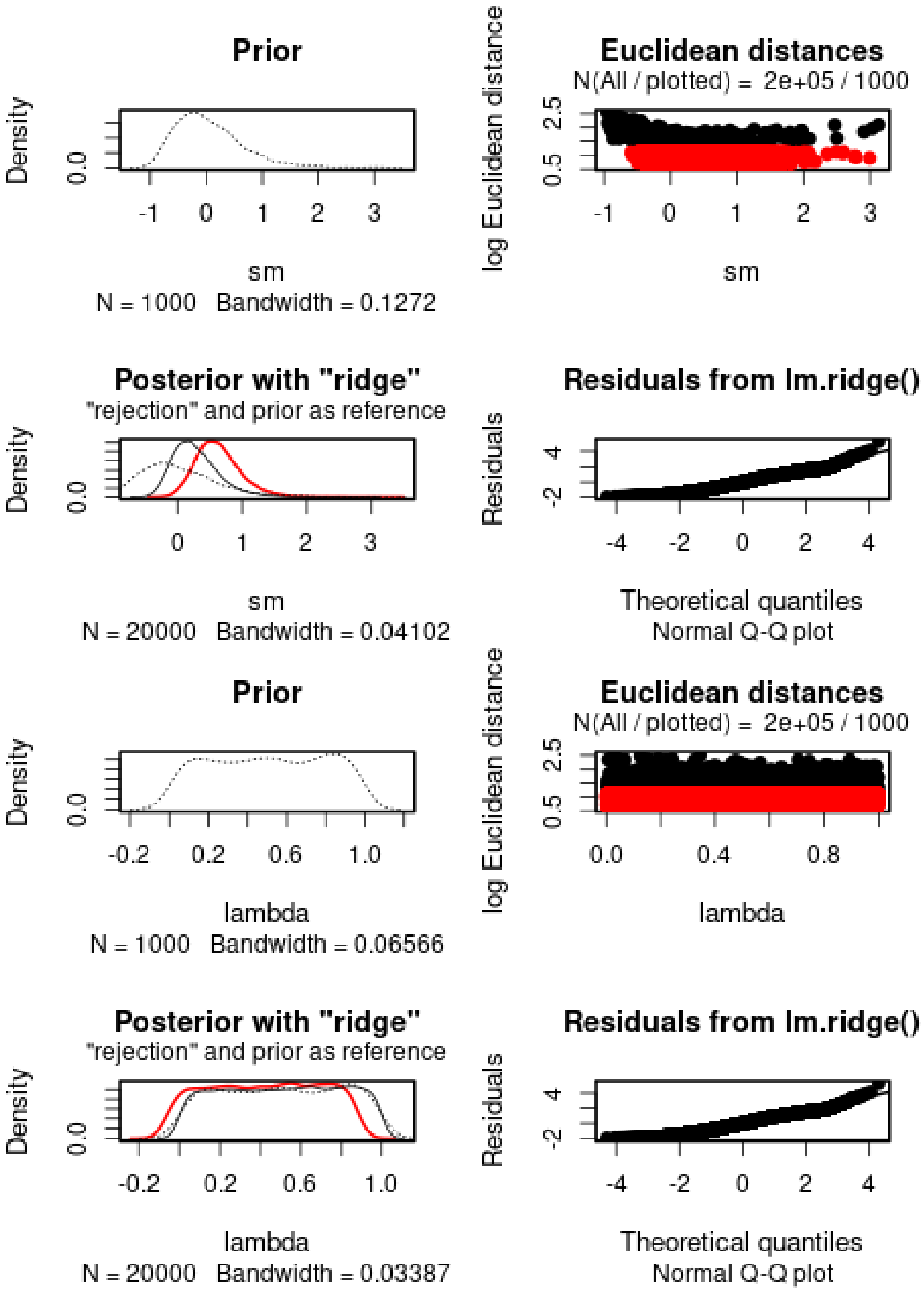
Prior
Density
0.0
-1 0 1 2 3
sm
N = 1000 Bandwidth = 0.1272
Euclidean distances
N(All / plotted) = 2e+05 / 1000
log Euclidean distance
0.5 2.5
-1 0 1 2 3
sm
Posterior with "ridge"
"rejection" and prior as reference
Density
0.0
0 1 2 3
sm
N = 20000 Bandwidth = 0.04102
Residuals from lm.ridge()
Residuals
-2 4
-4 -2 0 2 4
Theoretical quantiles
Normal Q-Q plot
Prior
Density
0.0
-0.2 0.2 0.6 1.0
lambda
N = 1000 Bandwidth = 0.06566
Euclidean distances
N(All / plotted) = 2e+05 / 1000
log Euclidean distance
0.5 2.5
0.0 0.4 0.8
lambda
Posterior with "ridge"
"rejection" and prior as reference
Density
0.0
-0.2 0.2 0.6 1.0
lambda
N = 20000 Bandwidth = 0.03387
Residuals from lm.ridge()
Residuals
-2 4
-4 -2 0 2 4
Theoretical quantiles
Normal Q-Q plot

$h$ cannot properly be estimated. $s$ is positive and the 95% posterior interval does not include 0. Lambda cannot be estimated.

We then check the distribution of summary statistics in the simulations:

```r
library(reshape2)
library(ggplot2)
library(ggpubr)
```

```
##
## Attaching package: 'ggpubr'

## The following object is masked from 'package:plyr':
##
##     mutate
```

```r
df.stats.sim <- melt(stat.sim.a, variable.name = "Statistic", value.name = "Frequency")
```

```
## No id variables; using all as measure variables
```

```r
l <- strsplit(as.character(df.stats.sim$Statistic), split = ".", fixed = TRUE)
df.stats.sim$Room <- sapply(l, function(x) x[1])
df.stats.sim$Generation <- sapply(
  strsplit(as.character(df.stats.sim$Statistic), split = ".", fixed = TRUE),
  function(x) x[2])

df.stats.obs <- as.data.frame(stat.obs.a)
names(df.stats.obs) <- "Frequency"
df.stats.obs$Statistic <- row.names(df.stats.obs)
l <- strsplit(as.character(df.stats.obs$Statistic), split = ".", fixed = TRUE)
df.stats.obs$Room <- sapply(l, function(x) x[1])
df.stats.obs$Generation <- sapply(
  strsplit(as.character(df.stats.obs$Statistic), split = ".", fixed = TRUE),
  function(x) x[2])

p <- ggplot(data = df.stats.sim, aes(x = Frequency, y = after_stat(density))) +
       geom_histogram(bins = 50) +
       geom_vline(data = df.stats.obs, aes(xintercept = Frequency), color = "orange") +
       facet_grid(Room~Generation) +
       theme_pubclean()
p
```

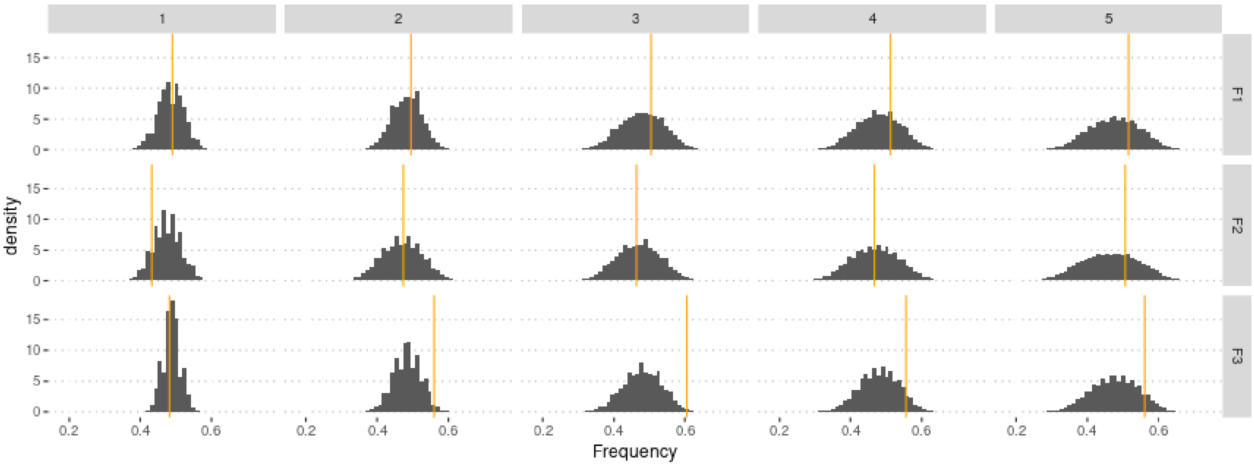

The *lambda* parameter is not identifiable when only allelic frequencies are used.

### 4 Using genotype frequencies

We estimate parameters using the ridge regression method:

```
if (redo) {
  m.r.g <- abc(target = stat.obs, param = param.sim, sumstat = stat.sim, tol = .1,
               method = "ridge", transf = c("none", "none", "none"))
  save(m.r.g, file = "Rdata/MalesOnlyNonRandomMating/backup_abc_genotype.Rdata")
} else {
  load("Rdata/MalesOnlyNonRandomMating/backup_abc_genotype.Rdata")
}
```

Now we display the results:

```
summary(m.r.g, intvl = .95)
```

```
## Call:
## abc(target = stat.obs, param = param.sim, sumstat = stat.sim,
##     tol = 0.1, method = "ridge", transf = c("none", "none", "none"))
## Data:
##  abc.out$adj.values (20000 posterior samples)
## Weights:
##  abc.out$weights
##
##                              hm      sm  lambda
## Min.:                   -0.1329 -0.4839 -2.3049
## Weighted 2.5 % Perc.:   -0.0162  0.0108 -0.5665
## Weighted Median:         0.4879  0.6313  0.4431
## Weighted Mean:           0.4905  0.6694  0.3890
## Weighted Mode:           0.1802  0.5999  0.5640
## Weighted 97.5 % Perc.:   0.9973  1.5483  1.0276
## Max.:                    1.0930  4.4139  1.3824
```

Distribution of parameter estimates:

```
hist(m.r.g, breaks = 30, caption = c(expression(h), expression(s), expression(lambda)))
```

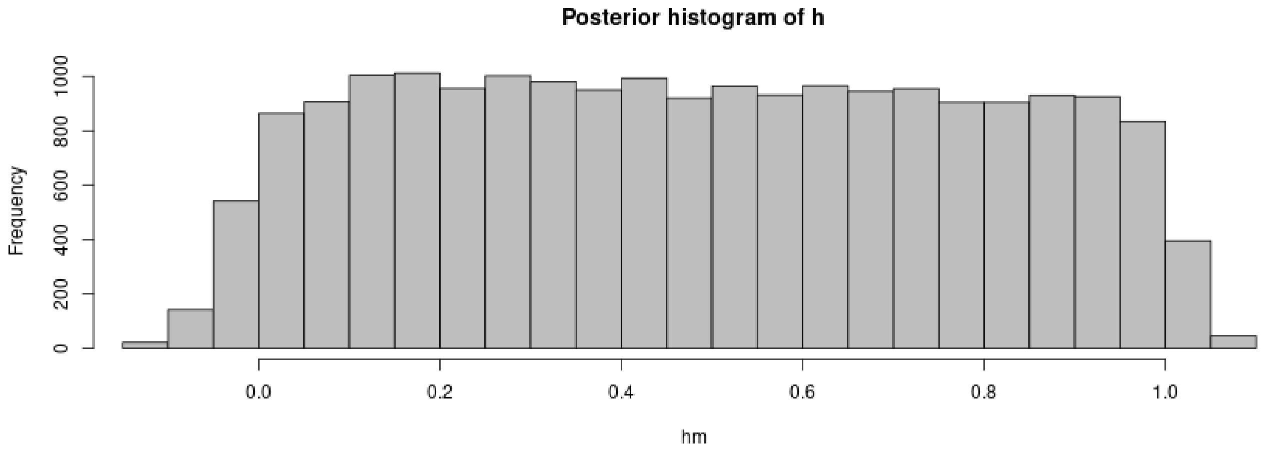

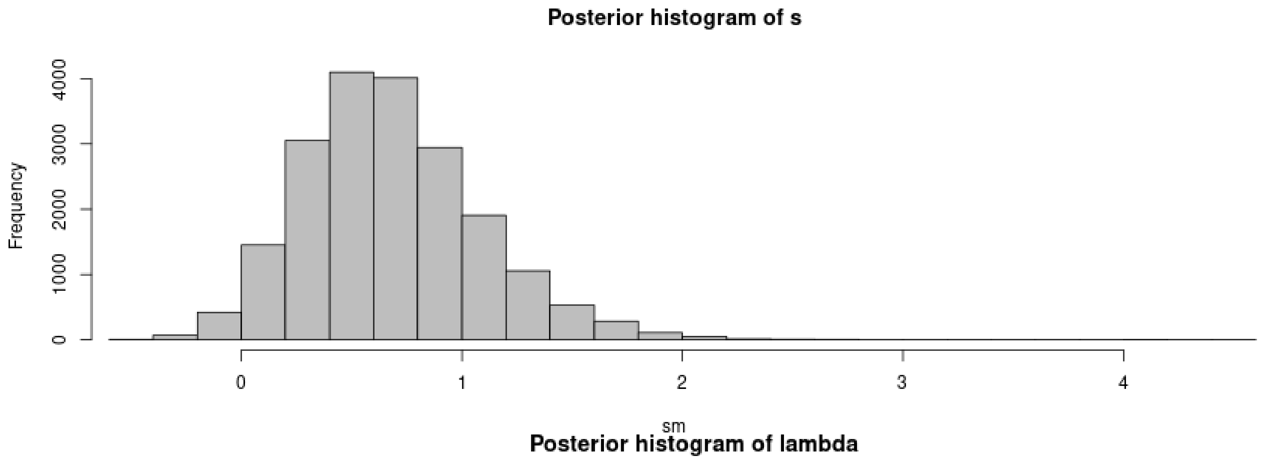

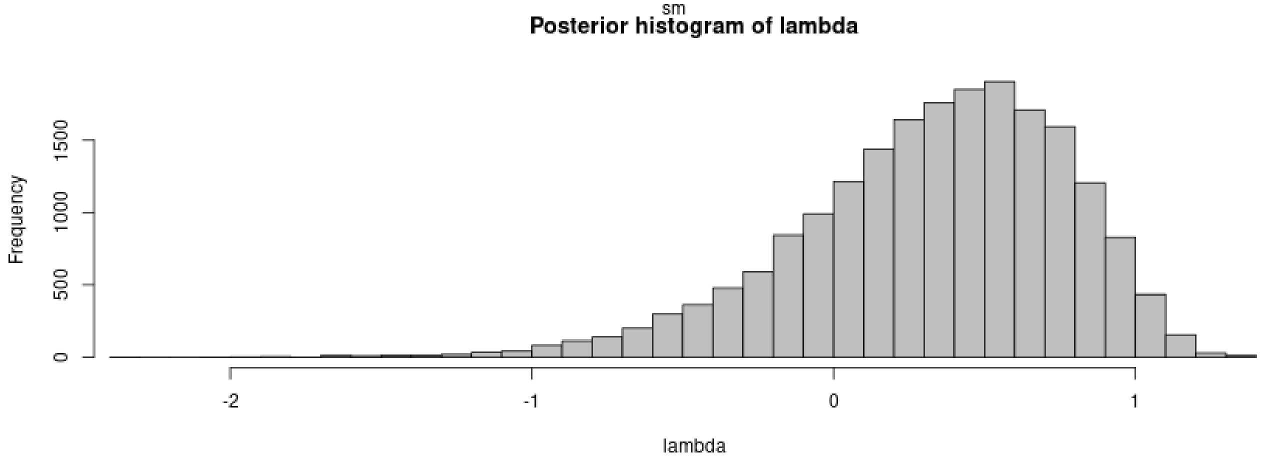

```
plot(m.r.g, param = param.sim)
```

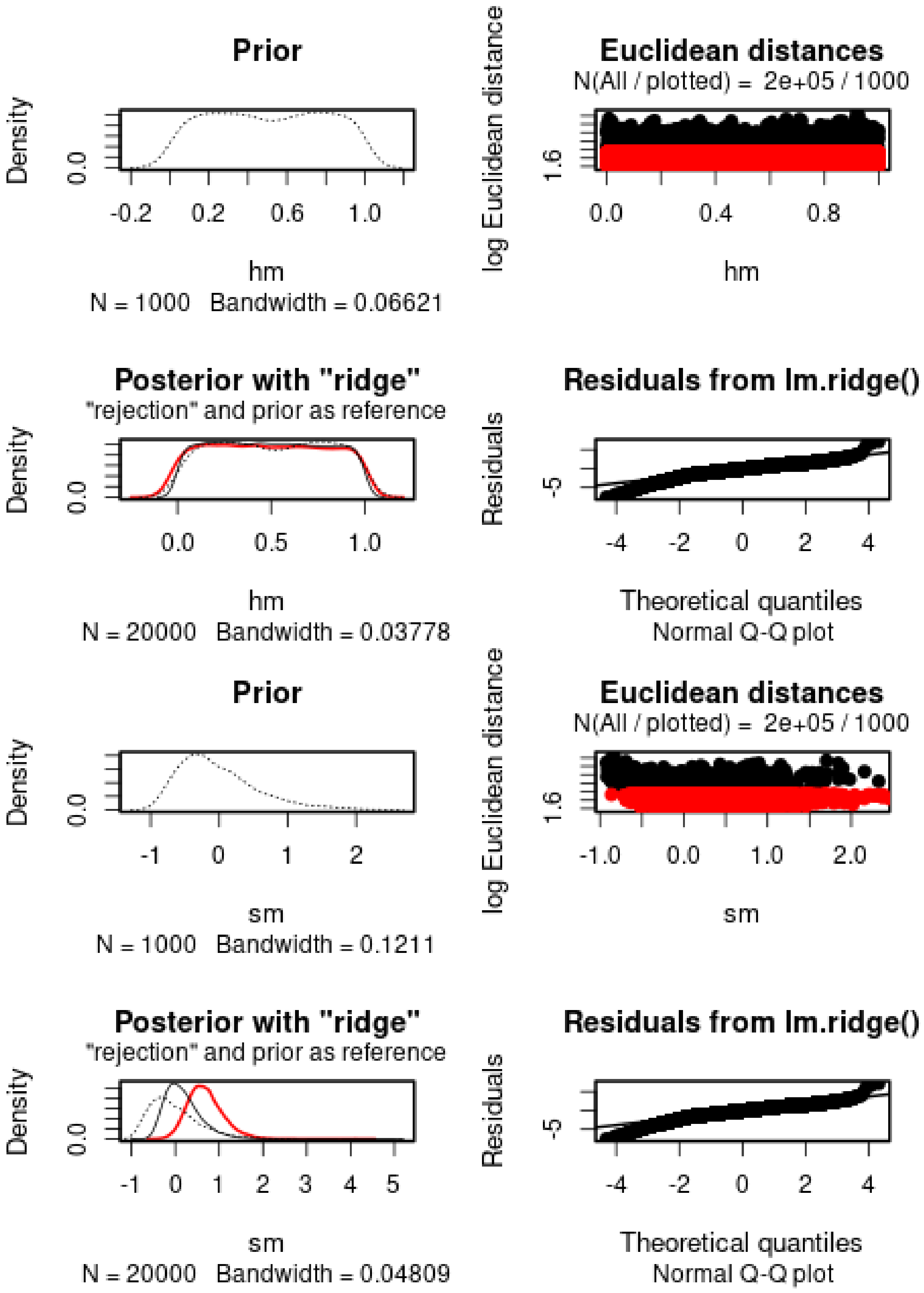
Prior
Density
0.0
-0.2
0.2
0.6
1.0
hm
N = 1000 Bandwidth = 0.06621
Euclidean distances
N(All / plotted) = 2e+05 / 1000
log Euclidean distance
1.6
0.0
0.4
0.8
hm
Posterior with "ridge"
"rejection" and prior as reference
Density
0.0
0.0
0.5
1.0
hm
N = 20000 Bandwidth = 0.03778
Residuals from lm.ridge()
Residuals
-5
-4
-2
0
2
4
Theoretical quantiles
Normal Q-Q plot
Prior
Density
0.0
-1
0
1
2
sm
N = 1000 Bandwidth = 0.1211
Euclidean distances
N(All / plotted) = 2e+05 / 1000
log Euclidean distance
1.6
-1.0
0.0
1.0
2.0
sm
Posterior with "ridge"
"rejection" and prior as reference
Density
0.0
-1
0
1
2
3
4
5
sm
N = 20000 Bandwidth = 0.04809
Residuals from lm.ridge()
Residuals
-5
-4
-2
0
2
4
Theoretical quantiles
Normal Q-Q plot

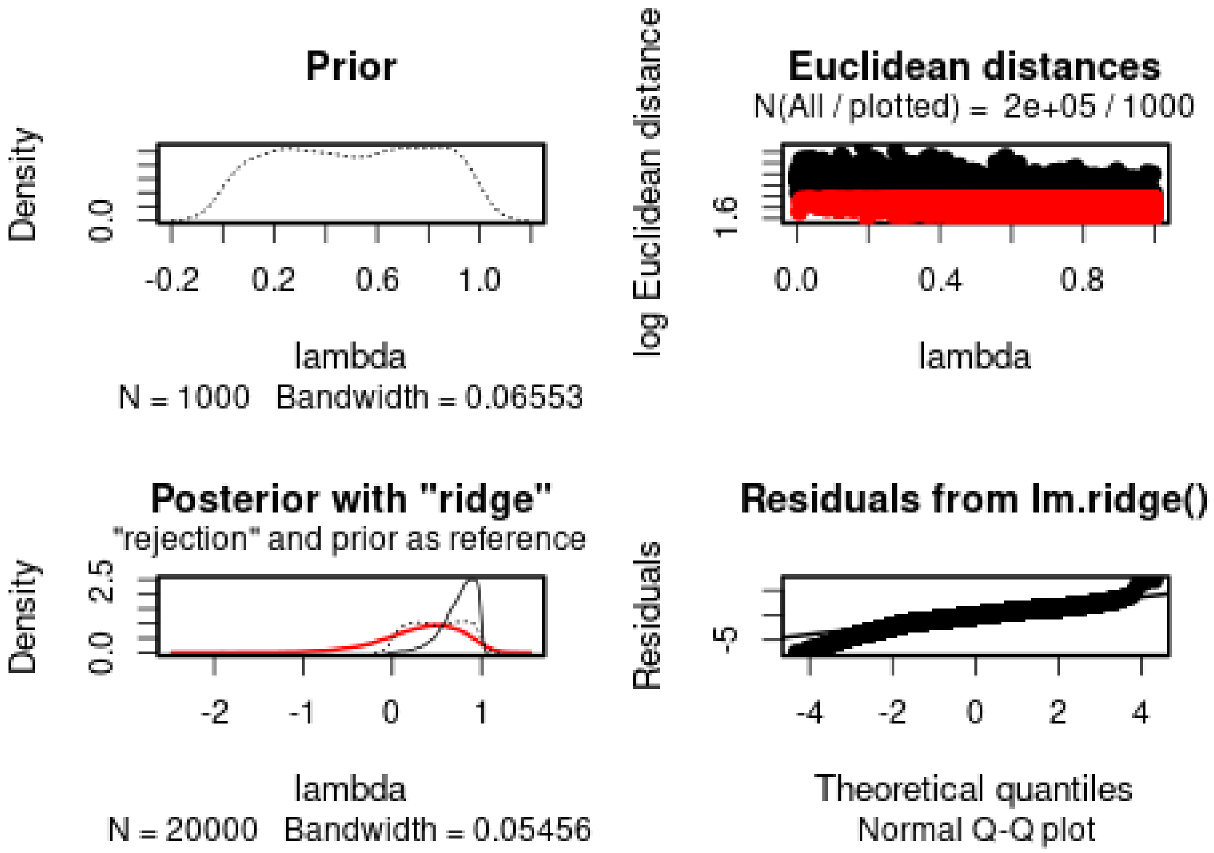

*h* cannot properly be estimated. *s* is positive (s = 0.6694), but has a large variance. The 95% posterior interval does not include 0. *lambda* is estimated to be 0.3890 (with a shift toward high values).

We then check the distribution of summary statistics in the simulations:

```
library(reshape2)
library(ggplot2)
library(ggpubr)

df.stats.sim <- melt(stat.sim, variable.name = "Statistic", value.name = "Frequency")
```

```
## No id variables; using all as measure variables
```

```
l <- strsplit(as.character(df.stats.sim$Statistic), split = ".", fixed = TRUE)
df.stats.sim$Room <- sapply(l, function(x) x[1])
df.stats.sim$Genotype <- sapply(l, function(x) x[2])
df.stats.sim$Generation <- sapply(
  strsplit(as.character(df.stats.sim$Statistic), split = ".", fixed = TRUE),
  function(x) x[3])

df.stats.obs <- as.data.frame(stat.obs)
names(df.stats.obs) <- "Frequency"
df.stats.obs$Statistic <- row.names(df.stats.obs)
l <- strsplit(as.character(df.stats.obs$Statistic), split = ".", fixed = TRUE)
df.stats.obs$Room <- sapply(l, function(x) x[1])
df.stats.obs$Genotype <- sapply(l, function(x) x[2])
df.stats.obs$Generation <- sapply(
  strsplit(as.character(df.stats.obs$Statistic), split = ".", fixed = TRUE),
  function(x) x[3])
```

```r
plot.obs <- function(gen) {
  p <- ggplot(data = subset(df.stats.sim, Generation == gen),
              aes(x = Frequency, y = ..density..)) +
    geom_histogram(bins = 50) +
    geom_vline(data = subset(df.stats.obs, Generation == gen),
               aes(xintercept = Frequency), color = "orange") +
    facet_grid(Genotype~Room) +
    theme_pubclean()
  return(p)
}
```

```r
plot.obs(1) + ggtitle("Timepoint 4")
```

```
## Warning: The dot-dot notation (`..density..`) was deprecated in ggplot2 3.4.0.
## i Please use `after_stat(density)` instead.
## This warning is displayed once every 8 hours.
## Call `lifecycle::last_lifecycle_warnings()` to see where this warning was
## generated.
```

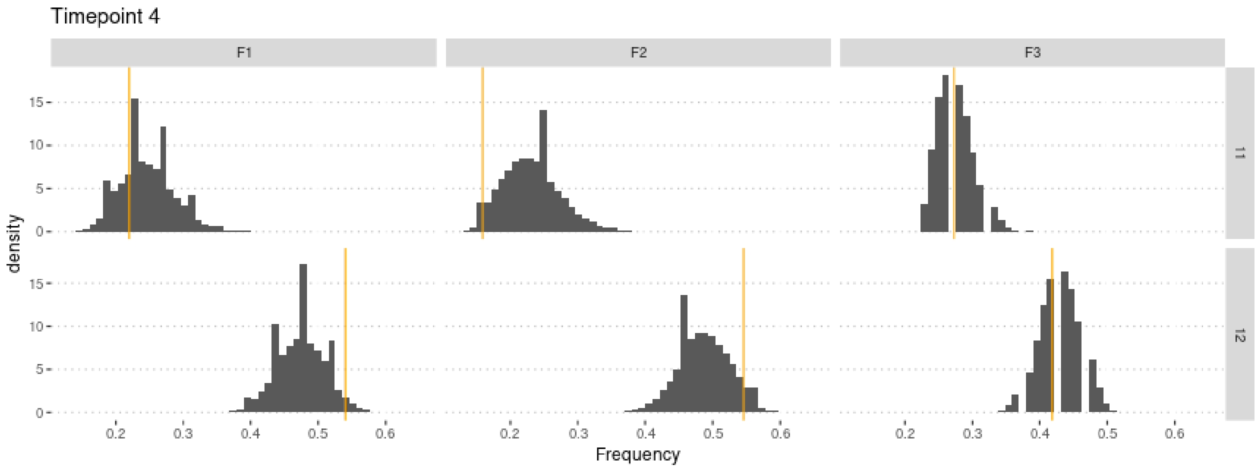

```r
plot.obs(2) + ggtitle("Timepoint 5")
```

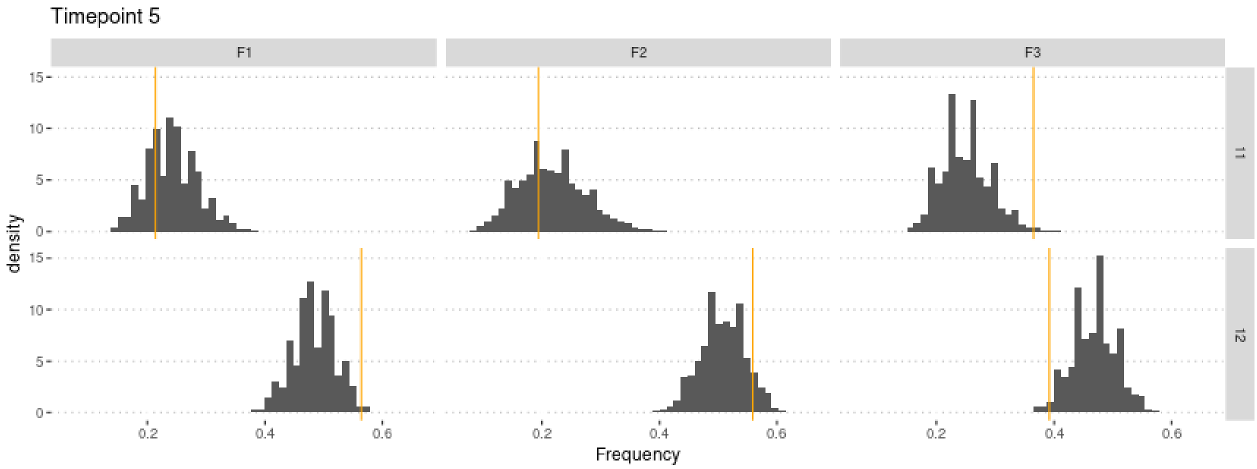

```
plot.obs(3) + ggtitle("Timepoint 6")
```

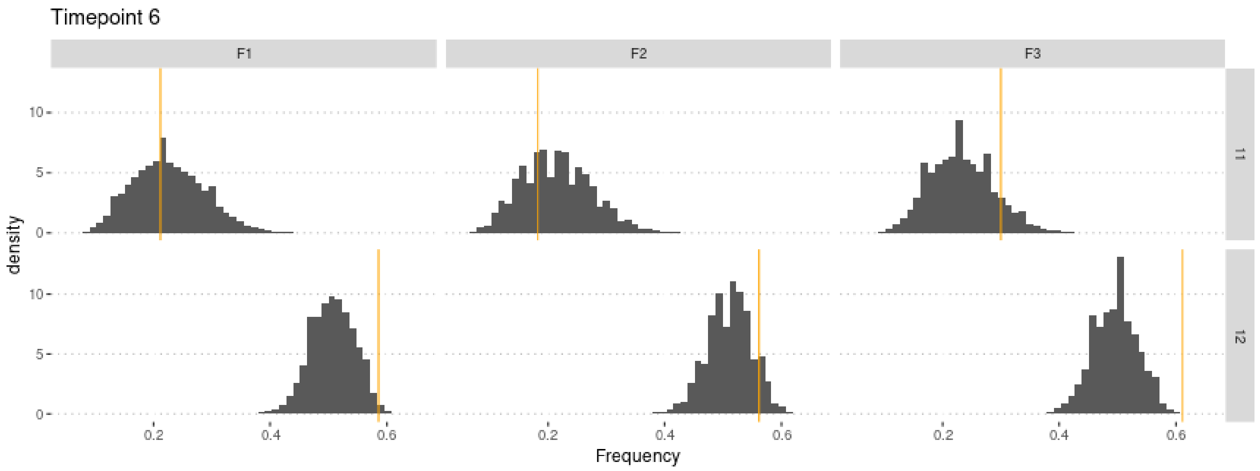

```
plot.obs(4) + ggtitle("Timepoint 7")
```

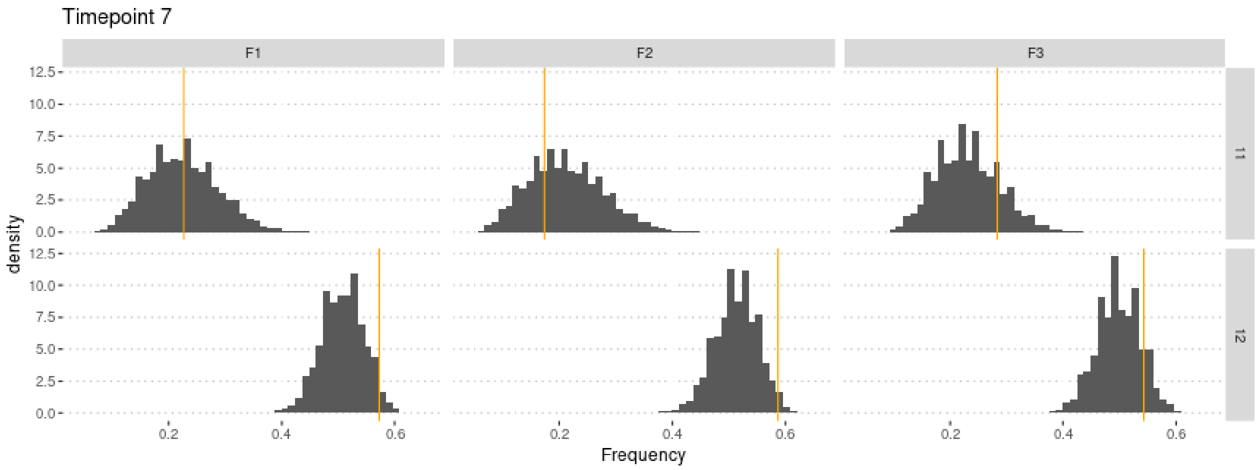

```
plot.obs(5) + ggtitle("Timepoint 8")
```

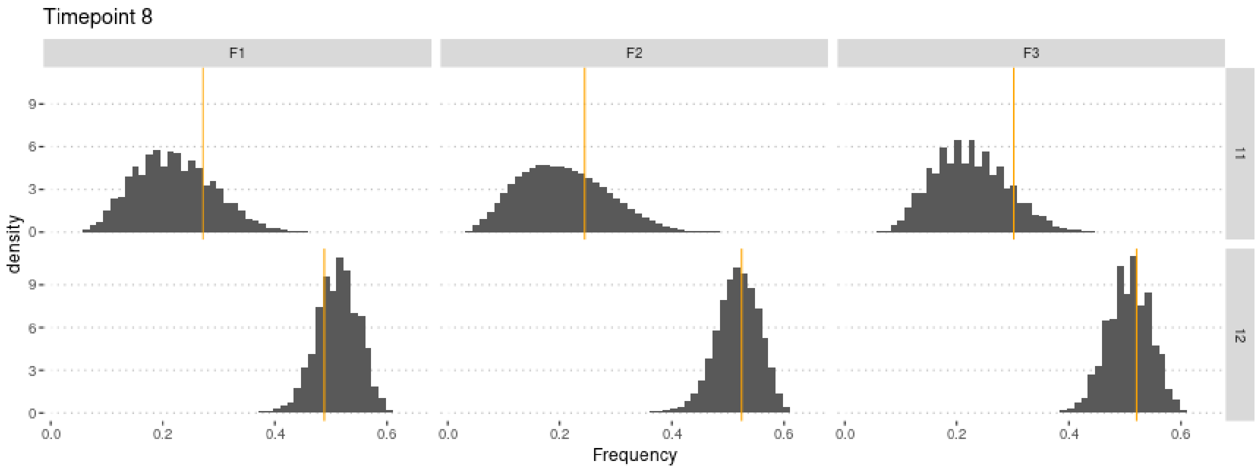

#### 4.1 Cross-Validation analysis

For memory-saving reasons, we only use 100,000 simulations in the following.

```r
sims2 <- sims[1:1e5,] #Not enough memory to use all simulations
param.sim2 <- param.sim[1:1e5,]
stat.sim2 <- stat.sim[1:1e5,]
m.r.g2 <- abc(target = stat.obs, param = param.sim2, sumstat = stat.sim2, tol = .1,
              method = "ridge", transf = c("none", "none", "none"))
```

Check that this does not affect the estimates too much:

```r
summary(m.r.g2, intvl = .95)
```

```
## Call:
## abc(target = stat.obs, param = param.sim2, sumstat = stat.sim2,
##     tol = 0.1, method = "ridge", transf = c("none", "none", "none"))
## Data:
##  abc.out$adj.values (10000 posterior samples)
## Weights:
##  abc.out$weights
##
##                              hm      sm  lambda
## Min.:                   -0.1115 -0.4983 -2.7073
## Weighted 2.5 % Perc.:    0.0168 -0.0233 -0.7487
## Weighted Median:         0.4834  0.5663  0.4168
## Weighted Mean:           0.4821  0.6004  0.3537
## Weighted Mode:           0.6748  0.5200  0.5093
## Weighted 97.5 % Perc.:   0.9432  1.4408  1.1106
## Max.:                    1.0870  4.1934  1.5592
```

Similar estimate, but slightly lower variance!

Compute predictions errors:

```r
if (redo) {
  cv.ridge.g <- cv4abc(
      param = param.sim2,
      sumstat = stat.sim2,
      abc.out = m.r.g2,
      nval = 1000,
      tols = c(.01, .1, .2))
  save(cv.ridge.g, file = "Rdata/MalesOnlyNonRandomMating/backup_cv_genotype.Rdata")
} else {
  load("Rdata/MalesOnlyNonRandomMating/backup_cv_genotype.Rdata")
}
summary(cv.ridge.g)
```

```
## Prediction error based on a cross-validation sample of 1000

##             hm        sm    lambda
## 0.01 1.0116961 0.1421933 0.2200749
## 0.1  0.9686981 0.1453140 0.2613666
## 0.2  0.9741045 0.1504224 0.2942033
```

```r
plot(cv.ridge.g)
```

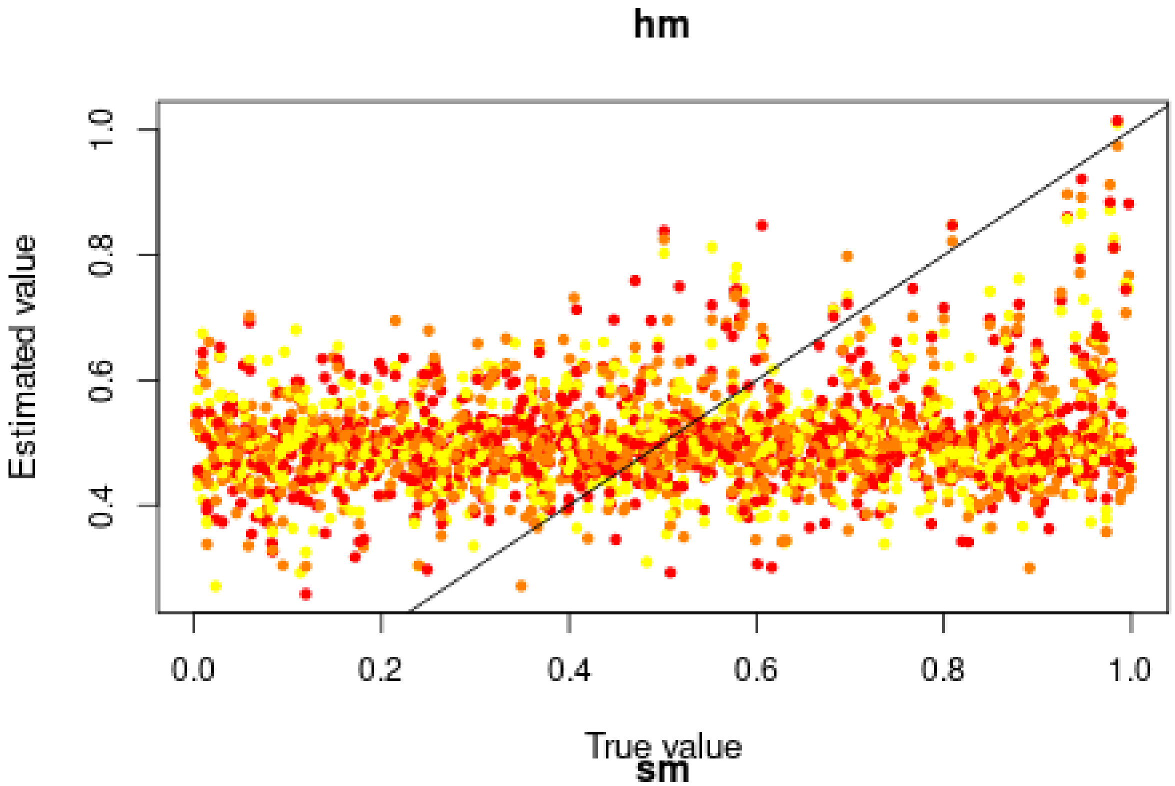
hm
Estimated value
1.0
0.8
0.6
0.4
0.0
0.2
0.4
0.6
0.8
1.0
True value

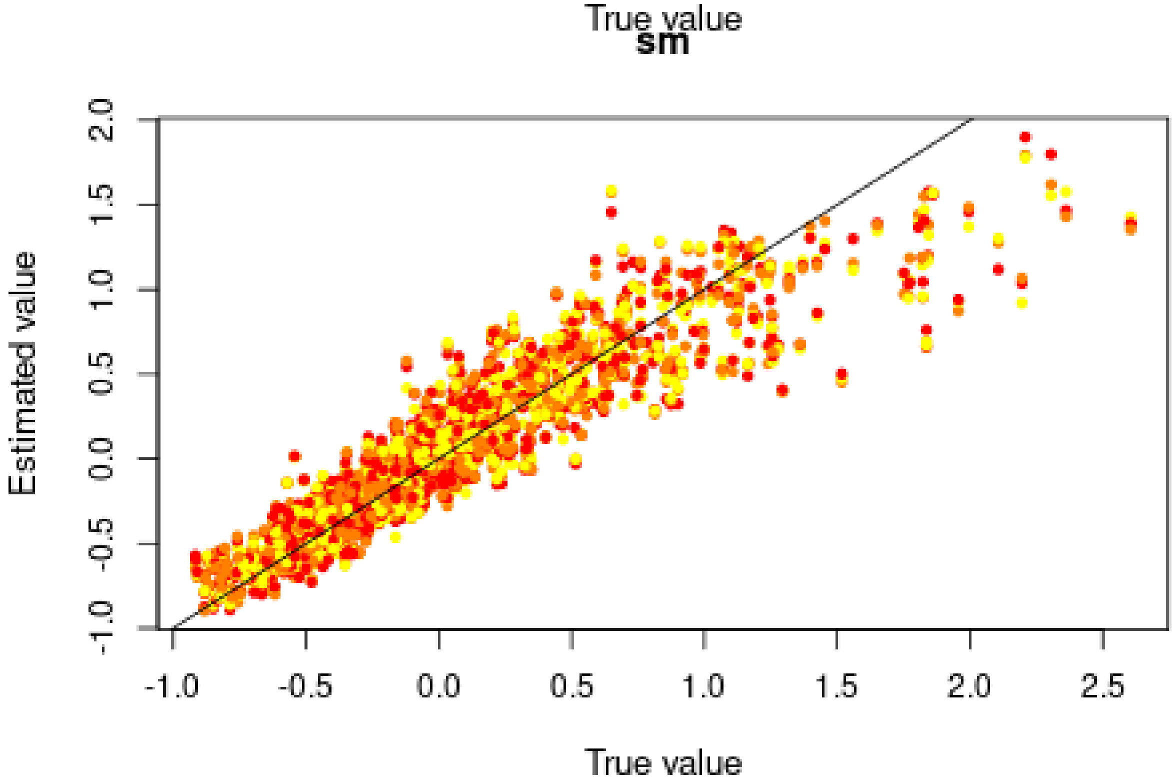
sm
Estimated value
2.0
1.5
1.0
0.5
0.0
-0.5
-1.0
-1.0
-0.5
0.0
0.5
1.0
1.5
2.0
2.5
True value

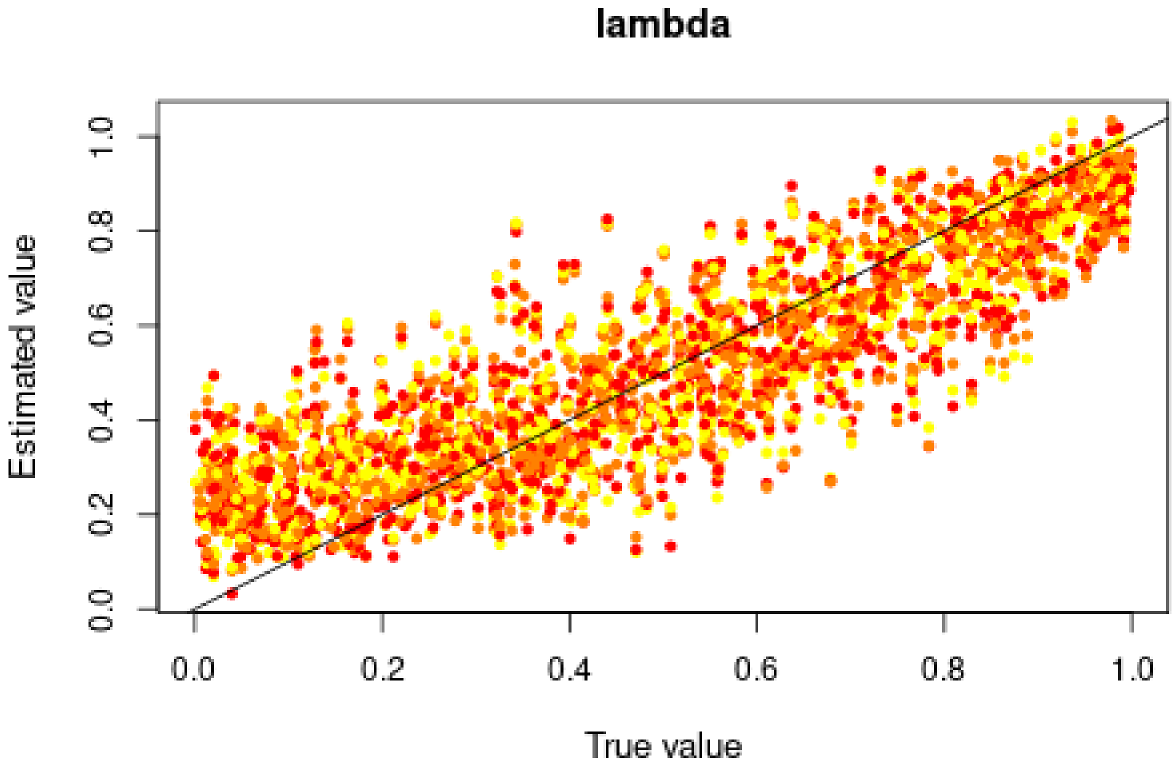

Good for *s*, but very bad for *h* and ok for *lambda*, in agreement with the posterior distributions.

##### 4.2 Misclassification errors

To test the power of the approach to distinguish between models, we also conduct a cross-validation experiment. We compare model M0 ($s = 0$) and M1 ($s = 0.6694 > 0$). $h$ is sampled over its prior distributions, and *lambda* is set to 0.3890. 100,000 simulations were conducted under each model.

```
sims0 <- read.csv("Simulations/Simulations0NonRandomMatingFromGen3.csv.gz")
param.sim0 <- subset(sims0, select = c("hm", "sm", "lambda"))
stat.sim0 <- subset(sims0, select = 2:46)
# Only consider the WT homozygotes and the heterozygotes,
# as the sum of the three genotypes is 1
stat.sim0 <- subset(stat.sim0, select = grep(".22.", names(stat.sim0), invert = TRUE))

sims1 <- read.csv("Simulations/Simulations1bNonRandomMatingMalesOnlyFromGen3.csv.gz")
param.sim1 <- subset(sims1, select = c("hm", "sm", "lambda"))
stat.sim1 <- subset(sims1, select = 2:46)
# Only consider the WT homozygotes and the heterozygotes,
# as the sum of the three genotypes is 1
stat.sim1 <- subset(stat.sim1, select = grep(".22.", names(stat.sim1), invert = TRUE))
```

We conduct a CV analysis (this takes some time… set nval = 100 for faster results):

```
models <- rep(c("Neutral", "Selection"), each = 100000)
if (redo) {
  cv.modsel.g <- cv4postpr(
    models, rbind(stat.sim0, stat.sim1),
    nval = 1000, tols = .1, method = "mnlogistic")
  save(cv.modsel.g, file = "Rdata/MalesOnlyNonRandomMating/backup_cv4postpr_genotype.Rdata")
} else {
```

```r
  load("Rdata/MalesOnlyNonRandomMating/backup_cv4postpr_genotype.Rdata")
}
```

We display the results:

```r
summary(cv.modsel.g)
```

```
## Confusion matrix based on 1000 samples for each model.
##
## $tol0.1
##           Neutral Selection
## Neutral       919        81
## Selection      73       927
##
##
## Mean model posterior probabilities (mnlogistic)
##
## $tol0.1
##           Neutral Selection
## Neutral    0.8854    0.1146
## Selection  0.1044    0.8956
```

```r
plot(cv.modsel.g, names.arg=c("Neutral", "Selection"))
```

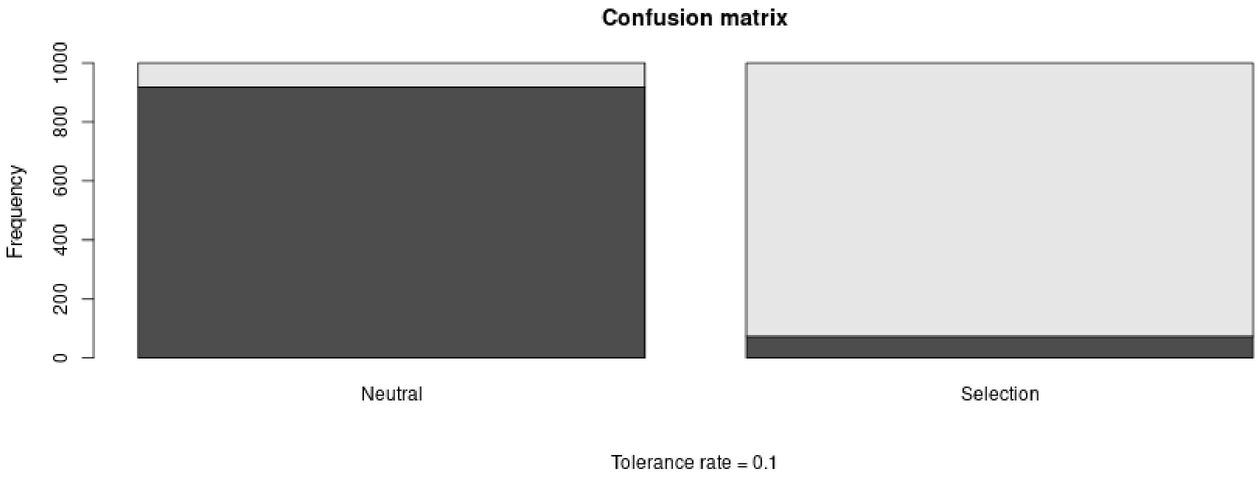

#### 4.3 Posterior prediction

```r
if (redo) {
  modsel.g <- postpr(
    stat.obs, models,
    rbind(stat.sim0, stat.sim1),
    tol = .1, method = "mnlogistic")
  save(modsel.g, file = "Rdata/MalesOnlyNonRandomMating/backup_postpr_genotype.Rdata")
} else {
  load("Rdata/MalesOnlyNonRandomMating/backup_postpr_genotype.Rdata")
}
summary(modsel.g)
```

```
## Call:
## postpr(target = stat.obs, index = models, sumstat = rbind(stat.sim0,
##     stat.sim1), tol = 0.1, method = "mnlogistic")
```

```
## Data:
##  postpr.out$values (20000 posterior samples)
## Models a priori:
##  Neutral, Selection
## Models a posteriori:
##  Neutral, Selection
##
## Proportion of accepted simulations (rejection):
##   Neutral Selection
##    0.6086    0.3914
##
## Bayes factors:
##           Neutral Selection
## Neutral    1.0000    1.5549
## Selection  0.6431    1.0000
##
##
## Posterior model probabilities (mnlogistic):
##   Neutral Selection
##    0.0651    0.9349
##
## Bayes factors:
##           Neutral Selection
## Neutral    1.0000    0.0696
## Selection 14.3692    1.0000
```

The model with selection is preferred.

##### 4.4 Goodness of fit

Under the neutral model:

```
if (redo) {
  res.gfit0.g <- gfit(
    target = stat.obs, sumstat = stat.sim0,
    statistic = median, nb.replicate = 1000)
  save(res.gfit0.g, file = "Rdata/MalesOnlyNonRandomMating/backup_gfit0_genotype.Rdata")
} else {
  load("Rdata/MalesOnlyNonRandomMating/backup_gfit0_genotype.Rdata")
}
plot(res.gfit0.g, main = "Histogram under M0")
```

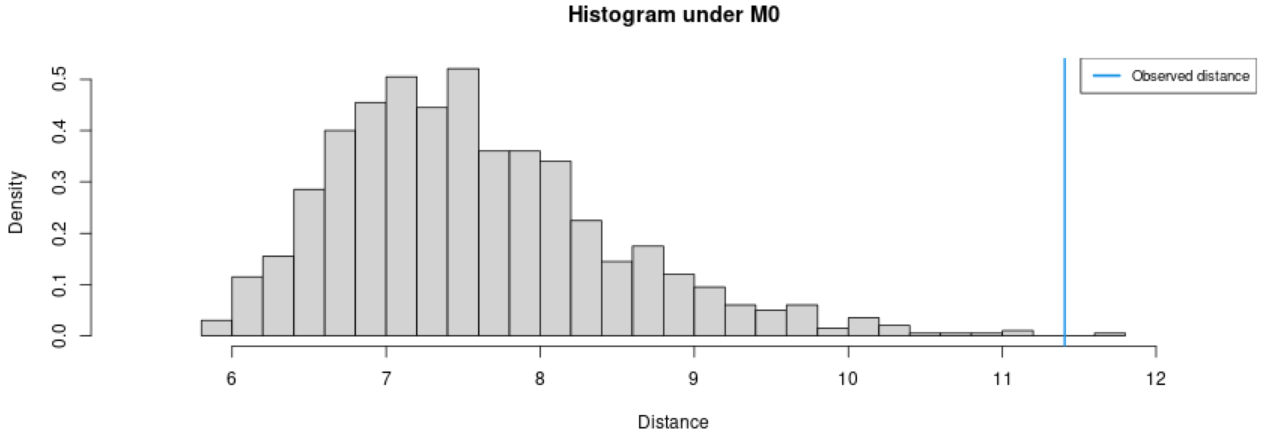

```r
summary(res.gfit0.g)
```

```
## $pvalue
## [1] 0.001
##
## $s.dist.sim
##    Min. 1st Qu.  Median    Mean 3rd Qu.    Max.
##   5.949   6.911   7.437   7.570   8.055  11.788
##
## $dist.obs
## [1] 11.40573
```

Under the selection model:

```r
if (redo) {
  res.gfit1.g <- gfit(
    target = stat.obs, sumstat = stat.sim1,
    statistic = median, nb.replicate = 1000)
  save(res.gfit1.g, file = "Rdata/MalesOnlyNonRandomMating/backup_gfit1_genotype.Rdata")
} else {
  load("Rdata/MalesOnlyNonRandomMating/backup_gfit1_genotype.Rdata")
}
plot(res.gfit1.g, main = "Histogram under M1")
```

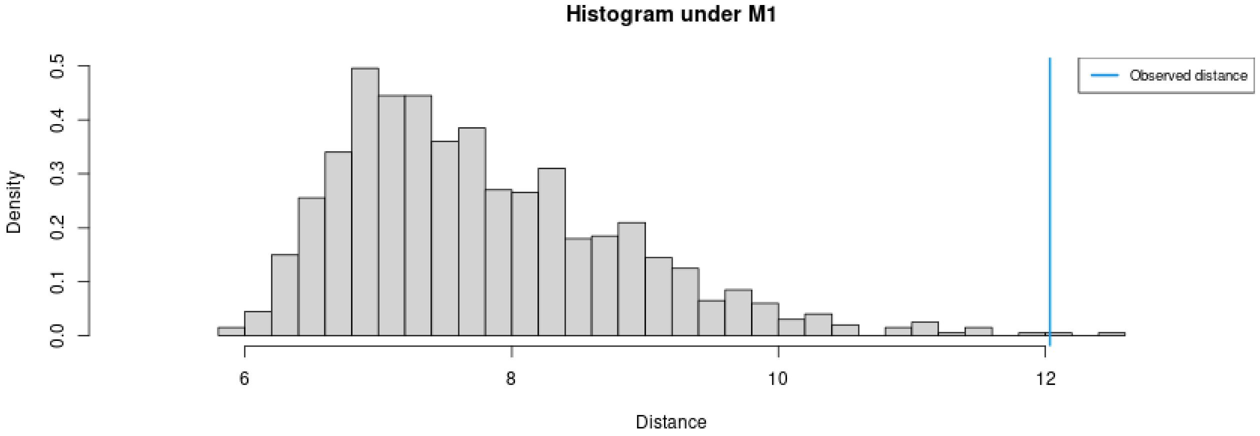

```r
summary(res.gfit1.g)
```

```
## $pvalue
## [1] 0.002
##
## $s.dist.sim
##    Min. 1st Qu.  Median    Mean 3rd Qu.    Max.
##   5.829   6.974   7.574   7.778   8.372  12.572
##
## $dist.obs
## [1] 12.03433
```

Both models provide a bad fit.

#### 4.5 Summary figure

Posterior distributions, with prior for comparison:

```
library(ggplot2)
library(ggpubr)
dat.prior.h <- data.frame(h = param.sim[,"hm"], Distribution = "Prior")
dat.post.h <- data.frame(h = m.r.g$adj.values[,"hm"], Distribution = "Posterior")
dat.prior.s <- data.frame(s = param.sim[,"sm"], Distribution = "Prior")
dat.post.s <- data.frame(s = m.r.g$adj.values[,"sm"], Distribution = "Posterior")
dat.prior.l <- data.frame(s = param.sim[,"lambda"], Distribution = "Prior")
dat.post.l <- data.frame(s = m.r.g$adj.values[,"lambda"], Distribution = "Posterior")
dat.h <- rbind(dat.prior.h, dat.post.h)
dat.s <- rbind(dat.prior.s, dat.post.s)
dat.l <- rbind(dat.prior.l, dat.post.l)
dat.h$Distribution <- factor(dat.h$Distribution, levels = c("Prior", "Posterior"))
dat.s$Distribution <- factor(dat.s$Distribution, levels = c("Prior", "Posterior"))
dat.l$Distribution <- factor(dat.l$Distribution, levels = c("Prior", "Posterior"))
names(dat.h)[1] <- "value"
dat.h$variable <- "h"
names(dat.s)[1] <- "value"
dat.s$variable <- "s"
names(dat.l)[1] <- "value"
dat.l$variable <- "lambda"
dat.dist <- rbind(dat.h, dat.s, dat.l)
dat.dist$variable <- factor(dat.dist$variable, levels = c("h", "s", "lambda"))

p.dist <- ggplot(data = dat.dist,
                 aes(x = value, linetype = Distribution, fill = Distribution)) +
          geom_density(alpha = 0.5) +
          scale_fill_brewer(type = "qual", palette = 3) +
          scale_linetype_manual(values = c(Prior = "dashed", Posterior = "solid")) +
          xlab("Parameter value") +
          facet_wrap(~variable, scales = "free") +
          ggtitle("Posterior and prior parameter distributions") +
          theme_pubclean() + theme(strip.background = element_blank())
```

Cross-validation:

```
d1<-rbind(as.data.frame(cv.ridge.g$estim$tol0.01),
          as.data.frame(cv.ridge.g$estim$tol0.1),
          as.data.frame(cv.ridge.g$estim$tol0.2))
d2<-rbind(cv.ridge.g$true, cv.ridge.g$true, cv.ridge.g$true)
d1<-melt(d1, value.name = "Estimated")
```

```
## No id variables; using all as measure variables
```

```
d1$Tolerance <- rep(c(0.01, 0.1, 0.2), each = 1000)
d1$Replicate <- rep(1:1000, 3)
d2<-melt(d2, value.name = "True")
```

```
## No id variables; using all as measure variables
```

```
d2$Replicate <- rep(1:1000, 3)
dat.cv <- merge(d1, d2, by = c("variable", "Replicate"))
dat.cv$variable <- factor(dat.cv$variable,
                          levels = c("hm", "sm", "lambda"),
                          labels = c("h", "s", "lambda"))
```

```r
p.cv <- ggplot(dat.cv, aes(x = True, y = Estimated)) +
        geom_point(aes(col = as.ordered(Tolerance))) +
        geom_abline(slope = 1) +
        facet_wrap(~variable, scales = "free") +
        theme_pubclean() + labs(color = "Tolerance") +
        ggtitle("Cross validation") +
        theme(strip.background = element_blank())
```

Confusion matrix:

```r
library(scales)
cv.sum <- summary(cv.modsel.g)
```

```
## Confusion matrix based on 1000 samples for each model.
##
## $tol0.1
##           Neutral Selection
## Neutral       919        81
## Selection      73       927
##
##
## Mean model posterior probabilities (mnlogistic)
##
## $tol0.1
##           Neutral Selection
## Neutral    0.8854    0.1146
## Selection  0.1044    0.8956
```

```r
dat.cv <- as.data.frame(cv.sum$conf.matrix$tol0.1/1000)
names(dat.cv) <- c("Real", "Inferred", "Frequency")
p.confmat <- ggplot(dat.cv, aes(x = Real, y = Frequency, fill = Inferred)) +
             geom_col() +
             scale_y_continuous(labels = scales::percent) +
             scale_fill_brewer(type = "qual", palette = 3) +
             ggtitle("Prediction errors") +
             theme_pubclean()
```

Model probabilities:

```r
p.mprob <- ggplot(as.data.frame(modsel.g$pred), aes(x = Var1, y = Freq, fill = Var1)) +
           geom_col() + ylab("Model posterior probability") + xlab("Model") +
           scale_fill_brewer(type = "qual", palette = 3) +
           theme_pubclean() +
           ggtitle("Model probabilities") +
           theme(legend.position = "none")
```

```r
library(cowplot)
```

```
##
## Attaching package: 'cowplot'

## The following object is masked from 'package:ggpubr':
##
##     get_legend
```

```r
p1 <- plot_grid(p.mprob, p.confmat, labels = c("C", "D"), nrow = 1)
p <- plot_grid(p.dist, p.cv, p1, labels = c("A", "B", ""), nrow = 3)
```

```r
p
```

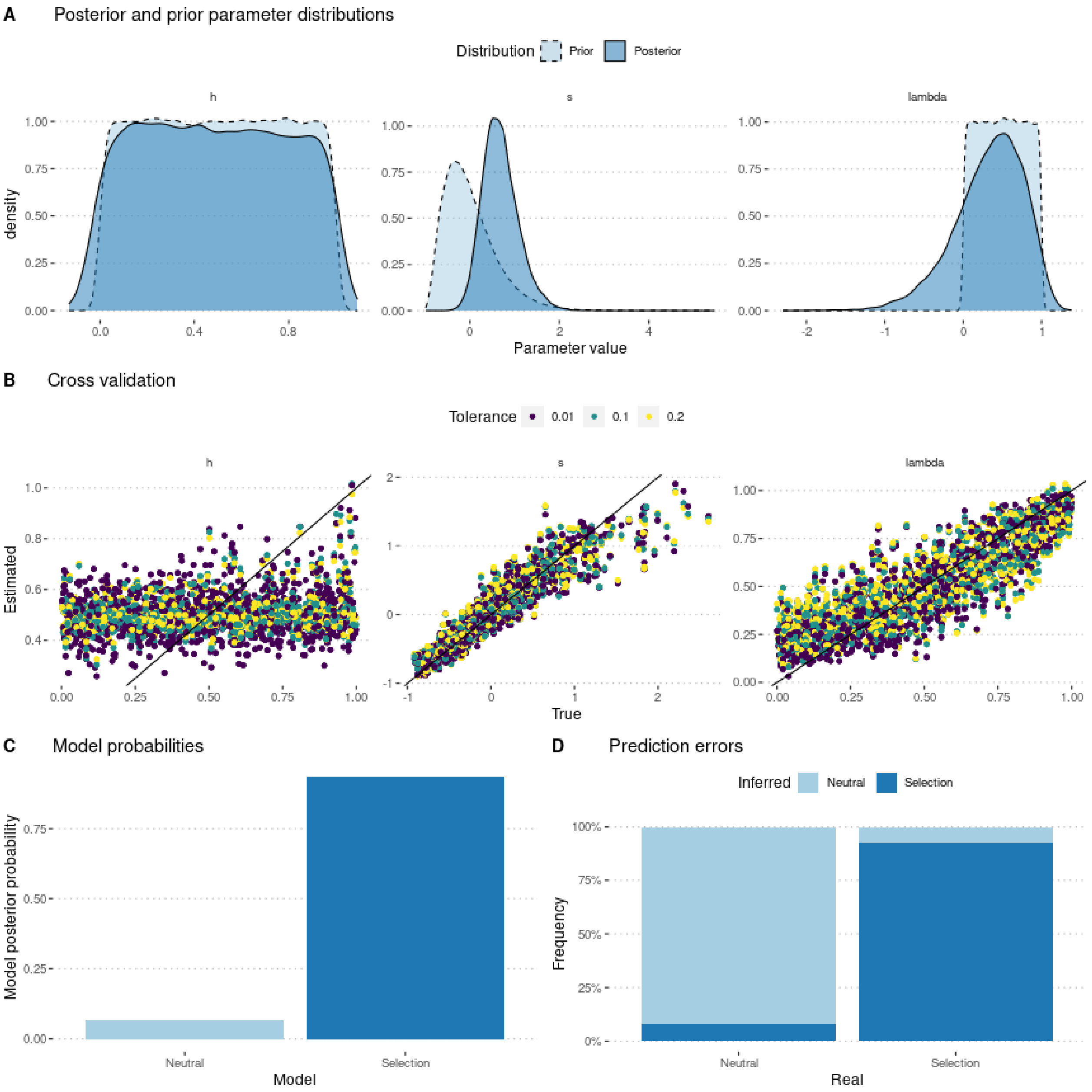

```r
ggsave(p, filename = "FigureABC-MalesOnlyNonRandomMating-Genotype.pdf",
       width = 8, height = 10)
```
