## Supplementary material for "Experimental evaluation of a direct fitness effect of the *de novo* evolved mouse gene *Pldi*": combined suppl files: SupplementaryFile7.pdf

### Estimating poldi's selection coefficient

Julien Y. Dutheil

08/03/2023

#### Contents

**1 Model parameterization** **1**

**2 Preamble** **1**

**3 Using allele frequencies** **2**

**4 Using genotype frequencies** **9**

### 1 Model parameterization

The model includes two parameters: $s$ the selection coefficient of the *poldi* allele compared to the knockout strain, which mimics the ancestral state, and $h$ the heterozygosity. Fitness values of each genotype are parameterized as follow:

$$\begin{aligned} poldi/poldi &: 1 \\ BL6/poldi &: 1 + h \cdot s \\ BL6/BL6 &: 1 + s \end{aligned}$$

where BL6 denotes the wild strain, and poldi the strain where the *poldi* gene is knocked out. Furthermore, we consider that the allele only has an effect in males, so that $h$ and $s$ are 0 in females (the two alleles have the same fitness). In this version, we model non random mating by adding three parameters $\{\lambda_1, \lambda_2, \lambda_3\}$, one for each replicate, so that parents are chosen according to their fitness multiplied by $(1 - \lambda)$ if they have the same two alleles, $(1 - \lambda/2)$ if they have only one allele in common, or 1 if both alleles are different.

We performed 300,000 simulations, using a gamma prior for $s$, a uniform prior between 0 and 1 for $h$, and a uniform prior between 0 and 1 for *lambda*.

### 2 Preamble

We load the data (observed and simulated):

```
sims <- read.csv("Simulations/SimulationsNonRandomMating3ParamMalesOnlyFromGen3.csv.gz")
param.sim <- subset(sims, select = c("hm", "sm", "lambda1", "lambda2", "lambda3"))
```

```r
stat.sim <- subset(sims, select = c(2:46))
# Only consider the WT homozygotes and the heterozygotes,
# as the sum of the three genotypes is 1
stat.sim <- subset(stat.sim, select = grep(".22.", names(stat.sim), invert = TRUE))

library(plyr, quietly = TRUE)
geno <- read.csv("GenotypeFrequencies.csv")
# We select the WT homozygote and heterozygote frequencies.
# Pldi homozygotes are 1 - sum(other two).
stat.obs <- subset(geno, TimePoint %in% 4:8, c("WT", "HET", "Replicate"))
stat.obs <- ddply(stat.obs, "Replicate", function(d) unlist(d[,1:2]))
stat.obs$Replicate <- NULL
stat.obs <- as.data.frame(t(stat.obs))
colnames(stat.obs) <- c("F1", "F2", "F3")
x <- rownames(stat.obs)
x <- gsub(x, pattern = "WT", replacement = ".11.")
x <- gsub(x, pattern = "HET", replacement = ".12.")
#x <- gsub(x, pattern = "Pldi", replacement = ".22.")
rownames(stat.obs) <- x
stat.obs1 <- subset(stat.obs, select = F1)
rownames(stat.obs1) <- paste0("F1", rownames(stat.obs1))
stat.obs2 <- subset(stat.obs, select = F2)
rownames(stat.obs2) <- paste0("F2", rownames(stat.obs2))
stat.obs3 <- subset(stat.obs, select = F3)
rownames(stat.obs3) <- paste0("F3", rownames(stat.obs3))
stat.obs <- cbind(t(stat.obs1), t(stat.obs2), t(stat.obs3))[1,]
```

### 3 Using allele frequencies

First, we need to compute the allelic frequencies from the genotype frequencies:

```r
compute.allelic.frequencies <- function(stat.sim) {
  stat.sim.a <- data.frame(row.names = row.names(stat.sim))
  #Pop1
  stat.sim.a$F1.1 <- stat.sim$F1.11.1 + stat.sim$F1.12.1 / 2
  stat.sim.a$F1.2 <- stat.sim$F1.11.2 + stat.sim$F1.12.2 / 2
  stat.sim.a$F1.3 <- stat.sim$F1.11.3 + stat.sim$F1.12.3 / 2
  stat.sim.a$F1.4 <- stat.sim$F1.11.4 + stat.sim$F1.12.4 / 2
  stat.sim.a$F1.5 <- stat.sim$F1.11.5 + stat.sim$F1.12.5 / 2
  #Pop2
  stat.sim.a$F2.1 <- stat.sim$F2.11.1 + stat.sim$F2.12.1 / 2
  stat.sim.a$F2.2 <- stat.sim$F2.11.2 + stat.sim$F2.12.2 / 2
  stat.sim.a$F2.3 <- stat.sim$F2.11.3 + stat.sim$F2.12.3 / 2
  stat.sim.a$F2.4 <- stat.sim$F2.11.4 + stat.sim$F2.12.4 / 2
  stat.sim.a$F2.5 <- stat.sim$F2.11.5 + stat.sim$F2.12.5 / 2
  #Pop3
  stat.sim.a$F3.1 <- stat.sim$F3.11.1 + stat.sim$F3.12.1 / 2
  stat.sim.a$F3.2 <- stat.sim$F3.11.2 + stat.sim$F3.12.2 / 2
  stat.sim.a$F3.3 <- stat.sim$F3.11.3 + stat.sim$F3.12.3 / 2
  stat.sim.a$F3.4 <- stat.sim$F3.11.4 + stat.sim$F3.12.4 / 2
  stat.sim.a$F3.5 <- stat.sim$F3.11.5 + stat.sim$F3.12.5 / 2

  return(stat.sim.a)
```

```
}
stat.sim.a <- compute.allelic.frequencies(stat.sim)
stat.obs.a <- numeric(15)
names(stat.obs.a) <- c("F1.1", "F1.2", "F1.3", "F1.4", "F1.5",
                       "F2.1", "F2.2", "F2.3", "F2.4", "F2.5",
                       "F3.1", "F3.2", "F3.3", "F3.4", "F3.5")
#Pop1
stat.obs.a[1]  <- stat.obs["F1.11.1"] + stat.obs["F1.12.1"] / 2
stat.obs.a[2]  <- stat.obs["F1.11.2"] + stat.obs["F1.12.2"] / 2
stat.obs.a[3]  <- stat.obs["F1.11.3"] + stat.obs["F1.12.3"] / 2
stat.obs.a[4]  <- stat.obs["F1.11.4"] + stat.obs["F1.12.4"] / 2
stat.obs.a[5]  <- stat.obs["F1.11.5"] + stat.obs["F1.12.5"] / 2
#Pop2
stat.obs.a[6]  <- stat.obs["F2.11.1"] + stat.obs["F2.12.1"] / 2
stat.obs.a[7]  <- stat.obs["F2.11.2"] + stat.obs["F2.12.2"] / 2
stat.obs.a[8]  <- stat.obs["F2.11.3"] + stat.obs["F2.12.3"] / 2
stat.obs.a[9]  <- stat.obs["F2.11.4"] + stat.obs["F2.12.4"] / 2
stat.obs.a[10] <- stat.obs["F2.11.5"] + stat.obs["F2.12.5"] / 2
#Pop3
stat.obs.a[11] <- stat.obs["F3.11.1"] + stat.obs["F3.12.1"] / 2
stat.obs.a[12] <- stat.obs["F3.11.2"] + stat.obs["F3.12.2"] / 2
stat.obs.a[13] <- stat.obs["F3.11.3"] + stat.obs["F3.12.3"] / 2
stat.obs.a[14] <- stat.obs["F3.11.4"] + stat.obs["F3.12.4"] / 2
stat.obs.a[15] <- stat.obs["F3.11.5"] + stat.obs["F3.12.5"] / 2
```

We estimate parameters using the ridge regression method:

```
library(abc, quietly = TRUE)
```

```
##
## Attaching package: 'SparseM'

## The following object is masked from 'package:base':
##
##     backsolve

## locfit 1.5-9.8    2023-06-11
```

```
if (redo) {
  m.r.a <- abc(target = stat.obs.a, param = param.sim, sumstat = stat.sim.a, tol=.1,
               method = "ridge", transf = c("none", "none", "none", "none", "none"))
  save(m.r.a, file = "Rdata/MalesOnlyNonRandomMating3Param/backup_abc_allelic.Rdata")
} else {
  load("Rdata/MalesOnlyNonRandomMating3Param/backup_abc_allelic.Rdata")
}
```

Now we display the results:

```
summary(m.r.a, intvl = .95)
```

```
## Call:
## abc(target = stat.obs.a, param = param.sim, sumstat = stat.sim.a,
##     tol = 0.1, method = "ridge", transf = c("none", "none", "none",
##         "none", "none"))
## Data:
##  abc.out$adj.values (30000 posterior samples)
## Weights:
```

```
##  abc.out$weights
##
##                               hm      sm lambda1 lambda2 lambda3
## Min.:                    -0.0771 -0.2591 -0.0398 -0.1148 -0.1702
## Weighted 2.5 % Perc.:     0.0133  0.1178  0.0607 -0.0187 -0.0704
## Weighted Median:          0.5098  0.5926  0.4869  0.4373  0.4192
## Weighted Mean:            0.5052  0.6249  0.4827  0.4395  0.4142
## Weighted Mode:            0.8110  0.5246  0.7633  0.3854  0.7558
## Weighted 97.5 % Perc.:    0.9855  1.3128  0.8957  0.9051  0.8893
## Max.:                     1.0540  4.0646  0.9645  1.0352  1.0171
```

Distribution of parameter estimates:

```
hist(m.r.a, breaks = 30, caption = c(expression(h), expression(s),
    expression(lambda[1]), expression(lambda[2]), expression(lambda[3])))
```

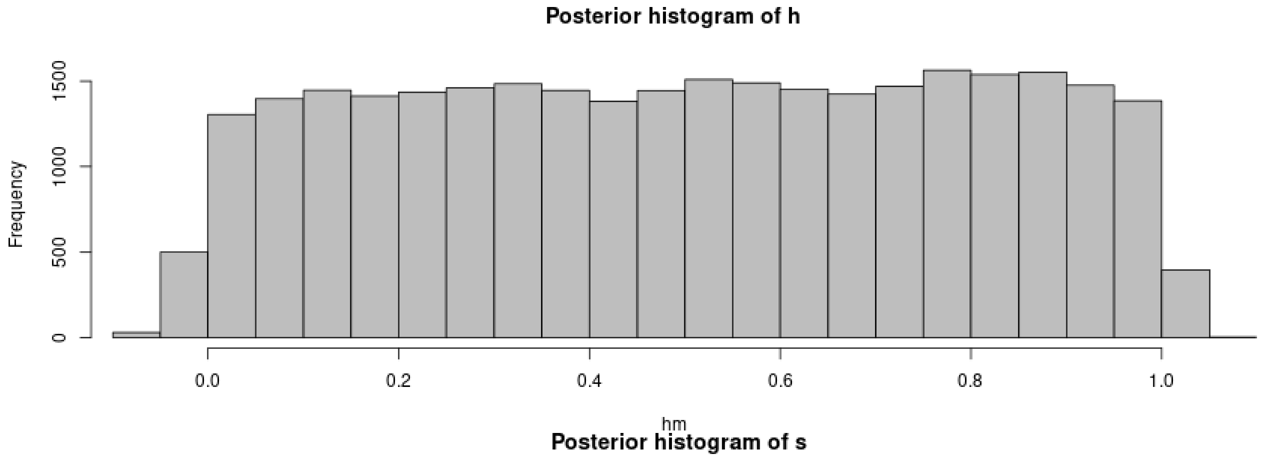

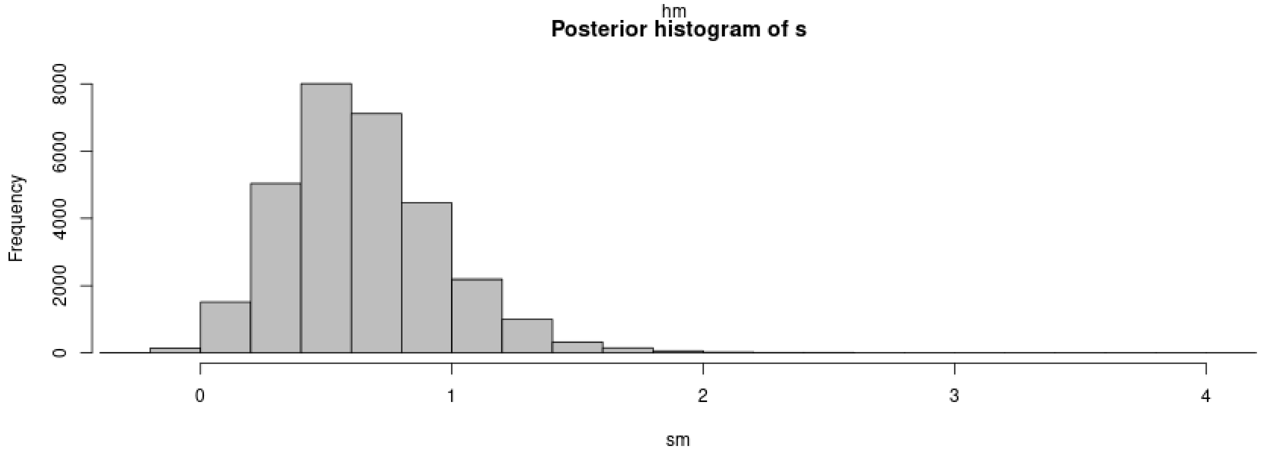

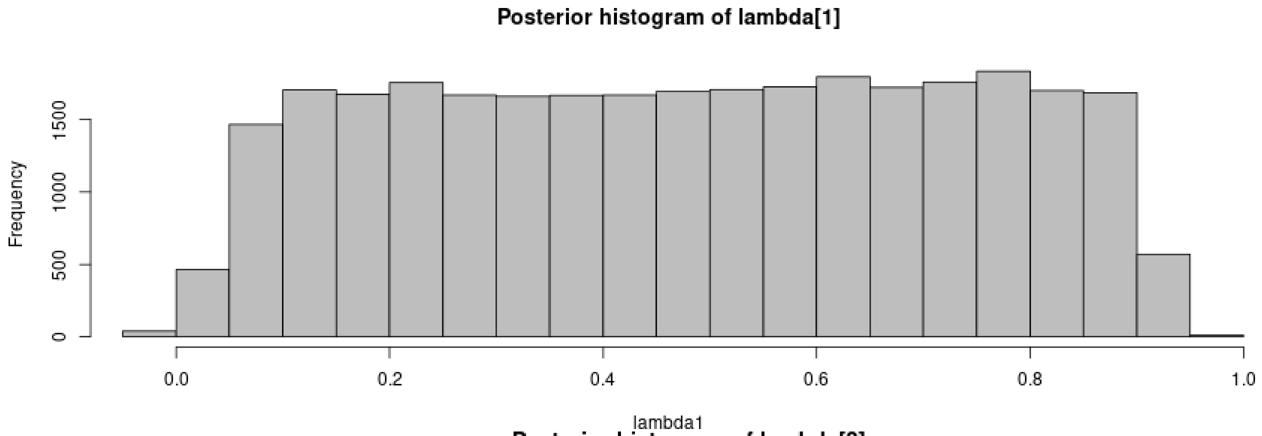

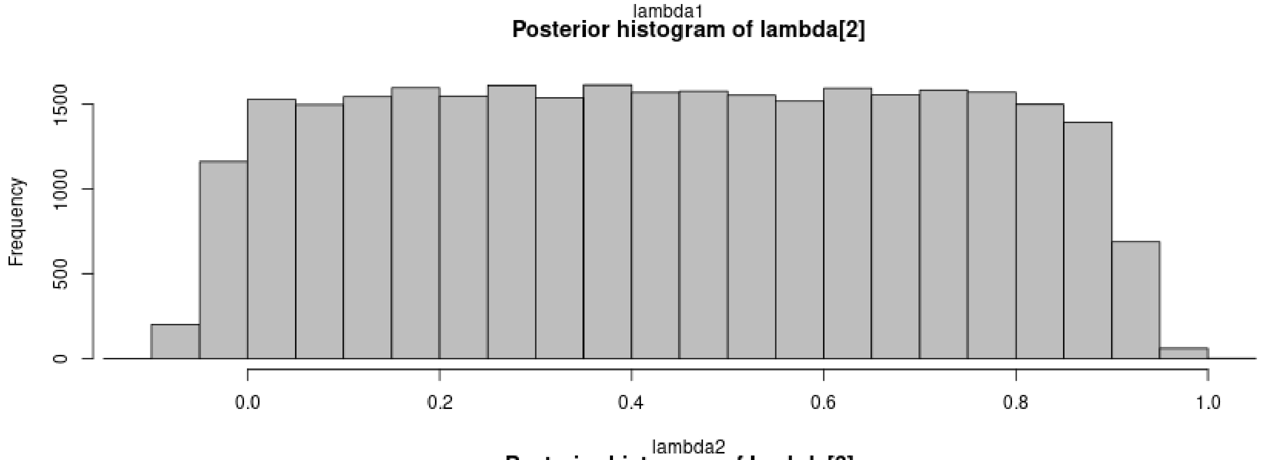

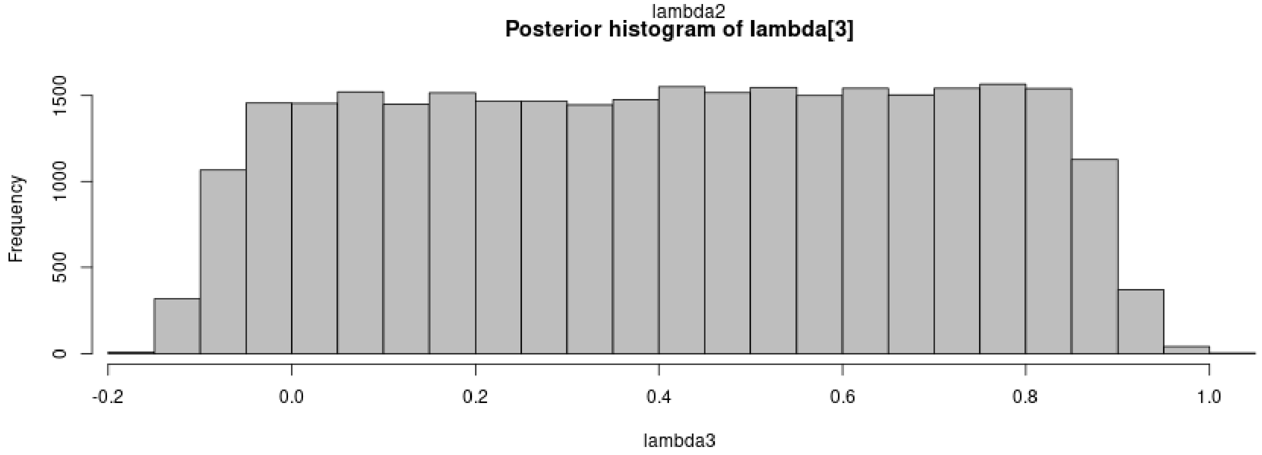

```
plot(m.r.a, param = param.sim)
```

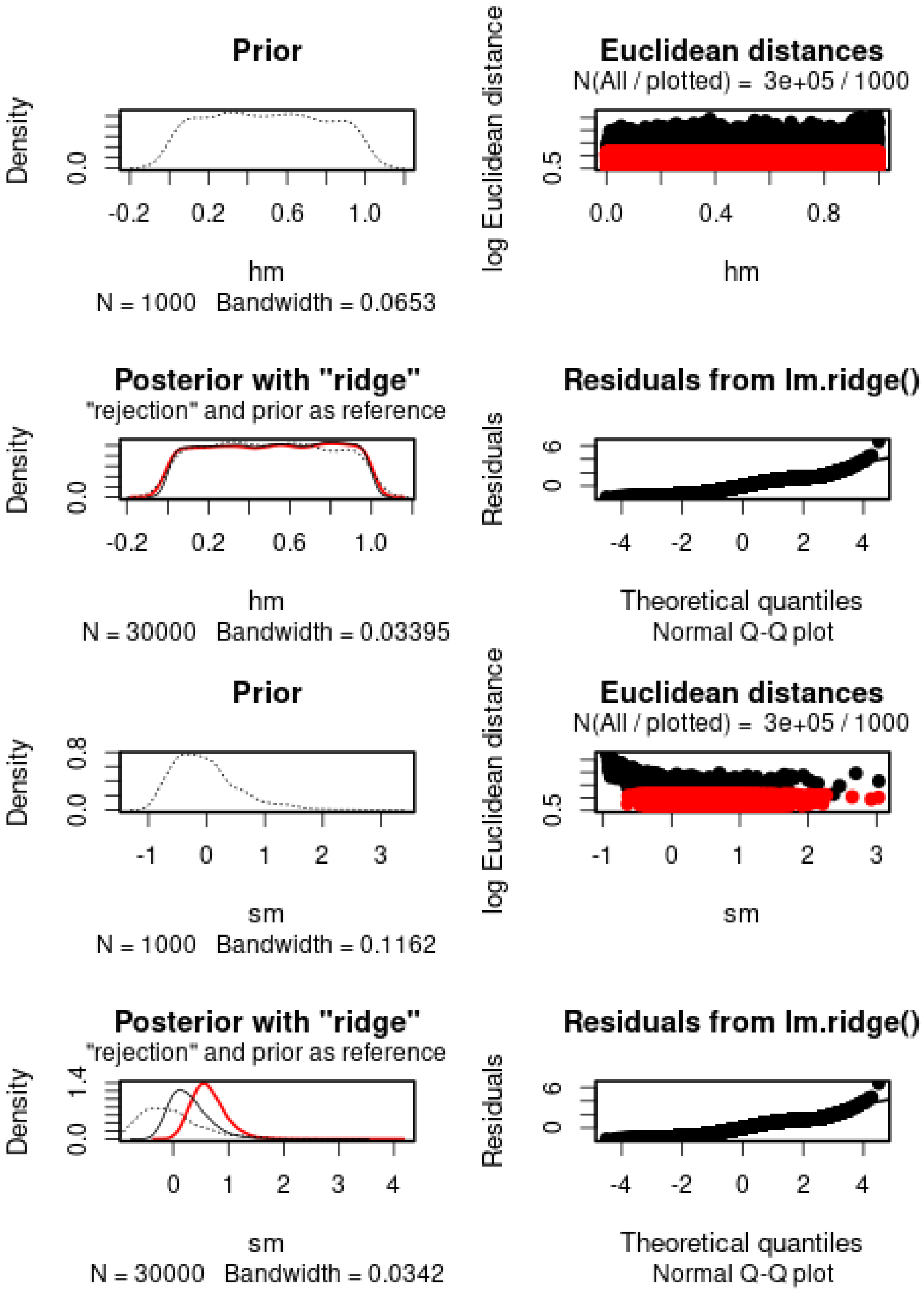
Prior
Density
0.0
-0.2 0.2 0.6 1.0
hm
N = 1000 Bandwidth = 0.0653
Euclidean distances
N(All / plotted) = 3e+05 / 1000
log Euclidean distance
0.5
0.0 0.4 0.8
hm
Posterior with "ridge"
"rejection" and prior as reference
Density
0.0
-0.2 0.2 0.6 1.0
hm
N = 30000 Bandwidth = 0.03395
Residuals from lm.ridge()
Residuals
0 6
-4 -2 0 2 4
Theoretical quantiles
Normal Q-Q plot
Prior
Density
0.0 0.8
-1 0 1 2 3
sm
N = 1000 Bandwidth = 0.1162
Euclidean distances
N(All / plotted) = 3e+05 / 1000
log Euclidean distance
0.5
-1 0 1 2 3
sm
Posterior with "ridge"
"rejection" and prior as reference
Density
0.0 1.4
0 1 2 3 4
sm
N = 30000 Bandwidth = 0.0342
Residuals from lm.ridge()
Residuals
0 6
-4 -2 0 2 4
Theoretical quantiles
Normal Q-Q plot

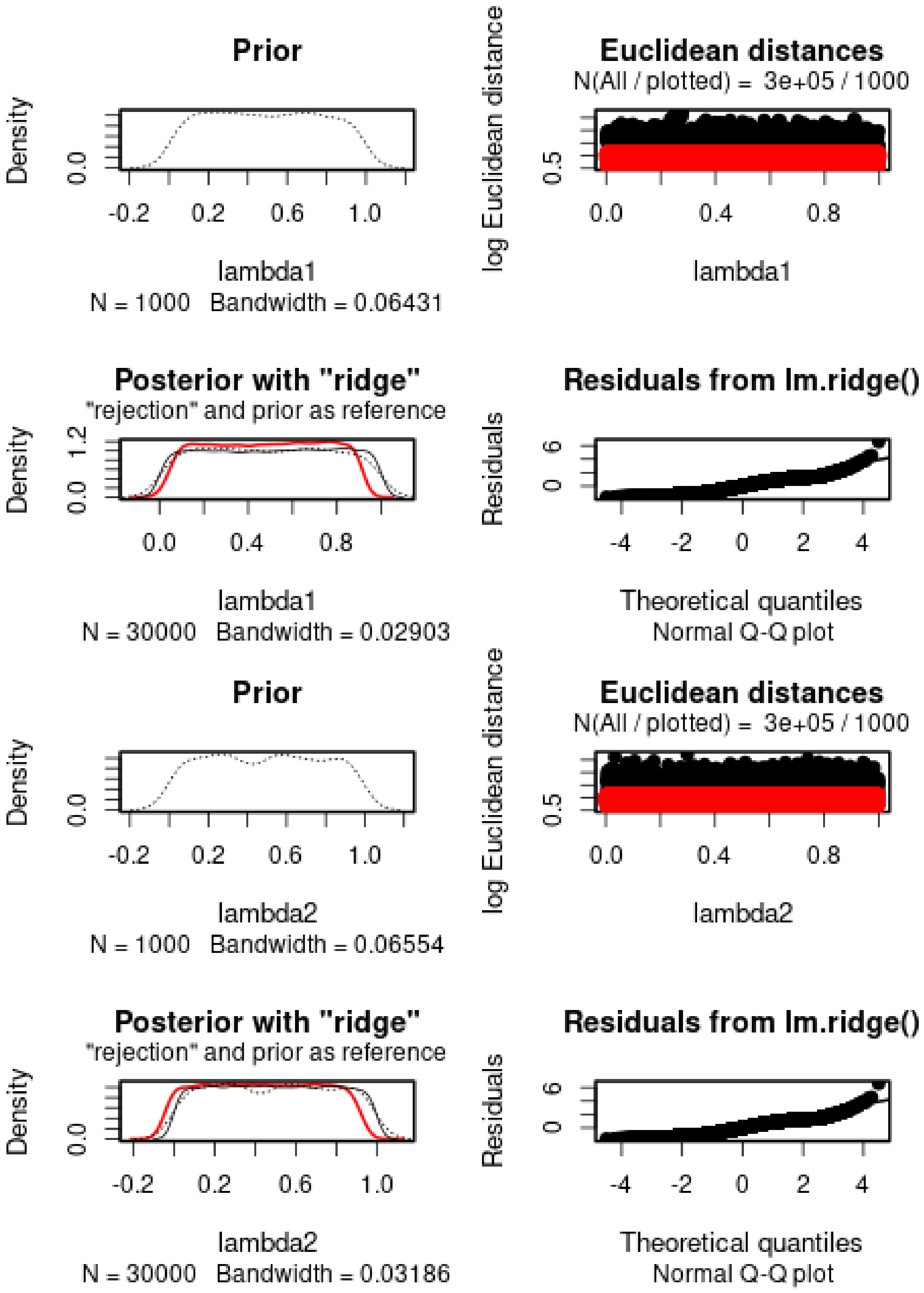

Prior
Density
0.0
-0.2
0.2
0.6
1.0
lambda1
N = 1000 Bandwidth = 0.06431
Euclidean distances
N(All / plotted) = 3e+05 / 1000
log Euclidean distance
0.5
0.0
0.4
0.8
lambda1
Posterior with "ridge"
"rejection" and prior as reference
Density
0.0
1.2
0.0
0.4
0.8
lambda1
N = 30000 Bandwidth = 0.02903
Residuals from lm.ridge()
Residuals
0
6
-4
-2
0
2
4
Theoretical quantiles
Normal Q-Q plot
Prior
Density
0.0
-0.2
0.2
0.6
1.0
lambda2
N = 1000 Bandwidth = 0.06554
Euclidean distances
N(All / plotted) = 3e+05 / 1000
log Euclidean distance
0.5
0.0
0.4
0.8
lambda2
Posterior with "ridge"
"rejection" and prior as reference
Density
0.0
-0.2
0.2
0.6
1.0
lambda2
N = 30000 Bandwidth = 0.03186
Residuals from lm.ridge()
Residuals
0
6
-4
-2
0
2
4
Theoretical quantiles
Normal Q-Q plot

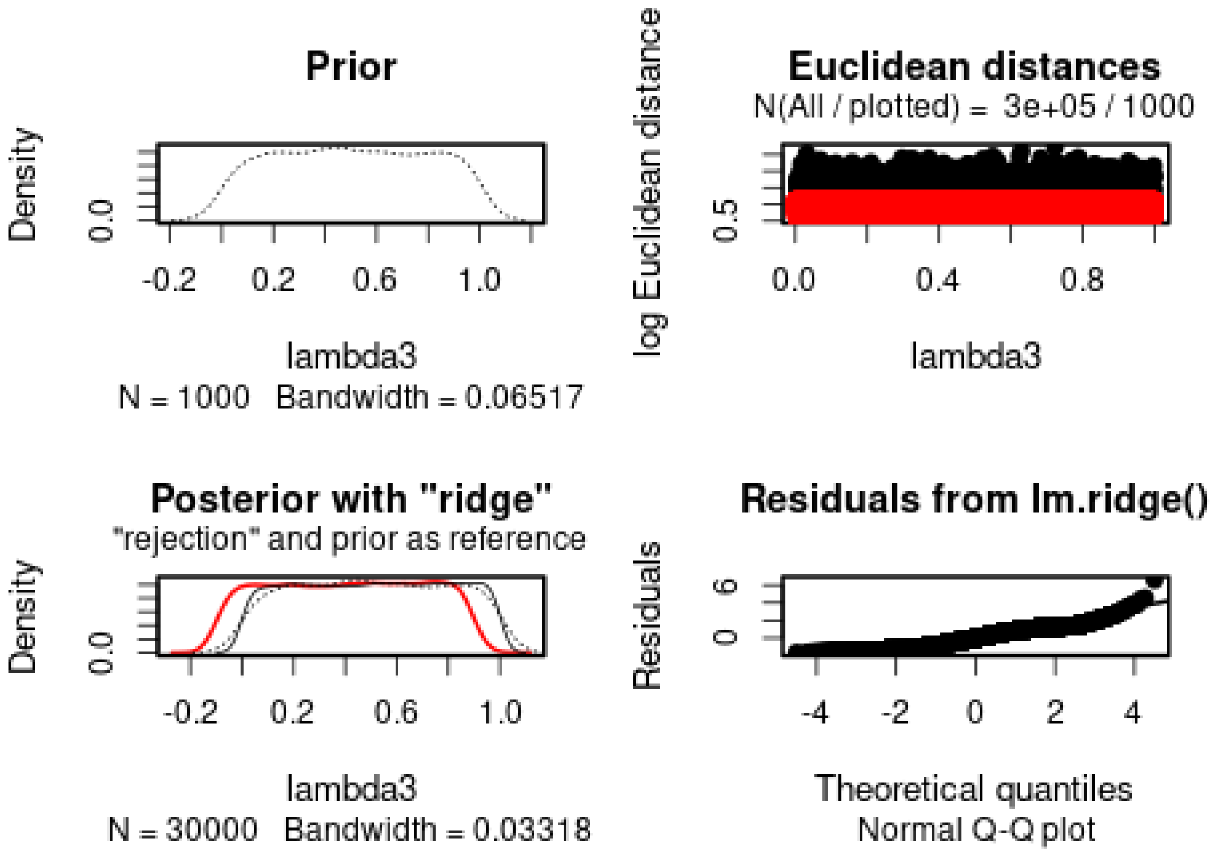

$h$ cannot properly be estimated. $s$ is positive and the 95% posterior interval does not include 0. Lambda cannot be estimated.

We then check the distribution of summary statistics in the simulations:

```
library(reshape2)
library(ggplot2)
library(ggpubr)
```

```
##
## Attaching package: 'ggpubr'

## The following object is masked from 'package:plyr':
##
##     mutate
```

```
df.stats.sim <- melt(stat.sim.a, variable.name = "Statistic", value.name = "Frequency")
```

```
## No id variables; using all as measure variables
```

```
l <- strsplit(as.character(df.stats.sim$Statistic), split = ".", fixed = TRUE)
df.stats.sim$Room <- sapply(l, function(x) x[1])
df.stats.sim$Generation <- sapply(
  strsplit(as.character(df.stats.sim$Statistic), split = ".", fixed = TRUE),
  function(x) x[2])

df.stats.obs <- as.data.frame(stat.obs.a)
names(df.stats.obs) <- "Frequency"
df.stats.obs$Statistic <- row.names(df.stats.obs)
l <- strsplit(as.character(df.stats.obs$Statistic), split = ".", fixed = TRUE)
```

```r
df.stats.obs$Room <- sapply(l, function(x) x[1])
df.stats.obs$Generation <- sapply(
  strsplit(as.character(df.stats.obs$Statistic), split = ".", fixed = TRUE),
  function(x) x[2])

p <- ggplot(data = df.stats.sim, aes(x = Frequency, y = after_stat(density))) +
       geom_histogram(bins = 50) +
       geom_vline(data = df.stats.obs, aes(xintercept = Frequency), color = "orange") +
       facet_grid(Room~Generation) +
       theme_pubclean()
p
```

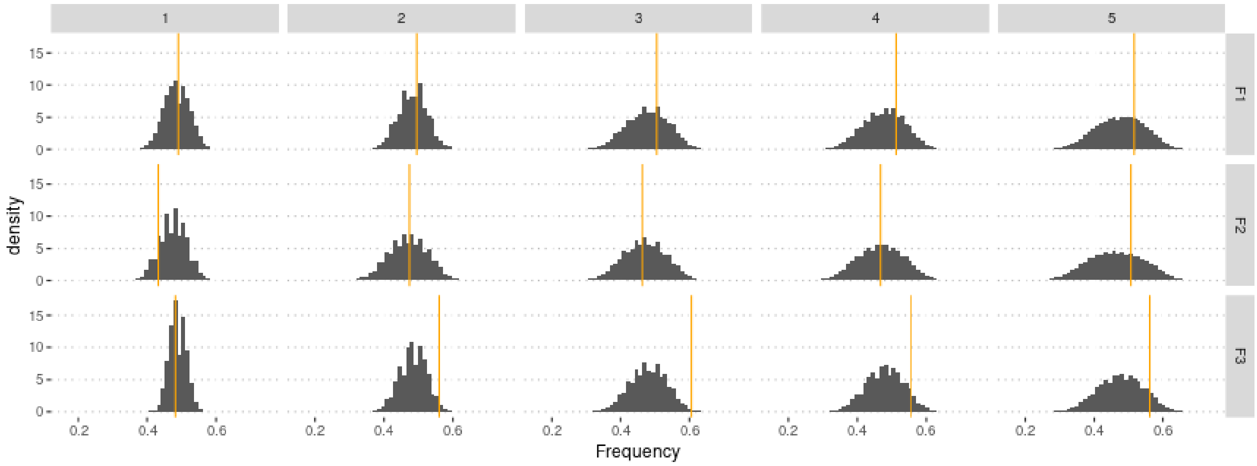

The *lambda* parameters are not identifiable when only allelic frequencies are used.

### 4 Using genotype frequencies

We estimate parameters using the ridge regression method:

```r
if (redo) {
  m.r.g <- abc(target = stat.obs, param = param.sim, sumstat = stat.sim, tol = .1,
               method = "ridge", transf = c("none", "none", "none", "none", "none"))
  save(m.r.g, file = "Rdata/MalesOnlyNonRandomMating3Param/backup_abc_genotype.Rdata")
} else {
  load("Rdata/MalesOnlyNonRandomMating3Param/backup_abc_genotype.Rdata")
}
```

Now we display the results:

```r
summary(m.r.g, intvl = .95)
```

```
## Call:
## abc(target = stat.obs, param = param.sim, sumstat = stat.sim,
##     tol = 0.1, method = "ridge", transf = c("none", "none", "none",
##         "none", "none"))
## Data:
##  abc.out$adj.values (30000 posterior samples)
## Weights:
##  abc.out$weights
##
##                              hm      sm lambda1 lambda2 lambda3
```

```
## Min.:                     -0.1032 -0.5254 -3.7586 -1.6424 -0.3732
## Weighted 2.5 % Perc.:      0.0146 -0.0067 -0.9672 -0.2827  0.0504
## Weighted Median:           0.4679  0.5912  0.3284  0.5342  0.4487
## Weighted Mean:             0.4714  0.6228  0.2524  0.4916  0.4350
## Weighted Mode:             0.2391  0.5926  0.4578  0.6181  0.5071
## Weighted 97.5 % Perc.:     0.9378  1.4528  1.0500  1.0265  0.7485
## Max.:                      1.0542  4.3889  1.3458  1.2782  0.9979
```

Distribution of parameter estimates:

```r
hist(m.r.g, breaks = 30, caption = c(expression(h), expression(s),
    expression(lambda[1]), expression(lambda[2]), expression(lambda[3])))
```

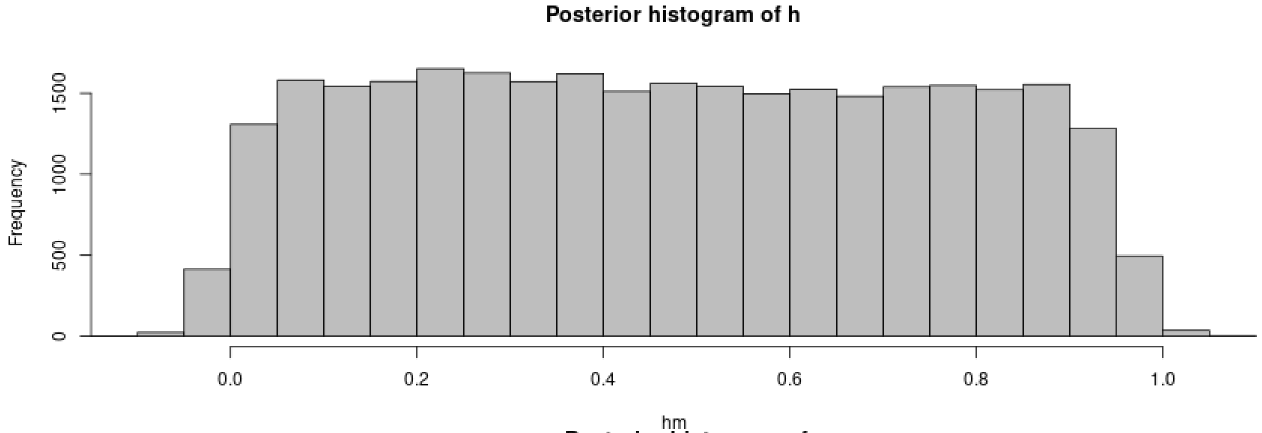

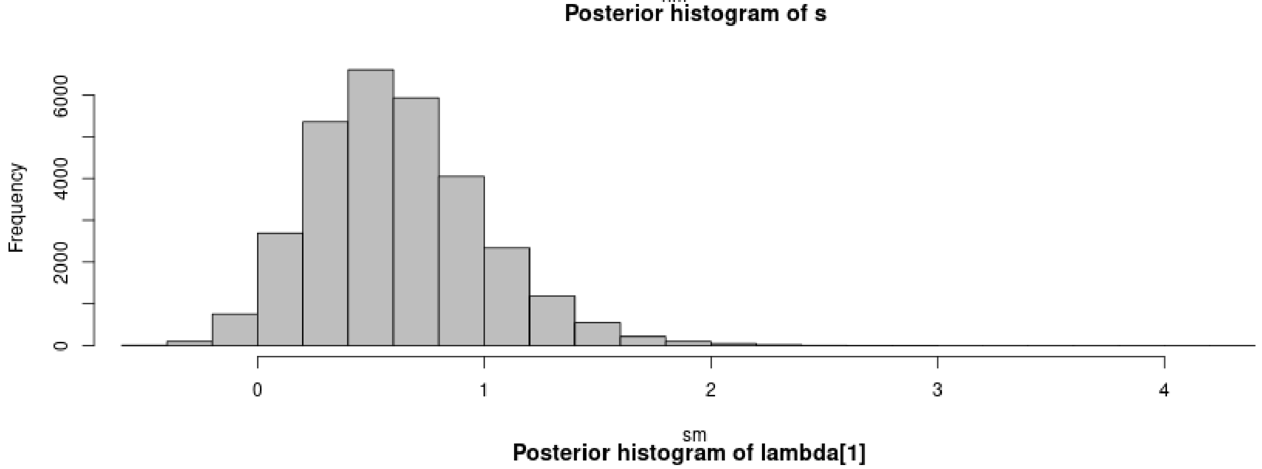

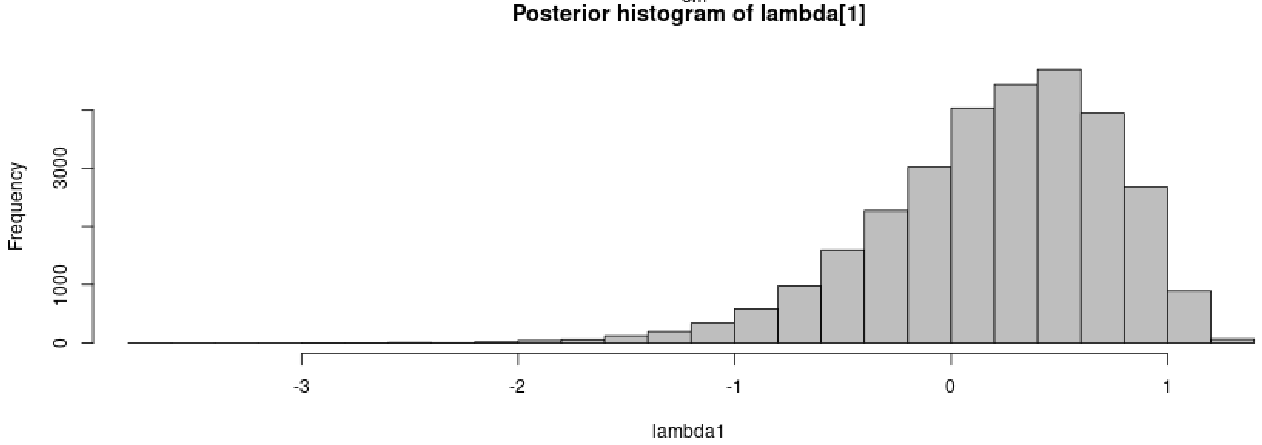

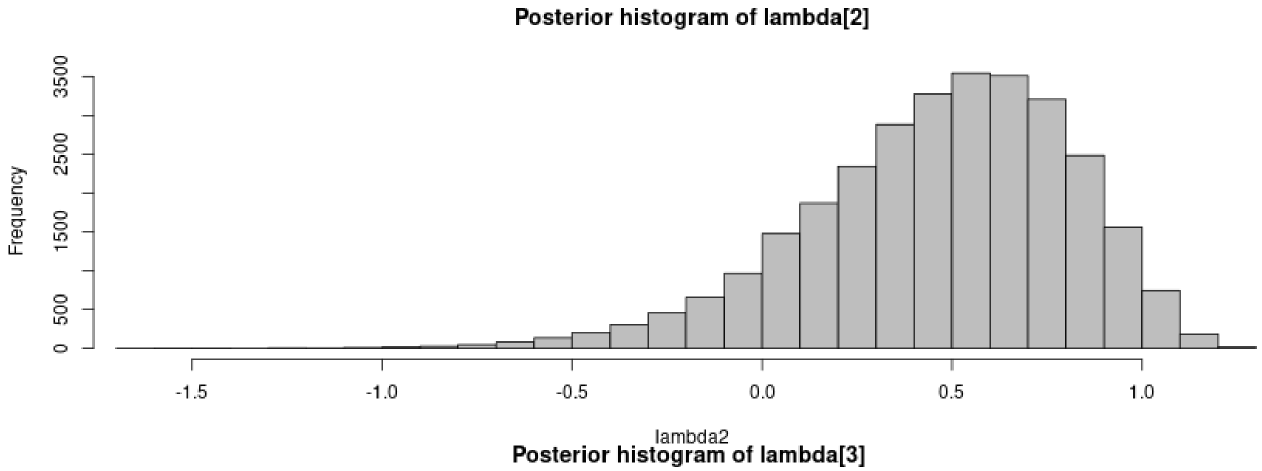

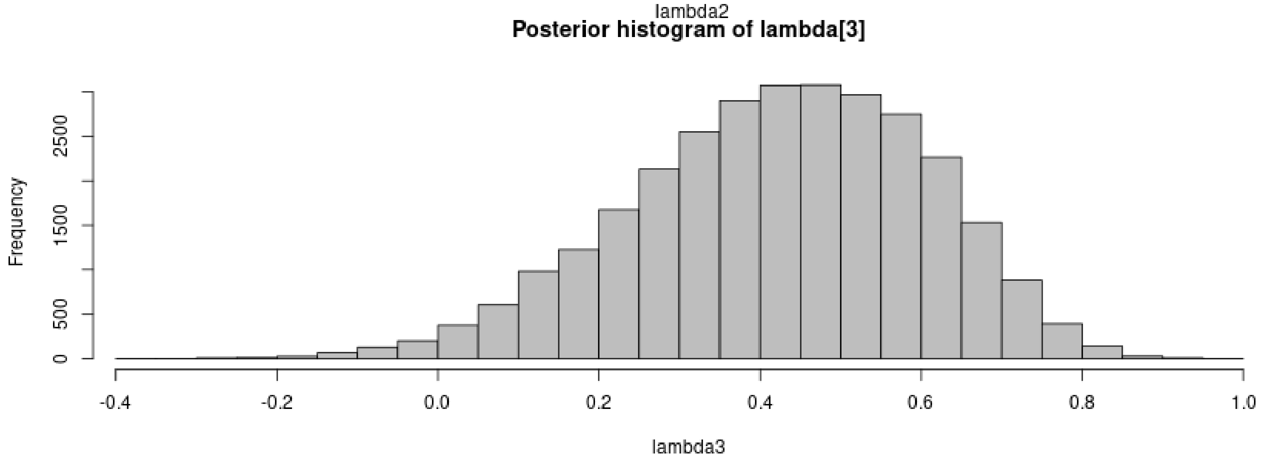

```
plot(m.r.g, param = param.sim)
```

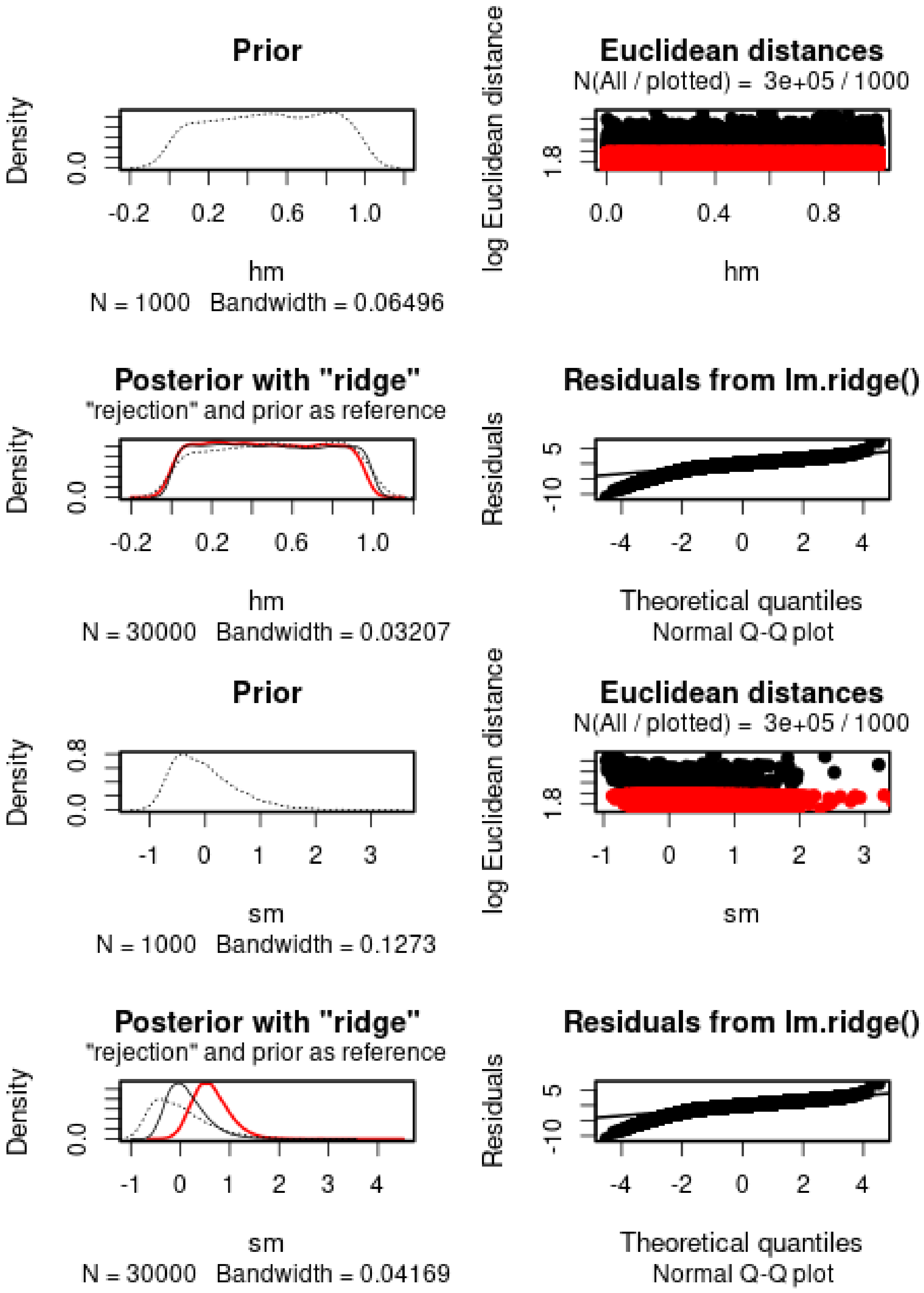
Prior
Density
0.0
-0.2
0.2
0.6
1.0
hm
N = 1000 Bandwidth = 0.06496
Euclidean distances
N(All / plotted) = 3e+05 / 1000
log Euclidean distance
1.8
0.0
0.4
0.8
hm
Posterior with "ridge"
"rejection" and prior as reference
Density
0.0
-0.2
0.2
0.6
1.0
hm
N = 30000 Bandwidth = 0.03207
Residuals from lm.ridge()
Residuals
5
-10
-4
-2
0
2
4
Theoretical quantiles
Normal Q-Q plot
Prior
Density
0.0
0.8
-1
0
1
2
3
sm
N = 1000 Bandwidth = 0.1273
Euclidean distances
N(All / plotted) = 3e+05 / 1000
log Euclidean distance
1.8
-1
0
1
2
3
sm
Posterior with "ridge"
"rejection" and prior as reference
Density
0.0
-1
0
1
2
3
4
sm
N = 30000 Bandwidth = 0.04169
Residuals from lm.ridge()
Residuals
5
-10
-4
-2
0
2
4
Theoretical quantiles
Normal Q-Q plot

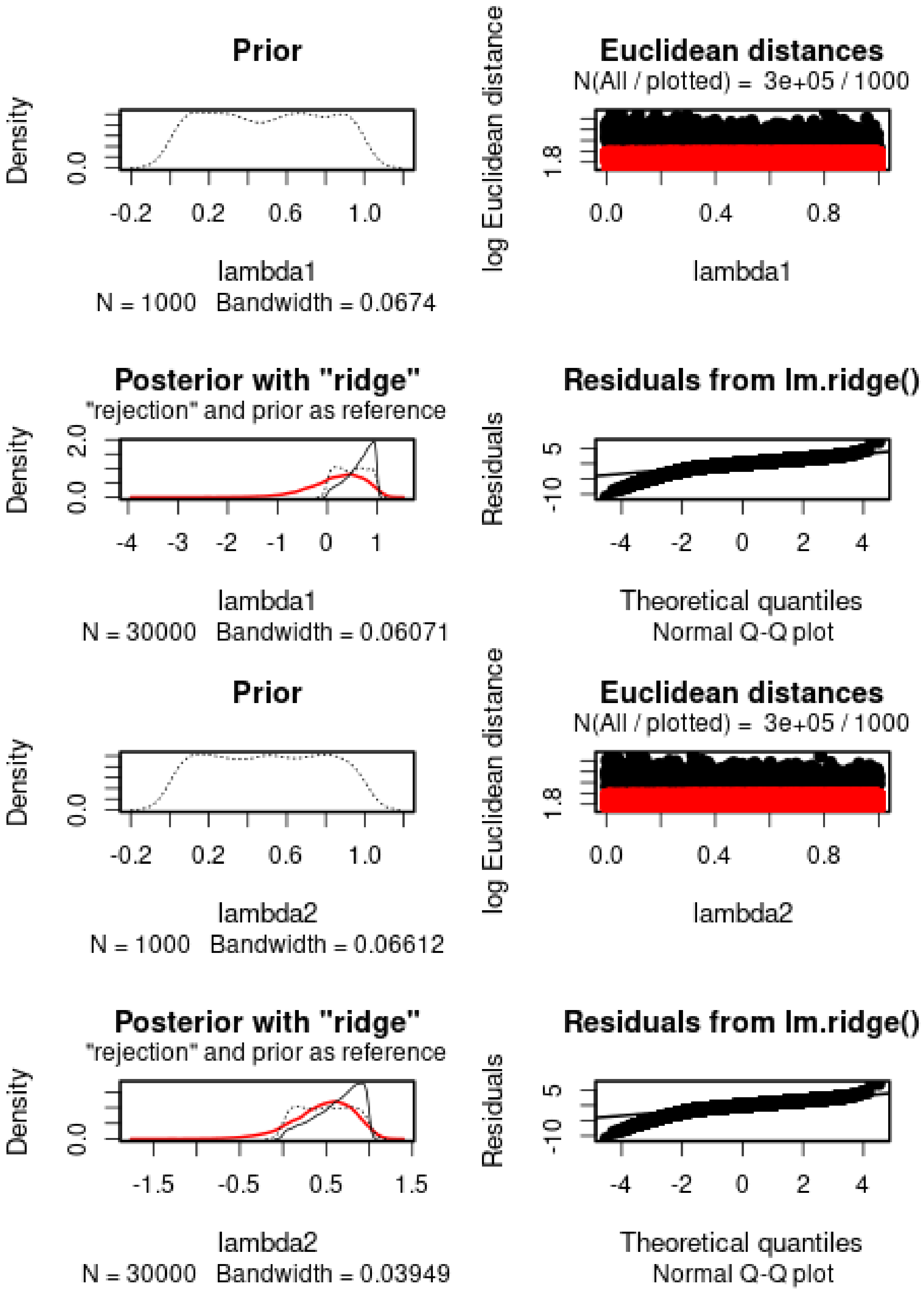
Prior
Density
0.0
-0.2 0.2 0.6 1.0
lambda1
N = 1000 Bandwidth = 0.0674
Euclidean distances
N(All / plotted) = 3e+05 / 1000
log Euclidean distance
1.8
0.0 0.4 0.8
lambda1
Posterior with "ridge"
"rejection" and prior as reference
Density
0.0 2.0
-4 -3 -2 -1 0 1
lambda1
N = 30000 Bandwidth = 0.06071
Residuals from lm.ridge()
Residuals
-10 5
-4 -2 0 2 4
Theoretical quantiles
Normal Q-Q plot
Prior
Density
0.0
-0.2 0.2 0.6 1.0
lambda2
N = 1000 Bandwidth = 0.06612
Euclidean distances
N(All / plotted) = 3e+05 / 1000
log Euclidean distance
1.8
0.0 0.4 0.8
lambda2
Posterior with "ridge"
"rejection" and prior as reference
Density
0.0
-1.5 -0.5 0.5 1.5
lambda2
N = 30000 Bandwidth = 0.03949
Residuals from lm.ridge()
Residuals
-10 5
-4 -2 0 2 4
Theoretical quantiles
Normal Q-Q plot

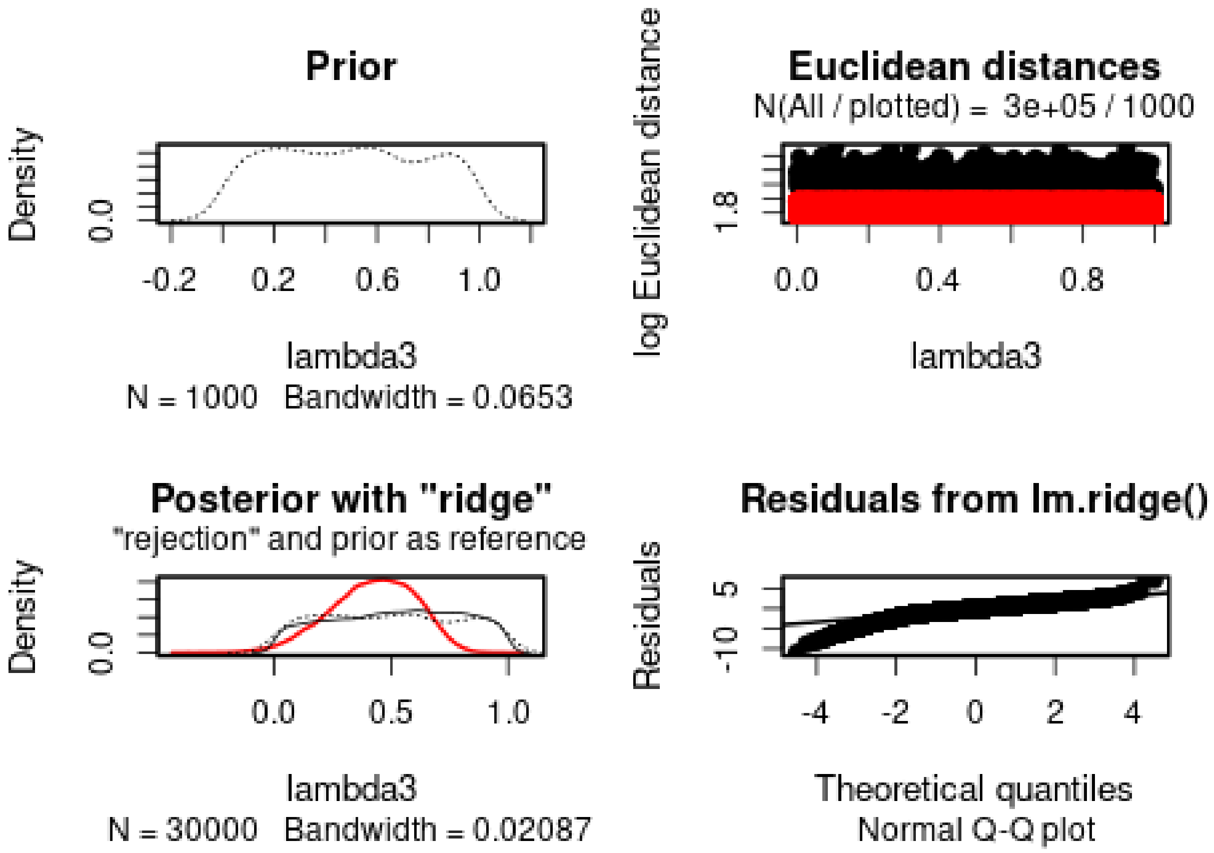

*h* cannot properly be estimated. *s* is positive (s = 0.6228). The 95% posterior interval includes 0. The *lambda* parameters are estimated to 0.2524, 0.4916, 0.4350, respectively.

We then check the distribution of summary statistics in the simulations:

```
library(reshape2)
library(ggplot2)
library(ggpubr)

df.stats.sim <- melt(stat.sim, variable.name = "Statistic", value.name = "Frequency")
```

```
## No id variables; using all as measure variables
```

```
l <- strsplit(as.character(df.stats.sim$Statistic), split = ".", fixed = TRUE)
df.stats.sim$Room <- sapply(l, function(x) x[1])
df.stats.sim$Genotype <- sapply(l, function(x) x[2])
df.stats.sim$Generation <- sapply(
  strsplit(as.character(df.stats.sim$Statistic), split = ".", fixed = TRUE),
  function(x) x[3])

df.stats.obs <- as.data.frame(stat.obs)
names(df.stats.obs) <- "Frequency"
df.stats.obs$Statistic <- row.names(df.stats.obs)
l <- strsplit(as.character(df.stats.obs$Statistic), split = ".", fixed = TRUE)
df.stats.obs$Room <- sapply(l, function(x) x[1])
df.stats.obs$Genotype <- sapply(l, function(x) x[2])
df.stats.obs$Generation <- sapply(
  strsplit(as.character(df.stats.obs$Statistic), split = ".", fixed = TRUE),
  function(x) x[3])
```

```
plot.obs <- function(gen) {
  p <- ggplot(data = subset(df.stats.sim, Generation == gen),
              aes(x = Frequency, y = ..density..)) +
    geom_histogram(bins = 50) +
    geom_vline(data = subset(df.stats.obs, Generation == gen),
               aes(xintercept = Frequency), color = "orange") +
    facet_grid(Genotype~Room) +
    theme_pubclean()
  return(p)
}
```

```
plot.obs(1) + ggtitle("Timepoint 4")
```

```
## Warning: The dot-dot notation (`..density..`) was deprecated in ggplot2 3.4.0.
## i Please use `after_stat(density)` instead.
## This warning is displayed once every 8 hours.
## Call `lifecycle::last_lifecycle_warnings()` to see where this warning was
## generated.
```

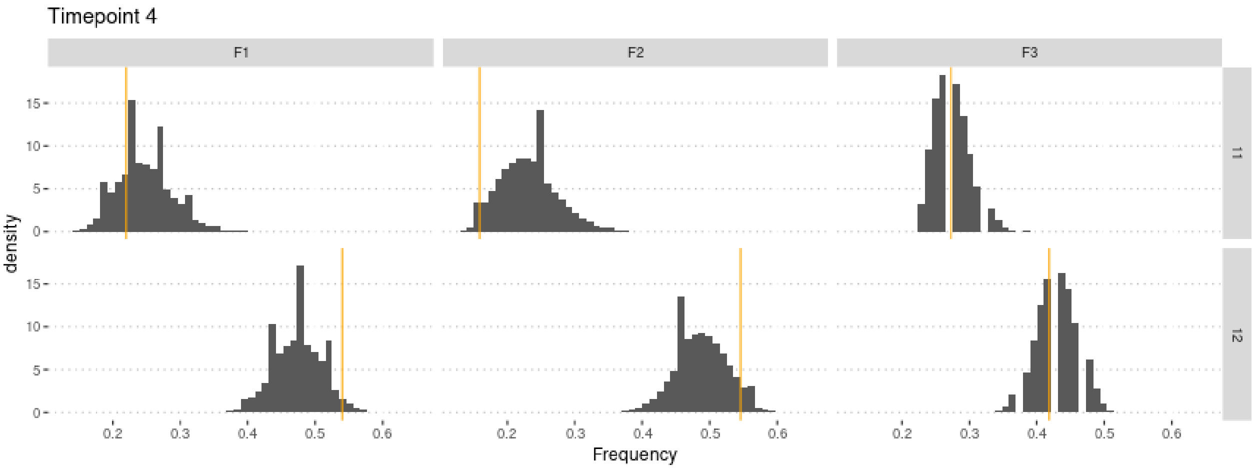

```
plot.obs(2) + ggtitle("Timepoint 5")
```

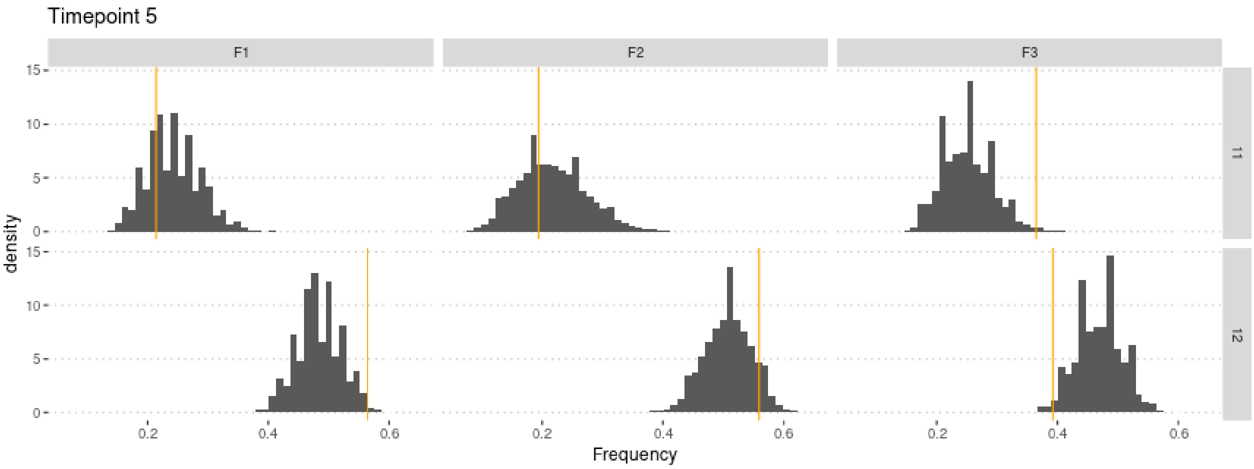

```
plot.obs(3) + ggtitle("Timepoint 6")
```

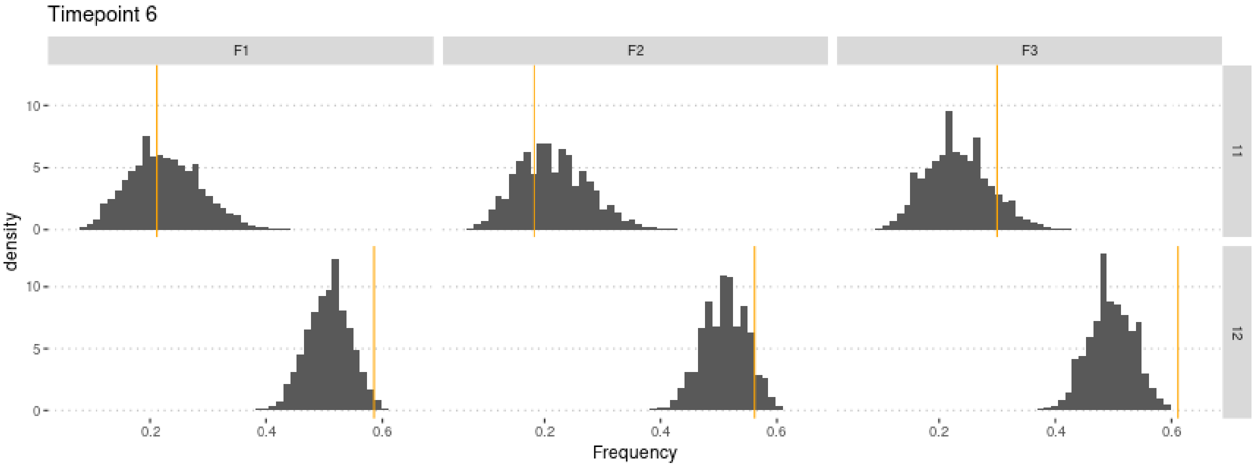

```
plot.obs(4) + ggtitle("Timepoint 7")
```

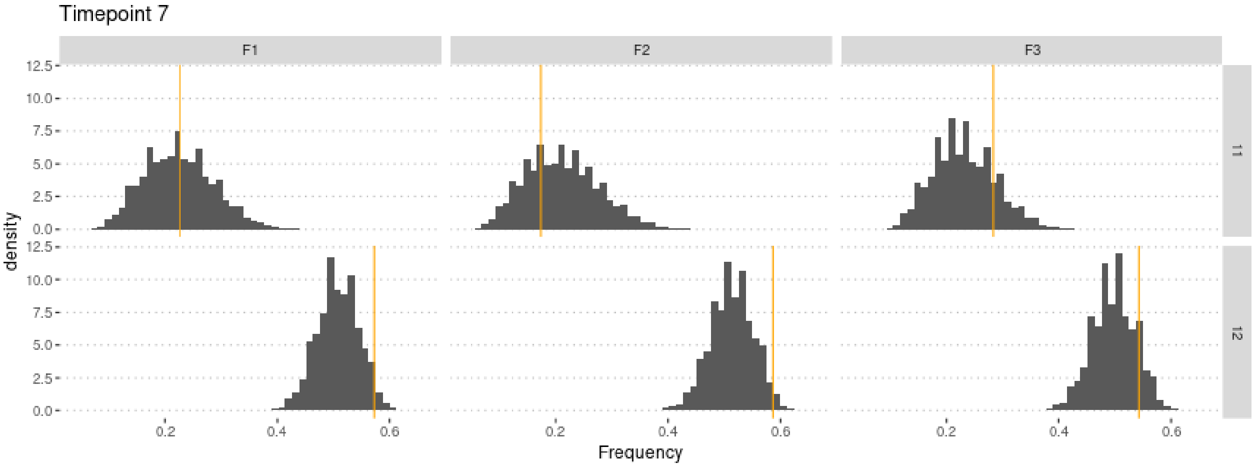

```
plot.obs(5) + ggtitle("Timepoint 8")
```

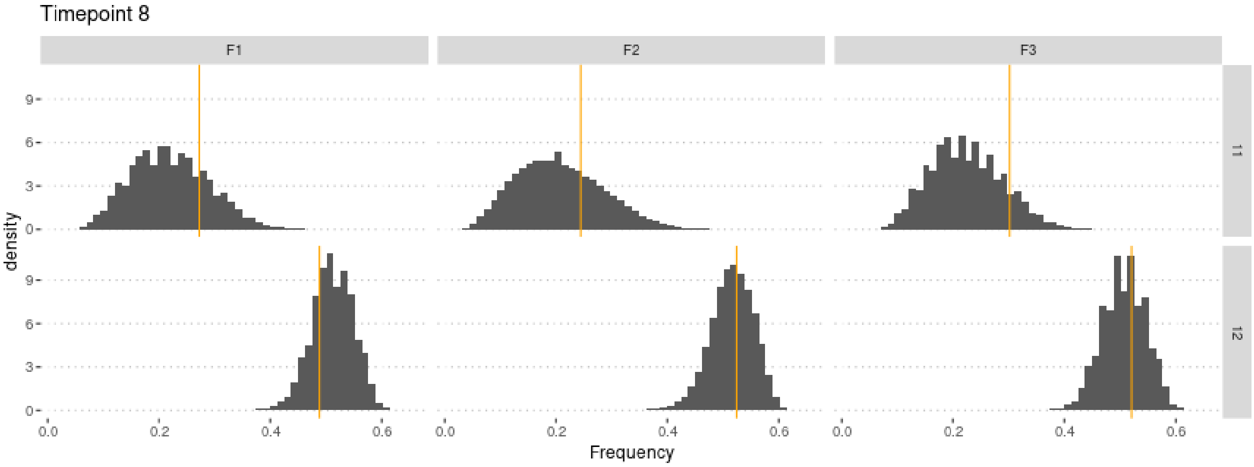

#### 4.1 Cross-Validation analysis

For memory-saving reasons, we only use 100,000 simulations in the following.

```r
sims2 <- sims[1:1e5,] #Not enough memory to use all simulations
param.sim2 <- param.sim[1:1e5,]
stat.sim2 <- stat.sim[1:1e5,]
m.r.g2 <- abc(target = stat.obs, param = param.sim2, sumstat = stat.sim2, tol = .1,
              method = "ridge", transf = c("none", "none", "none", "none", "none"))
```

Check that this does not affect the estimates too much:

```r
summary(m.r.g2, intvl = .95)
```

```
## Call:
## abc(target = stat.obs, param = param.sim2, sumstat = stat.sim2,
##     tol = 0.1, method = "ridge", transf = c("none", "none", "none",
##         "none", "none"))
## Data:
##  abc.out$adj.values (10000 posterior samples)
## Weights:
##  abc.out$weights
##
##                               hm      sm lambda1 lambda2 lambda3
## Min.:                     0.0343 -0.4026 -4.6987 -1.5110 -0.3662
## Weighted 2.5 % Perc.:     0.1487  0.0902 -1.1478 -0.2719  0.0375
## Weighted Median:          0.5384  0.6245  0.3479  0.5692  0.4411
## Weighted Mean:            0.5360  0.6531  0.2579  0.5246  0.4275
## Weighted Mode:            0.7862  0.5933  0.4662  0.7214  0.5178
## Weighted 97.5 % Perc.:    0.9286  1.3972  1.1854  1.0909  0.7416
## Max.:                     1.0429  3.8549  1.5084  1.4150  0.9399
```

Similar estimate, but slightly lower variance (95% CI of s no longer includes 0).

Compute predictions errors:

```r
if (redo) {
  cv.ridge.g <- cv4abc(
      param = param.sim2,
      sumstat = stat.sim2,
      abc.out = m.r.g2,
      nval = 1000,
      tols = c(.01, .1, .2))
  save(cv.ridge.g, file = "Rdata/MalesOnlyNonRandomMating3Param/backup_cv_genotype.Rdata")
} else {
  load("Rdata/MalesOnlyNonRandomMating3Param/backup_cv_genotype.Rdata")
}
summary(cv.ridge.g)
```

```
## Prediction error based on a cross-validation sample of 1000

##             hm        sm   lambda1   lambda2   lambda3
## 0.01 1.0061057 0.1283794 0.4059200 0.4160967 0.4903737
## 0.1  0.9621730 0.1271710 0.4258414 0.4268037 0.4900121
## 0.2  0.9639824 0.1307539 0.4445508 0.4451954 0.5010899
```

```r
plot(cv.ridge.g)
```

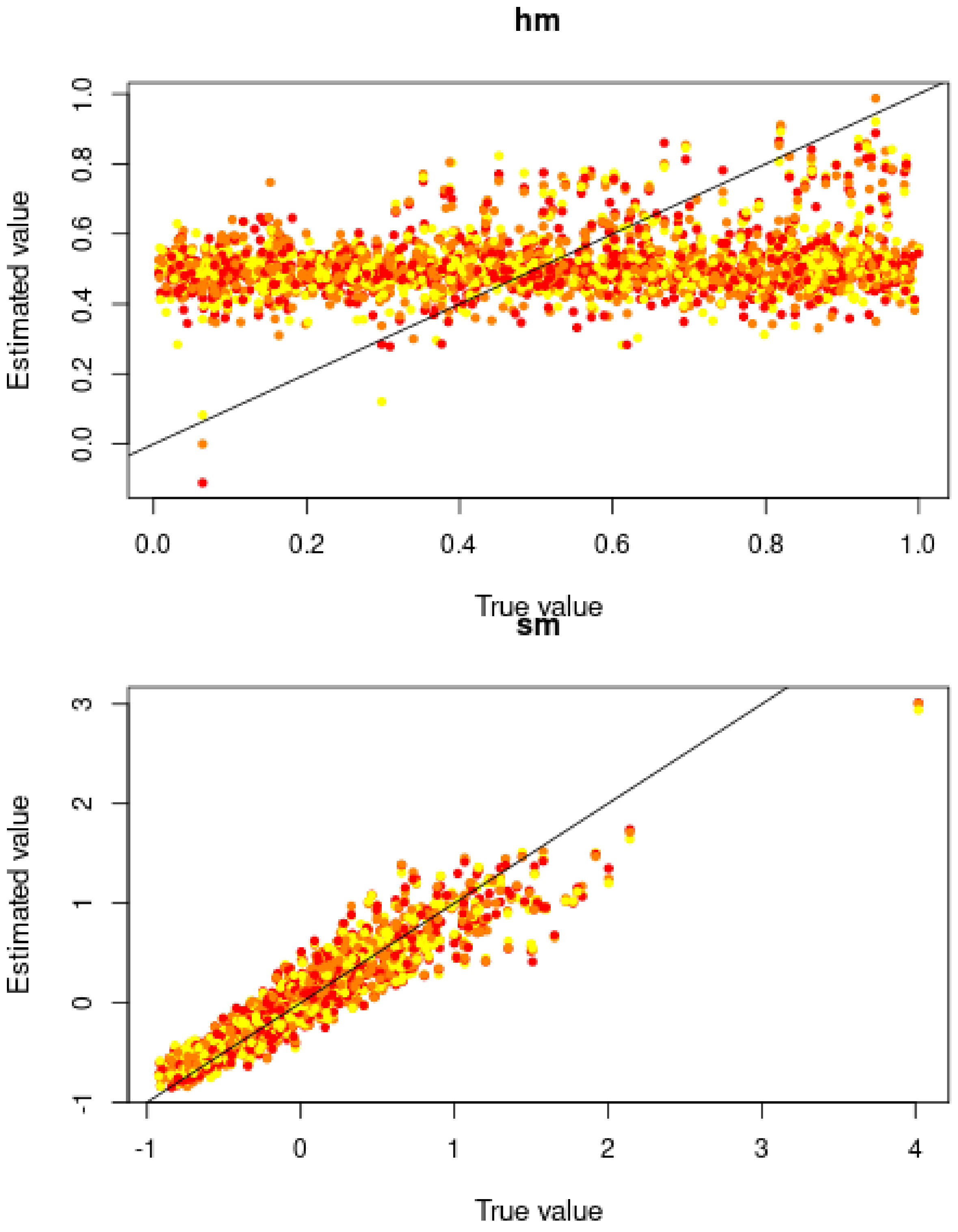

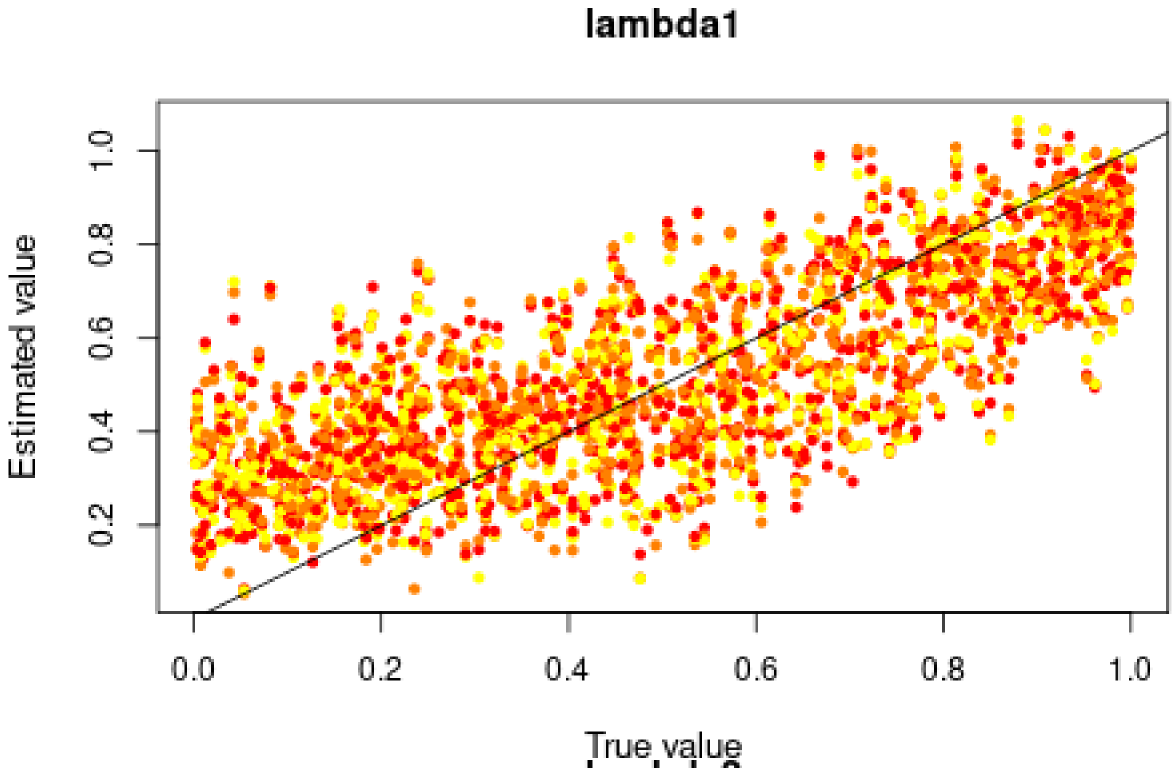
lambda1
Estimated value
True value
0.2
0.4
0.6
0.8
1.0
0.0
0.2
0.4
0.6
0.8
1.0

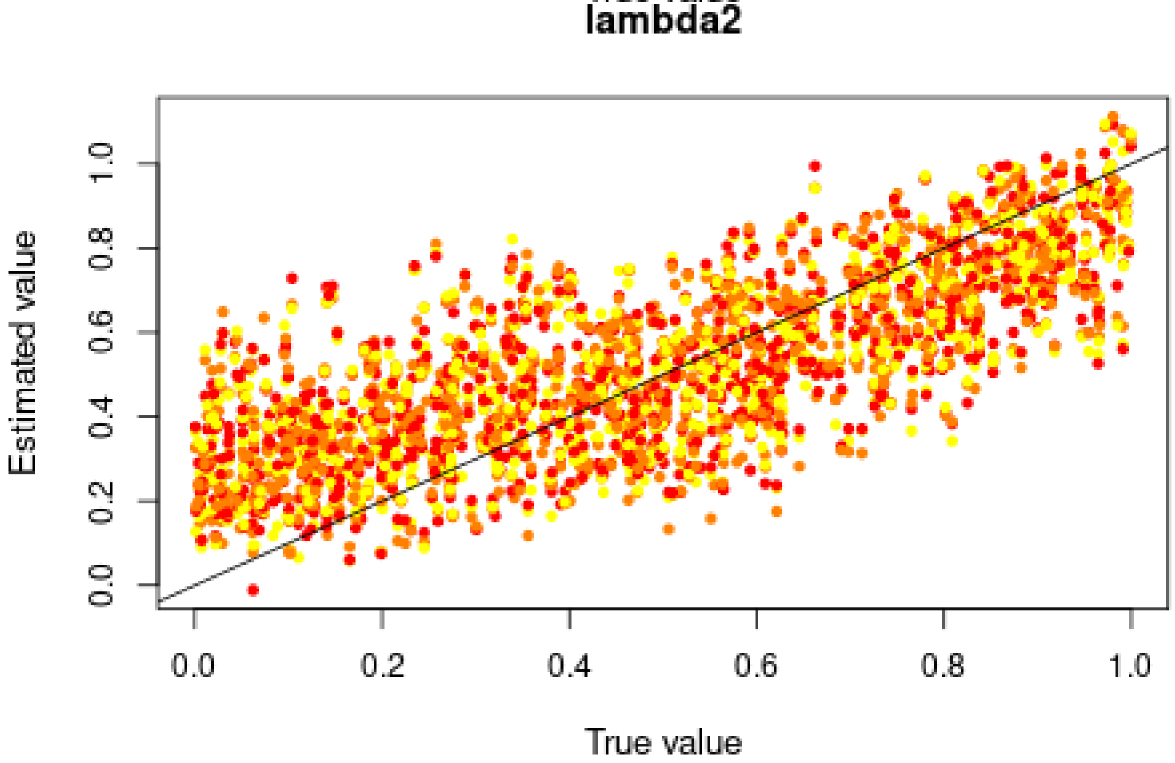
lambda2
Estimated value
True value
0.0
0.2
0.4
0.6
0.8
1.0
0.0
0.2
0.4
0.6
0.8
1.0

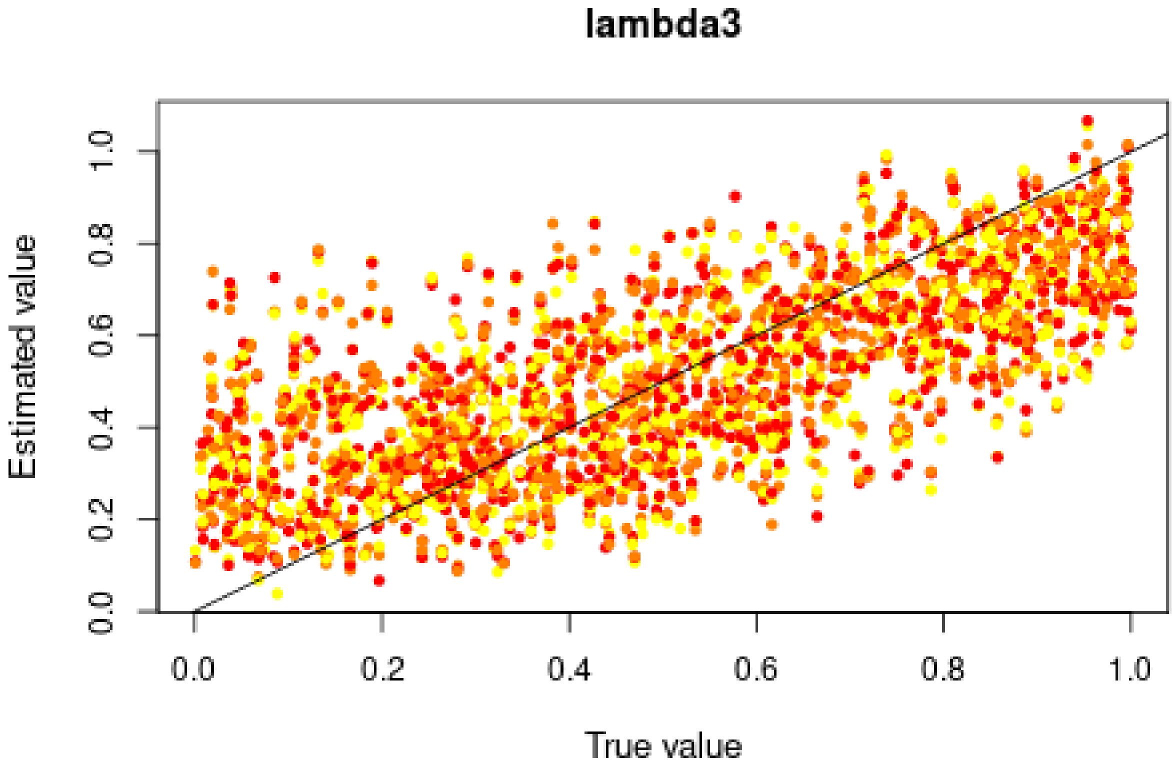

Good for $s$, but very bad for $h$ and intermediate for *lambda*. Results seem less good than when estimating a single *lambda*

#### 4.2 Misclassification errors

To test the power of the approach to distinguish between models, we also conduct a cross-validation experiment. We compare model M0 ($s = 0$) and M1 ($s = 0.6228 > 0$). $h$ is sampled over its prior distributions, and the *lambda*s are set to 0.2524, 0.4916, and 0.4350. 100,000 simulations were conducted under each model.

```
sims0 <- read.csv("Simulations/Simulations0NonRandomMating3ParamFromGen3.csv.gz")
param.sim0 <- subset(sims0, select = c("hm", "sm", "lambda1", "lambda2", "lambda3"))
stat.sim0 <- subset(sims0, select = 2:46)
# Only consider the WT homozygotes and the heterozygotes,
# as the sum of the three genotypes is 1
stat.sim0 <- subset(stat.sim0, select = grep(".22.", names(stat.sim0), invert = TRUE))

sims1 <- read.csv("Simulations/Simulations1bNonRandomMating3ParamMalesOnlyFromGen3.csv.gz")
param.sim1 <- subset(sims1, select = c("hm", "sm", "lambda1", "lambda2", "lambda3"))
stat.sim1 <- subset(sims1, select = 2:46)
# Only consider the WT homozygotes and the heterozygotes,
# as the sum of the three genotypes is 1
stat.sim1 <- subset(stat.sim1, select = grep(".22.", names(stat.sim1), invert = TRUE))
```

We conduct a CV analysis (this takes some time... set nval = 100 for faster results):

```
models <- rep(c("Neutral", "Selection"), each = 100000)
if (redo) {
  cv.modsel.g <- cv4postpr(
    models, rbind(stat.sim0, stat.sim1),
    nval = 1000, tols = .1, method = "mnlogistic")
  save(cv.modsel.g, file = "Rdata/MalesOnlyNonRandomMating3Param/backup_cv4postpr_genotype.Rdata")
```

```r
} else {
  load("Rdata/MalesOnlyNonRandomMating3Param/backup_cv4postpr_genotype.Rdata")
}
```

We display the results:

```r
summary(cv.modsel.g)
```

```
## Confusion matrix based on 1000 samples for each model.
##
## $tol0.1
##           Neutral Selection
## Neutral       909        91
## Selection      80       920
##
##
## Mean model posterior probabilities (mnlogistic)
##
## $tol0.1
##           Neutral Selection
## Neutral    0.8721    0.1279
## Selection  0.1205    0.8795
```

```r
plot(cv.modsel.g, names.arg=c("Neutral", "Selection"))
```

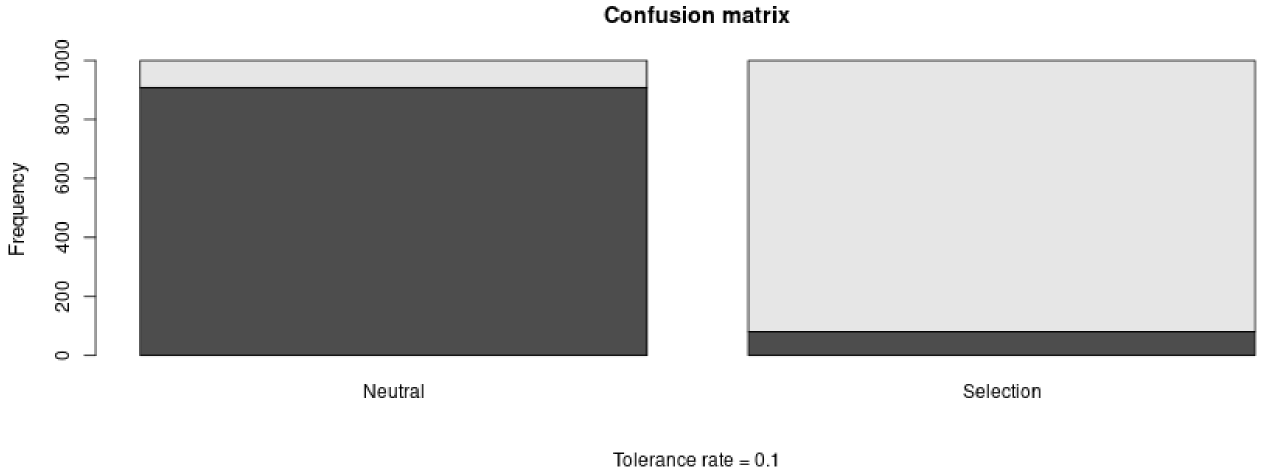

#### 4.3 Posterior prediction

```r
if (redo) {
  modsel.g <- postpr(
    stat.obs, models,
    rbind(stat.sim0, stat.sim1),
    tol = .1, method = "mnlogistic")
  save(modsel.g, file = "Rdata/MalesOnlyNonRandomMating3Param/backup_postpr_genotype.Rdata")
} else {
  load("Rdata/MalesOnlyNonRandomMating3Param/backup_postpr_genotype.Rdata")
}
summary(modsel.g)
```

```
## Call:
## postpr(target = stat.obs, index = models, sumstat = rbind(stat.sim0,
```

```
##     stat.sim1), tol = 0.1, method = "mnlogistic")
## Data:
##  postpr.out$values (20000 posterior samples)
## Models a priori:
##  Neutral, Selection
## Models a posteriori:
##  Neutral, Selection
##
## Proportion of accepted simulations (rejection):
##   Neutral Selection
##    0.5798    0.4201
##
## Bayes factors:
##           Neutral Selection
## Neutral    1.0000    1.3801
## Selection  0.7246    1.0000
##
##
## Posterior model probabilities (mnlogistic):
##   Neutral Selection
##    0.0226    0.9774
##
## Bayes factors:
##           Neutral Selection
## Neutral    1.0000    0.0231
## Selection 43.2589    1.0000
```

The model with selection is preferred.

#### 4.4 Goodness of fit

Under the neutral model:

```
if (redo) {
  res.gfit0.g <- gfit(
    target = stat.obs, sumstat = stat.sim0,
    statistic = median, nb.replicate = 1000)
  save(res.gfit0.g, file = "Rdata/MalesOnlyNonRandomMating3Param/backup_gfit0_genotype.Rdata")
} else {
  load("Rdata/MalesOnlyNonRandomMating3Param/backup_gfit0_genotype.Rdata")
}
plot(res.gfit0.g, main = "Histogram under M0")
```

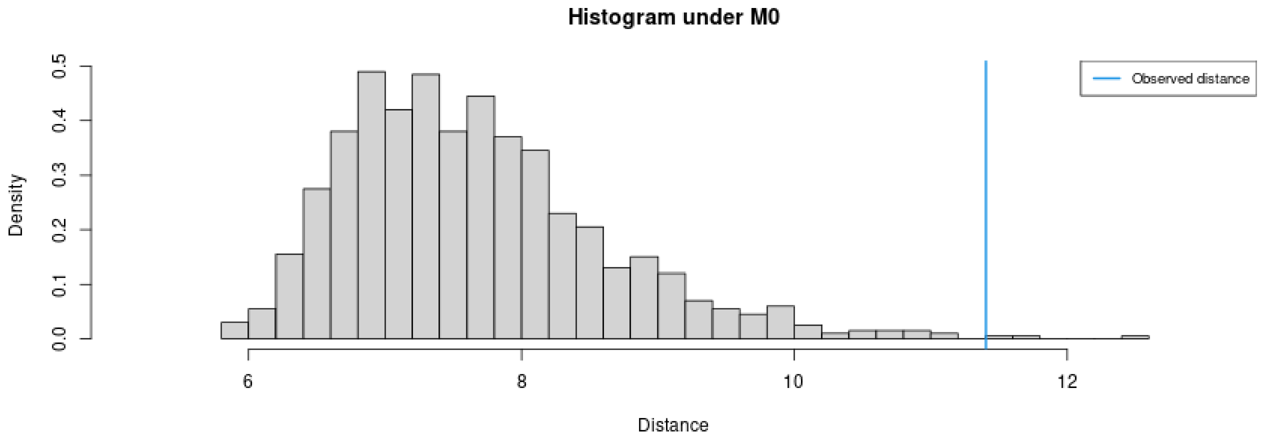

```
summary(res.gfit0.g)
```

```
## $pvalue
## [1] 0.003
##
## $s.dist.sim
##    Min. 1st Qu.  Median    Mean 3rd Qu.    Max.
##   5.821   6.954   7.505   7.652   8.158  12.413
##
## $dist.obs
## [1] 11.40589
```

Under the selection model:

```
if (redo) {
  res.gfit1.g <- gfit(
    target = stat.obs, sumstat = stat.sim1,
    statistic = median, nb.replicate = 1000)
  save(res.gfit1.g, file = "Rdata/MalesOnlyNonRandomMating3Param/backup_gfit1_genotype.Rdata")
} else {
  load("Rdata/MalesOnlyNonRandomMating3Param/backup_gfit1_genotype.Rdata")
}
plot(res.gfit1.g, main = "Histogram under M1")
```

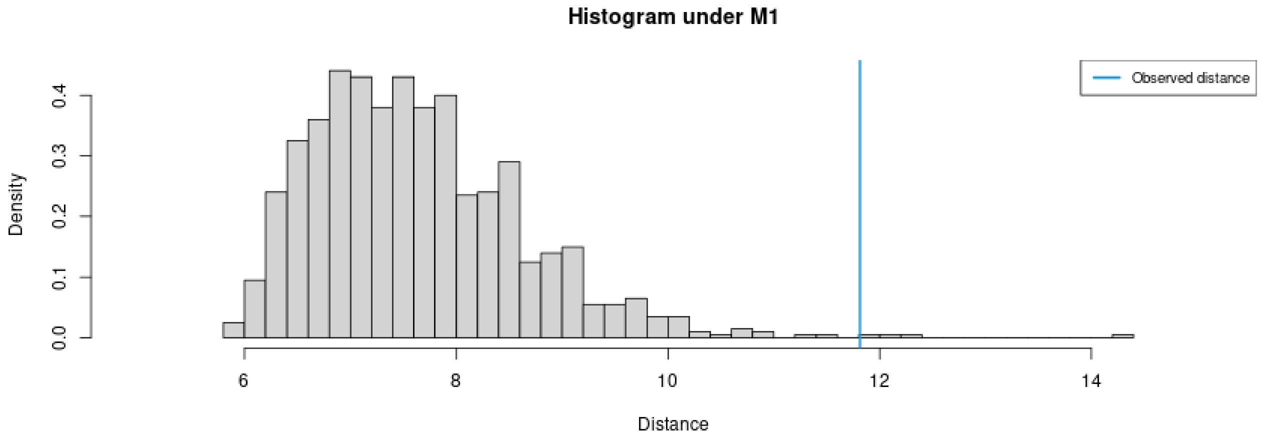

```
summary(res.gfit1.g)
```

```
## $pvalue
## [1] 0.004
```

```
##
## $s.dist.sim
##    Min. 1st Qu.  Median    Mean 3rd Qu.    Max.
##   5.809   6.904   7.501   7.630   8.202  14.304
##
## $dist.obs
## [1] 11.81235
```

M1 provides a better fit than M0.

#### 4.5 Summary figure:

Posterior distributions, with prior for comparison:

```r
library(ggplot2)
library(ggpubr)
dat.prior.h <- data.frame(h = param.sim[,"hm"], Distribution = "Prior")
dat.post.h <- data.frame(h = m.r.g$adj.values[,"hm"], Distribution = "Posterior")
dat.prior.s <- data.frame(s = param.sim[,"sm"], Distribution = "Prior")
dat.post.s <- data.frame(s = m.r.g$adj.values[,"sm"], Distribution = "Posterior")
dat.prior.l1 <- data.frame(s = param.sim[,"lambda1"], Distribution = "Prior")
dat.post.l1 <- data.frame(s = m.r.g$adj.values[,"lambda1"], Distribution = "Posterior")
dat.prior.l2 <- data.frame(s = param.sim[,"lambda2"], Distribution = "Prior")
dat.post.l2 <- data.frame(s = m.r.g$adj.values[,"lambda2"], Distribution = "Posterior")
dat.prior.l3 <- data.frame(s = param.sim[,"lambda3"], Distribution = "Prior")
dat.post.l3 <- data.frame(s = m.r.g$adj.values[,"lambda3"], Distribution = "Posterior")
dat.h <- rbind(dat.prior.h, dat.post.h)
dat.s <- rbind(dat.prior.s, dat.post.s)
dat.l1 <- rbind(dat.prior.l1, dat.post.l1)
dat.l2 <- rbind(dat.prior.l2, dat.post.l2)
dat.l3 <- rbind(dat.prior.l3, dat.post.l3)
dat.h$Distribution <- factor(dat.h$Distribution, levels = c("Prior", "Posterior"))
dat.s$Distribution <- factor(dat.s$Distribution, levels = c("Prior", "Posterior"))
dat.l1$Distribution <- factor(dat.l1$Distribution, levels = c("Prior", "Posterior"))
dat.l2$Distribution <- factor(dat.l2$Distribution, levels = c("Prior", "Posterior"))
dat.l3$Distribution <- factor(dat.l3$Distribution, levels = c("Prior", "Posterior"))
names(dat.h)[1] <- "value"
dat.h$variable <- "h"
names(dat.s)[1] <- "value"
dat.s$variable <- "s"
names(dat.l1)[1] <- "value"
dat.l1$variable <- "lambda1"
names(dat.l2)[1] <- "value"
dat.l2$variable <- "lambda2"
names(dat.l3)[1] <- "value"
dat.l3$variable <- "lambda3"
dat.dist <- rbind(dat.h, dat.s, dat.l1, dat.l2, dat.l3)
dat.dist$variable <- factor(dat.dist$variable, levels = c("h", "s", "g", "lambda1", "lambda2", "lambda3

p.dist <- ggplot(data = dat.dist, aes(x = value, linetype = Distribution, fill = Distribution)) +
          geom_density(alpha = 0.5) +
          scale_fill_brewer(type = "qual", palette = 3) +
          scale_linetype_manual(values = c(Prior = "dashed", Posterior = "solid")) +
          xlab("Parameter value") +
          facet_wrap(~variable, scales = "free") +
```

```
          ggtitle("Posterior and prior parameter distributions") +
          theme_pubclean() + theme(strip.background = element_blank())
```

Cross-validation:

```
d1<-rbind(as.data.frame(cv.ridge.g$estim$tol0.01),
          as.data.frame(cv.ridge.g$estim$tol0.1),
          as.data.frame(cv.ridge.g$estim$tol0.2))
d2<-rbind(cv.ridge.g$true, cv.ridge.g$true, cv.ridge.g$true)
d1<-melt(d1, value.name = "Estimated")
```

```
## No id variables; using all as measure variables
```

```
d1$Tolerance <- rep(c(0.01, 0.1, 0.2), each = 1000)
d1$Replicate <- rep(1:1000, 3)
d2<-melt(d2, value.name = "True")
```

```
## No id variables; using all as measure variables
```

```
d2$Replicate <- rep(1:1000, 3)
dat.cv <- merge(d1, d2, by = c("variable", "Replicate"))
dat.cv$variable <- factor(dat.cv$variable,
                          levels = c("hm", "sm", "lambda1", "lambda2", "lambda3"),
                          labels = c("h", "s", "lambda1", "lambda2", "lambda3"))
```

```
p.cv <- ggplot(dat.cv, aes(x = True, y = Estimated)) +
        geom_point(aes(col = as.ordered(Tolerance))) +
        geom_abline(slope = 1) +
        facet_wrap(~variable, scales = "free") +
        theme_pubclean() + labs(color = "Tolerance") +
        ggtitle("Cross validation") +
        theme(strip.background = element_blank())
```

Confusion matrix:

```
library(scales)
cv.sum <- summary(cv.modsel.g)
```

```
## Confusion matrix based on 1000 samples for each model.
##
## $tol0.1
##           Neutral Selection
## Neutral       909        91
## Selection      80       920
##
##
## Mean model posterior probabilities (mnlogistic)
##
## $tol0.1
##           Neutral Selection
## Neutral    0.8721    0.1279
## Selection  0.1205    0.8795
```

```
dat.cv <- as.data.frame(cv.sum$conf.matrix$tol0.1/1000)
names(dat.cv) <- c("Real", "Inferred", "Frequency")
p.confmat <- ggplot(dat.cv, aes(x = Real, y = Frequency, fill = Inferred)) +
             geom_col() +
```

```
             scale_y_continuous(labels = scales::percent) +
             scale_fill_brewer(type = "qual", palette = 3) +
             ggtitle("Prediction errors") +
             theme_pubclean()
```

Model probabilities:

```
p.mprob <- ggplot(as.data.frame(modsel.g$pred), aes(x = Var1, y = Freq, fill = Var1)) +
           geom_col() + ylab("Model posterior probability") + xlab("Model") +
           scale_fill_brewer(type = "qual", palette = 3) +
           theme_pubclean() +
           ggtitle("Model probabilities") +
           theme(legend.position = "none")
```

```
library(cowplot)
```

```
##
## Attaching package: 'cowplot'

## The following object is masked from 'package:ggpubr':
##
##     get_legend
```

```
p1 <- plot_grid(p.mprob, p.confmat, labels = c("C", "D"), nrow = 1)
p <- plot_grid(p.dist, p.cv, p1, labels = c("A", "B", ""), nrow = 3)
p
```

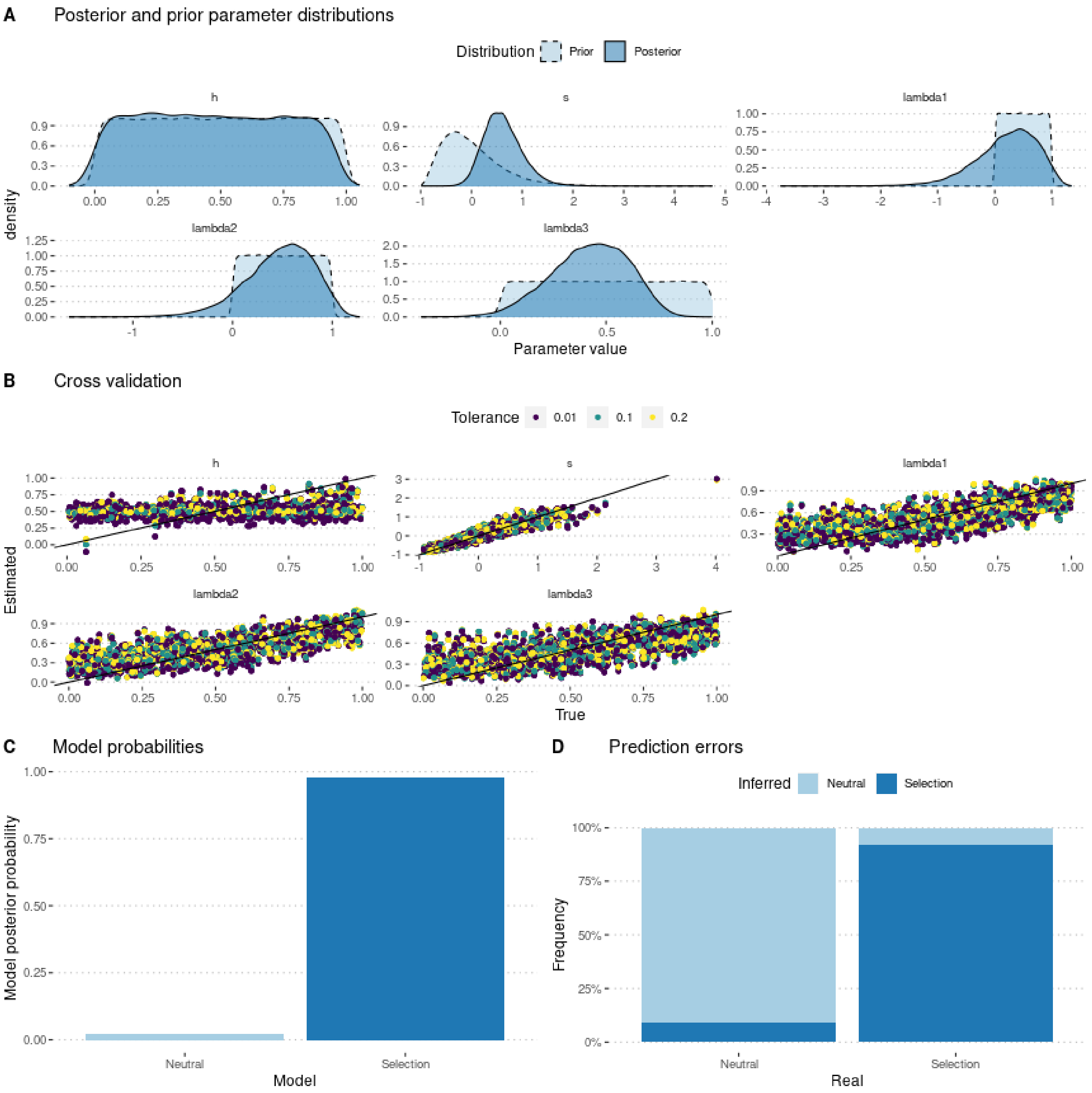

```r
ggsave(p, filename = "FigureABC-MalesOnlyNonRandomMating3Param-Genotype.pdf",
       width = 8, height = 10)
```
