## Supplementary material for "Experimental evaluation of a direct fitness effect of the *de novo* evolved mouse gene *Pldi*": combined suppl files: SupplementaryFile8.pdf

### Estimating poldi's selection coefficient

Julien Y. Dutheil

28/02/2023

### 1 Model parameterization

The model includes two parameters: $s$ the selection coefficient of the *poldi* allele compared to the knockout strain, which mimics the ancestral state, and $h$ the heterozygosity. Fitness values of each genotype are parameterized as follow:

$$\begin{aligned} poldi/poldi &: 1 \\ BL6/poldi &: 1 + h \cdot s \\ BL6/BL6 &: 1 + s \end{aligned}$$

where BL6 denotes the wild strain, and poldi the strain where the *poldi* gene is knocked out. Furthermore, we consider that the allele only has an effect in males, so that $h$ and $s$ are 0 in females (the two alleles have the same fitness). In this version, we model non random mating by adding a parameter $\lambda$, so that parents are chosen according to their fitness multiplied by $(1 - \lambda)$ if they have the same two alleles, $(1 - \lambda/2)$ if they have only one allele in common, or 1 if both alleles are different. We further account for non-genetic variance by modeling the fitness of an individual as a normal distribution with standard deviation $g$, and mean 1, $1 + h.s$, or $1 + s$, depending on its genotype.

We performed 300,000 simulations, using a gamma prior for $s$, a uniform prior between 0 and 1 for $h$, and a uniform prior between 0 and 1 for *lambda*.

### 2 Preamble

We load the data (observed and simulated):

```
sims <- read.csv(paste0("Simulations/",
    "SimulationsNonRandomMatingWithNonGeneticVarianceMalesOnlyFromGen3.csv.gz"))
```

```r
param.sim <- subset(sims, select = c("hm", "sm", "lambda", "g"))
stat.sim <- subset(sims, select = c(2:46))
# Only consider the WT homozygotes and the heterozygotes,
# as the sum of the three genotypes is 1
stat.sim <- subset(stat.sim, select = grep(".22.", names(stat.sim), invert = TRUE))

library(plyr, quietly = TRUE)
geno <- read.csv("GenotypeFrequencies.csv")
# We select the WT homozygote and heterozygote frequencies.
# Pldi homozygotes are 1 - sum(other two).
stat.obs <- subset(geno, TimePoint %in% 4:8, c("WT", "HET", "Replicate"))
stat.obs <- ddply(stat.obs, "Replicate", function(d) unlist(d[,1:2]))
stat.obs$Replicate <- NULL
stat.obs <- as.data.frame(t(stat.obs))
colnames(stat.obs) <- c("F1", "F2", "F3")
x <- rownames(stat.obs)
x <- gsub(x, pattern = "WT", replacement = ".11.")
x <- gsub(x, pattern = "HET", replacement = ".12.")
#x <- gsub(x, pattern = "Pldi", replacement = ".22.")
rownames(stat.obs) <- x
stat.obs1 <- subset(stat.obs, select = F1)
rownames(stat.obs1) <- paste0("F1", rownames(stat.obs1))
stat.obs2 <- subset(stat.obs, select = F2)
rownames(stat.obs2) <- paste0("F2", rownames(stat.obs2))
stat.obs3 <- subset(stat.obs, select = F3)
rownames(stat.obs3) <- paste0("F3", rownames(stat.obs3))
stat.obs <- cbind(t(stat.obs1), t(stat.obs2), t(stat.obs3))[1,]
```

Because *lambda* cannot be estimated with allelic frequencies, we only use genotype frequencies.

### 3 Using genotype frequencies

We estimate parameters using the ridge regression method:

```r
library(abc, quietly = TRUE)
```

```
##
## Attaching package: 'SparseM'

## The following object is masked from 'package:base':
##
##     backsolve

## locfit 1.5-9.8    2023-06-11
```

```r
if (redo) {
  m.r.g <- abc(target = stat.obs, param = param.sim, sumstat = stat.sim, tol = .1,
              method = "ridge", transf = c("none", "none", "none", "none"))
  save(m.r.g,
       file = "Rdata/MalesOnlyNonRandomMatingNonGeneticVariance/backup_abc_genotype.Rdata")
} else {
  load("Rdata/MalesOnlyNonRandomMatingNonGeneticVariance/backup_abc_genotype.Rdata")
}
```

Now we display the results:

```r
summary(m.r.g, intvl = .95)
```

```
## Call:
## abc(target = stat.obs, param = param.sim, sumstat = stat.sim,
##     tol = 0.1, method = "ridge", transf = c("none", "none", "none",
##         "none"))
## Data:
##  abc.out$adj.values (30000 posterior samples)
## Weights:
##  abc.out$weights
##
##                                hm      sm  lambda       g
## Min.:                     -0.0764 -0.5519 -2.6782  0.0195
## Weighted 2.5 % Perc.:      0.0163  0.0148 -0.6789  0.0601
## Weighted Median:           0.4647  0.6073  0.4524  0.2666
## Weighted Mean:             0.4692  0.6409  0.3895  0.2663
## Weighted Mode:             0.3122  0.5660  0.6459  0.3031
## Weighted 97.5 % Perc.:     0.9326  1.4576  1.1001  0.4748
## Max.:                      1.0480  3.9745  1.5044  0.5171
```

Distribution of parameter estimates:

```r
hist(m.r.g, breaks = 30,
     caption = c(expression(h), expression(s), expression(lambda), expression(g)))
```

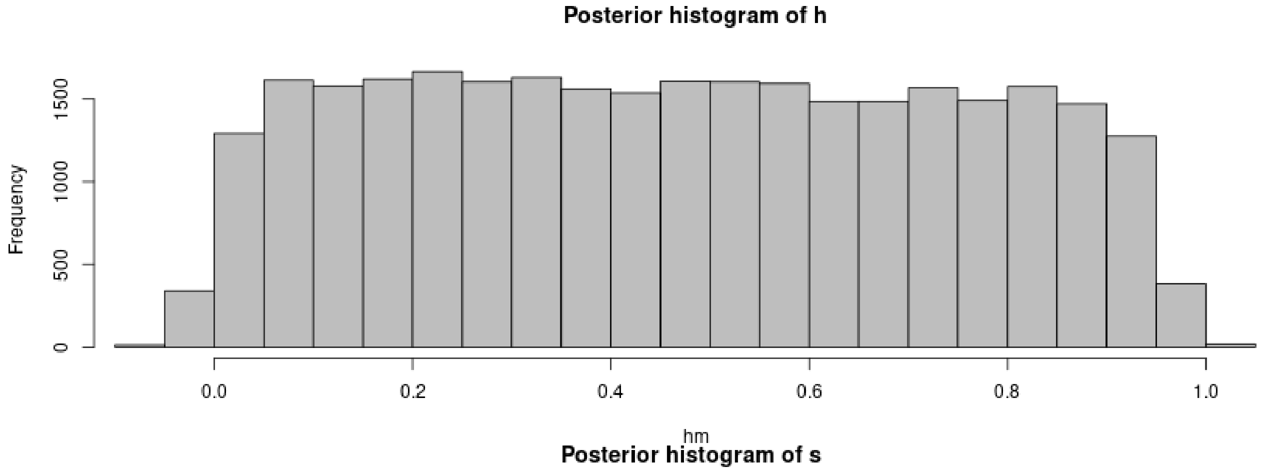

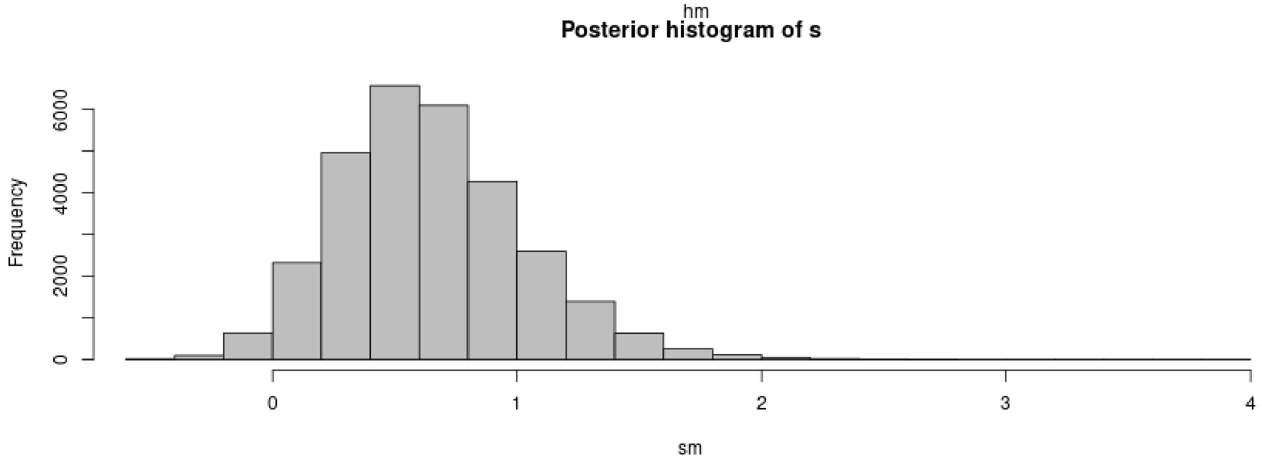

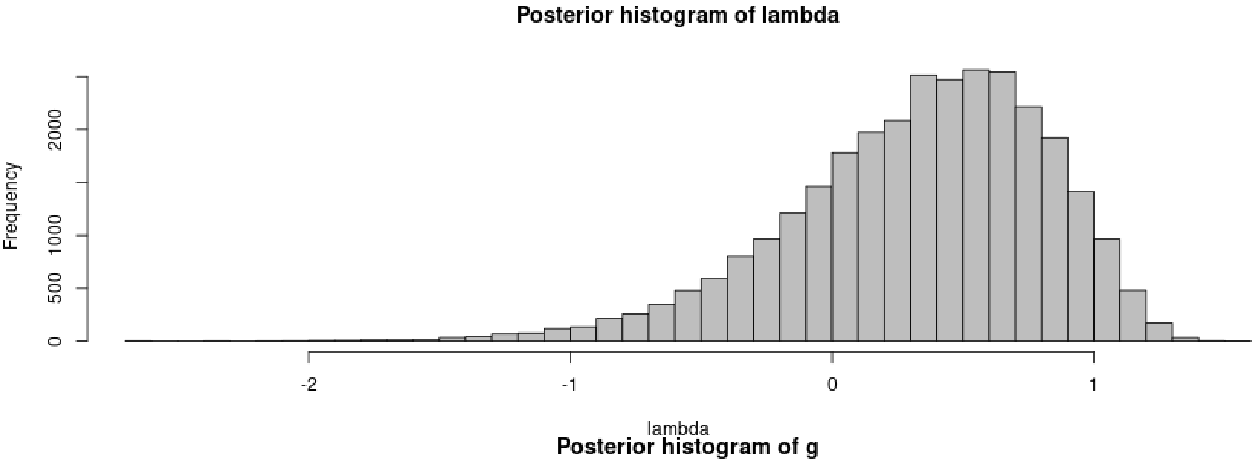

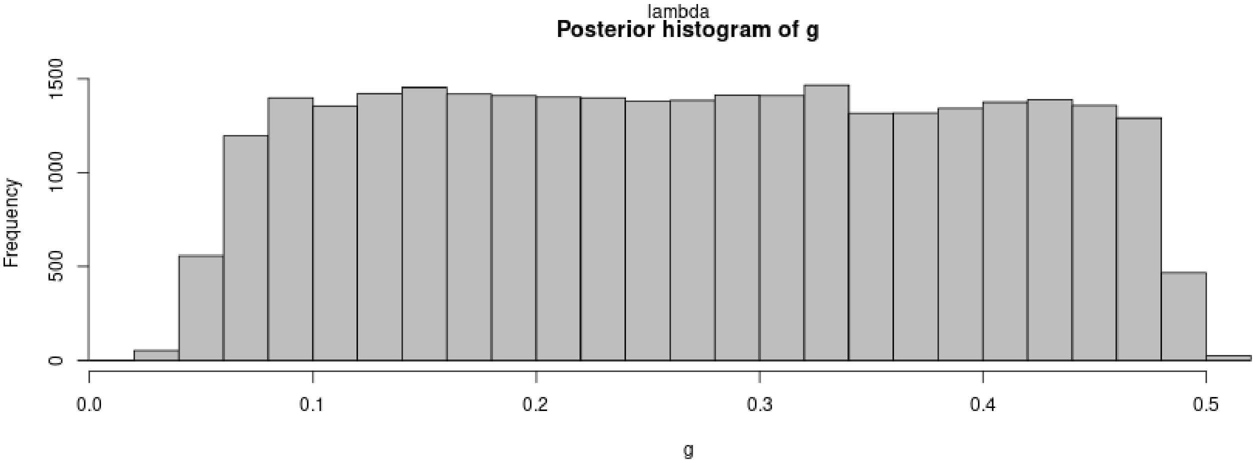

```
plot(m.r.g, param = param.sim)
```

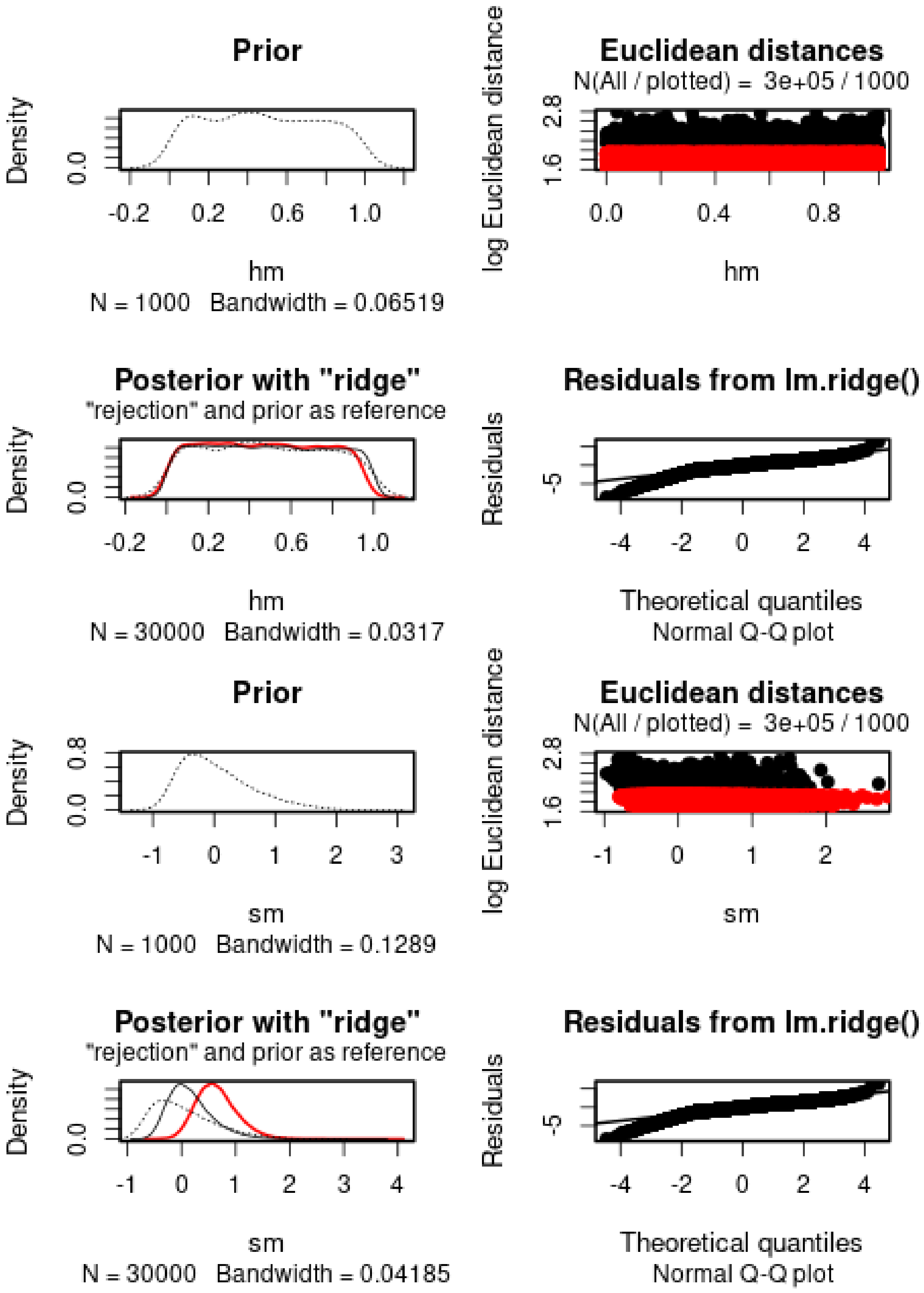
Prior
Density
0.0
-0.2 0.2 0.6 1.0
hm
N = 1000 Bandwidth = 0.06519
Euclidean distances
N(All / plotted) = 3e+05 / 1000
log Euclidean distance
1.6 2.8
0.0 0.4 0.8
hm
Posterior with "ridge"
"rejection" and prior as reference
Density
0.0
-0.2 0.2 0.6 1.0
hm
N = 30000 Bandwidth = 0.0317
Residuals from lm.ridge()
Residuals
-5
-4 -2 0 2 4
Theoretical quantiles
Normal Q-Q plot
Prior
Density
0.0 0.8
-1 0 1 2 3
sm
N = 1000 Bandwidth = 0.1289
Euclidean distances
N(All / plotted) = 3e+05 / 1000
log Euclidean distance
1.6 2.8
-1 0 1 2
sm
Posterior with "ridge"
"rejection" and prior as reference
Density
0.0
-1 0 1 2 3 4
sm
N = 30000 Bandwidth = 0.04185
Residuals from lm.ridge()
Residuals
-5
-4 -2 0 2 4
Theoretical quantiles
Normal Q-Q plot

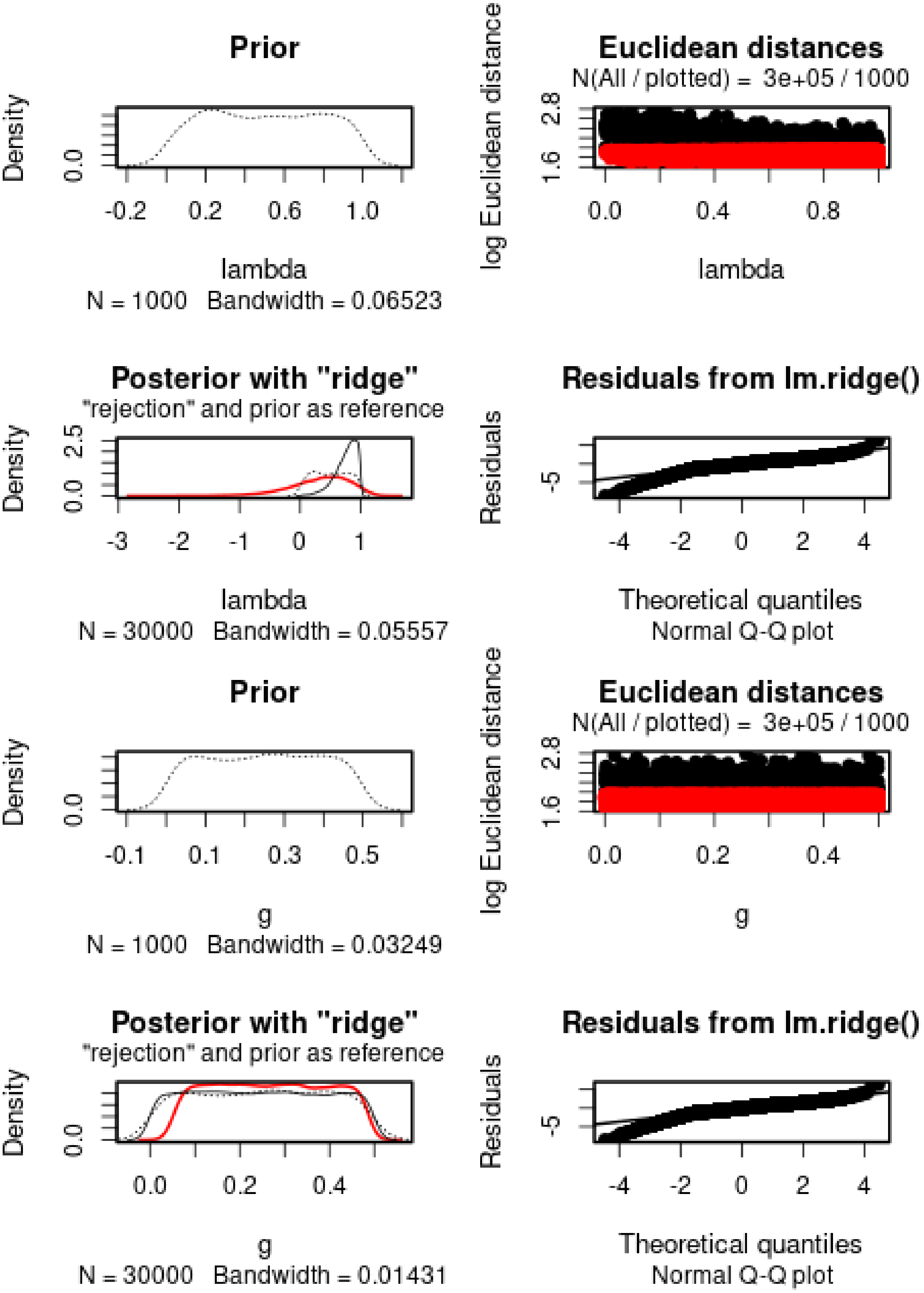
Prior
Density
0.0
-0.2
0.2
0.6
1.0
lambda
N = 1000 Bandwidth = 0.06523
Euclidean distances
N(All / plotted) = 3e+05 / 1000
log Euclidean distance
1.6
2.8
0.0
0.4
0.8
lambda
Posterior with "ridge"
"rejection" and prior as reference
Density
0.0
2.5
-3
-2
-1
0
1
lambda
N = 30000 Bandwidth = 0.05557
Residuals from lm.ridge()
Residuals
-5
-4
-2
0
2
4
Theoretical quantiles
Normal Q-Q plot
Prior
Density
0.0
-0.1
0.1
0.3
0.5
g
N = 1000 Bandwidth = 0.03249
Euclidean distances
N(All / plotted) = 3e+05 / 1000
log Euclidean distance
1.6
2.8
0.0
0.2
0.4
g
Posterior with "ridge"
"rejection" and prior as reference
Density
0.0
0.0
0.2
0.4
g
N = 30000 Bandwidth = 0.01431
Residuals from lm.ridge()
Residuals
-5
-4
-2
0
2
4
Theoretical quantiles
Normal Q-Q plot

$h$ cannot properly be estimated. $s$ is positive, estimated to 0.6409. The 95% posterior interval does not include 0. $g$ cannot be estimated. $lambda$ is estimated to 0.3895.

We then check the distribution of summary statistics in the simulations:

```
library(reshape2)
library(ggplot2)
library(ggpubr)
```

```
##
## Attaching package: 'ggpubr'

## The following object is masked from 'package:plyr':
##
##     mutate
```

```
df.stats.sim <- melt(stat.sim, variable.name = "Statistic", value.name = "Frequency")
```

```
## No id variables; using all as measure variables
```

```
l <- strsplit(as.character(df.stats.sim$Statistic), split = ".", fixed = TRUE)
df.stats.sim$Room <- sapply(l, function(x) x[1])
df.stats.sim$Genotype <- sapply(l, function(x) x[2])
df.stats.sim$Generation <- sapply(
  strsplit(as.character(df.stats.sim$Statistic), split = ".", fixed = TRUE),
  function(x) x[3])

df.stats.obs <- as.data.frame(stat.obs)
names(df.stats.obs) <- "Frequency"
df.stats.obs$Statistic <- row.names(df.stats.obs)
l <- strsplit(as.character(df.stats.obs$Statistic), split = ".", fixed = TRUE)
df.stats.obs$Room <- sapply(l, function(x) x[1])
df.stats.obs$Genotype <- sapply(l, function(x) x[2])
df.stats.obs$Generation <- sapply(
  strsplit(as.character(df.stats.obs$Statistic), split = ".", fixed = TRUE),
  function(x) x[3])

plot.obs <- function(gen) {
  p <- ggplot(data = subset(df.stats.sim, Generation == gen),
              aes(x = Frequency, y = ..density..)) +
    geom_histogram(bins = 50) +
    geom_vline(data = subset(df.stats.obs, Generation == gen),
               aes(xintercept = Frequency), color = "orange") +
    facet_grid(Genotype~Room) +
    theme_pubclean()
  return(p)
}
```

```
plot.obs(1) + ggtitle("Timepoint 4")
```

```
## Warning: The dot-dot notation (`..density..`) was deprecated in ggplot2 3.4.0.
## i Please use `after_stat(density)` instead.
## This warning is displayed once every 8 hours.
## Call `lifecycle::last_lifecycle_warnings()` to see where this warning was
## generated.
```

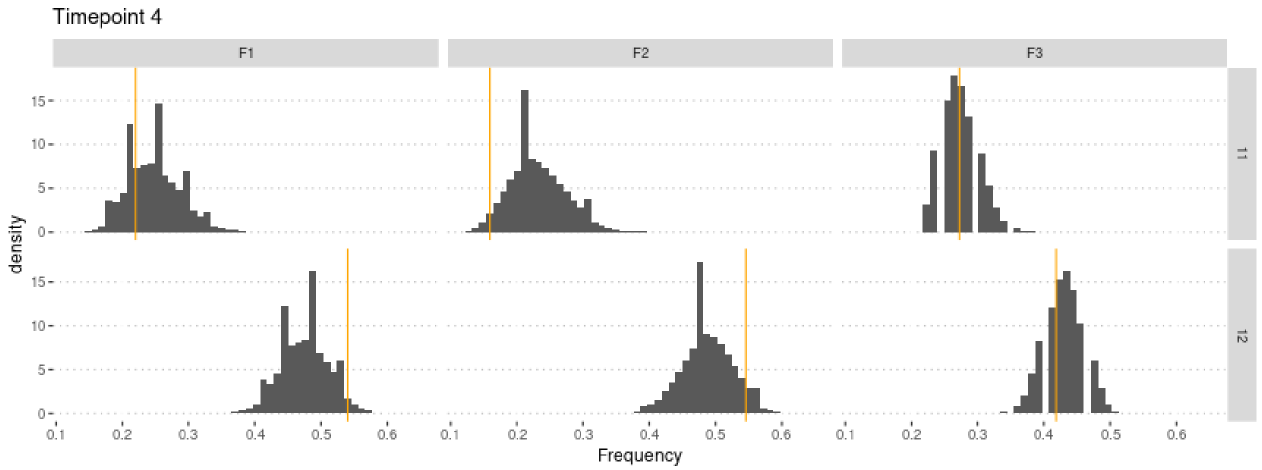

```
plot.obs(2) + ggtitle("Timepoint 5")
```

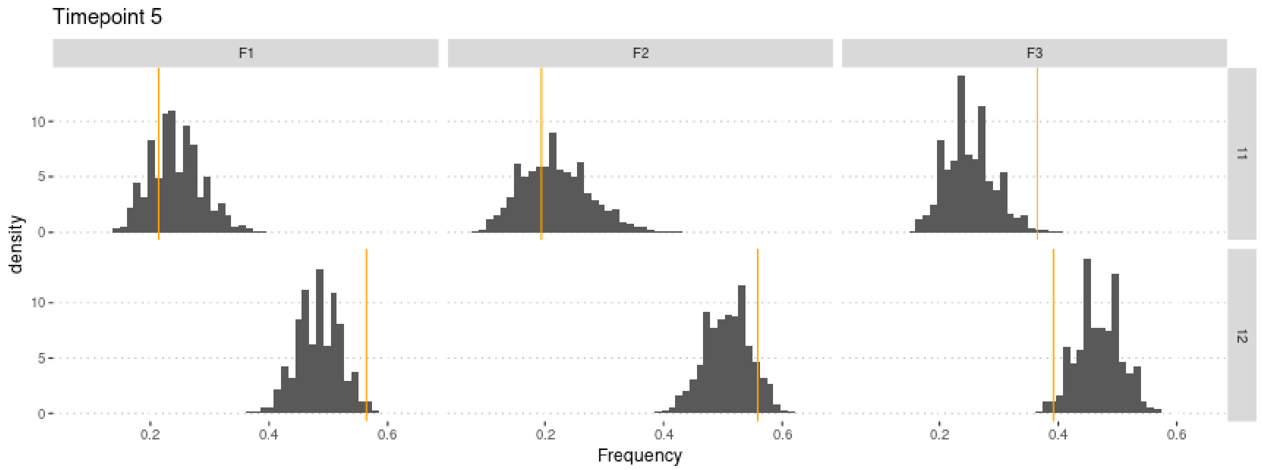

```
plot.obs(3) + ggtitle("Timepoint 6")
```

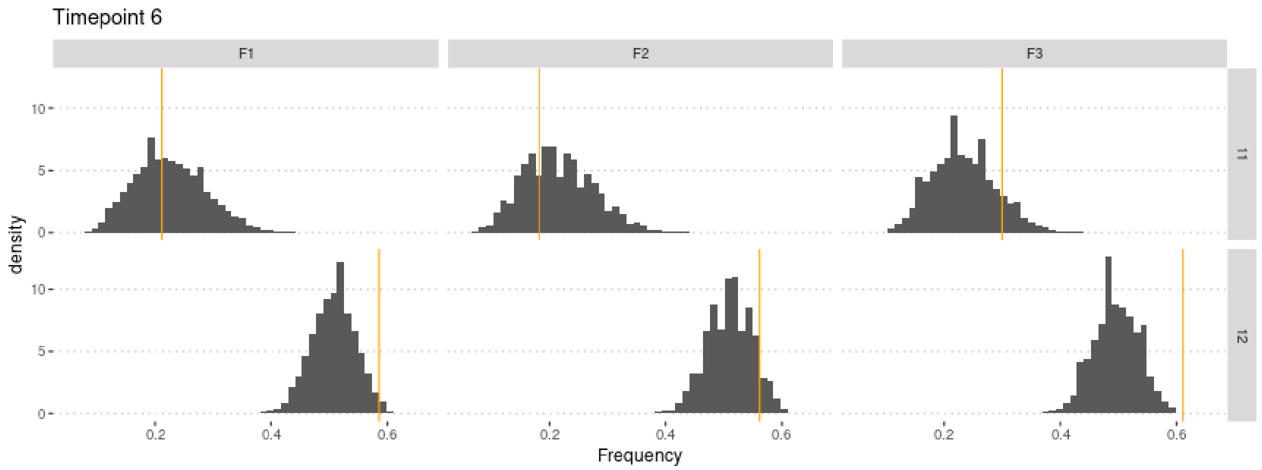

```
plot.obs(4) + ggtitle("Timepoint 7")
```

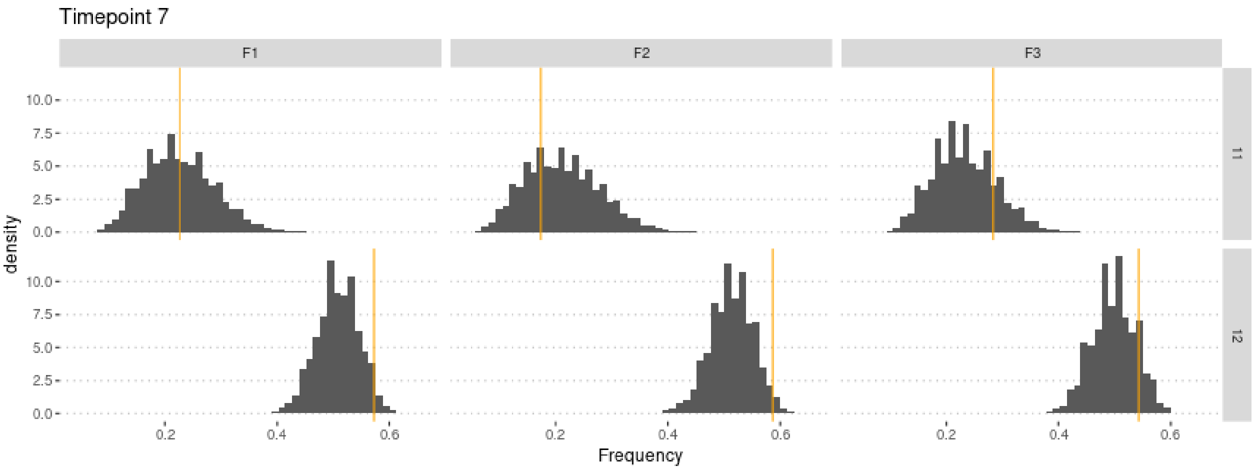

```
plot.obs(5) + ggtitle("Timepoint 8")
```

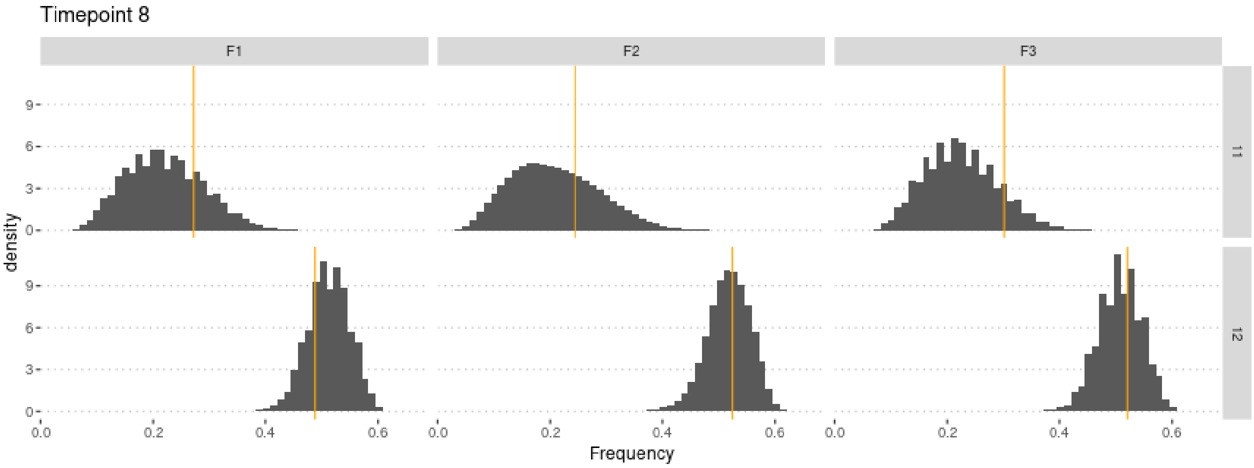

#### 3.1 Cross-Validation analysis

For memory-saving reasons, we only use 100,000 simulations in the following.

```
sims2 <- sims[1:1e5,] #Not enough memory to use all simulations
param.sim2 <- param.sim[1:1e5,]
stat.sim2 <- stat.sim[1:1e5,]
m.r.g2 <- abc(target = stat.obs, param = param.sim2, sumstat = stat.sim2, tol = .1,
              method = "ridge", transf = c("none", "none", "none", "none"))
```

Check that this does not affect the estimates too much:

```
summary(m.r.g2, intvl = .95)
```

```
## Call:
## abc(target = stat.obs, param = param.sim2, sumstat = stat.sim2,
##     tol = 0.1, method = "ridge", transf = c("none", "none", "none",
##         "none"))
## Data:
##  abc.out$adj.values (10000 posterior samples)
## Weights:
##  abc.out$weights
##
```

```
##                              hm      sm  lambda       g
## Min.:                   -0.3459 -0.4788 -2.4340  0.0339
## Weighted 2.5 % Perc.:  -0.1404  0.0645 -0.5732  0.0925
## Weighted Median:         0.4277  0.6036  0.5227  0.2559
## Weighted Mean:           0.4316  0.6318  0.4605  0.2574
## Weighted Mode:           0.0733  0.5877  0.6881  0.1476
## Weighted 97.5 % Perc.:  1.0260  1.3805  1.1740  0.4274
## Max.:                    1.2556  2.4402  1.5487  0.4617
```

Similar estimate, but slightly lower variance.

Compute predictions errors:

```
if (redo) {
  cv.ridge.g <- cv4abc(
      param = param.sim2,
      sumstat = stat.sim2,
      abc.out = m.r.g2,
      nval = 1000,
      tols = c(.01, .1, .2))
  save(cv.ridge.g,
       file = "Rdata/MalesOnlyNonRandomMatingNonGeneticVariance/backup_cv_genotype.Rdata")
} else {
  load("Rdata/MalesOnlyNonRandomMatingNonGeneticVariance/backup_cv_genotype.Rdata")
}
summary(cv.ridge.g)
```

```
## Prediction error based on a cross-validation sample of 1000
```

```
##             hm        sm    lambda         g
## 0.01 0.9777445 0.1579347 0.2412420 1.0320263
## 0.1  0.9433524 0.1609432 0.2845795 0.9924391
## 0.2  0.9444153 0.1660739 0.3196877 0.9913588
```

```
plot(cv.ridge.g)
```

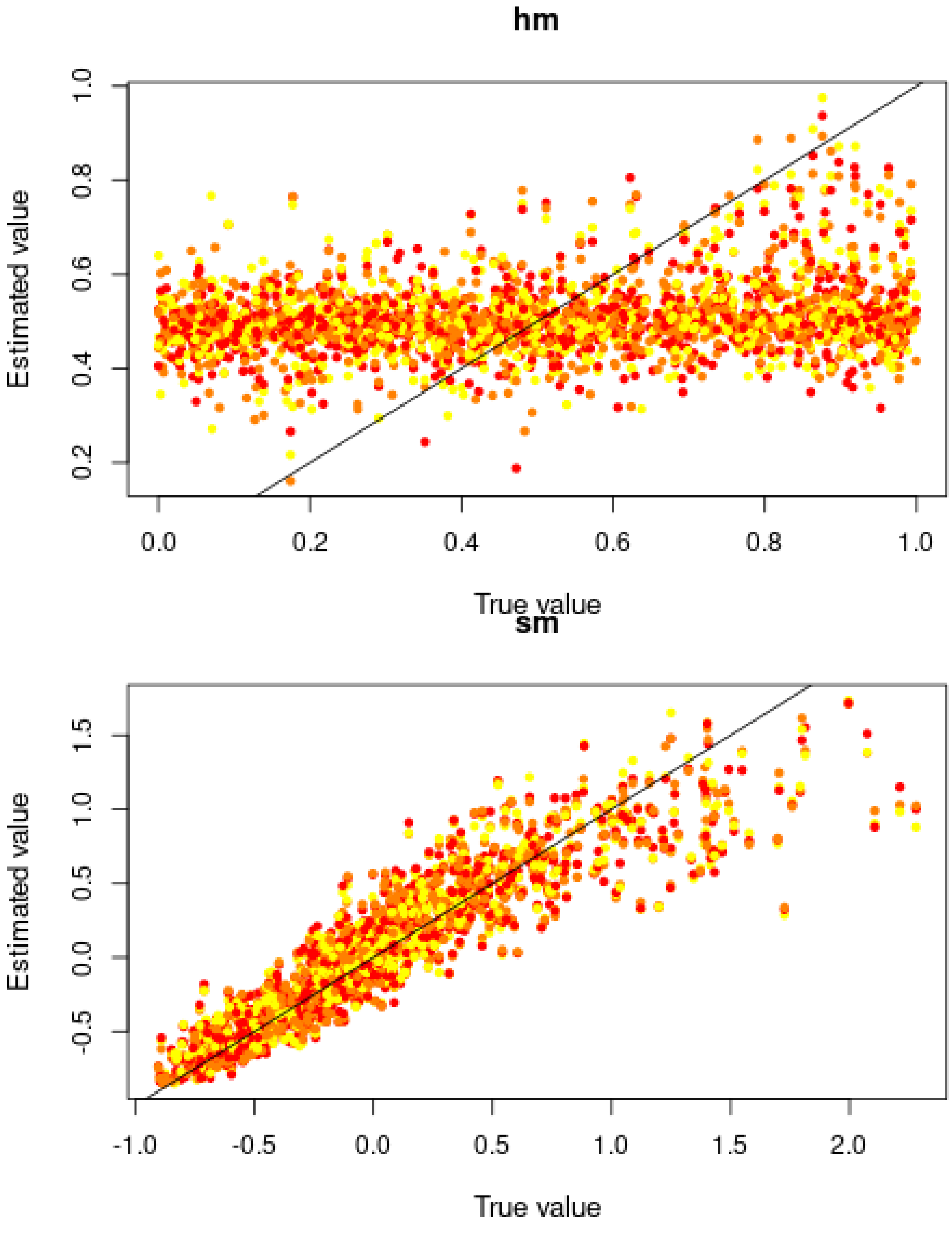
hm
Estimated value
0.2
0.4
0.6
0.8
1.0
0.0
0.2
0.4
0.6
0.8
1.0
True value
sm
Estimated value
-0.5
0.0
0.5
1.0
1.5
-1.0
-0.5
0.0
0.5
1.0
1.5
2.0
True value

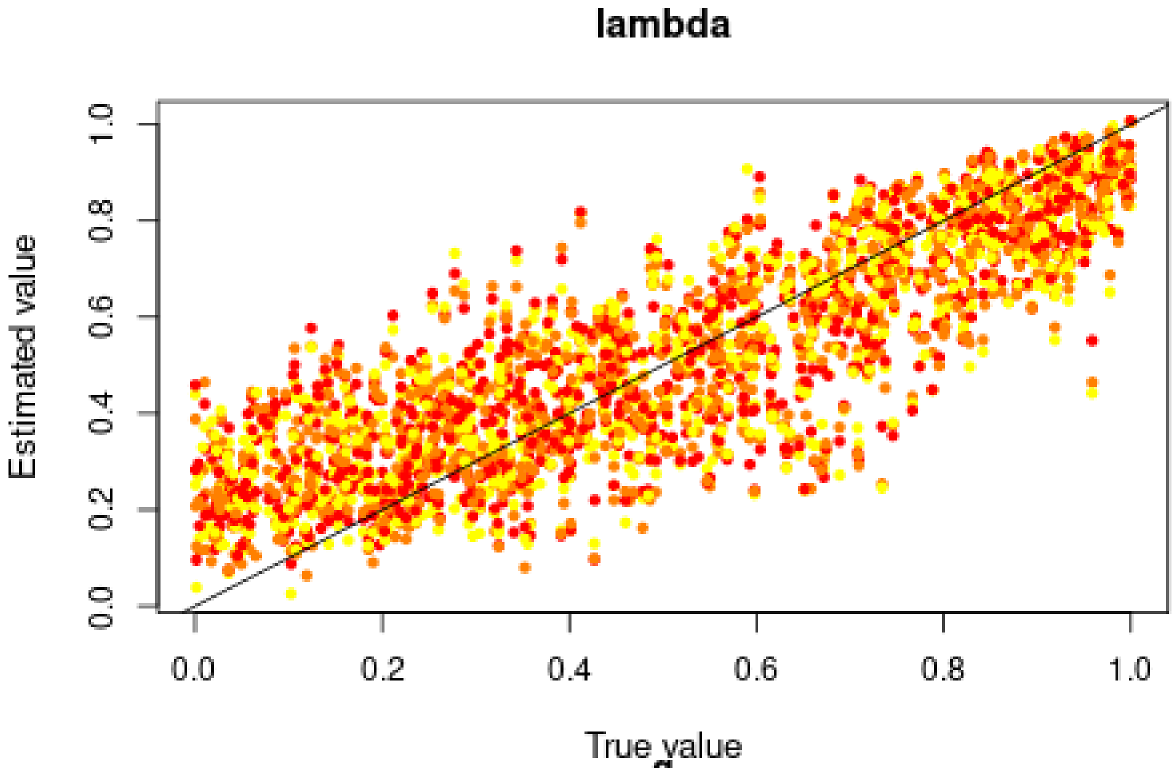

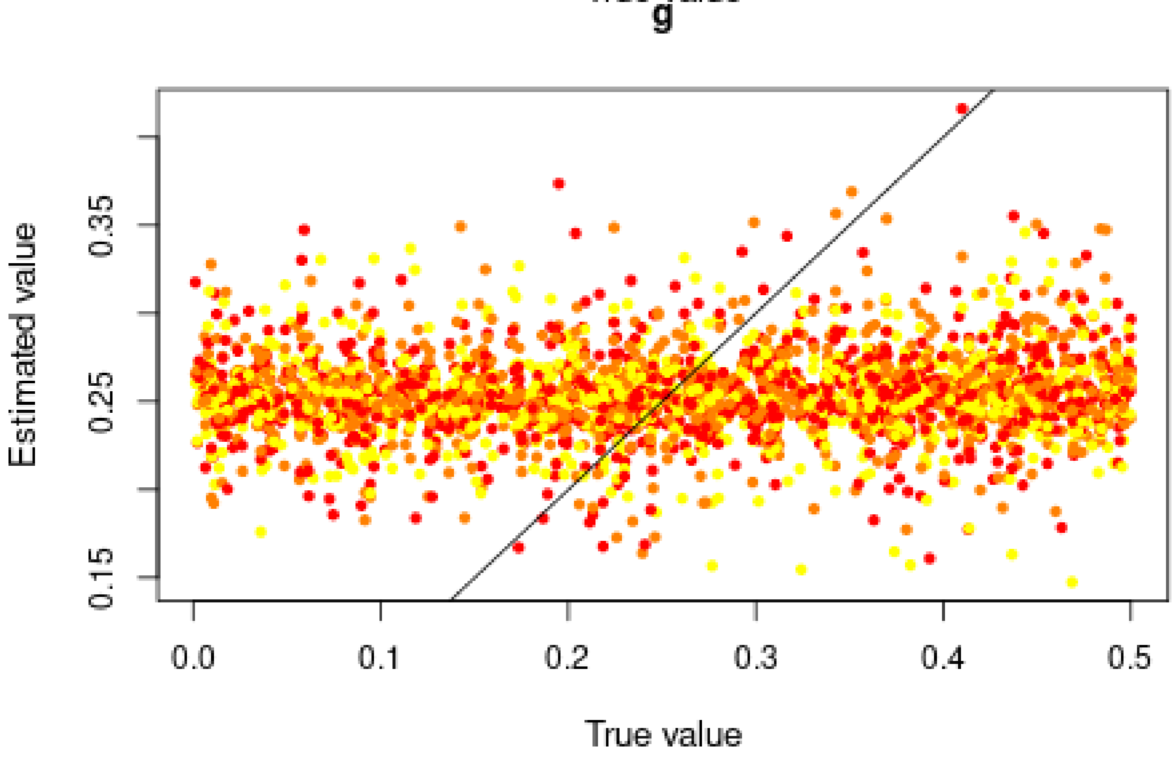

Good for $s$, ok for *lambda*, but very bad for $h$ and $g$, in agreement with the posterior distributions.

##### 3.2 Misclassification errors

To test the power of the approach to distinguish between models, we also conduct a cross-validation experiment. We compare model M0 ($s = 0$) and M1 ($s = 0.6409 > 0$). $h$ and $g$ are sampled over their prior distributions, and *lambda* is set to 0.3895. 100,000 simulations were conducted under each model.

```r
sims0 <- read.csv(paste0("Simulations/",
    "Simulations0NonRandomMatingWithNonGeneticVarianceFromGen3.csv.gz"))
param.sim0 <- subset(sims0, select = c("hm", "sm", "lambda"))
stat.sim0 <- subset(sims0, select = 2:46)
# Only consider the WT homozygotes and the heterozygotes,
# as the sum of the three genotypes is 1
stat.sim0 <- subset(stat.sim0, select = grep(".22.", names(stat.sim0), invert = TRUE))

sims1 <- read.csv(paste0("Simulations/",
    "Simulations1bNonRandomMatingWithNonGeneticVarianceMalesOnlyFromGen3.csv.gz"))
param.sim1 <- subset(sims1, select = c("hm", "sm", "lambda"))
stat.sim1 <- subset(sims1, select = 2:46)
# Only consider the WT homozygotes and the heterozygotes,
# as the sum of the three genotypes is 1
stat.sim1 <- subset(stat.sim1, select = grep(".22.", names(stat.sim1), invert = TRUE))
```

We conduct a CV analysis (this takes some time… set nval = 100 for faster results):

```r
models <- rep(c("Neutral", "Selection"), each = 100000)
if (redo) {
  cv.modsel.g <- cv4postpr(
    models, rbind(stat.sim0, stat.sim1),
    nval = 1000, tols = .1, method = "mnlogistic")
  save(cv.modsel.g,
      file = "Rdata/MalesOnlyNonRandomMatingNonGeneticVariance/backup_cv4postpr_genotype.Rdata")
} else {
  load("Rdata/MalesOnlyNonRandomMatingNonGeneticVariance/backup_cv4postpr_genotype.Rdata")
}
```

We display the results:

```r
summary(cv.modsel.g)
```

```
## Confusion matrix based on 1000 samples for each model.
##
## $tol0.1
##           Neutral Selection
## Neutral       903        97
## Selection      94       906
##
##
## Mean model posterior probabilities (mnlogistic)
##
## $tol0.1
##           Neutral Selection
## Neutral    0.8636    0.1364
## Selection  0.1346    0.8654
```

```r
plot(cv.modsel.g, names.arg=c("Neutral", "Selection"))
```

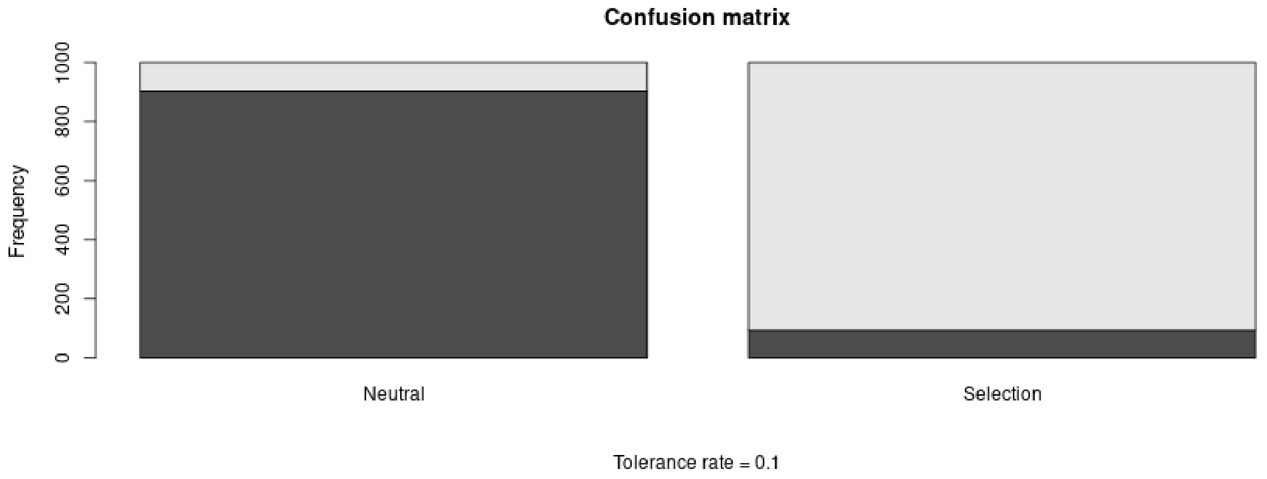

#### 3.3 Posterior prediction

```
if (redo) {
  modsel.g <- postpr(
    stat.obs, models,
    rbind(stat.sim0, stat.sim1),
    tol = .1, method = "mnlogistic")
  save(modsel.g,
       file = "Rdata/MalesOnlyNonRandomMatingNonGeneticVariance/backup_postpr_genotype.Rdata")
} else {
  load("Rdata/MalesOnlyNonRandomMatingNonGeneticVariance/backup_postpr_genotype.Rdata")
}
summary(modsel.g)
```

```
## Call:
## postpr(target = stat.obs, index = models, sumstat = rbind(stat.sim0,
##     stat.sim1), tol = 0.1, method = "mnlogistic")
## Data:
##  postpr.out$values (20000 posterior samples)
## Models a priori:
##  Neutral, Selection
## Models a posteriori:
##  Neutral, Selection
##
## Proportion of accepted simulations (rejection):
##   Neutral Selection
##    0.5972    0.4028
##
## Bayes factors:
##             Neutral Selection
## Neutral      1.0000    1.4826
## Selection    0.6745    1.0000
##
##
## Posterior model probabilities (mnlogistic):
##   Neutral Selection
##    0.0661    0.9339
##
```

```
## Bayes factors:
##           Neutral Selection
## Neutral    1.0000    0.0708
## Selection 14.1203    1.0000
```

The model with selection is preferred.

#### 3.4 Goodness of fit

Under the neutral model:

```
if (redo) {
  res.gfit0.g <- gfit(
    target = stat.obs, sumstat = stat.sim0,
    statistic = median, nb.replicate = 1000)
  save(res.gfit0.g,
       file = "Rdata/MalesOnlyNonRandomMatingNonGeneticVariance/backup_gfit0_genotype.Rdata")
} else {
  load("Rdata/MalesOnlyNonRandomMatingNonGeneticVariance/backup_gfit0_genotype.Rdata")
}
plot(res.gfit0.g, main = "Histogram under M0")
```

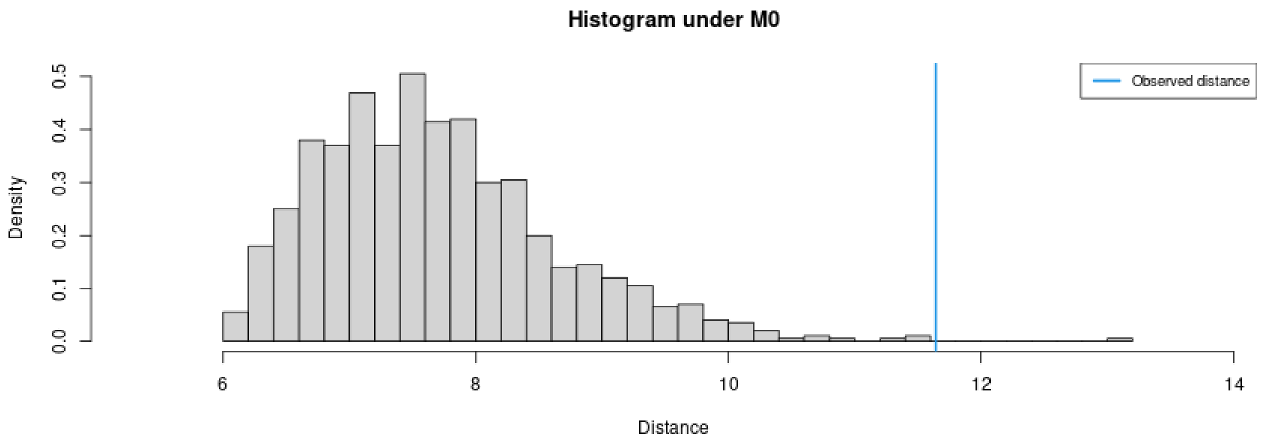

```
summary(res.gfit0.g)
```

```
## $pvalue
## [1] 0.001
##
## $s.dist.sim
##    Min. 1st Qu.  Median    Mean 3rd Qu.    Max.
##   6.009   7.008   7.571   7.696   8.232  13.127
##
## $dist.obs
## [1] 11.64144
```

Under the selection model:

```
if (redo) {
  res.gfit1.g <- gfit(
    target = stat.obs, sumstat = stat.sim1,
    statistic = median, nb.replicate = 1000)
  save(res.gfit1.g,
       file = "Rdata/MalesOnlyNonRandomMatingNonGeneticVariance/backup_gfit1_genotype.Rdata")
} else {
```

```r
  load("Rdata/MalesOnlyNonRandomMatingNonGeneticVariance/backup_gfit1_genotype.Rdata")
}
plot(res.gfit1.g, main = "Histogram under M1")
```

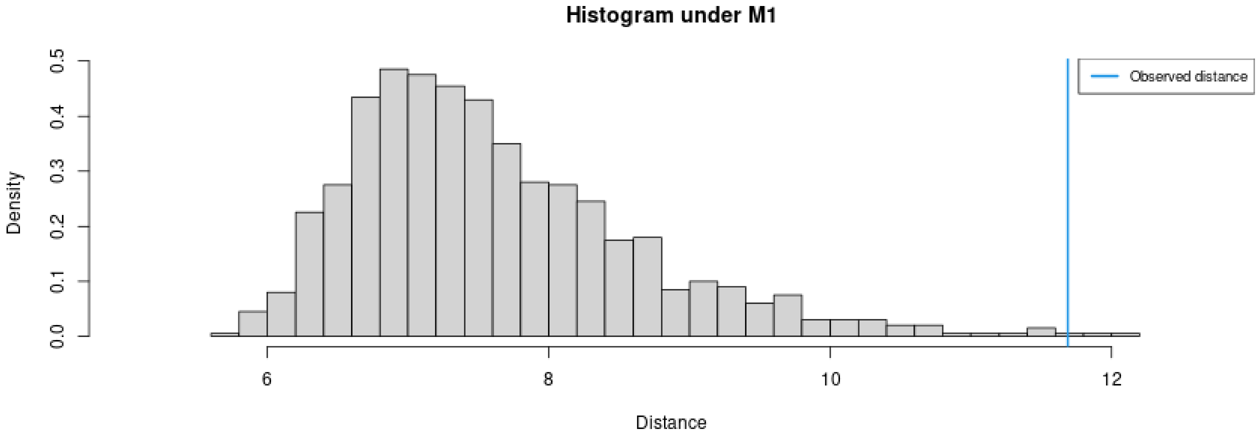

```r
summary(res.gfit1.g)
```

```
## $pvalue
## [1] 0.003
##
## $s.dist.sim
##    Min. 1st Qu.  Median    Mean 3rd Qu.    Max.
##   5.680   6.882   7.410   7.607   8.147  12.070
##
## $dist.obs
## [1] 11.68802
```

```
## No id variables; using all as measure variables
```

```r
d1$Tolerance <- rep(c(0.01, 0.1, 0.2), each = 1000)
d1$Replicate <- rep(1:1000, 3)
d2<-melt(d2, value.name = "True")
```

```
## No id variables; using all as measure variables
```

```r
d2$Replicate <- rep(1:1000, 3)
dat.cv <- merge(d1, d2, by = c("variable", "Replicate"))
dat.cv$variable <- factor(dat.cv$variable,
                          levels = c("hm", "sm", "g", "lambda"),
                          labels = c("h", "s", "g", "lambda"))
```

```r
p.cv <- ggplot(dat.cv, aes(x = True, y = Estimated)) +
        geom_point(aes(col = as.ordered(Tolerance))) +
        geom_abline(slope = 1) +
        facet_wrap(~variable, scales = "free") +
        theme_pubclean() + labs(color = "Tolerance") +
        ggtitle("Cross validation") +
        theme(strip.background = element_blank())
```

Confusion matrix:

```r
library(scales)
cv.sum <- summary(cv.modsel.g)
```

```
## Confusion matrix based on 1000 samples for each model.
##
```

```
## $tol0.1
##           Neutral Selection
## Neutral       903        97
## Selection      94       906
##
##
## Mean model posterior probabilities (mnlogistic)
##
## $tol0.1
##           Neutral Selection
## Neutral    0.8636    0.1364
## Selection  0.1346    0.8654
```

```
dat.cv <- as.data.frame(cv.sum$conf.matrix$tol0.1/1000)
names(dat.cv) <- c("Real", "Inferred", "Frequency")
p.confmat <- ggplot(dat.cv, aes(x = Real, y = Frequency, fill = Inferred)) +
             geom_col() +
             scale_y_continuous(labels = scales::percent) +
             scale_fill_brewer(type = "qual", palette = 3) +
             ggtitle("Prediction errors") +
             theme_pubclean()
```

Model probabilities:

```
p.mprob <- ggplot(as.data.frame(modsel.g$pred), aes(x = Var1, y = Freq, fill = Var1)) +
           geom_col() + ylab("Model posterior probability") + xlab("Model") +
           scale_fill_brewer(type = "qual", palette = 3) +
           theme_pubclean() +
           ggtitle("Model probabilities") +
           theme(legend.position = "none")
```

```
library(cowplot)
```

```
##
## Attaching package: 'cowplot'

## The following object is masked from 'package:ggpubr':
##
##     get_legend
```

```
p1 <- plot_grid(p.mprob, p.confmat, labels = c("C", "D"), nrow = 1)
p <- plot_grid(p.dist, p.cv, p1, labels = c("A", "B", ""), nrow = 3)
p
```

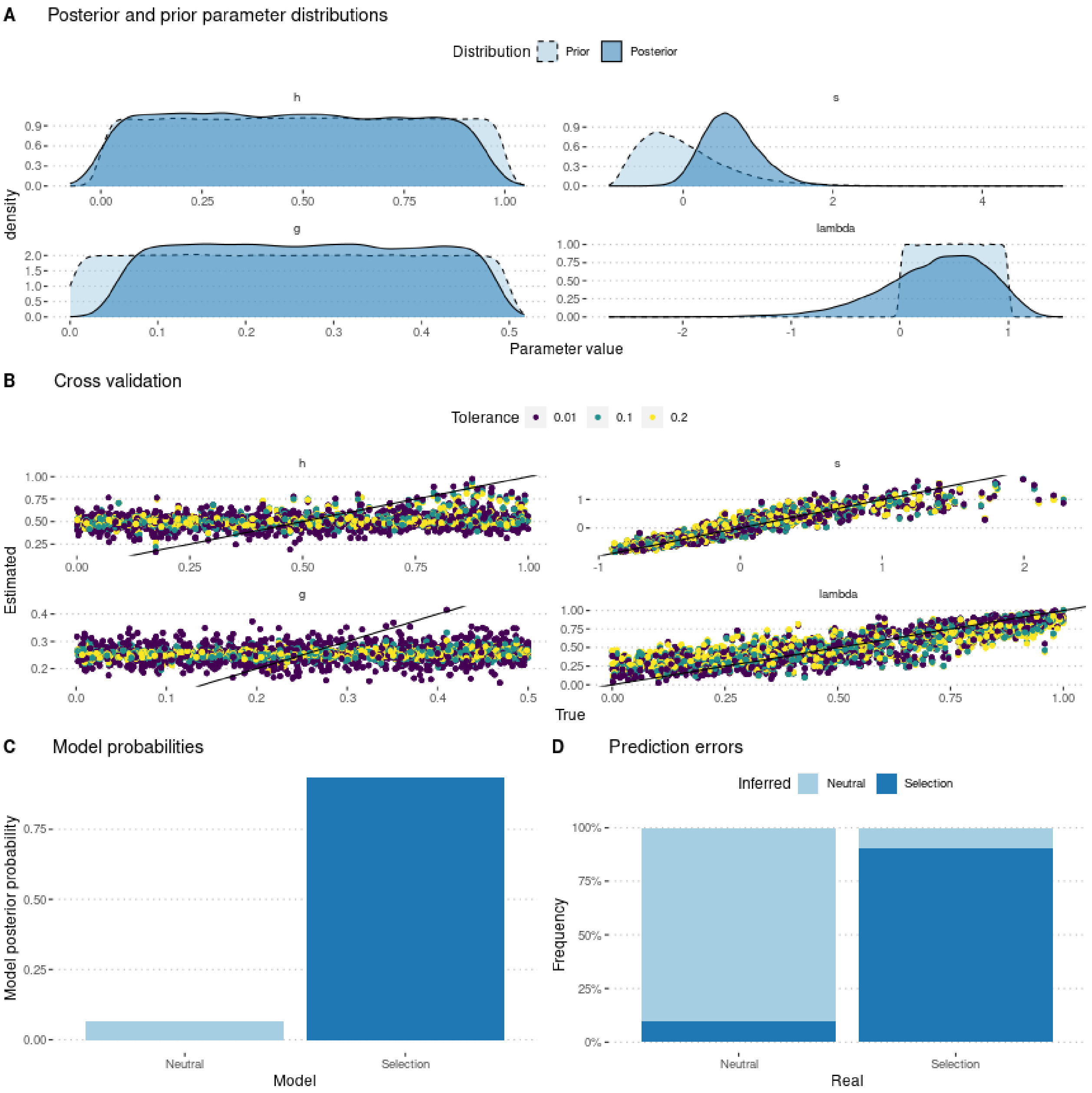

```r
ggsave(p,
       filename = "FigureABC-MalesOnlyNonRandomMatingNonGeneticVariance-Genotype.pdf",
       width = 8, height = 10)
```
