## Supplementary material for "Experimental evaluation of a direct fitness effect of the *de novo* evolved mouse gene *Pldi*": combined suppl files: SupplementaryFile10.pdf

Sequence alignments of the genomic regions coding for the *Pldi* RNA exons in *Mus* species and subspecies.

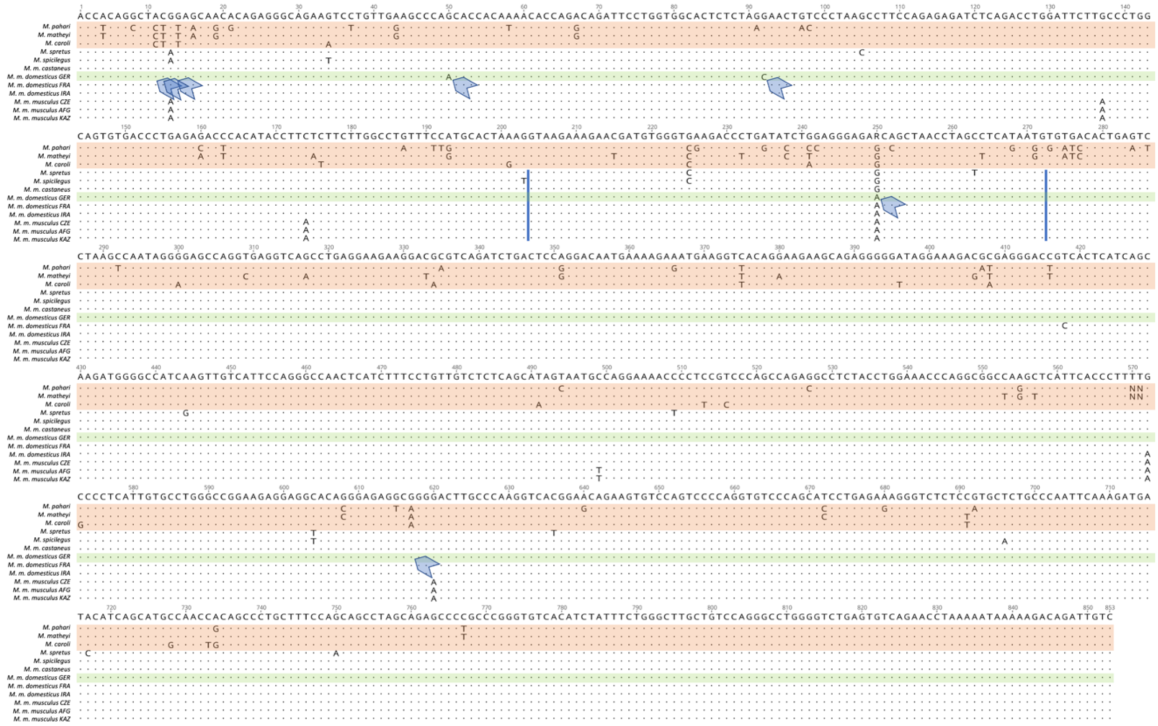

Outgroup species that do not express the *Pldi* RNA include *M. pahari*, *M. matheyi*, and *M. caroli* (shaded in red). The ingroup species are represented by population consensus sequences (obtained from the data described in (Harr et al., 2016) of *M. spretus*, *M. spicilegus*, as well as the subspecies *M. m. castaneus*, *M. m. domesticus*, and *M. m. musculus*. The latter two are represented by three populations each, labeled with 3-letter codes. The *M. m. domesticus* GER (shaded in green) corresponds to the *Mus musculus* reference sequence mm10, which is also the C57Bl6/J sequence that served here as the WT strain. Blue vertical lines indicate the positions of the introns. Blue arrows indicate substitutions in the *M. m. domesticus* GER sequence compared to the outgroups.
