## Supplementary figures and images for "Experimental evaluation of a direct fitness effect of the *de novo* evolved mouse gene *Pldi*"

### SupplementaryFile9.pdf

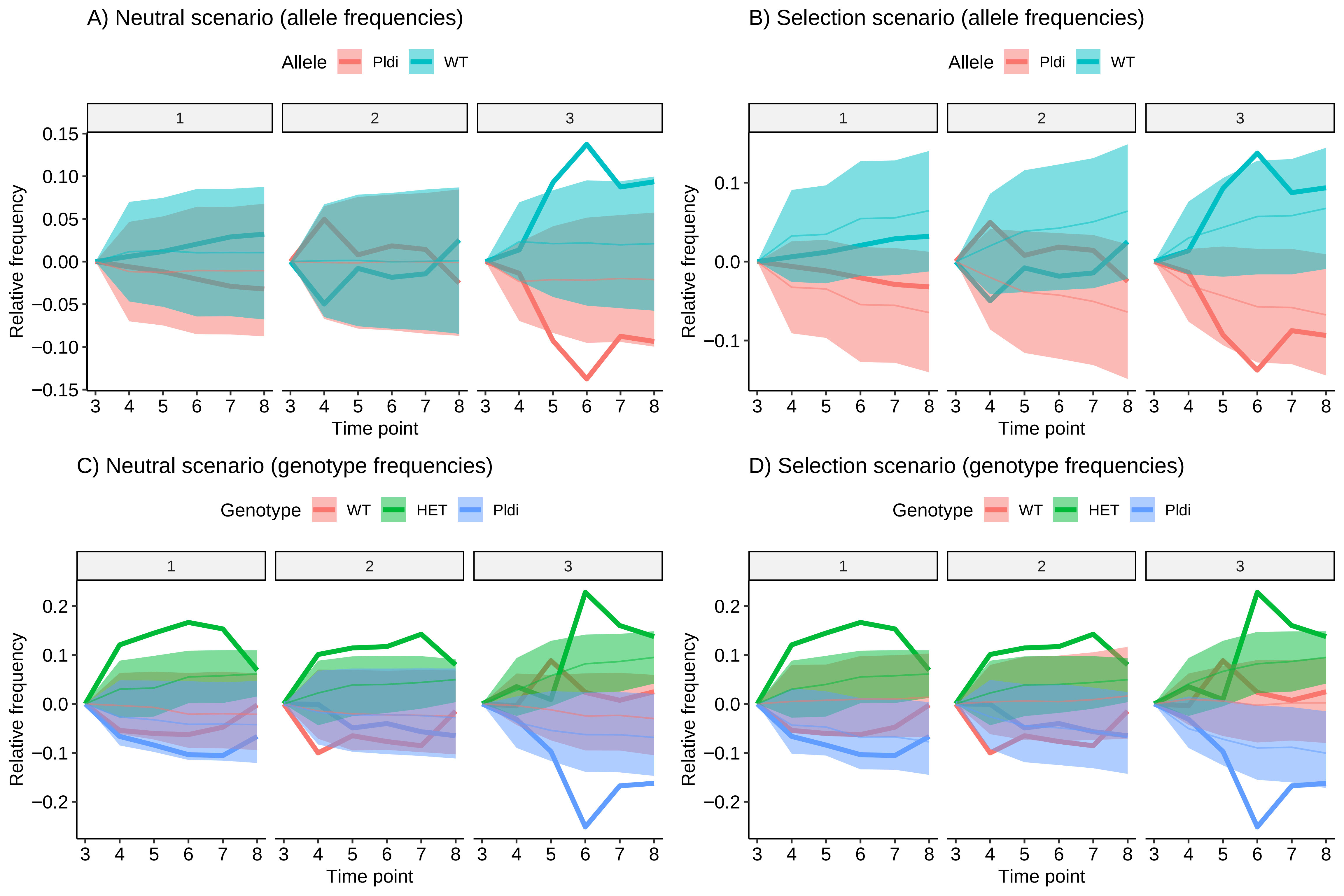
